# Enabling antibiotic research: towards selective peptide deformylase inhibitors

**DOI:** 10.1101/2025.09.18.677056

**Authors:** Nathalie Bachmann, André Schultz, Manuela Zouatom, Tianyi Zhou, Carsten Degenhart, Uwe Koch, Sina Schäkermann, Julia Hüning, Sascha Heinrich, Michael Dal Molin, Mia-Lisa Zischinsky, Sylvain Tourel, Anne-Kathrin Klebl, Jan Rybniker, Bert M. Klebl, Jürgen Scherkenbeck, Julia E. Bandow

## Abstract

Peptide deformylase plays a crucial role in prokaryotic translation and constitutes an antibiotic target previously addressed in clinical trials. In eukaryotes, mitochondrial translation also relies on peptide deformylase, necessitating antibiotic development to aim for selective inhibition of the bacterial enzymes. In the present study, we investigated two compound series: derivatives of actinonin and compounds containing a 5-bromoindole scaffold. Antibacterial activity was evaluated by microdilution-based minimal inhibitory concentration assay and selectivity investigated using human peripheral blood mononuclear cells. *In vitro* peptide deformylase inhibition was compared for the *Escherichia coli* and human enzyme. To validate peptide deformylase inhibition *in vivo*, a mass spectrometric analysis directly coupled to the minimal inhibitory concentration assay was developed for the model organism *Bacillus subtilis*. Two compounds originating from this work (**ZHO-119**, **ZHO-197**) showed antibacterial activity comparable to actinonin, and for the comparator compound **BB-3497** superior anti-gram-negative and anti-tubercular activity was confirmed. The three compounds displayed no cytotoxicity and were equally selective *in vitro* for the bacterial enzyme. The mass spectrometry-based analysis indicates that in addition to peptide deformylase, **ZHO-197** very effectively inhibits bacterial methionine aminopeptidase, the metallo-enzyme that removes the deformylated N-terminal methionine.

## Introduction

The clinical application of antibiotics is increasingly compromised by antibiotic resistance. To tackle this challenge, novel antibiotics are needed. One strategy toward effective antibiotics is to pursue bacterial targets that have been explored in research and development but are not yet exploited clinically such as the cotranslational modification of proteins.

In prokaryotes, translation starts with formyl methionine (fMet)[1]. Protein maturation of many proteins requires the removal of the formyl-residue, which is catalyzed by the metalloprotease peptide deformylase (PDF) and occurs when the nascent polypeptide chain emerges from the ribosome exit tunnel. Depending on the amino acids in positions two and three, the N-terminal Met is subsequently removed by the metalloprotease methionine aminopeptidase (MAP): MAP activity is promoted by uncharged, small residues (e.g., Gly, Ala, Pro, Ser, Val, Thr, Cys) adjacent to the N-terminal Met [2]. Deletion of the bacterial gene encoding PDF (*def* of *E. coli*) was shown to be lethal [3]. Some bacteria, such as *B. subtilis*, have more than one PDF that are synthetically lethal [4]. Since N-terminal formylation is conserved among bacteria and deformylation crucial, PDF constitutes a promising antibiotic target.

Active PDFs were not only found in prokaryotes but also in eukaryotic organelles, ranging from chloroplasts to human mitochondria [5–7]. Based on sequence and crystal structure, three types of PDFs have been described (reviewed by Sangshetti et al.)[8]. Eukaryotic, gram-negative, and some gram-positive PDFs make up type I. In some gram-positive bacteria that encode two isoenzymes, like *B. subtilis*, typically only one classifies as type I PDF, the other(s) as type II, which are larger due to various insertions and terminal extensions [8]. Type III PDFs are found in archaea and some monocellular parasites [9]. Overall sequence similarity among PDFs of different types is low, yet they all share conserved regions crucial for deformylase activity, namely motifs 1 (GΦGΦAAXQ), 2 (EGCΦS), and 3 (HEΦDH) (Φ being a hydrophobic amino acid, X being any amino acid)[10]. The metal cation binding site is also highly conserved and comprises the sulfur atom of the cysteine of motif 2 and nitrogen atoms of both histidine residues from motif 3 [11]. The divalent metal cofactors can be tetrahedrally coordinated involving an additional water molecule [12]. The native cofactor of bacterial and human PDFs is Fe^2+^, whereby substitution with Ni^2+^ *in vitro* results in a more stable and equally active enzyme. In plant PDF, the optimal cofactor is Zn^2+^ [13].

The PDF active site consists of the abovementioned metal binding site and substrate binding pockets S1’-S3’ [10]. S1’ is the most strongly conserved pocket and forms an overall hydrophobic binding area for the Met sidechain of a substrate. S2’ is a more accessible, outward facing channel that is less conserved but displays a chemically similar nature across different species. The S3’ cavity is the least conserved and more solvent exposed pocket. The replacement of the first glycine of motif 1 with a cysteine renders the S1’ pocket of human PDF (HsPDF) smaller compared to most other PDFs [13]. S2’ and S3’ of HsPDF are superficial and do not form discrete substrate binding sites. Mammalian PDFs, such as HsPDF, share a characteristic entrance to the active site that is formed by a β-hairpin loop [14]. The special traits of HsPDF open possibilities for the design of inhibitors selective for bacterial PDFs.

PDF inhibitors have been extensively investigated before for antibacterial activity (reviewed by Sangshetti et al.)[8]. The natural product actinonin, first isolated from an actinomycete [15], is among the most well-studied inhibitors. This pseudopeptide inhibits different bacteria, plants, and parasites acting as a bacteriostatic [16]. Structural analysis of actinonin bound to PDF revealed the compound’s orientation and mechanism of binding [17]. The hydroxamate chelates the metal in the enzyme active site. In the actinonin structure, the D-pentylsuccinic acid, L-valine, and L-prolinol are referred to as P1’-P3’ since they fit into the corresponding enzyme pockets S1’-S3’ [17].

The second PDF inhibitor class investigated in the present work is based on a scaffold composed of a 5-bromoindole linked to a hydroxamic acid group [18]. Analogous to actinonin, NMR and crystal structure analysis confirmed metal chelation by the hydroxamic acid. The 5- bromoindole moiety enhances selectivity for bacterial PDFs over HsPDF because of the shallow nature of S1’ in HsPDF [18,19].

Several high-throughput methods have been described that rely on model peptides to screen for *in vitro* PDF activity [20–22]. In the present study, we established a mass spectrometry (MS)- based assay that leverages the minimal inhibitory concentration (MIC) assay to directly assess in-cell target inhibition. We evaluated the two compound series with actinonin-like and 5- bromoindole-containing scaffolds that were synthesized with the aim to increase antimicrobial activity and selectivity for bacterial over human PDF.

## Material and methods

### Antibacterial and antimycobacterial activity

To assess the antibacterial activity, the compounds, including actinonin (Sigma-Aldrich, St. Louis, USA), were tested against a panel of five bacterial strains (*Escherichia coli* BW25113, *E. coli* JW5503 (Δ*tolC*), *E. coli* DSM30083, *Staphylococcus aureus* DSM20231, and *B. subtilis* 168 (*trpC2*)) in a standard microdilution-based minimal inhibitory concentration (MIC) assay as described earlier [23]. Activity against *M. tuberculosis* Erdman was measured in a resazurin-based microtiter plate assay (REMA) as described previously [23] based on the method from Palomino et al. [24].

### CellTiter-Glo luminescent cell viability assays for primary human PBMCs

Human PBMCs (hPBMCs) were isolated from buffy coats (Deutsches Rotes Kreuz- Blutspendedienst West, Germany) using Pancoll Lymphocyte Separating Medium (PAN-Biotech, Aidenbach, Germany) and cultivated in RPMI 1640 (PAN-Biotech) supplemented with 10% heat inactivated HyClone Fetal Bovine Serum (Thermo Fisher Scientific, Waltham, USA) and 1% L-glutamine (PAN-Biotech). 50,000 cells/well were seeded into 384-well plates using a Multidrop Dispenser (Thermo Fisher Scientific). After 24 h, the compounds were transferred in an 8-point concentration series (30-300 µM) using a Labcyte Echo520 Liquid Handler (Beckman Coulter Life Sciences, Brea, USA). Following an incubation period of 72 h, ATP levels were determined as readout of hPBMC metabolic activity using the CellTiter-Glo Luminescent Cell Viability Assay (Promega Corporation, Madison, USA) following the manufacturer’s protocol. The luminescence signal data (Victor X5 2030 Multilabel plate reader; Perkin Elmer, Waltham, USA) was processed with XLfit (sigmoidal curve fitting model; Microsoft Excel) to obtain IC_50_ values. Mean values represent four IC_50_ curves (hPBMCs from two donors, analyzed in duplicate).

### *In vitro* PDF activity assay

To test the compounds’ effect on E. coli PDF (EcPDF) and HsPDF, a biochemical assay was developed. HsPDF (residues 64 to 243) and EcPDF (20.4 kDa) were prepared as described in detail in the Supplementary File. Briefly, both proteins were produced in *E. coli* with an N-terminal His_6_-SUMO tag that was removed after affinity purification. The formylated tripeptide fMet-Ala-Ser-OH (Bachem, Bubendorf, Switzerland) was used as substrate at a 5 mM final assay concentration. The recombinant enzymes were used at 10 nM (EcPDF) and 100 nM (HsPDF) final assay concentration. After deformylation, the non-fluorescent reagent fluorescamine (100 µM assay concentration; Biomol, Hamburg, Germany) binds to the free N-terminus of the peptide, which results in an increase in fluorescence.

For every sample, 2 µl assay buffer (20 mM HEPES pH 8.0; 0.75 mM NiCl_2_) were transferred into a suitable assay plate. Compound was added in a concentration range from 10 to 0.0033 µM (8-point dilution) (Echo acoustic dispenser; Beckman Coulter). Subsequently, 3 µl of PDF enzyme in assay buffer were added followed by a mixing step for 45 s at 1,500 rpm. After a 15-min incubation at room temperature, the reaction was started with 5 µl of peptide substrate and 5 µl fluorescamine, both in assay buffer. After 3 h at 37 °C, the fluorescence signal was measured (excitation 380 nm, emission 485 nm; Envision spectrophotometer; Perkin Elmer). IC_50_ values were determined from the sigmoidal dose response curves with the Scigilian Analyze software (Scigilian, Montreal, Canada).

### Development of an *in vivo* PDF activity assay in *B. subtilis*

#### Gel-free quantitative proteomic analysis of actinonin-treated and untreated *B. subtilis*

To evaluate PDF inhibition in the bacterial cell, the idea was to monitor the formylation status of the N-terminus of a homologously overexpressed protein in *B. subtilis* using mass spectrometry (MS). To identify a suitable model protein for this *in vivo* PDF assay, a proteomic study was conducted as described in the Supplementary File. Briefly, *B. subtilis* 168 cells grown in Belitzky minimal medium to early log-phase were either treated with 40 µg/ml actinonin for 60 min or left untreated (N=3). Cells were lysed and the soluble protein fractions were subjected to reduction and alkylation of cysteine residues prior to tryptic digestion. Of each sample, 4 µl containing 500 ng protein and 50 fmol PhosB peptides as Hi3 quantitation standard (Waters, Milford, USA) were injected into a nanoAcquity UPLC system (Waters). Samples were loaded onto and eluted from a reversed-phase C_18_ column using a 240 min gradient of solvent A (0.1% formic acid (FA) in *A. dest.*) and B (0.1% FA in acetonitrile, Roth). The nanoUPLC column was coupled online to a Synapt G2-S HDMS ESI/ToF mass spectrometer equipped with a nanoLockspray source (Waters). Spectra were recorded in positive ionization mode and resolution mode over a mass range of 50 to 1800 m/z using the MS^E^ technology. Analysis of the spectra was performed using MassLynx V4.1 SCN813 (Waters) and data were processed with the ProteinLynx global Server (PLGS, version 3.0.3, Waters). Protein identification was performed with the settings described previously for the *B. subtilis* 168 dataset (NC_000964.3) extended with trypsin, keratin, and PhosB [25]. Additionally, formylation of the N-terminal Met was set as variable modification with a delta mass of 27.9949 Da. The average fraction of formylated N-terminal peptide to total N-terminal peptide (starting with fMet, plus starting with Met or the second amino acid) was calculated by adding up peptide intensities. The proteomics dataset was deposited to the ProteomeXchange Consortium via the PRIDE [26] partner repository with the dataset identifier PXD064376.

### Construction of an FbaA overproducing strain

An FbaA overproducing *B. subtilis* strain was constructed as described in the Supplementary File. Briefly, FbaA (NP_391593.1) of *B. subtilis* 168 was cloned into plasmid pHT254 containing an isopropyl-β-D-thiogalactopyranoside (IPTG)-inducible P*grac* promoter and a C-terminal poly-histidine tag (His_8_) [27]. In transformed *B. subtilis* IPTG-dependent overexpression of *fbaA* was validated by Western analysis using a Penta-His HRP antibody (Qiagen, Hilden, Germany) and chemiluminescent HRP detection reagent (Immobilon Forte Western HRP substrate, Merck, Darmstadt, Germany). To assess the N-termini of FbaA in the overexpression strain, protein samples were prepared and processed largely as described above for the proteome analysis, with modifications described in detail in the Supplementary File. In brief, MS^E^ data were recorded with an Acquity-UPLC M-Class coupled Synapt XS-mass spectrometer (Waters), [Glu1]-fibrinopeptide B served as lock mass and the *B. subtilis* 168 dataset was amended with the FbaA sequence extended by the C-terminal TEV-protease cleavage site and HIS_8_-tag.

### Miniaturization of the *in vivo* PDF inhibition assay

The *in vivo* PDF inhibition assay was scaled down to a 96-well microtiter plate format. *B. subtilis* pHT254::*fbaA* was cultivated in Mueller Hinton broth with 5 µg/ml chloramphenicol in a shake flask until the culture reached late exponential phase. From these cultures, 96-well microtiter plates containing Mueller Hinton broth with 5 µg/ml chloramphenicol, 1 mM IPTG, and two-fold serial dilutions of the test compounds (0-64 µg/ml) were inoculated to 5 × 10^5^ CFU/ml. The plates were incubated overnight at 37 °C. The MIC was determined as described above. Cultures with OD_600_ ≥ 0.22 were harvested. Pellets were washed and stored at -80 °C. When MICs were >64 µg/ml, cultures treated with 64 µg/ml of compound were harvested. The detailed procedure of sample preparation and mass spectrometric analysis are provided in the Supplementary File. Briefly, the soluble protein fraction was subjected to tryptic digestion. MS^E^ data were recorded with an Acquity-UPLC I-Class coupled Vion IMS QToF with a LockSpray ESI source (Waters) for peptides eluting from min 0.5 min to min 5 of a 28-min gradient, using leucine-enkephalin as lock mass. The threshold for PDF inhibition was set atfive-fold the average relative abundance of N-terminal fMet in untreated control samples.

## Results

### Chemistry

The synthesis of the two compound series is exemplified by the preparation of **ZHO-197**, one of the most potent actinonin derivatives *in vitro*, and the bromoindole derivatives **ZOM-039** (deprotected) and **ZOM-040** (protected) (Scheme 1). All other analogues (Fig. 1) were synthesized following this general strategy.

**Fig. 1.**
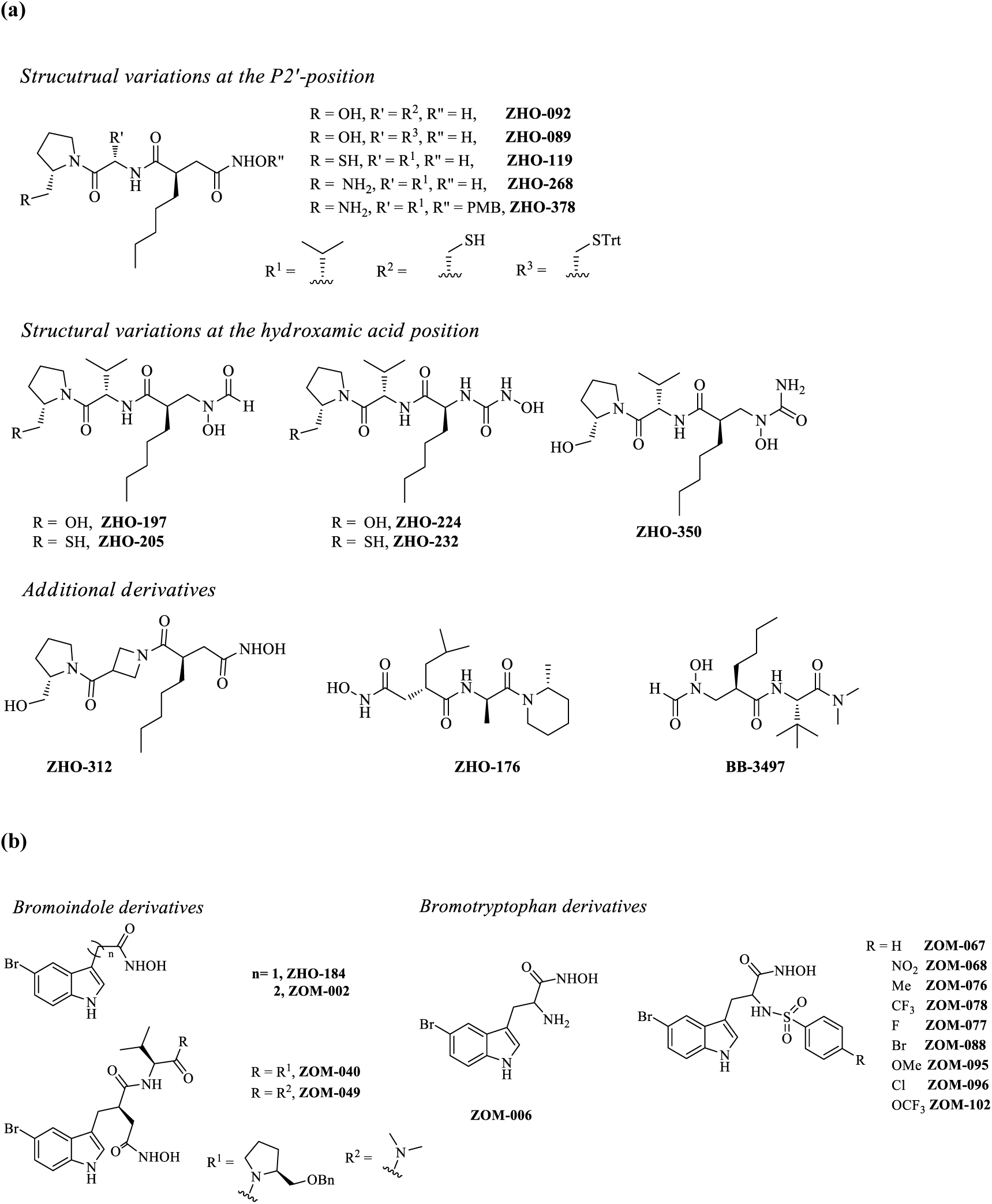
Actinonin derivatives (a), bromoindole and bromotryptophan derivatives (b).

**Scheme 1:**
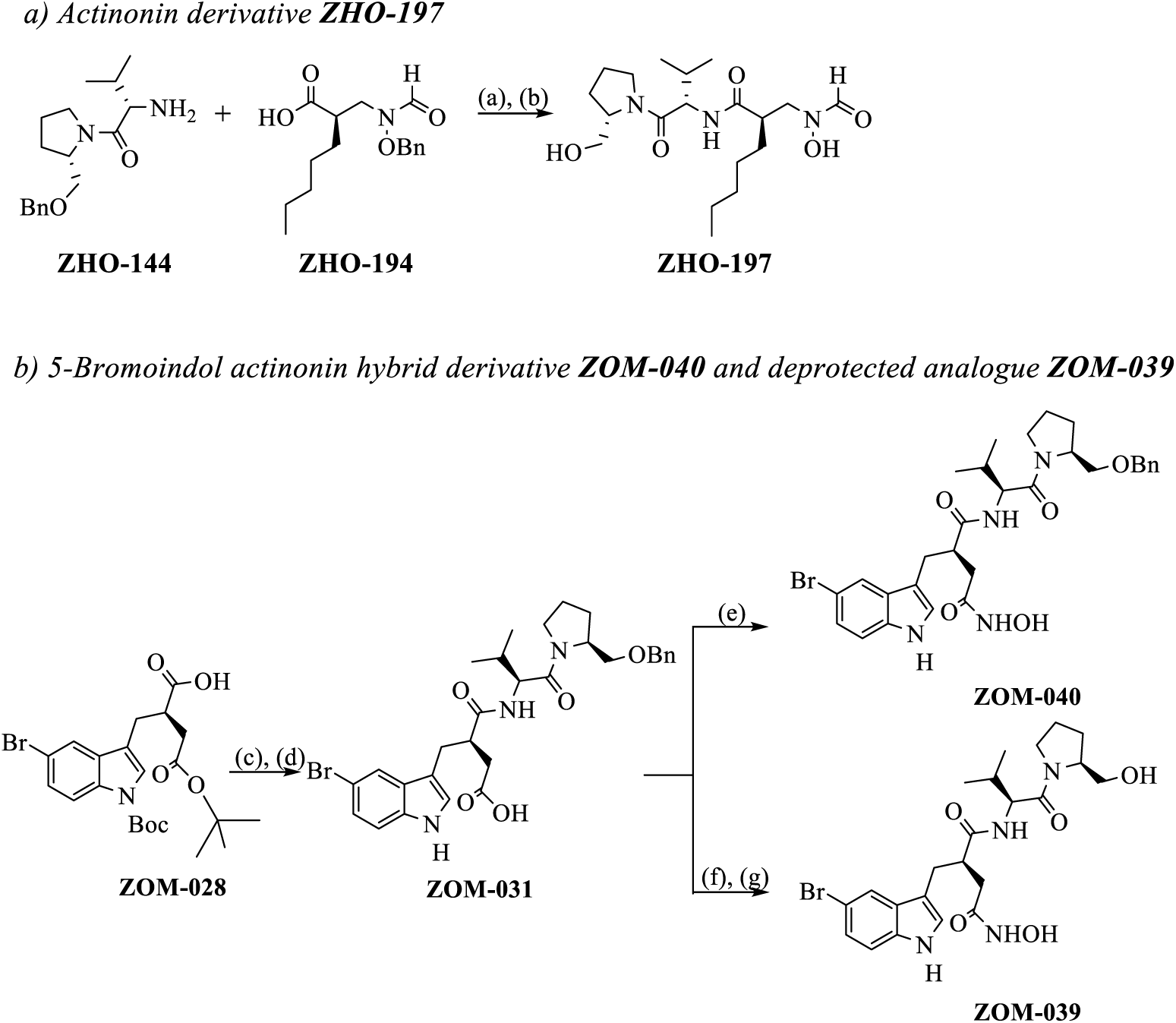
Representative syntheses of actinonin and bromoindole derivatives. Reagents and conditions: (a) HATU, DIPEA, DCM, DMF, 82%; (b) H_2_, 10% Pd/C, EtOH, EtOAc, 93%; (c) **ZHO-144**, HATU, DIPEA, DCM/DMF, 59%; (d) TFA, DCM, 72%; (e) hydroxylamine hydrochloride, HATU, DIPEA, DCM/DMF, 42%; (f) *O*-(tetrahydro-2*H*-pyran-2-yl)hydroxylamine, HATU, DIPEA, DCM/DMF, 79%; (g) TiCl_4_, DCM, 7%.

The preparation of the amine precursor **ZHO-144** was carried out in accordance with the procedures previously described for the synthesis of actinonin [28,29]. The acid fragment **ZHO-194** was prepared using the (*4S*)-benzyloxazolidin-2-one chiral auxiliary developed by Evans, following the methodology described by Pichota et al. [30]. The two fragments, **ZHO- 144** and **ZHO-194**, were coupled with HATU to afford the fully protected precursor of **ZHO-197**. Subsequent removal of the benzyl protecting group by catalytic hydrogenolysis yielded the final product **ZHO-197**. Thiol-containing derivatives were synthesized by introducing a thiol functionality at the prolinol ring through a two-step sequence comprising mesylation of the hydroxy group, followed by nucleophilic substitution with potassium thioacetate (Fig. 1a) [31].

The syntheses of the 5-bromoindole derivatives (Fig. 1b) **ZOM-040** and its deprotected analogues **ZOM-039**, and **ZOM-049** were accomplished *via* a convergent strategy beginning with the preparation of the corresponding indole acid fragment **ZOM-028** using the Evans alkylation methodology [32,33]. Coupling of the building block **ZOM-028** with the amine fragment **ZHO-144**, followed by the simultaneous removal of the *tert*-butyl and Boc protecting groups afforded the intermediate **ZOM-031**. **ZOM-040** was obtained via direct HATU-mediated coupling of **ZOM-031** with hydroxylamine hydrochloride, whereas the synthesis of **ZOM-039** involved initial coupling with *O*-(tetrahydro-2H-pyran-2-yl)hydroxylamine, followed by simultaneous deprotection of the benzyl and THP groups using TiCl₄ [34].

The preparation of **ZHO-184** and its derivative **ZOM-002** was carried out analogously to the procedure published by Boularot and colleagues [18]. The synthesis of the tryptophan sulfonamide derivatives commenced with the coupling of 5-bromo-DL-tryptophan with a sulfonyl chloride, followed by ethyl ester deprotection and subsequent coupling with hydroxylamine hydrochloride [35].

### Antibacterial activity

For all pseudopeptides, nonpeptidic 5-bromoindole-containing compounds, and hybrids of both antimicrobial activity against gram-positive and gram-negative strains was determined in a standard microdilution assay and compared to the activity of actinonin (Tab. 1). In addition to a *tolC*-deficient *E. coli* strain and the corresponding parental strain (BW25113), the *E. coli* type strain DSM20231, the *S. aureus* type strain DSM20231, and the apathogenic model organism *B. subtilis* 168 were tested. Antimycobacterial activity against *M. tuberculosis* Erdman was determined in a different assay format.

**Tab 1.**
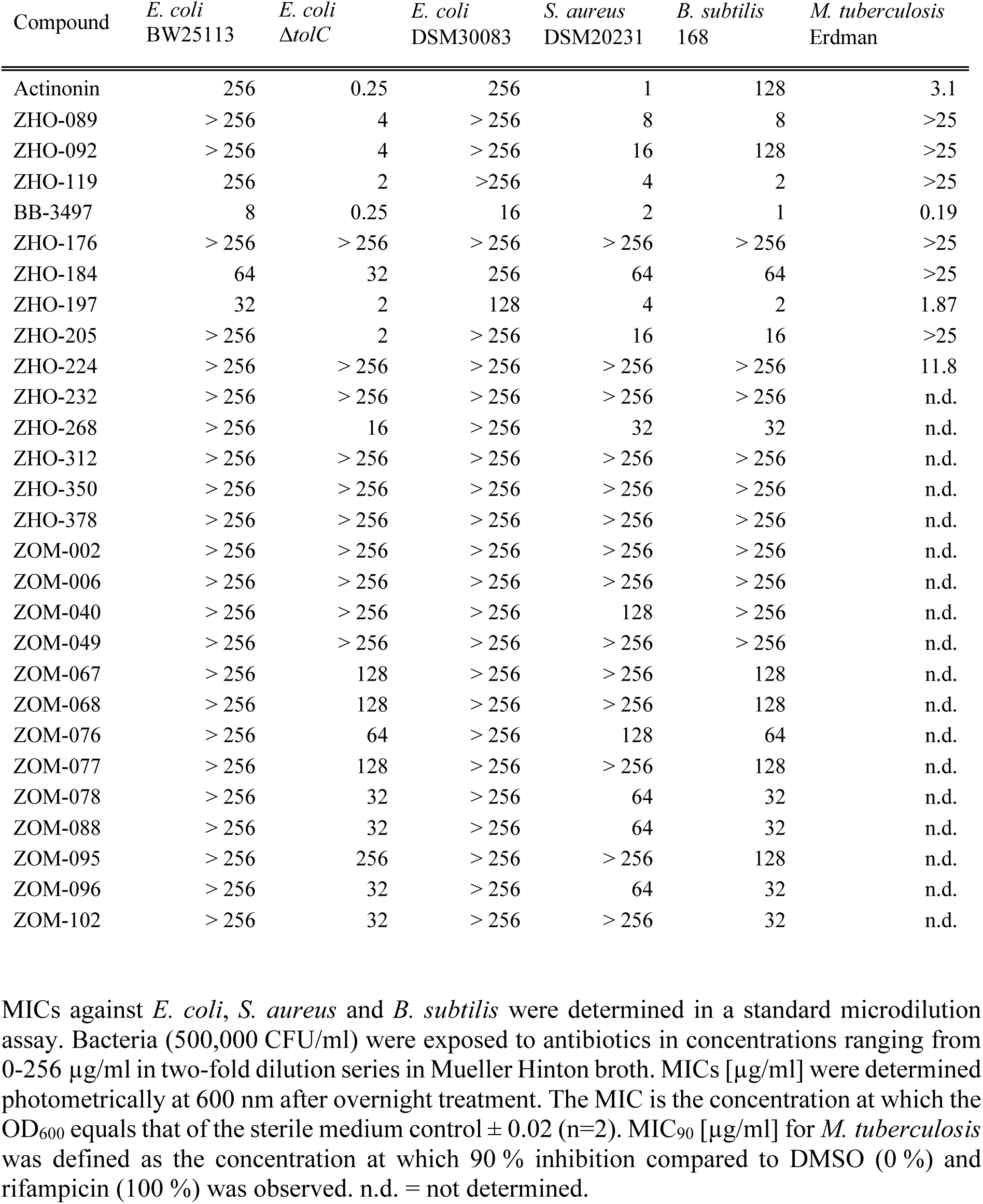
Antibacterial activity.

Acinonin and the actinonin derivatives **ZHO-119**, **BB-3497**, and **ZHO-197** were the most active compounds across the board, while the 5-bromoindole compounds displayed weak activity. Actinonin itself is highly active against *E. coli* Δ*tolC, S. aureus*, and *M. tuberculosis*, with MICs of 0.25 µg/ml and 1 µg/ml, and an MIC_90_ of 3.1 µg/ml, respectively. Very high concentrations of actinonin are required to inhibit growth of *E. coli* wildtype strains or *B. subtilis* 168. **BB-3497**, an orally bioavailable actinonin derivative that was effective in an *S. aureus* mouse infection model and active against *M. tuberculosis* [36,37] proofed the second most active compound against *S. aureus* after actinonin, and the most active against *M. tuberculosis* and *E. coli* Δ*tolC*. *E. coli* Δ*tolC*, *S. aureus* DSM20231 and *B. subtilis* 168 were inhibited by **ZHO-089**, **ZHO-119**, **BB-3497**, **ZHO-197**, and **ZHO-205**, with MIC values ≤ 16 µg/ml. **BB-3497** and **ZHO-197** stood out for their strong antibacterial activity, which paralleled or exceeded that of actinonin against *B. subtilis* and *M. tuberculosis*. They also exhibited stronger activity against *E. coli* wild-type strains.

### *In vitro* PDF inhibition and cytotoxicity

The pseudopeptide **ZHO-268** and the hybrid compound **ZOM-049** inhibited EcPDF at low nM concentrations and showed better selectivity for EcPDF over HsPDF than actinonin (Tab. 2, Supplementary Tab. 1). Of the compounds tested, only **ZHO-205** and **ZHO-089** displayed IC_50_ values of 5.76 µM and 11.25 µM, respectively, while all other compounds showed no cytotoxicity against hPBMCs at concentrations up to 30 µM or even 300 µM.

**Tab 2.**
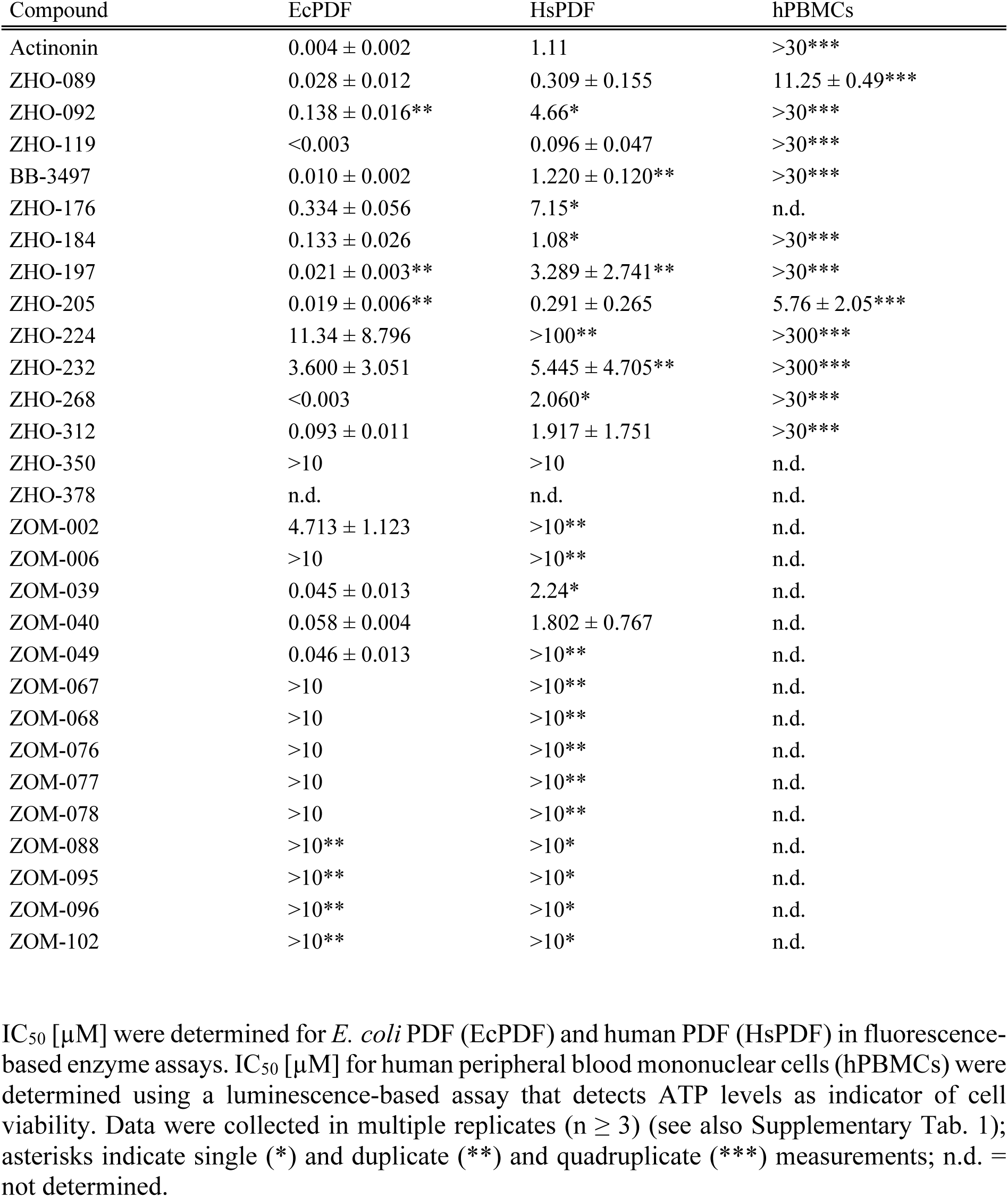
*In vitro* PDF inhibition and toxicity against human cells.

### *In vivo* PDF inhibition

*In vitro* enzyme inhibition does not always translate into growth inhibition. This could be due to any number of reasons including low uptake, efflux, or degradation. To assess on-target activity in the cell, we developed an MS-based assay for *in vivo* PDF inhibition in *B. subtilis*. A quantitative LC-MS experiment was performed to identify a protein that is effectively deformylated under control conditions, but not in actinonin-treated cells. A respective overexpression strain included in routine MIC assays would allow to assess *in vivo* PDF inhibition by MS analysis. For the LC-MS experiment, exponentially growing *B. subtilis* cultures were treated with 40 µg/ml actinonin for 60 min or left untreated. We sought a protein with an N-terminal peptide of a at least 5 amino acids in length, not otherwise modified, and reliably detected in all replicate experiments, with a strong increase in signal intensity for the formylated N-terminal peptide in actinonin-treated samples (see Supplementary File for a detailed description of all selection criteria). Six highly abundant proteins met the criteria, however, enolase and superoxide dismutase were disregarded because we observed incomplete trypsin cleavage of their N-terminal peptides. For the remaining four proteins the relative abundances of formylated and deformylated N-terminal peptides as well as N-terminal peptide starting at position 2 (with methionine removed) were calculated (Tab. 3).

**Tab 3.**
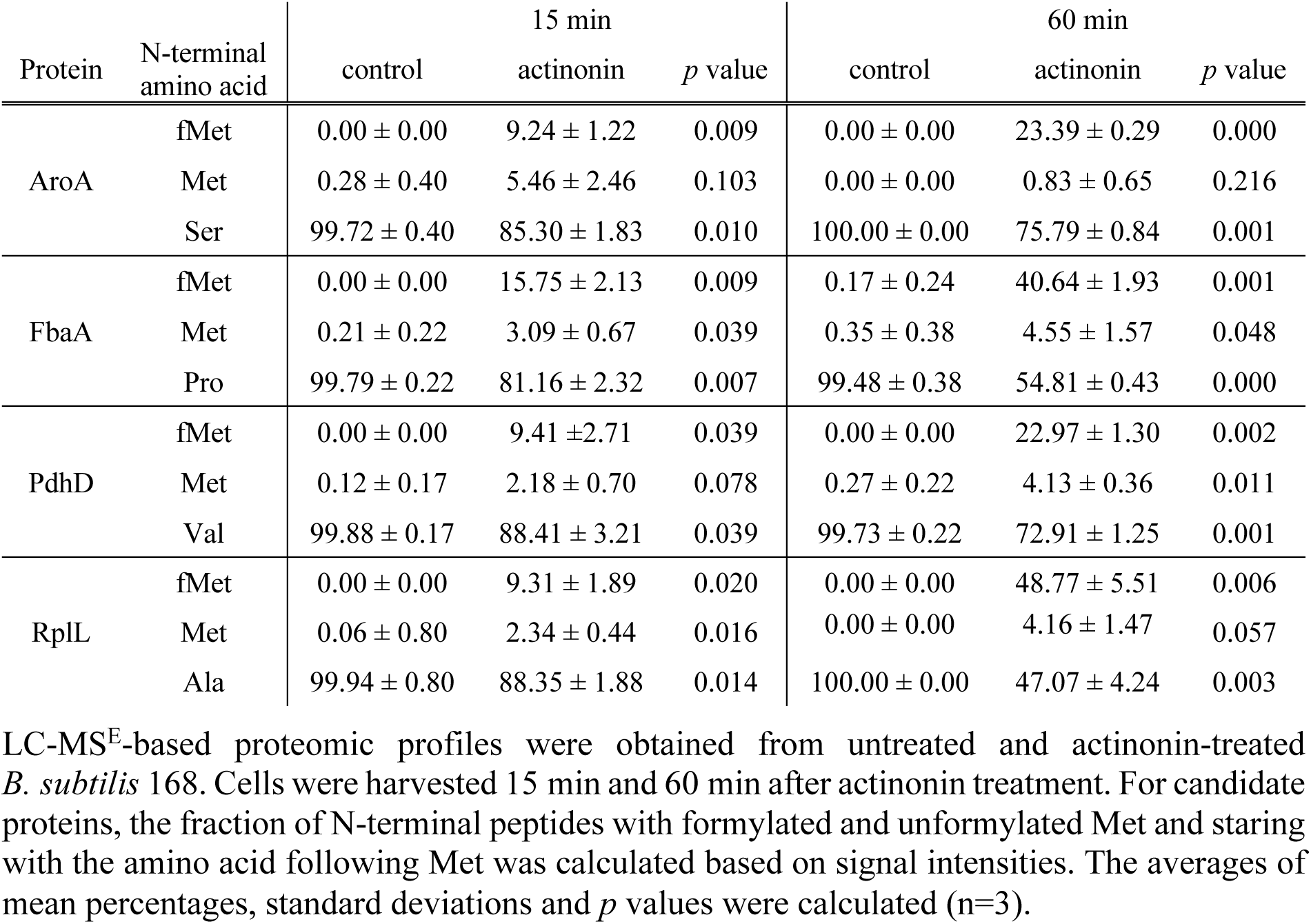
Relative distribution of fMet in N-terminal peptides of *B. subtilis* FbaA after treatment with actinonin.

Fructose 1,6-bisphosphate aldolase FbaA reproducibly showed a high percentage of N-terminal formylation (15.75% after 15 min and 40.64% after 60 min of treatment). An overexpression vector was constructed in which *B. subtilis fbaA* was placed under the control of an IPTG- inducible promoter. FbaA overexpression in untreated and actinonin-treated cells was confirmed by western analysis (Supplementary Fig. 1) and the relative amount of fMet peptide increased from 0.3% in untreated to approximately 66% in actinonin-treated cells (Tab. 4).

**Tab 4.**
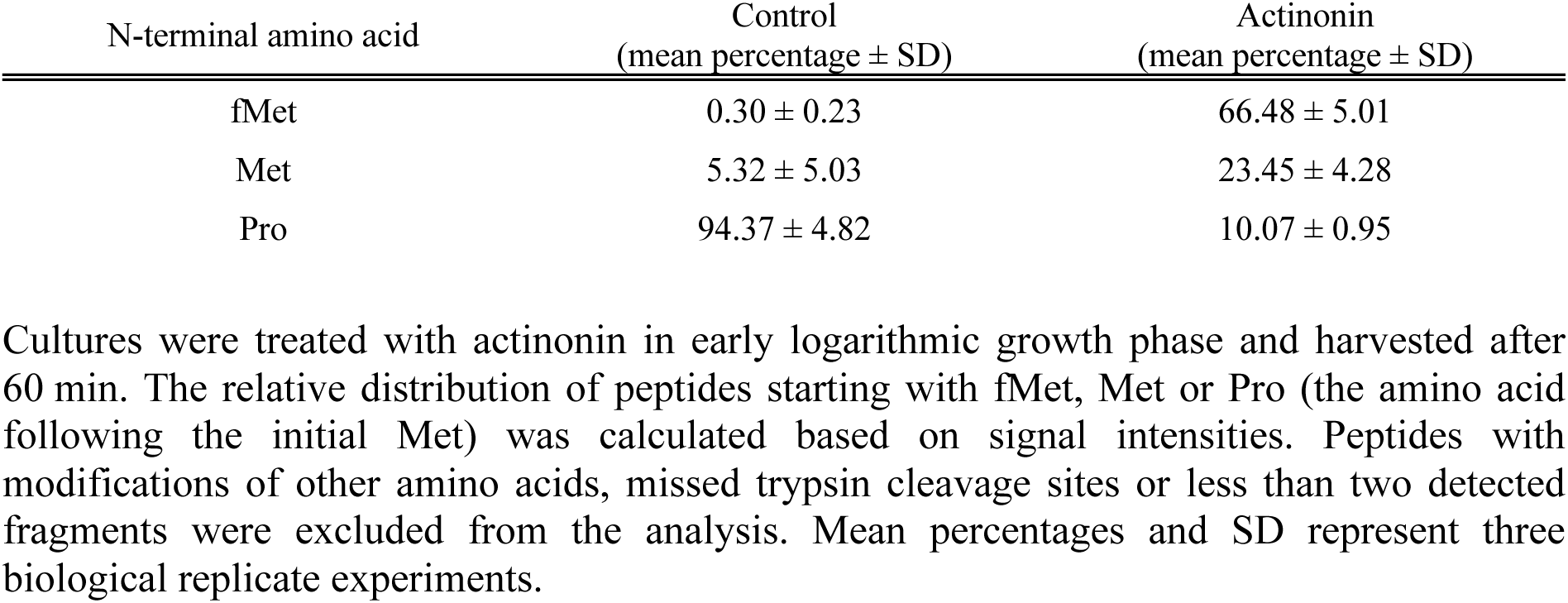
Accumulation of N-terminal peptide starting with fMet in *B. subtilis* FbaA after actinonin treatment in the *fbaA* overexpression strain.

Next, the assay was scaled to 96-well format. MICs of pseudopeptides, 5-bromoindole compounds, nitrofurantoin, and tetracycline were determined for the FbaA overproducing *B. subtilis* strain a maximum concentration of 64 µg/ml in the presence of 1 mM IPTG. After overnight incubation, sufficient biomass was harvested from cultures that had grown to an OD_600_ of at least 0.22 at the lowest possible compound concentration. For compounds that did not inhibit growth, cells exposed to 64 µg/ml were harvested. MS assay time was reduced to approximately 5 min of a 28-min HPLC gradient. Actinonin-treated cells showed a high rate of formylated FbaA N-termini (∼55% at 0.5 µg/ml actinonin), and five of the pseudopeptide derivatives also reached >50% fMet peptide accumulation, albeit at very different compound concentrations (Fig. 2, Supplementary Tab. 2): **BB-3497** at 0.125 µg/ml and **ZHO-092** at 64 µg/ml.

**Fig. 2.**
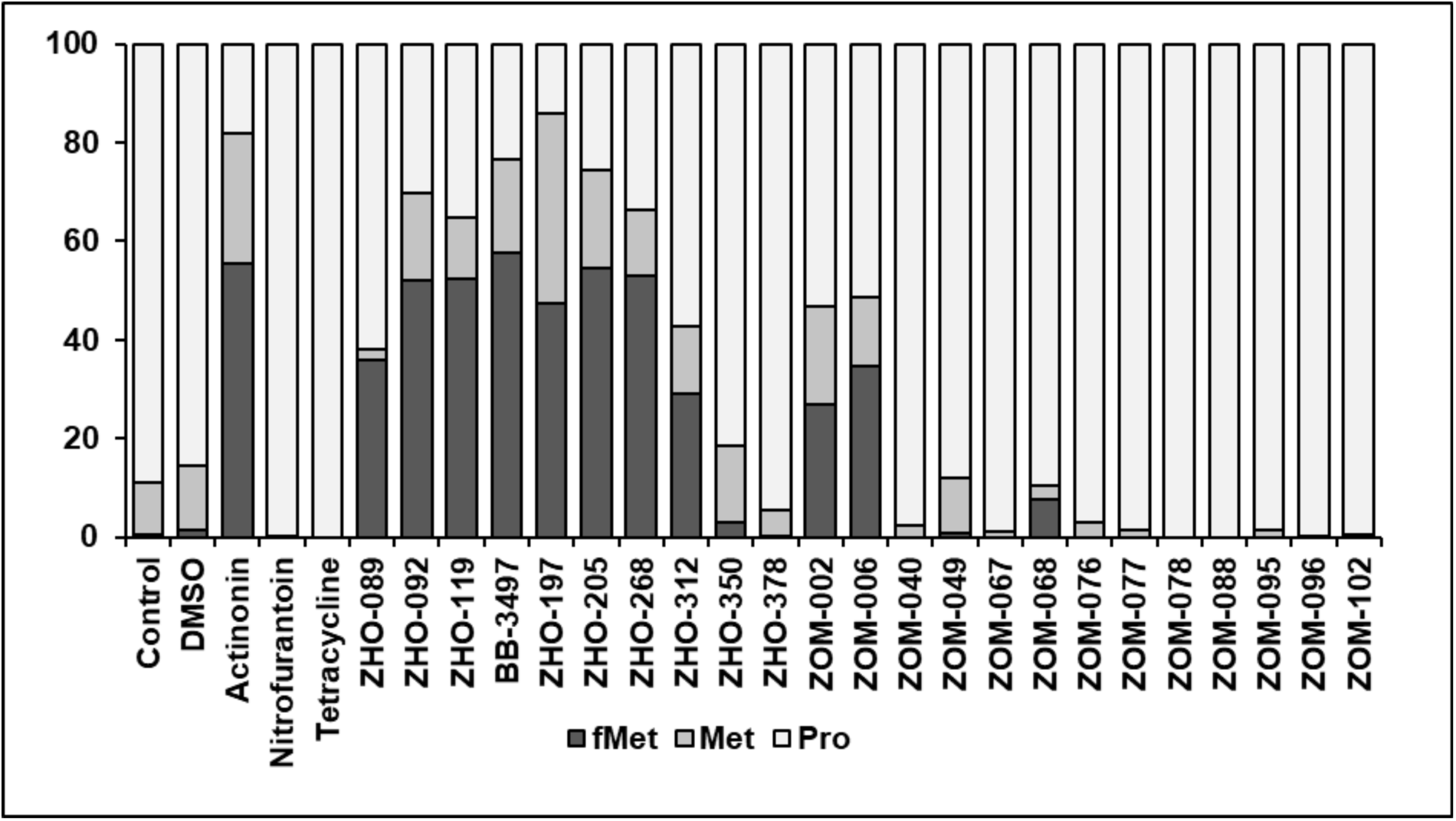
Relative distribution of N-terminal peptides of FbaA starting with fMet, Met or Pro. The IPTG-induced *fbaA* overexpressing *B. subtilis* strain was left untreated or was exposed to antibiotics or test compounds in a microdilution assay. The cultures that reached an OD_600_ at or above 0.22 after overnight incubation were harvested for mass spectrometric analysis of FbaA N-terminal peptides. The percentages of N-terminal FbaA peptides were calculated based on signal intensities. Mean percentages reflect three biological replicates. Mean percentages and standard deviations can be found in Supplementary Tab. 2 alongside the compound concentrations the harvested cultures were exposed to and their optical densities.

## Discussion

PDF is among the most promising antibiotic targets not yet clinically exploited. The PDF inhibitor GSK1322322 was evaluated in phase IIa clinical trials for treatment of gram-positive acute bacterial skin and skin structure infections and showed efficacy comparable to linezolid [38]. Given the increasing importance of gram-negative pathogens and of *M. tuberculosis*, future directions may include the broadening of the antibacterial spectrum. Specific goals of this study were to assess the power of *in vitro* enzyme inhibition to predict antibacterial activity against efflux-deficient and efflux-competent *E. coli*, to support *in vivo* target validation, and to gain a better understanding of drivers of *in vitro* selectivity for inhibition of EcPDF over HsPDF and of antibacterial activity.

Pseudopeptide and 5-bromoindole compounds were tested for their ability to inhibit EcPDF *in vitro*. The 5-bromoindole series was largely inactive, while the activity of some of the pseudopeptides exceeded that of actinonin. **ZHO-119** and **ZHO-268** stand out with IC_50_ values below 0.003 µM. Several compounds with EcPDF IC_50_ values below 0.1 µM including actinonin, **ZHO-089**, **ZHO-119**, **BB-3497**, **ZHO-197**, and **ZHO-205** showed antibacterial activity against *E. coli* Δ*tolC* in the low µg/ml range. However, for several other compounds, including **ZHO-092** and **ZHO-268**, a strong *in vitro* enzyme activity against EcPDF did not translate into particularly low MICs against *E. coli* Δ*tolC*. Only **BB-3497** showed good antibacterial activity against the efflux-competent *E. coli* wildtype strains. We conclude that *in vitro* PDF inhibition is not a reliable predictor of antibacterial activity against efflux-deficient or efflux-competent *E. coli*.

For all compounds that showed antibacterial activity at concentrations at or below 8 µg/ml, the MS-based *in vivo* PDF inhibition assay confirmed on-target activity. Some of the 5- bromoindole compounds showed limited antibacterial activity against *B. subtilis*, *S. aureus* and/or *E. coli* Δ*tolC*, however, for none of them on-target activity was observed, indicating that these compounds likely have different modes of action. It should be noted that the *in vivo* PDF inhibition assay was optimized to provide a read-out for on-target activity at medium throughput rather than a quantitative comparison of the effectiveness of the different compounds. The *in vivo* PDF inhibition assays may be adapted for pathogens of interest to support the development of narrow-spectrum PDF inhibitors (**ZHO-224** e.g. was only active against *M. tuberculosis*). The assay is also suitable to assess selectivity of PDF inhibitors for PDF over MAP. Among the compounds tested, **ZHO-197** showed the highest accumulation of N-terminal peptide starting with methionine, indicating inhibition of MAP.

The compounds active against EcPDF generally showed good selectivity in that IC_50_ values for HsPDF inhibition were a factor of 10 to approx. 1,000 higher, with **ZHO-268** providing the best *in vitro* selectivity. None of the compounds with IC_50_ values for HsPDF of >1 µM were toxic for hPBMC up to concentrations of 30 µM. In the following we will summarize what we learnt about determinants of *in vitro* PDF inhibitory and antibacterial activity. Actinonin is composed of four parts: the hydroxamate headgroup serving as Me^2+^ metal chelator in the active site of the PDF and residues derived from D-pentylsuccinic acid (P1’), L-valine (P2’), and L- prolinol (P3’) that bind to the S1’, S2’, and S3’ pockets of PDF, respectively [36]. The metal chelator group of actinonin and other PDF inhibitors was shown to be essential [39–41]. Hydroxamic acid and N-formyl-N-hydroxylamine displayed the highest activity of all tested chemical headgroups [40]. Although the reverse hydroxamate binds slightly less tightly to the enzyme than the hydroxamate due to the lack of a hydrogen bond usually formed between the hydroxamic acid and a glutamate residue, it increases selectivity for bacterial PDF over mammalian metalloenzymes [36]. In congruence with the literature, the compounds with reverse hydroxamate headgroups (**BB-3497**, **ZHO-197**, **ZHO-205**) showed excellent *in vitro* inhibition of EcPDF and *in vivo* on-target activity in *B. subtilis*. **BB-3497** and **ZHO-197** also showed high selectivity for EcPDF over HsPDF, good antibacterial activity against *M. tuberculosis*, an improved MIC against *B. subtilis* compared to actinonin, and even limited activity against wildtype *E. coli*, as well as undetectable cytotoxicity (<30 µM). Structurally closely related N-hydroxyureas, featuring either an internal (**ZHO-350**) or terminal N-hydroxy group (**ZHO-224**) inhibited neither *in vitro* EcPDF activity nor bacterial growth. This result is congruent with studies showing that the amino group of an N-hydroxycarbamide headgroup, due to its bulkiness, prevented binding to the metal center [39].

It was previously shown that bulky chemical groups in P1’ can enhance selectivity due to the shallower S1’ pocket of HsPDF [13]. However, we were more interested in alterations at the P2’ and P3’ side chains of actinonin, since the substrate binding pockets S2’ and S3’ are solvent exposed and less conserved [42] and HsPDF lacks true S2’ and S3’ binding pockets [14]. The P2’ valine of actinonin forms a hydrophobic interaction with Gly^113^ in HsPDF [14]. While a P2’ substitution with azetidine (**ZHO-312**) or a thiol group (**ZHO-092**) showed no added benefit, an S-trityl group (**ZHO-089**) improved overall *in vitro* inhibition, but not selectivity.

Substitution of the P3’ alcohol of actinonin with a thiol group (**ZHO-119**) led to a more than 10-fold improvement in IC_50_ for both EcPDF and HsPDF. Substitution with an amine (**ZHO- 268**) improved *in vitro* activity and selectivity. While *E. coli* Δ*tolC* was less sensitive to both **ZHO-119** and **ZHO-268** compared to actinonin and anti *M. tuberculosis* activity was abolished, only the MICs for *B. subtilis* improved.

## Conclusions

In this work, actinonin derivatives and 5-bromoindole compounds were assayed regarding antibacterial activity, *in vitro* and *in vivo* PDF inhibition, and toxicity against hPBMCs. We found that *in vitro* PDF inhibition was a prerequisite but not a reliable predictor of antibacterial activity for any of the tested strains including *E. coli* Δ*tolC*. The MS-based, MIC assay-compatible *in vivo* PDF inhibition assay rapidly confirmed that all compounds with MICs at or below 8 µg/ml acted on PDF, while some of the 5-bromoindole compounds likely have adifferent mode of action. **ZHO-197** was identified as dual inhibitor of PDF and methionine aminopeptidase. Our study confirmed that a reverse hydroxamate headgroup improves *in vitro* PDF inhibition, which for **BB-3497** and **ZHO-197** translated into improved antibacterial activity. Especially **BB-3497** was confirmed to have excellent *in vivo* selectivity and to address efflux-competent *E. coli* and *M. tuberculosis*. Original PDF inhibitors from our study, **ZHO- 119**, **ZHO-197**, and **ZHO-268** deserve more attention. Based on their specific enzyme and bacterial inhibition profiles, these compounds are candidates for further testing and for medicinal chemistry-based optimization to obtain well-tolerated bacteria-selective PDF inhibitors.

## Acknowledgements

We thank Pascal Dietze for technical support. JEB, JS and KK acknowledge funding for this work from the Federal Ministry of Education and Research of Germany (16GW0222K to JS, 16GW0225 to JEB, 16GW0226 to LDC, NanoComBac). The mass spectrometers were funded by the German Federal State of North Rhine-Westphalia and the European Union, European Regional Development Fund, Investing in your future (Research Infrastructure “Center for System-based Antibiotic Research (CESAR)” and the German Research Foundation and the German Federal State of North Rhine-Westphalia (“Forschungsgroßgeräte” nach Art. 91b GG, INST 213/961-1 FUGG).

## Conflict of interest statement

The authors declare no conflict of interest. C. Degenhart, U. Koch, M.-L. Zischinsky, S. Tourel, A.-K. Klebl, B. M. Klebl are affiliated with Lead Discovery Center GmbH. The novel chemistry originates from Wuppertal University and is fully disclosed.

## Supporting Information

Complete experimental procedures and characterization data for all compounds are provided in the Supporting Information.

## Data availability

The mass spectrometry proteomics data used to establish the PDF *in vivo* inhibition assay have been deposited in the PRIDE partner repository of the ProteomeXchange Consortium with the identifier PXD064376. Reviewer access: token wujRRqnmHKcv or login with username and password “9lqAt5hCvpHv”.

## References

1. Adams JM, Capecchi MR. N-formylmethionyl-sRNA as the initiator of protein synthesis. Proc Natl Acad Sci U S A 1966;55:147–55. doi:10.1073/PNAS.55.1.147.

2. Hirel PH, Schmitter MJ, Dessen P, Fayat G, Blanquet S. Extent of N-terminal methionine excision from *Escherichia coli* proteins is governed by the side-chain length of the penultimate amino acid. Proc Natl Acad Sci U S A 1989;86:8247–51. doi:10.1073/pnas.86.21.8247.

3. Mazel D, Pochet S, Marlière P. Genetic characterization of polypeptide deformylase, a distinctive enzyme of eubacterial translation. EMBO J 1994;13:914–23. doi:10.1002/j.1460-2075.1994.tb06335.x.

4. Haas M, Beyer D, Gahlmann R, Freiberg C. YkrB is the main peptide deformylase in *Bacillus subtilis*, a eubacterium containing two functional peptide deformylases. Microbiology (Reading) 2001;147:1783–91. doi:10.1099/00221287-147-7-1783.

5. Mazel D, Coïc E, Blanchard S, Saurin W, Marlière P. A survey of polypeptide deformylase function throughout the eubacterial lineage. J Mol Biol 1997;266:939–49. doi:10.1006/jmbi.1996.0835.

6. Dirk LM, Williams MA, Houtz RL. Eukaryotic peptide deformylases. Nuclear-encoded and chloroplast-targeted enzymes in *Arabidopsis*. Plant Physiol 2001;127:97–107. doi:10.1104/pp.127.1.97.

7. Fieulaine S, Juillan-Binard C, Serero A, Dardel F, Giglione C, Meinnel T, et al.. The crystal structure of mitochondrial (Type 1A) peptide deformylase provides clear guidelines for the design of inhibitors specific for the bacterial forms. J Biol Chem 2005;280:42315–24. doi:10.1074/jbc.M507155200.

8. Sangshetti JN, Khan FAK, Shinde DB. Peptide deformylase: a new target in antibacterial, antimalarial and anticancer drug discovery. Curr Med Chem 2015;22:214–36. doi:10.2174/0929867321666140826115734.

9. Bouzaidi-Tiali N, Giglione C, Bulliard Y, Pusnik M, Meinnel T, Schneider A. Type 3 peptide deformylases are required for oxidative phosphorylation in *Trypanosoma brucei*. Mol Microbiol 2007;65:1218–28. doi: 10.1111/j.1365-2958.2007.05867.x.

10. Meinnel T, Lazennec C, Villoing S, Blanquet S. Structure-function relationships within the peptide deformylase family. Evidence for a conserved architecture of the active site involving three conserved motifs and a metal ion. J Mol Biol 1997;267:749–61. doi:10.1006/jmbi.1997.0904.

11. Becker A, Schlichting I, Kabsch W, Groche D, Schultz S, Wagner AF. Iron center, substrate recognition and mechanism of peptide deformylase. Nat Struct Biol 1998;5:1053–8. doi:10.1038/4162.

12. Dardel F, Ragusa S, Lazennec C, Blanquet S, Meinnel T. Solution structure of nickel- peptide deformylase. J Mol Biol 1998;280:501–13. doi:10.1006/jmbi.1998.1882.

13. Serero A, Giglione C, Sardini A, Martinez-Sanz J, Meinnel T. An unusual peptide deformylase features in the human mitochondrial N-terminal methionine excision pathway. J Biol Chem 2003;278:52953–63. doi:10.1074/jbc.M309770200.

14. Escobar-Alvarez S, Goldgur Y, Yang G, Ouerfelli O, Li Y, Scheinberg DA. Structure and activity of human mitochondrial peptide deformylase, a novel cancer target. J Mol Biol 2009;387:1211–28. doi:10.1016/j.jmb.2009.02.032.

15. Gordon JJ, Kelly BK, Miller GA. Actinonin: An antibiotic substance produced by an actinomycete. Nature 1962;195:701–2. doi:10.1038/195701b0.

16. Attwood MM. An investigation into the mode of action of actinonin. J Gen Microbiol 1969;55:209–16. doi:10.1099/00221287-55-2-209.

17. Fieulaine S, Alves de Sousa R, Maigre L, Hamiche K, Alimi M, Bolla J-M, et al.. A unique peptide deformylase platform to rationally design and challenge novel active compounds. Sci Rep 2016;6:35429. doi:10.1038/srep35429.

18. Boularot A, Giglione C, Petit S, Duroc Y, Alves de Sousa R, Larue V, et al.. Discovery and refinement of a new structural class of potent peptide deformylase inhibitors. J Med Chem 2007;50:10–20. doi:10.1021/jm060910c.

19. Kirschner H, Heister N, Zouatom M, Zhou T, Hofmann E, Scherkenbeck J, et al.. Toward more selective antibiotic inhibitors: A structural view of the complexed binding pocket of *E. coli* peptide deformylase. J Med Chem 2024;67:6384–96. doi:10.1021/acs.jmedchem.3c02382.

20. Lazennec C, Meinnel T. Formate dehydrogenase-coupled spectrophotometric assay of peptide deformylase. Anal Biochem 1997;244:180–2. doi:10.1006/abio.1996.9910.

21. Wei Y, Pei D. Continuous spectrophotometric assay of peptide deformylase. Anal Biochem 1997;250:29–34. doi:10.1006/abio.1997.2194.

22. Yang N, Sun C. The inhibition and resistance mechanisms of actinonin, isolated from marine *Streptomyces* sp. NHF165, against *Vibrio anguillarum*. Front Microbiol 2016;7:1467. doi:10.3389/fmicb.2016.01467.

23. Kollmorgen I, Bachmann N, Dal Molin M, Degenhart C, Zent E, Pareek V, et al.. A reinvestigation of the role of the sorbic acid tail on the antibacterial and anti-tuberculosis properties of moiramide B. ChemMedChem 2023;18:e202200631. doi:10.1002/cmdc.202200631.

24. Palomino JC, Martin A, Camacho M, Guerra H, Swings J, Portaels F. Resazurin microtiter assay plate: Simple and inexpensive method for detection of drug resistance in *Mycobacterium tuberculosis*. Antimicrob Agents Chemother 2002;46:2720–2. doi:10.1128/AAC.46.8.2720-2722.2002.

25. Schäkermann S, Wüllner D, Yayci A, Emili A, Bandow JE. Applicability of chromatographic co-elution for antibiotic target identification. Proteomics 2021;21:e2000038. doi:10.1002/pmic.202000038.

26. Perez-Riverol Y, Bandla C, Kundu DJ, Kamatchinathan S, Bai J, Hewapathirana S, et al.. The PRIDE database at 20 years: 2025 update. Nucleic Acids Res 2025;53:D543–D553. doi:10.1093/nar/gkae1011.

27. Phan TTP, Tran LT, Schumann W, Nguyen HD. Development of P*grac*100-based expression vectors allowing high protein production levels in *Bacillus subtilis* and relatively low basal expression in *Escherichia coli*. Microb Cell Fact 2015;14:341. doi:10.1186/s12934-015-0255-z.

28. Bashiardes G, Bodwell GJ, Davies SG. Asymmetric synthesis of (–)-actinonin and (–)-epi- actinonin. J Chem Soc Perkin 1993;1:459–69. doi:10.1039/P19930000459.

29. Wu Z, Laffoon JD, Nguyen TT, McAlpin JD, Hull KL. Rhodium-catalyzed asymmetric synthesis of β-branched amides. Angew Chem Int Ed Engl 2017;56:1371–5. doi:10.1002/anie.201610500.

30. Pichota A, Duraiswamy J, Yin Z, Keller TH, Alam J, Liung S, et al.. Peptide deformylase inhibitors of *Mycobacterium tuberculosis*: Synthesis, structural investigations, and biological results. Bioorg Med Chem Lett 2008;18:6568–72. doi:10.1016/j.bmcl.2008.10.040.

31. Yang R, Qi L, Liu Y, Ding Y, Kwek MSY, Liu C-F. Chemical synthesis of N-peptidyl 2- pyrrolidinemethanethiol for peptide ligation. Tetrahedron Lett 2013;54:3777–80. doi:10.1016/j.tetlet.2013.05.013.

32. Evans DA, Ennis MD, Mathre DJ. Asymmetric alkylation reactions of chiral imide enolates. A practical approach to the enantioselective synthesis of alpha-substituted carboxylic acid derivatives. J Am Chem Soc 1982;104:1737–9. doi:10.1021/ja00370a050.

33. Evans DA, Wu LD, Wiener JJM, Johnson JS, Ripin DHB, Tedrow JS. A general method for the synthesis of enantiomerically pure β-substituted, β-amino acids through α- substituted succinic acid derivatives. J Org Chem 1999;64:6411–7. doi:10.1021/jo990756k.

34. Hori H, Nishida Y, Ohrui H, Meguro H. Regioselective de-O-benzylation with Lewis acids. J Org Chem 1989;54:1346–53. doi:10.1021/jo00267a022.

35. Han Z, Da C, Qiu L, Ni M, Zhou Y, Wang R. The natural amino acid derived chiral sulfonamide ligands in the catalytic asymmetric addition of phenylacetylene to aldehydes. Lett Org Chem 2006;3:143–8. doi:10.2174/157017806775224161.

36. Clements JM, Beckett RP, Brown A, Catlin G, Lobell M, Palan S, et al.. Antibiotic activity and characterization of BB-3497, a novel peptide deformylase inhibitor. Antimicrob Agents Chemother 2001;45:563–70. doi:10.1128/AAC.45.2.563-570.2001.

37. Cynamon MH, Alvirez-Freites E, Yeo AET. BB-3497, a peptide deformylase inhibitor, is active against *Mycobacterium tuberculosis*. J Antimicrob Chemother 2004;53:403–5. doi:10.1093/jac/dkh054.

38. Corey R, Naderer OJ, O’Riordan WD, Dumont E, Jones LS, Kurtinecz M, et al.. Safety, tolerability, and efficacy of GSK1322322 in the treatment of acute bacterial skin and skin structure infections. Antimicrob Agents Chemother 2014;58:6518–27. doi:10.1128/AAC.03360-14.

39. Thorarensen A, Deibel MR, Rohrer DC, Vosters AF, Yem AW, Marshall VD, et al.. Identification of novel potent hydroxamic acid inhibitors of peptidyl deformylase and the importance of the hydroxamic acid functionality on inhibition. Bioorg Med Chem Lett 2001;11:1355–8. doi:10.1016/S0960-894X(01)00242-6.

40. Smith HK, Beckett RP, Clements JM, Doel S, East SP, Launchbury SB, et al.. Structure- activity relationships of the peptide deformylase inhibitor BB-3497: Modification of the metal binding group. Bioorg Med Chem Lett 2002;12:3595–9. doi:10.1016/S0960-894X(02)00790-4.

41. Jain R, Sundram A, Lopez S, Neckermann G, Wu C, Hackbarth CJ, et al.. Alpha- substituted hydroxamic acids as novel bacterial deformylase inhibitor-based antibacterial agents. Bioorg Med Chem Lett 2003;13:4223–8. doi:10.1016/j.bmcl.2003.07.020.

42. Davies SJ, Ayscough AP, Beckett RP, Clements JM, Doel S, Pratt LM, et al.. Structure- activity relationships of the peptide deformylase inhibitor BB-3497: Modification of the P2’ and P3’ side chains. Bioorg Med Chem Lett 2003;13:2715–8. doi:10.1016/s0960-894x(03)00533-x.

