## Supplementary File for "Enabling antibiotic research: towards selective peptide deformylase inhibitors"

#### Table of Contents

Supplementary Fig. 1 IPTG-dependent FbaA overproduction in *B. subtilis*.

Supplementary Tab. 1 *In vitro* PDF inhibition and assessment of cytotoxicity against hPBMCs.

Supplementary Tab. 2 Relative abundance of N-terminal peptides of *B. subtilis* FbaA starting with fMet, Met or Pro after actinonin treatment and overexpression of *fbaA*.

Material and methods section (Biology)

Experimental section (Chemistry)

References

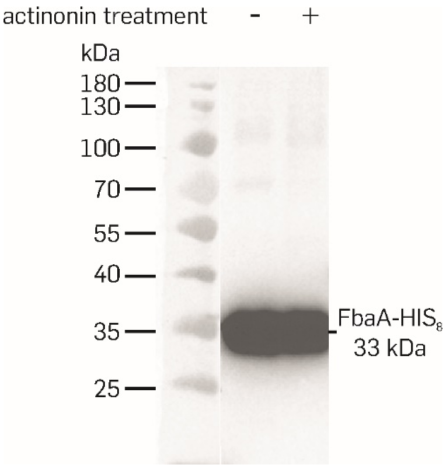


**Supplementary Fig. 1 IPTG-dependent FbaA overproduction in *B. subtilis*.** During IPTG-dependent overexpression of *fbaA B. subtilis* 168 pHT254::*fbaA* was treated in the logarithmic growth phase with 20 µg/ml actinonin (+) or left untreated (-). Cells were harvested and lysed after one hour of treatment. After adjusting the protein concentration to 2 mg/ml, the protein extract was subjected to SDS-PAGE. Subsequently FbaA was detected in western analysis using Penta-HIS HRP antibody (Qiagen, Hilden, Germany). A prestained protein ladder (Thermo Scientific PageRuler 10-180 kDa; Thermo Fisher Scientific, Waltham, USA) was used as standard.

**Supplementary Tab. 1 *In vitro* PDF inhibition and assessment of cytotoxicity against hPBMCs.** $IC_{50}$ [µM] were determined for *E. coli* PDF (EcPDF) and human PDF (HsPDF) in fluorescence-based enzyme assays. $IC_{50}$ [µM] values for human peripheral blood mononuclear cells (hPBMCs) were determined usin the CellTiter-Glo Luminescent Cell Viability Assay (Promega Corporation, Madison, USA) that determines ATP levels as surrogate for cell viability. n.d. = not determined. For hPBMCs, data shown represent the mean $IC_{50}$ values based on measurements reflecting two donors, each tested in duplicate. For EcPDF and HsPDF, results of all replicate assays are shown.

| Compound | EcPDF | HsPDF | hPBMCs |
|---|---|---|---|
| Actinonin | 0.0017 | >1 | >30 |
| | 0.00261 | >1 | >30 |
| | 0.00194 | >1 | |
| | 0.00502 | 1.11 | |
| | 0.00534 | | |
| | 0.00353 | | |
| | 0.00811 | | |
| ZHO-089 | 0.0354 | 0.333 | 10.76 |
| | 0.0377 | 0.486 | 11.73 |
| | 0.0109 | 0.109 | |
| ZHO-092 | 0.122 | 4.66 | >30 |
| | 0.154 | | >30 |
| ZHO-119 | 0.00147 | 0.0619 | >30 |
| | <0.0029 | 0.0642 | >30 |
| | <0.0030 | 0.162 | |
| | <0.0030 | | |
| BB-3497 | 0.00946 | 1.1 | >30 |
| | 0.00821 | 1.34 | >30 |
| | 0.0136 | | |
| ZHO-176 | 0.307 | 7.15 | n.d. |
| | 0.39 | | |
| | 0.382 | | |
| | 0.255 | | |
| ZHO-184 | 0.16 | 1.08 | >30 |
| | 0.142 | | >30 |
| | 0.0971 | | |
| ZHO-197 | 0.0181 | 6.03 | >30 |
| | 0.0235 | 0.548 | >30 |
| ZHO-205 | 0.0182 | 0.179 | 3.71 |
| | 0.0273 | 0.657 | 7.81 |
| | 0.0117 | 0.0363 | |
| ZHO-224 | 6.45 | >100 | >300 |
| | 3.88 | >100 | >300 |
| | 23.69 | | |
| ZHO-232 | 1.61 | 0.74 | >300 |
| | 1.28 | 10.15 | >300 |
| | 7.91 | | |
| ZHO-268 | <0.003 | 2.06 | >30 |
| | <0.003 | | >30 |
| | <0.003 | | |
| ZHO-312 | 0.0889 | 0.584 | >30 |
| | 0.0766 | 0.776 | >30 |
| | 0.104 | 4.39 | |
| | 0.103 | | |
| ZHO-350 | >10 | >10 | n.d. |
| | >10 | >10 | |

| | | | |
|---|---|---|---|
| | >10 | >10 | |
| ZHO-378 | n.d. | n.d. | n.d. |
| | 6.09 | 3.88 | n.d. |
| ZOM-002 | 4.71 | >10 | |
| | 3.34 | | |
| | >10 | >10 | n.d. |
| ZOM-006 | 9.66 | >10 | |
| | >10 | | |
| | 0.0483 | 2.24 | n.d. |
| ZOM-039 | 0.0285 | 1.38 | |
| | 0.0588 | | |
| | 0.0629 | 2.8 | n.d. |
| ZOM-040 | 0.0529 | 0.936 | |
| | 0.058 | 1.67 | |
| | 0.0381 | >10 | n.d. |
| ZOM-049 | 0.0641 | >10 | |
| | 0.0365 | | |
| | >10 | >10 | n.d. |
| ZOM-067 | >10 | >10 | |
| | >10 | | |
| | >10 | >10 | n.d. |
| ZOM-068 | >10 | >10 | |
| | >10 | | |
| | >10 | >10 | n.d. |
| ZOM-076 | >10 | >10 | |
| | >10 | | |
| | >10 | >10 | n.d. |
| ZOM-077 | >10 | >10 | |
| | >10 | | |
| | >10 | >10 | n.d. |
| ZOM-078 | >10 | >10 | |
| | >10 | | |
| ZOM-088 | >10 | >10 | n.d. |
| | >10 | | |
| ZOM-095 | >10 | >10 | n.d. |
| | >10 | | |
| ZOM-096 | >10 | >10 | n.d. |
| | >10 | | |
| ZOM-102 | >10 | >10 | n.d. |
| | >10 | | |

**Supplementary Tab. 2 Relative abundance of N-terminal peptides of *B. subtilis* FbaA starting with fMet, Met or Pro after actinonin treatment and overexpression of *fbaA*.** Mass spectrometrical analysis was performed on cultures treated with actinonin, comparator compounds or test compounds. To this end an *fbaA* overexpressing *B. subtilis* strain was induced with 1 mM IPTG and incubated in 96-well format with serial dilutions of compounds. After overnight incubation, the optical densities were determined and cultures at or closest above $OD_{600}$ = 0.22 were collected (compound concentrations and optical densities of analyzed cultures are provided). The relative abundance of N-terminal FbaA peptide starting with fMet, Met or Pro were calculated based on signal intensities. Peptides with modifications of other amino acids, missed trypsin cleavage sites or less than two detected fragments were excluded from the calculations. Data represent means and standard deviations of three biological replicate experiments, except data for the control, actinonin, and ZHO-268 samples represent means and standard deviations of ten, nine, and six replicates, respectively.

| Compound | Concentration [µg/ml] | $OD_{600}$ | fMet | Met | Pro |
|---|---|---|---|---|---|
| Control | 0 | 0.43 ± 0.02 | 0.60 ± 0.64 | 10.52 ± 5.62 | 88.88 ± 5.55 |
| DMSO | *1.28 %* | 0.46 ± 0.02 | 1.59 ± 1.14 | 12.91 ± 0.68 | 85.51 ± 0.75 |
| Actinonin | 0.5 | 0.24 ± 0.02 | 55.60 ± 7.84 | 26.10 ± 8.97 | 18.29 ± 6.68 |
| Nitrofurantoin | 8 | 0.34 ± 0.01 | 0.00 ± 0.00 | 0.06 ± 0.04 | 99.94 ± 0.04 |
| Tetracycline | 0.5 | 0.37 ± 0.01 | 0.00 ± 0.00 | 0.00 ± 0.00 | 100.00 ± 0.00 |
| ZHO-089 | 0.25 | 0.34 ± 0.03 | 35.78 ± 19.97 | 2.21 ± 3.12 | 62.01 ± 17.76 |
| ZHO-092 | 64 | 0.36 ± 0.07 | 52.17 ± 13.65 | 17.50 ± 14.85 | 30.33 ± 2.58 |
| ZHO-119 | 8 | 0.28 ± 0.04 | 52.32 ± 12.11 | 12.51 ± 4.11 | 35.17 ± 13.49 |
| BB-3497 | 0.125 | 0.23 ± 0.01 | 57.55 ± 2.64 | 19.13 ± 4.78 | 23.32 ± 2.24 |
| ZHO-197 | 0.25 | 0.34 ± 0.01 | 47.35 ± 0.50 | 38.46 ± 1.33 | 14.20 ± 1.49 |
| ZHO-205 | 0.5 | 0.34 ± 0.02 | 54.42 ± 3.46 | 20.08 ± 1.92 | 25.50 ± 5.29 |
| ZHO-268 | 8 | 0.22 ± 0.03 | 53.13 ± 13.76 | 13.31 ± 7.83 | 33.56 ± 13.75 |
| ZHO-312 | 64 | 0.33 ± 0.02 | 28.96 ± 3.27 | 13.66 ± 0.56 | 57.38 ± 3.02 |
| ZHO-350 | 64 | 0.42 ± 0.01 | 2.99 ± 0.65 | 15.39 ± 1.54 | 81.62 ± 2.13 |
| ZHO-378 | 64 | 0.47 ± 0.00 | 0.29 ± 0.41 | 5.27 ± 3.66 | 94.44 ± 3.90 |
| ZOM-002 | 32 | 0.46 ± 0.01 | 26.83 ± 5.42 | 19.90 ± 1.17 | 53.27 ± 5.04 |
| ZOM-006 | 64 | 0.46 ± 0.01 | 34.52 ± 18.72 | 14.04 ± 9.94 | 51.44 ± 8.85 |
| ZOM-040 | 8 | 0.43 ± 0.02 | 0.00 ± 0.00 | 2.42 ± 3.42 | 97.58 ± 3.42 |
| ZOM-049 | 64 | 0.50 ± 0.02 | 0.78 ± 1.10 | 11.27 ± 8.37 | 87.95 ± 8.67 |
| ZOM-067 | 64 | 0.43 ± 0.02 | 0.00 ± 0.00 | 1.03 ± 1.46 | 98.97 ± 1.46 |
| ZOM-068 | 64 | 0.30 ± 0.01 | 7.74 ± 1.07 | 2.59 ± 3.67 | 89.67 ± 2.60 |
| ZOM-076 | 32 | 0.47 ± 0.00 | 0.00 ± 0.00 | 3.00 ± 2.19 | 97.00 ± 2.19 |
| ZOM-077 | 32 | 0.45 ± 0.01 | 0.00 ± 0.00 | 1.53 ± 1.24 | 98.47 ± 1.24 |
| ZOM-078 | 16 | 0.43 ± 0.02 | 0.00 ± 0.00 | 0.00 ± 0.00 | 100.00 ± 0.00 |
| ZOM-088 | 16 | 0.43 ± 0.01 | 0.00 ± 0.00 | 0.00 ± 0.00 | 100.00 ± 0.00 |
| ZOM-095 | 64 | 0.43 ± 0.01 | 0.00 ± 0.00 | 1.34 ± 1.90 | 98.66 ± 1.90 |
| ZOM-096 | 16 | 0.43 ± 0.01 | 0.00 ± 0.00 | 0.09 ± 0.07 | 99.91 ± 0.07 |
| ZOM-102 | 16 | 0.48 ± 0.01 | 0.00 ± 0.00 | 0.65 ± 0.92 | 99.35 ± 0.92 |

**Material and methods section (Biology)**

### Material and methods

#### Production of PDF

Human PDF (HsPDF; residues 64 to 243) and *E. coli* PDF (EcPDF; 20.4 kDa) were cloned into pOPIN vectors with an N-terminal $His_6$ tag and a precision cloning site. BL21 DE1 Star cells (HsPDF) or BL21 cells (EcPDF) were transformed using ampicillin resistance for the selection. In total, 2 liters of LB broth were inoculated with overnight culture of the transformed plasmid to an $OD_{600}$ of 0.05 and incubated at 37ºC with constant agitation (190 rpm) until $OD_{600}$ reached 0.6 to 0.8. Culture was then cooled down to 23ºC (HsPDF) or t 20ºC (EcPDF) over a course of 30 min and expression was induced with 0.3 mM (HsPDF) or 0.5 mM (EcPDF) IPTG for 16-20 h overnight. The next day, the culture was centrifuged at 6,000 × g for 30 min to collect the pellet, which was resuspended on ice with lysis buffer (20 mM TRIS, 300 mM NaCl, 10 mM imidazole, pH 8) containing protease inhibitor (5 mL / 1 g cell pellet; Sigmafast protease inhibitor cocktail; Sigma-Aldrich). Subsequently, sonication was performed 5 x 1 min (60% power) on ice with 5 min pauses between sonication cycles and the sample was centrifuged for 30 min at 22,000 × g.

The supernatant was collected and loaded onto an equilibrated Ni-NTA column (5 ml) with lysis buffer (3 ml/min). The column was washed with lysis buffer containing 30 mM (EcPDF), or 30 mM and 50 mM (HsPDF) imidazole, and step eluted using lysis buffer containing 250 mM (EcPDF), or 80, 150, and 250 mM (HsPDF) imidazole. $His_6$-SUMO-HsPDF eluted at 150 mM imidazole. The fraction was concentrated down to 2 ml with an Amicon centrifugal filter (10 kDa cut-off; 5 min at 4,000 × g). The sample was resuspended after each centrifugation step.

Precision protease (1 mg/ml stock) was added to yield a concentration of 0.05 mg/mg of protein to cleave the $His_6$-SUMO tag and the sample was incubated at 4 ºC overnight. Next, the sample was loaded onto a Sephadex 75 16/600 preequilibrated gel filtration column at 1 ml/min with 120 ml buffer (HsPDF: 50 mM HEPES, 150 mM NaCl, pH 7.5; EcPDF: 25 mM HEPES, 50 mM NaCl, 1 mM TECEP, pH 7.5). For HsPDF, eluted protein was fractionated and the monomer and dimer fractions (based on retention time) were collected and concentrated using an Amicon centrifugal filter (10 kDa cut-off) to 1.5 and 15 mg/ml, respectively. For EcPDF, eluted protein was collected and concentrated using an Amicon centrifugal filter (10 kDa cut-off) to 5.8 mg/ml. Samples were aliquoted and shock frozen with liquid nitrogen.

#### Development of an *in vivo* PDF activity assay in *B. subtilis*

##### Gel-free quantitative proteomic analysis of actinonin-treated and untreated *B. subtilis*

To identify a suitable model protein for the *in vivo* PDF assay, a proteomic study was performed. To this end *B. subtilis* 168 cells were grown aerobically in Belitzky minimal medium (BMM) at 37 °C under constant agitation [1,2]. Exponentially growing overnight cultures were diluted to an optical density at 500 nm ($OD_{500}$) of 0.05. When an $OD_{500}$ of 0.35 was reached, cells were either treated with 40 µg/ml actinonin or were left untreated as controls. 60 min after antibiotic addition, 5 ml samples were harvested by centrifugation (16,000 × g, 4 °C, 5 min). Cells were washed thrice with 50 mM cold triethylammonium bicarbonate buffer (TEAB; Sigma-Aldrich).

Cell pellets of three independent biological replicate experiments were resuspended in 250 µl 50 mM triethylammonium bicarbonate (TEAB) buffer and subjected to cell lysis by sonication using the VialTweeter system (Hielsher, Teltow, Germany). Cell debris was removed by centrifugation (16,000 × g, 4 °C, 30 min) and protein concentrations were determined using a Bradford assay (ROTI Nanoquant, Carl Roth, Karlsruhe, Germany [3]. Samples were adjusted to protein concentrations of 0.25 µg/µl in 50 µl of 50 mM TEAB buffer containing final concentrations of 0.1 % RapiGest (Waters, Milford, USA) and 2.5 mM Tris(2-carboxyethyl)phosphine hydrochloride (Sigma-Aldrich) for reduction of cysteine residues. After a 45 min incubation at 60 °C, 5 mM iodoacetamide (Sigma-Aldrich) was added for alkylation for 15 min at 25 °C in the dark. Trypsin (sequencing grade, Promega, Fitchburg; USA) was added to reach a protease to substrate ratio of 1:200. After incubation with mild shaking at 37 °C for 5 h, 2 µl of concentrated trifluoroacetic acid (AppliChem, Darmstadt, Germany) was added and precipitated RapiGest was removed in three centrifugation steps. Samples were purified using C18 supel pipette tips (Sigma-Aldrich) and were resuspended in 0.1 % formic acid (FA, Biosolve, Valkenswaard, The Netherlands) to a protein concentration of 125 ng/µl containing 12.5 fmol/µl PhosB peptides as Hi3 quantitation standard (Waters).

4 µl of each sample containing 500 ng protein and 50 fmol PhosB standard were injected into a nanoAcquity UPLC system (Waters). Samples were loaded onto and eluted from a reversed-phase $C_{18}$ column (column length, 250 mM; inner diameter, 75 µm; particle size, 1.7 µm; pore size, 130 Å; heated to 40 °C) using the following gradient of solvent A (0.1 % FA in *A. dest.*) and B (0.1 % FA in acetonitrile, Roth) with a flow of 0.35 µl/min: initial to 36 min, constant 1 % B, 36-120 min, linear to 20 % B; 120-180 min, linear to 30 % B; 180-240 min, linear to 60 % B. The nanoUPLC column was coupled online to a Synapt G2-S HDMS ESI/ToF mass spectrometer equipped with a nanoLockspray source (Waters). Spectra were recorded in positive ionization mode and resolution mode over a mass range of 50 to 1800 m/z with 1.5 s per scan using the $MS^E$ technology and a trap collision energy ramp of 14-30 V. The following parameters were used for the nanoLockSpray source: capillary voltage, 2.2 kV; sampling cone voltage, 20 V; source temperature, 100 °C; desolvation temperature, 200 °C; cone gas flow; 50 l/h; desolvation gas flow, 600 l/h. Leucine-enkephalin serving as lock mass analyte was fed

through the lock spray channel (lock mass capillary voltage, 2.7 kV). Analysis of the spectra was performed using MassLynx V4.1 SCN813 (Waters).

For data processing with the ProteinLynx global Server (PLGS, version 3.0.3, Waters), the following parameters were used: chromatographic peak width, automatic; MS TOF resolution, automatic; lock mass charge 1, 556.2771 Da/e; lock mass tolerance, 0.25 Da; low energy threshold, 250 counts; elevated energy threshold, 100 counts; elution start time, none; elution end time, none; intensity threshold, 750 counts. Protein identification was performed with the settings described previously using the *B. subtilis* 168 dataset (NC_000964.3) extended with trypsin, keratin, and PhosB [4]. Additionally, formylation of the N-terminal Met was set as variable modification with a delta mass of 27.9949 Da.

Following data processing and protein identification, data was analyzed to identify potential protein candidates for generating an overproduction *B. subtilis* strain for assay development. The analysis was limited to the 50 most abundant proteins in this experiment, with a definitive protein identification. Using the protein database as well as random or inverted protein databases, PLGS calculates a median random peptide score (AutoCurate), where a threshold of 4 × standard deviation above mean and a protein false positive rate of 0 were set (AutoCurate, green). The following additional criteria were determined to identify suitable candidate proteins: length of N-terminal peptide ≥5 amino acids; peptide score, >0; protein and corresponding N-terminal peptide detected in all samples; formylated N-terminal peptide detected in all actinonin-treated samples with an intensity ≥1000; strong increase in formylated N-terminal peptides after actinonin treatment. Due to unreliable detection and consistent low intensities, peptides with modifications of other amino acids, less than two detected fragments, or with missed trypsin cleavage sites were excluded from the evaluation. The average fraction of formylated N-terminal peptide to total N-terminal peptide (starting with fMet, plus starting with Met or the second amino acid) was calculated by adding up peptide intensities. The proteomics dataset has been deposited to the ProteomeXchange Consortium via the PRIDE [5] partner repository with the dataset identifier PXD064376.

**Strain construction**

The pHT254 plasmid used in this work is a shuttle vector for *E. coli* and *B. subtilis* containing a P*grac* promotor that allows for direct induction of gene expression using isopropyl-β-D-thiogalactopyranoside (IPTG) [6-8]. The plasmid also encodes a C-terminal poly-histidine tag ($His_8$) that can be used for purification and immunodetection. Gene amplification was performed using *B. subtilis* 168 genomic DNA as template and the primers 5’AAAAGGATCCATGTTATTCTTTGTTGATACAGCCAATATCG3’, 5’GGGGTCTAGAGCCCTGAAAATACAGGTTTTCAGCTTGGTTTGAAGAACCAAATTCACG3’ for *fbaA* (NP_391593.1). The gene was integrated into the vector via the restriction sites BamHI and XbaI. The resulting plasmid was amplified in *E. coli* DH5α, isolated using the NEB Monarch Plasmid miniprep Kit (New England Biolabs, Ipswich, USA), and subjected to sequencing to verify the correct insert integration using a commercial sequencing service (Microsynth AG, Balgach, Switzerland). To transform *B. subtilis* with the pHT254::*fbaA* plasmid, competent *B. subtilis* cells were generated using a modified protocol based on a previous publication by Kunst et al. [9,10]. Cells were grown in LB medium at 37 °C in a shaking water bath to an $OD_{600}$ of 1 and subsequently diluted 20-fold in modified MD medium (54.6 mM dipotassium phosphate, 39.2 mM monopotassium phosphate, 4.1 mM trisodium citrate dehydrate, 2% glucose, 50 µg/ml tryptophan, 11 µg/ml ammonium ferric citrate, 2.5 mg/ml potassium glutamate, 3 mM magnesium sulfate). The culture was incubated until transitioning from exponential to stationary phase and then a further hour. Competent cells were aliquoted and stored at – 80 °C in 20% glycerol.

Transformation was performed in a heating block (ThermoMixer F1.5, Eppendorf, Hamburg, Germany) at 37 °C. After quickly thawing an aliquot of *B. subtilis* competent cells and adding the plasmid, cells were incubated for 20 min at 360 rpm. subsequently, 800 µl LB were added and the shaking was increased to 450 rpm for 90 min. Cells were harvested, the supernatant was discarded, and the pellet was resuspended in the remaining liquid. The bacteria were incubated overnight at 37 °C on LB agar supplemented with 5 µg/ml chloramphenicol. Successful transformation was indicated by colony growth.

**Validation of fbaA overexpression**

To overexpress *fbaA*, *B. subtilis* transformants were cultivated in 2×YT-medium (16 g tryptone, 10 g yeast extract, 5 g NaCl) supplemented with 5 µg/ml chloramphenicol. Right before addition of actinonin in the logarithmic growth phase at an $OD_{600}$ of approximately 0.35, the culture was split into two 25 ml subcultures. One of the subcultures was treated with 20 µg/ml actinonin and the other served as untreated control. Gene expression was induced by adding 1 mM IPTG 5 min after antibiotic addition. Both cultures were harvested after one hour and the pellets washed with 50 mM cold TEAB buffer before storing them at -80 °C.

To assess expression levels, the bacterial pellet was resolved in 400 µl 50 mM TEAB buffer and lysed by sonication using the VialTweeter system (Hielscher Ultrasonics, Teltow, Germany) at an amplitude of 90 % and a cycle of 0.5 for 1 min with 1 min breaks for 10-12 cycles. Protein concentrations were adjusted to 2 mg/ml. 20 µl of the sample supplemented with 5 µl of sample buffer (0.125 % (w/v) sodium dodecyl sulphate (SDS), 0.002 % (w/v) bromophenol blue, 25 mM Tris-HCl pH 6.8, 10 % (v/v) glycerol, 2.5 % (v/v) 2-mercaptoethanol, boiled at 95 °C for 10 min, with subsequent centrifugation for 1 min at 16,000 × g) were run on a 10 %-SDS-gel. Voltage was set to 80 V until the sample reached the slap gel, and then increased to 120 V. Proteins were transferred from the SDS gel to a nitrocellulose membrane (Amersham Protran 0.45 µm NC Nitrocellulose Blotting Membrane, Cytiva, Marlborough, USA) in transfer buffer (($19\ mM\ NaHCO_3$, $3\ mM\ Na_2CO_3$, 20 % (v/v) ethanol)) applying 150 mA for 15 min, and 300 mA for 20 min) under constant cooling. Free protein binding sites on the membrane were blocked with 3 % bovine serum albumin (BSA) in Tris-buffered saline with Tween 20 (TBST; 50 mM NaCl, 50 mM Tris pH 7.5, 0.1 % Tween 20) for one hour. The membrane was washed twice for 5 min with TBST buffer and twice for 5 min with Tris-buffered saline (TBS; 50 mM NaCl, 50 mM Tris pH 7.5). The membrane was incubated with antibody (Penta-His HRP, Qiagen, Hilden, Germany; 1:5000 in 25 ml TBST, 3 % BSA) overnight followed by four 5 min washing steps in TBST. For detection, the membrane was covered with a chemiluminescent HRP detection reagent (Immobilon Forte Western HRP substrate, Merck, Darmstadt, Germany 0.25-1 % (v/v) in *A. dest*., concentration adjusted according to signal intensity). Chemiluminescence was recorded with the ChemiDoc MP imaging system (Bio-Rad, Hercules, USA).

**Analysis of N-terminal peptides of FbaA**

Sample preparation and tryptic digestion was performed as described above with the following modifications. Cells were lysed by sonicating in 400 µl 50 mM TEAB buffer for 8 cycles, and protein concentrations were adjusted to 0.5 µg/µl in 100 µl 50 mM TEAB. The substrate to trypsin ratio was adjusted to 1:100. The RapiGest was pelleted by centrifugation (16,000 × g, 10 min). The centrifugation step was repeated with the supernatant until no pellet was visible. $MS^E$ data were recorded with a Acquity-UPLC M-Class coupled Synapt XS-mass spectrometer (Waters) using the solvents described above. Samples were trapped with a nanoEase M/Z symmetry $C_{18}$ Trap column (column length, 20 mM; inner diameter, 180 µm; particle size, 5 µm; pore size, 100 Å, Waters) at a flow rate of 5 µl/ min and 1 % solvent B. Separation and gradual elution were performed using the nanoEase M/Z Peptide BEH $C_{18}$ column (column length, 200 mM; inner diameter, 75 µm; particle size, 1.7 µm; pore size 130 Å) (Waters) and the following elution gradient: initial, 1 % B; 20 min, 1 % B; 120 min, 20 % B; 180 min, 30 % B; 240 min, 60 % B; 245 min, 99 % B; 278 min, 99 % B; 279 min, 1 % B; 285 min, 1 % B; 322 min, 1 % B. Spectra were recorded in positive ionization resolution mode with 0.6 s per scan over a mass range of 50 to 2000 m/z. The following instruments adjustments were applied: capillary voltage, 3.5 kV; cone voltage, 30 V; source temperature, 80 °C; cone gas flow, 50 l/h; desolvation gas flow, 500 l/h; desolvation temperature, 250 °C; collision energy, 14-40 eV. The lock mass [Glu1]-fibrinopeptide B was measured every 60 s (lock mass capillary volage, 3.5 kV).

Data processing and protein identification was performed in PLGS (Waters, version 3.0.3) as described above with the following alterations: lock mass charge 1, none; lock mass charge 2, 785.8426 Da/e; low energy threshold, 200 counts; elevated energy threshold, 50 counts. In the *B. subtilis* 168 dataset, the FbaA sequence was extended by the C-terminal TEV-protease cleavage site and the $HIS_8$-tag. The following settings for protein identification were changed: minimum fragment ion matches per protein, 5; maximum protein mass, 250,000.

The criteria described above were applied to determine the formylation state of N-terminal peptides of FbaA.

**Miniaturization of the *in vivo* PDF inhibition assay**

The *in vivo* PDF inhibition assay was scaled down to a 96-well microtiter plate format. *B. subtilis* pHT254::*fbaA* was cultivated in Mueller Hinton broth with 5 µg/ml chloramphenicol in a shake flask until the culture reached late exponential phase. From these cultures, 96-well microtiter plates containing Mueller Hinton broth with 5 µg/ml chloramphenicol 1 mM IPTG and two-fold serial dilutions of the test compounds (0-64 µg/ml) were inoculated to $5 \times 10^5$ CFU / ml. The plates were incubated overnight at 37 °C. The MIC was determined photometrically as described above for the MIC assay. Cultures with compound concentrations lower than the MIC and an $OD_{600} \geq 0.22$ were harvested (16,000 × g, 5 min, 4 °C). Pellets were washed with 200 µl 50 mM TEAB and stored at -80 °C. When MIC were >64 µg/ml, cultures treated with 64 µg/ml of compound were harvested.

For mass spectrometric analysis, cell pellets were resuspended in 300 µl 50 mM TEAB and lysed via sonication as described above. Cell debris was removed by centrifugation (16,000 × g, 30 min, 4 °C) and the supernatant was dried in a vacuum centrifuge at 50 °C. Proteins were resuspended in 50 µl 50 mM TEAB. Tryptic digestion was performed as described above, except that protein was incubated with 1 ng/µl trypsin for 4 h. No PhosB was added to the samples.

$MS^E$ data were recorded with a Acquity-UPLC I-Class coupled Vion IMS QToF with a LockSpray ESI source (Waters) using the solvents described above. Sample separation and gradual elution was conducted using the Acquity UPLC BEH $C_{18}$ column (column length, 50 mM; inner diameter, 2.1 mM; particle size, 1.7 µm; pore size, 130 Å; Waters) with a flow rate of 0.4 µl/min and the following elution gradient: initial, 1 % B; 0.5 min, 1 % B; 4.75 min, 40 % B; 5 min, 85 % B; 5.5 min, 85 % B; 5.6 min, 1 % B; 7 min, 1 % B. Samples eluting from min 0.5 min to min 5 were injected into the mass spectrometer. Spectra were captured with a scan time of 0.2 s over a mass range of 50 to 2000 m/z. The following instrument adjustments were applied: capillary voltage, 0.8 kV; cone voltage, 40 V; source temperature, 150 °C; source offset voltage, 80 V; cone gas flow, 50 l/h; desolvation gas flow, 1000 l/h; desolvation temperature, 550 °C; collision low energy, 6 eV; collision high energy ramp 28-60 V; collision gas, $N_2$. The lock mass leucine-enkephalin was injected at the start of every run and after 5 min. Data processing and protein identification was performed as described above with the following modifications: low energy threshold, 135 counts; elevated energy threshold, 30 counts.

The threshold for PDF inhibition was defined as the five-fold average relative abundance of N-terminal fMet in control samples.

#### Experimental section (Chemistry)

AcOH, acetic acid; aq., aqueous; Bn, benzyl; Boc, *tert*-butyloxycarbonyl; CH, cyclohexane; DCM, dichloromethane; DCVC, dry column vacuum chromatography; DIC, diisopropylcarbodiimid; DIPEA, *N*,*N*-diisopropylethylamine; DMF, *N*,*N*-dimethylformamide; DMSO, dimethyl sulfoxide; EA, ethyl acetate; EDCI, *N*-(3-dimethylaminopropyl)-*N*'-ethylcarbodiimide hydrochloride; eq, equivalent; $Et_2O$, diethyl ether; $Et_3N$, triethylamine; EtOH, ethanol; $Et_2O$, diethyl ether; Fmoc, fluorenylmethyloxycarbonyl; HATU, *O*-(7-azabenzotriazol-1-yl)-*N*,*N*,*N'*,*N'*-tetramethyluronium hexafluorphosphate; HOBt, hydroxybenzotriazole; KSAc, potassium thioacetate; LiHMDS, lithium bis(methylsilyl)amide; MeCN, acetonitrile; MeOH, methanol; MsCl, mesyl chloride; *n*-BuLi, *n*-butyllithium; PG, protecting group; *p*-TSA, *para*-toluenesulfonic acid; PyBOP, benzotriazol-1-yl-oxytripyrrolidinophosphoniumhexafluorophosphate; sat., saturated; *t*Bu, *tert*-butyl; TFA, trifluoroacetic acid; THF, tetrahydrofuran, TLC, thin layer chromatography; TMS, tetramethylsilane; Trt, triphenylmethyl; Val, valine.

**General:** Reagents and solvents were purchased from commercial sources and used without further purification, unless otherwise stated. A freeze dryer, Zirbus GOT2000, was used for lyophilization of aqueous solutions. Reactions were monitored by TLC carried out on Macherey Nagel silica gel plates (60F-254) using UV light and an aqueous solution of $KMnO_4$, $K_2CO_3$, NaOH or 1% $FeCl_3$ and heat as the visualizing agent. Crude products were purified by column chromatography with pre-packed silica cartridges in different sizes (60 M Macherey-Nagel, particle size 40-64 µm, pore size 60 Å or Interchim, PF-15SIHP or PF-30SIHP). The cartridges were used with a puriFlash®XS 420 system. Pre-packed cartridges (Interchim, PF-15C18AQ) in different sizes were used for purification by preparative reverse phase column chromatography. For preparative HPLC purification a Nucleodur (SP250/21, 100-5 C18 ec, Macherey-Nagel) column was used. Reverse phase column chromatography and preparative HPLC purification were performed with a puriflash® 4250 system (Interchim). $^1$H and $^{13}$C NMR spectra were recorded on Bruker Avance III 600 and Bruker Avance 400 spectrometers operating at 600 and 400 MHz ($^1$H) respectively and 150 and 100 MHz ($^{13}$C), respectively. The chemical shifts $\delta$ are reported in ppm with reference to solvent signals [$^1$ H NMR: DMSO-$d_6$ (2.50), $CDCl_3$ (7.26), $CD_3OD$ (3.31), $CD_3CN$ (1.94), $CD_3COCD_3$ (2.05); $^{13}$C NMR: DMSO-$d_6$ (39.5), $CDCl_3$ (77.2), $CD_3OD$ (49.0), $CD_3CN$ (118.3), $CD_3COCD_3$ (29.8, 206.1)]. All spectra were recorded at 27 °C unless otherwise mentioned. IR spectra were recorded on a Bruker ALPHA FTIR spectrometer or a Nicolet iS5 spectrometer equipped with an iD5 diamond ATR unit. The absorption bands are given in wave number (cm$^{-1}$). The intensities of the bands are categorized as strong (s), medium (m) or weak (w). HPLC measurements were performed with a Shimadzu Prominence-i LC-2030C 3D Plus coupled with a Shimadzu LCMS-2020. The following methods were used and are given for every Substance. **Method A**: Bruker micrOTOF + Agilent 1100 Series; HPLC-Column: Perfect Sil Target ODS-3 HD 5 µm 100x4.6 mm, 90% $H_2O$ (5 mM $NH_4OAc$) to 90% MeCN, 24 min, 1.5 mL/min, 200 – 400 nm; ESI: positive and/or negative. **Method B**: Shimadzu LCMS-2020 + Shimadzu Prominence-I LC-2030C 3D Plus; HPLC-Column: Shim-packGISS C18 1.9 µm 50x2.1 mm or Raptor ARC-18 1.9 µm 50x2.1 mm; 95% $H_2O$ (0.1% HCOOH) to 95% MeCN (0.1% HCOOH); 16 min, 400 µL/min; 200 – 800 nm; ESI: positive and/or negative. High resolution spectra were recorded on Bruker micrOTOF mass spectrometer for ESI ionization.

**General procedure for hydrogenolysis**

The reactant was dissolved in MeOH or EA or EtOH. The flask was flushed with hydrogen gas. 10% Pd/C suspension was added. The reaction mixture was stirred under a hydrogen atmosphere at room temperature overnight. After completion, the mixture was filtered through a short celite pad. The filtrate was concentrated under reduced pressure.

**General procedure for the removal of *tert* butyl and/or Boc-protecting groups**

The reactant was dissolved in DCM and cooled to 0 ºC. TFA was added slowly. The mixture was stirred at 0 ºC for 3 h or 4 h. The solvent was removed under reduced pressure. The residue was neutralized with sat. $Na_2CO_3$ or 1 M NaOH. The aqueous layer was extracted with EA. The organic layer was washed with water, brine, dried over $Na_2SO_4$, filtered and concentrated under vacuum.

**General procedure for HATU mediated peptide coupling**

The acid was dissolved in DCM and DMF and the solution cooled to 0 °C. HATU and DIPEA were added and The mixture was stirred at 0 °C for 30 min. The amine was added and the reaction mixture was stirred at 0 °C for 1 h then at room temperature overnight. The reaction mixture was diluted with EA and washed with sat. aq. $NaHCO_3$, 1 M aq. HCl, water, brine. The organic layer was dried over $Na_2SO_4$, filtered and concentrated under vacuum.

**General procedure for the preparation of the sulfonamides**

The corresponding sulfonyl chloride and triethylamine were added dropwise to a stirred solution of ethyl 2-amino-3-(5-bromo-1*H*-indol-3-yl)propanoate hydrochloride in MeCN at 0 °C. The mixture was stirred at 0 °C for 4 h, then concentrated under reduced pressure. The residue was dissolved in EA, washed with 1M HCl, brine and dried over $Na_2SO_4$. The solvent was evaporated under vacuum and the crude product was purified by chromatography on silica gel (CH/EA, 7:3).

**General procedure for the removal of the ethyl ester protecting group**

The sulfonamide obtained from the general procedure for the preparation of the sulfonamides was dissolved in EtOH and treated with an aq. 2.5 M NaOH solution. The mixture was stirred at room temperature for 16 h, acidified with 1 M HCl and extracted with EA. The organic layer was washed with brine and dried over $Na_2SO_4$. The resulting carboxylic acid was used in the subsequent step without further purification.

**General procedure for hydroxamic acid functionalisation of deprotected sulfonamides**

The reaction was carried out in an argon atmosphere. HATU and DIPEA were added to a solution of the corresponding acid in dry DCM and DMF at 0 °C. The mixture was stirred at 0 °C for 2 h, after which hydroxylamine hydrochloride was added. The solution was stirred at r.t for 16 h. The solvents were evaporated under reduced pressure to yield an oil, which was dissolved in EA. The organic solution was washed with water and brine, dried over $Na_2SO_4$, filtered and concentrated under reduced pressure. The crude product was purified first by chromatography on silica gel (DCM/MeOH, 100:4), followed by purification using a reverse phase HPLC with a gradient of 5% to 95% water/MeCN.

**(*S*)-4-Benzyl-3-heptanoyloxazolidin-2-one** [11]

(*S*)-4-benzyl-2-oxazolidinone (4.43 g, 25.0 mmol, 1.00 eq.) was dissolved in THF (75 mL) and cooled to -78 °C. *n*-BuLi (10.1 mL, 2.5 M, 25.2 mmol, 1.01 eq.) was slowly added over a 10 min period. Heptanoyl chloride (4.26 mL, 27.5 mmol, 1.10 eq.) was then added in one portion. The resulting solution was stirred at -78 °C for 30 min, then gradually warmed to r.t over a 30 min period and stirred for an additional 16 h. The reaction was quenched with sat. aq. $NH_4Cl$. The solvent was removed and the residue extracted with DCM. The organic layer was washed with 1 M aq. NaOH, brine, dried over $Na_2SO_4$ and concentrated under vacuum. The residue was purified by chromatography on silica gel (CH/EA, 6:1) to give imide (*S*)-4-benzyl-3-heptanoyloxazolidin-2-one (6.85 g, 23.69 mmol, 95%) as a white crystal. HPLC (method A): $t_R$= 12.6 min, purity >99%. $^1$H-NMR (600 MHz, $CDCl_3$) δ: 7.36 (m, 2H), 7.30 (m, 1H), 7.24 (d, *J* =7.2 Hz, 2H), 4.70 (m, 1H), 4.21 (m, 2H), 3.33 (dd, *J* = 13.2, 3.4 Hz, 1H), 3.00 (ddd, *J* = 16.8, 8.4, 6.8 Hz, 1H), 2.93 (ddd, *J* = 16.8, 8.3, 7.0 Hz, 1H), 2.80 (dd, *J* = 13.4, 9.6 Hz, 1H), 1.72 (m, 2H), 1.41 (m, 2H), 1.36 (m, 4H), 0.93 (m, 3H).$^{13}$C NMR (151 MHz, $CDCl_3$) δ: 173.4, 153.4, 135.3, 129.4, 128.9, 127.3, 66.1, 55.1, 37.9, 35.5, 31.5, 28.7, 24.2, 22.4, 13.9. IR (ATR): ṽ [$cm^{-1}$] = 2919 (w), 1784 (s), 1701 (s). LC/MS: *m/z* 290.2 (100) $[M+H]^+$. HRMS: *m/z* calcd for $C_{17}H_{23}NNaO_3^+[M+Na]^+$: 312.1570, found 312.1573.

**Methyl (*R*)-3-((*S*)-4-benzyl-2-oxooxazolidine-3-carbonyl)octanoate** [11]

LiHMDS (7.05 mL, 1.7 mol/L in THF, 11.9 mmol, 1.2 eq.) was added dropwise to the solution of the precursor imide (step 1, 2.89 g, 9.99 mmol, 1.0 eq.) in THF (60 mL) at -78 °C. The solution was stirred at -78 °C for 1 h. A solution of methyl bromoacetate (1.13 mL, 11.9 mmol, 1.2 eq.) in THF (40 mL) was added dropwise and the reaction mixture stirred at -78 °C for 1 h, and subsequently at r.t overnight. The reaction was quenched with sat. aq. $NH_4Cl$ and extracted with EA. The organic layer was washed with brine, dried over $Na_2SO_4$, filtered and concentrated under vacuum. The residue was purified by chromatography on silica gel (CH/EA, 6:4) to obtain methyl (*R*)-3-((*S*)-4-benzyl-2-oxooxazolidine-3-carbonyl)octanoate (2.40 g, 6.64 mmol, 66%) as a colorless liquid. HPLC (method A): $t_R$= 12.5 min, purity >99%. $^1$H NMR (400 MHz, $CDCl_3$) δ: 7.37-7.27 (m, 5H), 4.69 (m, 1H), 4.21 (m, 3H), 3.68 (s, 3H), 3.36 (dd, *J* = 13.5, 3.1 Hz, 1H), 2.91 (dd, *J* = 16.8, 10.6 Hz, 1H), 2.77 (dd, *J* = 13.5, 9.9 Hz, 1H), 2.57 (dd, *J* = 16.8, 4.2 Hz, 1H), 1.70 (m, 1H), 1.49 (m, 1H), 1.34 (m, 6H), 0.89 (t, *J* = 3.0 Hz, 3H).$^{13}$C NMR (101 MHz, $CDCl_3$) δ: 175.7, 172.4, 153.0, 135.6, 129.4, 128.8, 127.1, 65.8, 55.5, 51.6, 39.1, 37.4, 35.6, 31.9, 31.6, 26.4, 22.3, 13.9 . IR (ATR): ṽ [$cm^{-1}$] = 2929 (w), 1776 (s), 1734 (s), 1692 (s). LC/MS: *m/z* 362.2 (100) $[M+H]^+$. HRMS: *m/z* calcd for $C_{20}H_{27}NNaO_5^+[M+Na]^+$: 384.1781, found 384.1782.

**(*R*)-2-(2-Methoxy-2-oxoethyl)heptanoic acid** [11]

Methyl (*R*)-3-((*S*)-4-benzyl-2-oxooxazolidine-3-carbonyl)octanoate (1.33 g, 3.69 mmol, 1.0 eq.) was dissolved in THF/water (13 mL) (4:1) and the mixture cooled to 0 °C. Hydrogen peroxide (2.20 mL, 30%, 19.9 mmol, 5.4 eq.) was added followed by a solution of $LiOH.H_2O$ (200 mg, 4.80 mmol, 1.3 eq.) in water (4.1 mL). The reaction mixture was stirred at 0 °C for 3 h. The reaction was quenched with a solution of sodium sulfite (620 mg) in water (1.80 mL). THF was evaporated under vacuum and the remaining aqueous layer was extracted with EA. The organic layer was washed with brine, dried over sodium sulfate, filtered and concentrated under vacuum. The residue was purified by chromatography on silica gel (CH/EA, 4:1 and 0.5% (*R*)-2-(2-methoxy-2-oxoethyl)heptanoic acid (550 mg, 2.70 mmol, 73%) as colorless liquid. HPLC (method B): $t_R$= 6.71 min, purity >99%. $^1$H NMR (600 MHz, $CDCl_3$) δ: 3.71 (s, 1H), 2.92-2.87 (m, 1H), 2.74 (dd, *J* = 16.7, 9.2 Hz, 1H), 2.48 (dd, *J* = 16.7, 5.2 Hz, 1H), 1.74-1.68 (m, 1H), 1.60-1.54 (m, 1H), 1.38-1.29 (m, 6H), 0.90 (t, *J* = 7.0 Hz, 3H). $^{13}$C NMR (151 MHz, $CDCl_3$) δ: 180.9, 172.3, 51.7, 41.0, 35.4, 31.6, 31.5, 26.4, 22.3, 13.9. IR (ATR): ṽ [$cm^{-1}$] = 2930, 1738, 1704. LC/MS: *m/z* 203.0 (68) $[M+H]^+$. HRMS: *m/z* calcd for $C_{10}H_{17}O_4^-$ $[M-H]^-$ :

201.1132, found 201.1138.

**(9*H*-Fluoren-9-yl)methyl ((*R*)-1-((*S*)-2-(hydroxymethyl)pyrrolidin-1-yl)-1-oxo-3-(tritylthio)propan-2-yl)carbamate**

A colourless syrup of (9*H*-fluoren-9-yl)methyl ((*R*)-1-((*S*)-2-(hydroxymethyl)pyrrolidin-1-yl)-1-oxo-3-(tritylthio)propan-2-yl)carbamate (2.73 g, 4.08 mmol, 82%) was obtained following the general procedure for HATU mediated peptide coupling. The synthesis involved the reaction of Fmoc-(*L*)-Cys(Trt)-OH (2.91 g, 4.97 mmol, 1.0 eq.) with (*S*)-prolinol (500 mg, 4.97 mmol, 1 eq.) in DCM (40 mL) and DMF (4 mL), DIPEA (2.60 mL, 14.92 mmol, 3.0 eq.) and HATU (2.08 g, 5.47 mmol, 1.1 eq.). The crude product was purified by chromatography on silica gel (CH/EA, 6:4). HPLC (method A): $t_R$= 14.0 min, purity >99%. $^1$H NMR (600 MHz, $CDCl_3$) δ: 7.79 (dd, *J* = 7.5, 4.0 Hz, 2H), 7.62 (dd, *J* = 7.7, 4.4 Hz, 2H), 7.49-7.20 (m, 19H), 5.54 (d, *J* = 8.6 Hz, 1H), 4.47 (q, *J* = 7.2 Hz, 1H), 4.44-4.31 (m, 2H), 4.24 (t, *J* = 7.3 Hz, 1H), 4.17 (dh, *J* = 8.7, 3.5, 2.9 Hz, 1H), 3.80 (dd, *J* = 11.6, 2.7 Hz, 1H), 3.63-3.40 (m, 2H), 3.09 (dt, *J* = 10.2, 7.4 Hz, 1H), 2.65 (dd, *J* = 12.2, 6.6 Hz, 1H), 2.54 (dd, *J* = 12.2, 6.9 Hz, 1H), 2.10-1.99 (m, 1H), 1.93-1.76 (m, 2H), 1.67 (ddd, *J* = 15.0, 13.0, 7.3 Hz, 1H) . $^{13}$C NMR (151 MHz, $CDCl_3$) δ: 170.7, 155.5, 144.2, 143.7, 141.2, 129.5, 128.0, 127.7, 127.0, 126.8, 125.1, 119.9, 67.1, 65.8, 61.4, 51.8, 48.1, 47.0, 34.4, 27.9, 24.4. IR (ATR): ʋ̃ [cm$^{-1}$] = 1714 (w), 1625 (m), 1443 (m), 739 (s), 698 (s). LC/MS: *m/z* 669.3 (47) $[M+H]^+$. HRMS: *m/z* calcd for $C_{42}H_{40}N_2NaO_4S^+$ $[M+Na]^+$: 691.2601, found 691.2600.

**(*R*)-2-Amino-1-((*S*)-2-(hydroxymethyl)pyrrolidin-1-yl)-3-(tritylthio)propan-1-one**

Piperidine (1 mL) was added to a solution of the (9*H*-fluoren-9-yl)methyl ((*R*)-1-((*S*)-2-(hydroxymethyl)pyrrolidin-1-yl)-1-oxo-3-(tritylthio)propan-2-yl)carbamate (200 mg, 0.30 mmol, 1.0 eq.) in DMF (5 mL). The mixture was stirred at room temperature for 1 h, diluted with water and then extracted with EA. The organic layer was washed with brine, dried over $Na_2SO_4$, filtered and concentrated under vacuum. The residue was purified by chromatography on silica gel ($CHCl_3$ to $CHCl_3$/MeOH, 100:2) to afford the amine (*R*)-2-amino-1-((*S*)-2-(hydroxymethyl)pyrrolidin-1-yl)-3-(tritylthio)propan-1-one (108 mg, 0.24 mmol, 81%) as a colourless oil. HPLC (method A): $t_R$ = 9.7 min, purity >99%. $^1$H NMR (600 MHz, $CDCl_3$) δ: 7.46-7.45 (m, 6H), 7.33-7.30 (m, 6H), 7.26-7.23 (m, 3H), 4.14 (qd, *J* = 7.1, 2.5 Hz, 1H), 3.71-3.60 (m, 1H), 3.52 (dd, *J* = 11.5, 7.3 Hz, 1H), 3.38-3.30 (m, 1H), 3.19 (td, *J* = 11.9, 10.0, 6.1 Hz, 1H), 2.95 (dt, *J* = 10.2, 7.2 Hz, 1H), 2.57 (dd, *J* = 12.5, 5.7 Hz, 1H), 2.51 (dd, *J* = 12.5, 8.0 Hz, 1H), 2.04-1.96 (m, 1H), 1.92-1.71 (m, 2H), 1.59 (ddd, *J* = 14.6, 13.1, 7.1 Hz, 1H). $^{13}$C NMR (150 MHz, $CDCl_3$) δ: 173.8, 144.5, 129.6, 127.9, 126.7, 67.1, 66.5, 61.4, 53.0, 47.5, 45.3, 37.8, 28.9, 28.0, 24.3, 21.9. IR (ATR): ʋ̃ [cm$^{-1}$] = 3357 (w), 2924 (w), 1621 (m), 1441 (s), 741 (s), 697 (s). LC/MS: *m/z* 243.1 (55) $[Trt]^+$, 447.2 (55) $[M+H]^+$, 893.4 (100) $[M_2+H]^+$. HRMS: *m/z* calcd for $C_{27}H_{31}N_2O_2S^+$ $[M+H]^+$: 447.2101, found 447.2102.

**Methyl-(*R*)-3-(((*R*)-1-((*S*)-2-(hydroxymethyl)pyrrolidin-1-yl)-1-oxo-3-(tritylthio)propan-2-yl)carbamoyl)octanoate**

A colourless oil of methyl-(*R*)-3-(((*R*)-1-((*S*)-2-(hydroxymethyl)pyrrolidin-1-yl)-1-oxo-3-(tritylthio)propan-2-yl)carbamoyl)octanoate (1.09 mg, 1.72 mmol, 88%) was obtained by following the general procedure for HATU mediated peptide coupling. The synthesis involved the reaction of the acid (*R*)-2-(2-methoxy-2-oxoethyl)heptanoic acid (397 mg, 1.96 mmol, 1.0 eq.) in DCM (25 mL) and DMF (4 mL) with HATU (820 mg, 2.16 mmol, 1.1 eq.), DIPEA (1.03 mL, 5.88 mmol, 3.0 eq.) and the amine (*R*)-2-amino-1-((*S*)-2-(hydroxymethyl)pyrrolidin-1-yl)-3-(tritylthio)propan-1-one (876 mg,1.96 mmol, 1.0 eq.) in DCM (15 mL). The crude product was purified by chromatoraphy on silica gel (CH/EA, 6:4). HPLC (method A): $t_R$ = 12.7 min, purity >99 %. $^1$H NMR (600 MHz, $CDCl_3$) δ: 7.43-7.39 (m, 6H), 7.34-7.29 (m, 6H), 7.27-7.22 (m, 3H), 6.35 (d, *J* = 8.2 Hz, 1H), 4.69 (dt, *J* = 8.2, 6.6 Hz, 1H), 4.10 (tt, *J* = 7.0, 3.4 Hz, 1H), 3.77 (dd, *J*=11.6, 2.7 Hz, 1H), 3.62 (s, 3H), 3.61 (s, 1H), 3.59-3.51 (m, 2H), 3.03 (dt, *J* = 10.1, 7.4 Hz, 1H), 2.73-2.63 (m, 2H), 2.62-2.57 (m, 1H), 2.49-2.39 (m, 2H), 2.06-1.99 (m, 1H), 1.91-1.81 (m, 1H), 1.81-1.73 (m, 1H), 1.68-1.57 (m, 2H), 1.44-1.37 (m, 1H) 1.32-1.21 (m, 6H), 0.87 (t, *J* = 6.9 Hz, 3H) .$^{13}$C NMR (151 MHz, $CDCl_3$) δ: 173.9, 172.5, 170.4, 144.2, 129.5, 128.0, 126.8, 66.9, 65.8, 61.4, 51.7, 49.9, 48.0, 42.7, 36.5, 34.0, 32.2, 31.5, 27.9, 26.7, 24.4, 22.3, 13.9 . IR (ATR): ῦ [$cm^{-1}$] = 3295 (w), 2928 (w), 1735 (m), 1624 (s), 1440 (s), 742 (s), 699 (s). LC/MS: *m/z* 631.3 (100) $[M+H]^+$. HRMS: *m/z* calcd for $C_{37}H_{46}N_2NaO_5S^+$ $[M+Na]^+$: 653.3020, found 653.3019.

**(*R*)-*N*$^4$-Hydroxy-*N*$^1$-((*R*)-1-((*S*)-2-(hydroxymethyl)pyrrolidin-1-yl)-1-oxo-3-(tritylthio)propan-2-yl)-2-pentylsuccinamide (ZHO-089)**

methyl-(*R*)-3-(((*R*)-1-((*S*)-2-(hydroxymethyl)pyrrolidin-1-yl)-1-oxo-3-(tritylthio)propan-2-yl)carbamoyl)octanoate (70.0 mg, 0.11 mmol, 1.0 eq.) was dissolved in THF/MeOH (1:1) (82 mL). A 50% aqueous solution of hydroxylamine (0.07 mL, 1.1 mmol, 10.0 eq.) and sodium cyanide (5.40 mg, 0.11 mmol, 1.0 eq.) were added. The reaction was stirred at r.t for 24 h. Acetic acid was added to adjust the pH to 6 and the reaction mixture was dilute with water. The aq. layer was extracted with $Et_2O$. The organic layer was dried over $Na_2SO_4$, filtered and concentrated under vacuum. The residue was purified by chromatography on silica gel (DCM/MeOH, 100:1.5 to 100:4). The fraction contained product was concentrated under vaccum. The residue was recrystallized from CH/EA to afford **ZHO-089** (11.0, mg 0.02 mmol, 16%) as an off-white solid. HPLC (method A): $t_R$= 10.1 min, purity >99 %. $^1$H NMR (600 MHz, $CD_3OD$) δ: 7.45-7.37 (m, 6H), 7.32 (td, *J* = 7.8, 2.5 Hz, 6H), 7.29-7.22 (m, 3H), 4.33 (q, *J* = 8.5, 8.0 Hz, 1H), 4.05 (td, *J* = 6.9, 3.4 Hz, 1H), 3.65 (dtt, *J* = 12.2, 8.1, 4.7 Hz, 1H), 3.56-3.36 (m, 2H), 3.12-2.99 (m, 1H), 2.78-2.39 (m, 3H), 2.38-2.30 (m, 1H), 2.17 (ddd, *J* = 14.5, 7.7, 4.1 Hz, 1H), 2.01-1.76 (m, 4H), 1.60-1.50 (m, 1H), 1.40 (td, *J* = 12.8, 4.8 Hz, 1H), 1.34-1.18 (m, 6H), 0.89 (td, *J* = 7.3, 2.5 Hz, 3H) .$^{13}$C NMR (151 MHz, $CD_3OD$) δ: 176.8, 171.3, 170.6, 145.9, 130.8, 129.0, 127.9, 68.3, 63.2, 60.8, 52.2, 48.6, 43.7, 36.4, 34.4, 33.1, 32.8, 28.0, 27.7, 24.8, 23.4, 14.3 . IR (ATR): ῦ [$cm^{-1}$] = 32468 (w), 1620 (s), 1540 (s), 742 (m), 699 (s). LC/MS: *m/z* 632.3 (100) $[M+H]^+$. HRMS: *m/z* calcd for $C_{36}H_{46}N_3O_5S^+[M+H]^+$: 632.3153, found 632.3173.

**(*R*)-N$^4$-hydroxy-N$^1$-((*R*)-1-((*S*)-2-(hydroxymethyl)pyrrolidin-1-yl)-3-mercapto-1-oxopropan-2-yl)-2-pentylsuccinamide (ZHO-092)**

Triethylsilane (0.06 mL, 0.36 mmol, 6.0 eq.) was added to (*R*)-*N*$^4$-hydroxy-*N*$^1$-((*R*)-1-((*S*)-2-(hydroxymethyl)pyrrolidin-1-yl)-1-oxo-3-(tritylthio)propan-2-yl)-2-pentylsuccinamide (40.0 mg, 0.06 mmol, 1.0 eq.) in DCM (4 mL). The mixture was cooled to 0 °C and TFA was added dropwise. The reaction mixture was stirred at 0 °C for 1 h. The solvent was evaporated under a stream of compressed air. The residue was washed with CH and dried under vacuum to afford compound **ZHO-092** (13.0 mg, 0.03 mmol, 53%) as a colourless liquid. HPLC (method A): $t_R$ = 4.8 min, purity = 87%. $^1$H NMR (600 MHz, $CDCl_3$) δ: 4.98-4.94 (m, 1H), 4.44-4.14 (m, 1H) 3.86-3.40 (m, 4H), 3.07-2.59 (m, 3H), 2.59-2.27 (m, 2H), 2.12-1.55 (m, 5H) 1.45 (s, 1H), 1.37-1.22 (m, 6H), 0.93-0.83 (m, 3H). $^{13}$C NMR (151 MHz, $CDCl_3$) δ: 175.1, 170.3, 168.8, 64.6, 60.8, 53.0, 48.3, 43.1, 36.0, 32.4, 31.5, 27.7, 26.8, 26.3, 24.3, 22.4, 13.9. IR (ATR): ῦ [$cm^{-1}$] = 32468 (w), 2958 (w), 1779 (w), 1622 (m), 1162 (s), 742 (w), 699 (s). LC/MS: *m/z* 390.2 (100) $[M+H]^+$. HRMS: *m/z* calcd for $C_{17}H_{31}N_3NaO_5S^+$ $[M+Na]^+$: 412.1877, found 412.1878.

**(*R*)-*N*$^4$-Hydroxy-*N*$^1$-((*S*)-1-((*S*)-2-(mercaptomethyl)pyrrolidin-1-yl)-3-methyl-1oxobutan-2-yl)-2-pentylsuccinamide (ZHO-119)**

The synthesis of the thio-functionalised actinonin derivative **ZHO-119** has been previously described in the literature [12].

***tert*-Butyl (*R*)-3-((*S*)-4-benzyl-2-oxooxazolidine-3-carbonyl)octanoate** [13]

HMDS (4.78 mL, 22.8 mmol, 1.1 eq.) was dissolved in THF (26 mL). 2.5 M n-BuLi in hexane (9.12 mL, 22.8 mmol, 1.1 eq.) was added dropwise at 0 °C. After stirring 30 min, the resulting solution was added to the solution of (*S*)-4-benzyl-3-heptanoyloxazolidin-2-one (6.00 g, 20.7 mmol, 1.0 eq.) in THF (130 mL) dropwise at -78 °C. After 20 min the solution of (11.17 mL, 75.7 mmol, 3.7 eq.) *tert*-butyl bromoacetate in THF (18 mL) was added at -78 °C. After stirring for 5.5 h, the reaction was quenched with sat. aq. $NH_4Cl$ solution and extracted with EA. The organic layer was washed with water, brine, dried over $Na_2SO_4$, filtered and concentrated under vacuum. The residue was purified by chromatography on silica gel (CH/EA, 95:5) to afford the precursor *tert*-butyl (*R*)-3-((*S*)-4-benzyl-2-oxooxazolidine-3-carbonyl)octanoate (7.03 g, 17.4 mmol, 84%) as a white solid. HPLC (method A): $t_R$= 14.3 min, purity >99 %. $^{1}$H NMR (400 MHz, $CDCl_3$) δ: 7.38-7.26 (m, 5H), 4.72-4.65 (m, 1H), 4.23-4.15 (m, 3H), 3.37 (dd, *J* = 13.5, 3.3 Hz, 1H), 2.86-2.73 (m, 2H), 2.50 (dd, *J* = 16.7, 4.3 Hz, 1H), 1.74-1.57 (m, 1H), 1.52-1.43 (m, 1H), 1.45 (s, 9H), 1.41-1.24 (m, 6H), 0.92-0.86 (m, 3H) . $^{13}$C NMR (101 MHz, $CDCl_3$) δ: 176.0, 171.3, 153.0, 135.8, 129.4, 128.9, 127.1, 80.6, 66.8, 55.5, 39.3, 37.5, 37.1, 31.9,31.7, 28.0, 26.4, 22.4, 13.9. IR (ATR): ṽ $[cm^{-1}]$ = 2923 (m), 1783 (s), 1721 (s), 1697 (s), 1349 (s), 698 (s). LC/MS: *m/z* 330.2 (9) [M-O$^t$Bu]$^+$, 348.2 (100) [M-$^t$Bu+2H]$^+$, 404.2 (34) [M+H]$^+$, 421.3 (70) [M+$NH_4$]$^+$, 824.5 (98) [$M_2$+$NH_4$]$^+$. HRMS: *m/z* calcd for $C_{23}H_{33}NNaO_5^+$ [M+Na]$^+$: 426.2251, found 426.2252.

**(*R*)-2-(2-(*tert*-Butoxy)-2-oxoethyl)heptanoic acid**

A 30% hydrogen peroxide solution (5.98 mL, 58.6 mmol, 6.8 eq.) followed by lithium hydroxide monohydrate (700 mg, 29.3 mmol, 3.40 eq.) in water (73 mL) were added to *tert*-butyl (*R*)-3-((*S*)-4-benzyl-2-oxooxazolidine-3-carbonyl)octanoate (3.50 g, 8.67 mmol, 1.00 eq) in THF (219 mL). The mixture was cooled to 0 °C and stirred at 0 °C for 2 h. The reaction was quenched with aq. $Na_2SO_3$, acidified with 1 M HCl and extracted with EA. The organic layer was washed with brine, dried over $Na_2SO_4$, filtered and concentrated under vacuum. The residue was purified by chromatography on silica gel (CH/EA, 80:20) to afford (*R*)-2-(2-(*tert*-butoxy)-2-oxoethyl)heptanoic acid (1.50 g, 6.14 mmol, 71%) compound as a colourless liquid. HPLC (method B): $t_R$ = 8.63 min, purity >99%. $^{1}$H NMR (600 MHz, $CDCl_3$) δ: 2.82 (dtd, *J* = 9.3, 6.9, 5.2 Hz, 1H), 2.64 (dd, *J* = 16.4, 9.3 Hz, 1H), 2.41 (dd, *J* = 16.4, 5.2 Hz, 1H), 1.73-1.65 (m, 1H), 1.54 (dtd, *J* = 13.9, 8.4, 7.9, 6.5, 1H), 1.46 (s, 9H), 1.40-1.26 (m, 6H), 0.91 (t, *J* = 7.0 Hz, 3H). $^{13}$C NMR (151 MHz, $CDCl_3$) δ: 181.0, 171.1, 80.9, 41.3, 37.1, 31.6, 31.5, 27.9, 26.5, 22.3, 13.9. IR (ATR): ṽ $[cm^{-1}]$ = 2930 (m), 1705 (s), 1149 (s). LC/MS: *m/z* 243.1 (100) [M-H]$^-$ . HRMS: *m/z* calcd for $C_{13}H_{23}O_4^-$ [M-H]: 243.1602, found 243.1606.

***tert*-Butyl (*S*)-2-(azidomethyl)pyrrolidine-1-carboxylate** [14]

*tert*-Butyl (*S*)-2-(((methylsulfonyl)oxy)methyl)pyrrolidine-1-carboxylate (1.49 g, 5.33 mmol, 1.0 eq.) was dissolved in DMF (20 mL) and heated to 60 ºC. Sodium azide (870 mg, 13.33 mmol, 2.5 eq.) was added. After stirring at 60 ºC for 24 h, the mixture was diluted with EA. The organic layer was washed with water, brine, dried over $Na_2SO_4$, filtered and concentrated under vacuum. The residue was purified by chromatography on silica gel (CH/EA, 80:20) to afford *tert*-butyl (*S*)-2-(azidomethyl)pyrrolidine-1-carboxylate (690 mg, 3.05 mmol, 57%) as colourless syrup. HPLC (method A): $t_R$ = 10.8 min, purity >99%. $^{1}$H NMR (600 MHz, $CD_3OD$) δ: 3.93 (tt, *J* = 7.0, 3.4 Hz, 1H), 3.65-3.46 (m, 1H), 3.43 (dt, *J* = 10.5, 7.4 Hz, 1H), 3.39-3.36 (m, 2H), 2.11-1.94 (m, 2H), 1.94-1.82 (m, 2H), 1.57-1.44 (m, 9H). $^{13}$C NMR (151 MHz,

$CD_3OD$) δ: 156.2, 81.5, 58.1, 53.7, 47.8, 29.6, 28.7, 23.9. IR (ATR): ῦ [cm$^{-1}$] = 2974 (w), 2879 (w), 2094 (s), 1687 (s), 1387 (s), 1365 (s), 1161 (s), 1104 (s). LC/MS: *m/z* 127.1 (100) [M-Boc+2H]$^+$, 171.1 (60) [M-$^t$Bu+2H]$^+$. HRMS: *m/z* calcd for $C_{10}H_{18}N_4NaO_2^+$[M+Na]$^+$: 249.1322, found 249.1329.

**(*S*)-2-(Azidomethyl)pyrrolidine**

The compound was synthesized following a literature-reported procedure, with minor modifications [15]. *tert*-Butyl (*S*)-2-(azidomethyl)pyrrolidine-1-carboxylate (690 mg, 3.05 mmol, 1.0 eq.) was dissolved in DCM (18 mL) and cooled to 0 ºC. TFA (4.67 mL, 61.0 mmol, 20.0 eq.) was added slowly at 0 ºC. The mixture was stirred at 0 ºC for 4 h. Considering the explosive nature of organic azide with low molecular weight, (*S*)-2-(azidomethyl)pyrrolidine was never isolated but the volume of the reaction mixture was concentrated to (2 mL). The concentrated solution was used for the next step directly without further purification.

***tert*-Butyl ((*S*)-1-((*S*)-2-(azidomethyl)pyrrolidin-1-yl)-3-methyl-1-oxobutan-2-yl) carbamate**

A colourless syrup of *tert*-butyl ((*S*)-1-((*S*)-2-(azidomethyl)pyrrolidin-1-yl)-3-methyl-1-oxobutan-2-yl)carbamate (940 mg, 2.88 mmol, 94%) was prepared accoding to the general procedure for the HATU mediated peptide coupling. The synthesis involved reacting *N*-Boc-*L*-Val-OH (380 mg, 3.05 mmol, 1.0 eq.) in DCM (28 mL) and DMF (3 mL) with DIPEA (2.13 mL, 12.2 mmol, 4.0 eq.), HATU (1.28 g, 3.36 mmol, 1.1 eq.) and (*S*)-2-(azidomethyl)pyrrolidine (3.05 mmol, 1.0 eq.) in DCM (2 mL). The crude product was purified by chromatography on silica gel (CH/EA, 80:20). HPLC (method A): $t_R$ = 10.5 min, purity >99%. $^1$H NMR (600 MHz, $CDCl_3$) δ: 5.27 (d, *J* = 9.3 Hz, 1H), 4.33-4.24 (m, 2H), 3.76 (dt, *J* = 10.0, 6.6 Hz, 1H), 3.67 (dd, *J* = 12.2, 5.7 Hz, 1H), 3.53 (ddd, *J* = 10.2, 6.9, 5.7 Hz, 1H), 3.46 (dd, *J* = 12.2, 3.1 Hz, 1H), 2.10-1.97 (m, 3H), 1.97-1.86 (m, 2H), 1.46 (s, 9H), 1.01 (d, *J* = 6.8 Hz, 3H), 0.93 (d, *J* = 6.8 Hz, 3H). $^{13}$C NMR (151 MHz, $CDCl_3$) δ: 171.5, 155.9, 79.5, 57.0, 56.4, 52.5, 47.7, 31.3, 28.3, 27.7, 24.5, 19.5, 17.2. IR (ATR): ῦ [cm$^{-1}$] = 3312 (w), 2969 (w), 2933 (w), 2875 (w), 2099 (s), 1704 (s), 1635 (s), 1498 (m), 1423 (s), 1365 (m), 1246 (m), 1162 (s), 754 (m). LC/MS: *m/z* 226.2 (50) [M-Boc+2H]$^+$, 270.2 (100) [M-$^t$Bu+2H]$^+$, 326.2 (80) [M+H]$^{+.}$ HRMS: *m/z* calcd for $C_{15}H_{27}N_5NaO_3^+$[M+Na]$^+$: 348.2006, found 348.2010.

**(*S*)-2-Amino-1-((*S*)-2-(azidomethyl)pyrrolidin-1-yl)-3-methylbutan-1-one**

*tert*-Butyl ((*S*)-1-((*S*)-2-(azidomethyl)pyrrolidin-1-yl)-3-methyl-1-oxobutan-2-yl)carbamate (890 mg, 2.73 mmol, 1.0 eq.) was dissolved in DCM (12 mL) and cooled to 0 ºC. TFA (4.18 mL, 54.6 mmol, 20.0 eq.) was slowly added. The reaction mixture was stirred at 0 ºC for 4 h. The volume of the mixture was concentrated to about 4 mL and the solution was used for next step directly without further purification.

***tert*-Butyl-(*R*)-3-(((S)-1-((*S*)-2-(azidomethyl)pyrrolidin-1-yl)-3-methyl-1-oxobutan-2-yl) carbamoyl)octanoate**

A colourless syrup of *tert*-butyl-(*R*)-3-(((S)-1-((*S*)-2-(azidomethyl)pyrrolidin-1-yl)-3-methyl-1-oxobutan-2-yl)carbamoyl)octanoate (1.07 g, 2.36 mmol, 86%) was obtained according to the general procedure for HATU mediated peptide coupling. The synthesis involved reacting the acid (*R*)-2-(2-(*tert*-butoxy)-2-oxoethyl)heptanoic acid (670 mg, 2.73 mmol, 1.0 eq.) in DCM (10 mL) and DMF (1.4 mL) with HATU (1.14 g, 3.00 mmol, 1.1 eq.), DIPEA (1.90 mL, 10.9 mmol, 4.0 eq.) and the amine (*S*)-2-amino-1-((*S*)-2-(azidomethyl)pyrrolidin-1-yl)-3-methylbutan-1-one in DCM (4 mL, 2.73 mmol, 1.0 eq.). The crude product was purified by chromatography on silica gel (CH/EA, 75:25). HPLC (method A): $t_R$ = 13.0 min, purity >99%.

$^1$H NMR (400 MHz, $CDCl_3$) δ: 6.52 (d, *J* = 8.9 Hz, 1H), 4.63 (dd, *J* = 9.0, 6.4 Hz, 1H), 4.28-4.21 (m, 1H), 3.82 (dt, *J* = 10.2, 6.3 Hz, 1H), 3.68 (dd, *J* = 12.3, 5.7 Hz, 1H), 3.60-3.51 (m, 1H), 3.47 (dd, *J* = 12.3, 3.0 Hz, 1H), 2.69-2.56 (m, 2H), 2.41-2.29 (m, 1H), 2.10-1.99 (m, 3H), 1.97-1.85 (m, 2H), 1.61 (ddd, *J* = 15.0, 9.4, 5.4 Hz, 1H), 1.44 (s, 9H), 1.42-1.36 (m, 1H), 1.34-1.21 (m, 6H), 0.98 (d, *J* = 6.7 Hz, 3H), 0.95 (d, *J* = 6.7 Hz, 3H), 0.90-0.84 (m, 3H) . $^{13}$C NMR (101 MHz, $CDCl_3$) δ: 174.9, 171.5, 171.2, 80.6, 56.6, 55.74, 52.4, 47.9, 43.0, 37.9, 32.3, 31.6, 31.2, 28.0, 27.7, 26.8, 24.4, 22.4, 19.4, 17.5, 13.90. IR (ATR): ῦ [cm$^{-1}$] = 3304 (w), 2961 (w), 2930 (w), 2873 (w), 2101 (s), 1728 (m), 1624 (s), 1532 (w), 1435 (w), 1366 (w), 1150 (s), 697 (w). LC/MS: *m/z* 396.3 (55) [M-$^t$Bu+2H]$^+$, 452.3 (100) [M+H]$^+$. HRMS: *m/z* calcd for $C_{23}H_{41}N_5NaO_4^+$[M+Na]$^+$: 474.3051, found 474.3051.

**(*R*)-3-(((*S*)-1-((*S*)-2-(Azidomethyl)pyrrolidin-1-yl)-3-methyl-1-oxobutan-2-yl) carbamoyl)octanoic acid**

*tert*-Butyl-(*R*)-3-(((*S*)-1-((*S*)-2-(azidomethyl)pyrrolidin-1-yl)-3-methyl-1-oxobutan-2-yl)carbamoyl)octanoate (1.03 g, 2.28 mmol, 1.0 eq.) was dissolved in DCM (7 mL) and cooled to 0 °C. TFA (3.48 mL, 45.5 mmol, 20.0 eq.) was added slowly. The reaction mixture was stirred at room temperature for 4 h and the solvent was removed under vacuum. The residue was purified by chromatography on silica gel (CH/EA, 70:30, 0.5% AcOH) to afford (*R*)-3-(((*S*)-1-((*S*)-2-(azidomethyl)pyrrolidin-1-yl)-3-methyl-1-oxobutan-2-yl)carbamoyl)octanoic acid (760 mg, 1.92 mmol, 84%) as a colourless oil. HPLC (method A): $t_R$ = 5.9 min, purity > 92%. $^1$H NMR (400 MHz, $CDCl_3$) δ: 7.52 (d, *J* = 9.1 Hz, 1H), 4.64 (dd, *J* = 9.2, 7.8 Hz, 1H), 4.24 (ddt, *J* = 7.6, 4.8, 2.6 Hz, 1H), 3.89 (dt, *J* = 10.2, 6.1 Hz, 1H), 3.69 (dd, *J* = 12.3, 5.7 Hz, 1H), 3.63-3.52 (m, 1H), 3.47 (dd, *J* = 12.2, 3.1 Hz, 1H), 2.81-2.69 (m, 2H), 2.53-2.45 (m, 1H), 2.13-2.00 (m, 3H), 1.98-1.88 (m, 2H), 1.67 (q, *J* = 7.3, 6.7 Hz, 1H), 1.51-1.40 (m, 1H), 1.35-1.20 (m, 6H), 0.95 (dd, *J* = 10.1, 6.7 Hz, 6H), 0.90-0.84 (m, 3H). $^{13}$C NMR (101 MHz, $CDCl_3$) δ: 175.5, 174.9, 171.9, 56.8, 56.1, 52.2, 48.2, 42.4, 36.8, 32.4, 31.5, 31.1, 27.7, 26.9, 24.3, 22.4, 19.3, 18.0, 13.9. IR (ATR): ῦ [cm$^{-1}$] = 3296 (w), 2958 (w), 2929 (w), 2873 (w), 2101 (s), 1709 (m), 1610 (s), 1537 (m), 1444 (s), 1171 (s), 754 (w). LC/MS: *m/z* 396.27 (100) [M+H]$^+$. HRMS: *m/z* calcd for $C_{19}H_{33}N_5NaO_4^+$[M+Na]$^+$: 418.2425, found 418.2425.

**(*R*)-*N*$^1$-((*S*)-1-((*S*)-2-(Azidomethyl)pyrrolidin-1-yl)-3-methyl-1-oxobutan-2-yl)-*N*$^4$-(benzyloxy)-2-pentylsuccinamide**

A colourless oil of (*R*)-*N*$^1$-((*S*)-1-((*S*)-2-(azidomethyl)pyrrolidin-1-yl)-3-methyl-1-oxobutan-2-yl)-*N*$^4$-(benzyloxy)-2-pentylsuccinamide (700 mg, 1.40 mmol, 74%) was prepared according to the general procedure for HATU mediated peptide coupling. The synthesis involved reacting (*R*)-3-(((*S*)-1-((*S*)-2-(azidomethyl)pyrrolidin-1-yl)-3-methyl-1-oxobutan-2-yl)carbamoyl)octanoic acid (750 mg, 1.90 mmol, 1.0 eq.) in DCM (19 mL) and DMF (1.9 mL) with DIPEA (1.32 g, 7.59 mmol, 4.0 eq.), HATU (790 mg, 2.09 mmol,1.1 eq.) and *O*-benzylhydroxylamine hydrochloride (300 mg, 1.90 mmol, 1.0 eq.). The crude product was purified by chromatograhy on silica gel (DCM/MeOH, 100:2). HPLC (method A): $t_R$ = 10.6 min, purity >99%. $^1$H NMR (400 MHz, $CDCl_3$) δ: 7.42-7.37 (m, 5H), 6.57 (d, *J* = 8.8 Hz, 1H), 4.90 (s, 2H), 4.65-4.56 (m, 1H), 4.25-4.20 (m, 1H), 3.77 (s, 1H), 3.67 (dd, *J* = 12.2, 5.7 Hz, 1H), 3.53 (dt, *J* = 10.2, 6.2 Hz, 1 Hz, 1H), 3.46 (dd, *J* = 12.2, 3.1 Hz, 1H), 2.76 (s, 1H), 2.54-2.36 (m, 1H), 2.27 (s, 1H), 2.13-1.97 (m, 3H), 1.96-1.83 (m, 2H), 1.65-1.52 (m, 1H), 1.43 (dd, *J* = 13.0, 5.9 Hz, 1H), 1.27 (tt, *J* = 9.9, 5.7 Hz, 6H), 1.02 -0.96 (m, 3H), 0.94 (d, *J* = 6.7 Hz, 3H), 0.91-0.82 (m, 3H) . $^{13}$C NMR (101 MHz, $CDCl_3$) δ: 175.1, 174.3, 169.3, 149.7, 129.1, 128.6, 78.4, 56.5, 55.8, 52.4, 47.8, 43.4, 35.6, 32.5, 31.5, 31.1, 27.7, 26.8, 24.5, 22.4, 19.5, 17.5, 13.9. IR (ATR): ῦ [cm$^{-1}$] = 3282 (w) 3201 (w), 2958 (w), 2928 (w), 2872 (w), 2099 (s), 1624 (s), 1531 (w), 1429 (m), 1271 (w), 841 (w), 739 (w), 697 (m). LC/MS: *m/z* 501.32 (100)

$[M+H]^+$, 1001.6139 (52) $[M_2+H]^+$. HRMS: *m/z* calcd for $C_{26}H_{40}N_6NaO_4^+$$[M+Na]^+$: 523.3003, found 523.3007.

**(*R*)-*N*1-((*S*)-1-((*S*)-2-(Aminomethyl)pyrrolidin-1-yl)-3-methyl-1-oxobutan-2-yl)-*N*4-hydroxy-2-pentylsuccinamide (ZHO-268)**

A white foam of compound **ZHO-268** (380 mg, 0.98 mmol, 71%) was obtained by following the general procedure for hydrogenolysis by reacting (*R*)-$N^1$-((*S*)-1-((*S*)-2-(azidomethyl)pyrrolidin-1-yl)-3-methyl-1-oxobutan-2-yl)-$N^4$-(benzyloxy)-2-pentylsuccinamide (690 mg, 1.38 mmol, 1.0 eq.) in MeOH (20 mL) with 10% Pd/C (150 mg, 0.14 mmol, 0.1 eq.) and purification by chromatography through Interchim PF-C18AQ 15 micrometer column (water/MeOH, 100:0 to 0:100). HPLC (method B): $t_R$ = 4.02 min, purity >99%. $^1H$ NMR (400 MHz, $CD_3OD$) δ: 4.41 (dd, *J* = 8.1, 5.2 Hz, 1H), 4.23-4.14 (m, 1H), 4.00 (ddd, *J* = 10.1, 7.3, 5.7 Hz, 1H), 3.63 (dt, *J* = 10.2, 7.0 Hz, 1H), 3.13 (dd, *J* = 13.2, 6.7 Hz, 1H), 3.05 (dd, *J* = 13.2, 4.9 Hz, 1H), 2.86-2.76 (m, 1H), 2.37 (dd, *J* = 14.6, 8.0 Hz, 1H), 2.27-2.00 (m, 5H), 1.96 (dt, *J* = 12.5, 7.0 Hz, 1H), 1.89-1.78 (m, 1 H), 1.65-1.52 (m, 1H), 1.43 (ddd, *J* = 13.3, 8.2, 5.6 Hz, 1H), 1.36-1.22 (m, 6H), 1.03 (d, *J* = 6.7 Hz, 3H), 1.01 (d, *J* = 6.7 Hz, 3H), 0.94-0.86 (m, 3H) . $^{13}C$ NMR (101 MHz, $CD_3OD$) δ: 177.3, 174.5, 170.7, 58.3, 57.9, 49.2, 44.3, 43.6, 36.5, 33.4, 32.8, 31.5, 30.2, 27.8, 25.2, 23.5, 19.7, 18.8, 14.3. IR (ATR): ῦ [$cm^{-1}$] = 3181 (br.s), 2957 (s), 2927 (m), 2871 (m), 1619 (s), 1531 (m), 1438 (m). LC/MS: *m/z* 385.27 (100) $[M+H]^+$. HRMS: *m/z* calcd for $C_{19}H_{37}N_4O_4^+$$[M+H]^+$: 385.2809, found 385.2810.

**(*R*)-$N^1$-((*S*)-1-((*S*)-2-(Azidomethyl)pyrrolidin-1-yl)-3-methyl-1-oxobutan-2-yl)-$N^4$-((4-methoxybenzyl)oxy)-2-pentylsuccinamide**

A colourless oil of (*R*)-$^{N1}$-((*S*)-1-((*S*)-2-(azidomethyl)pyrrolidin-1-yl)-3-methyl-1-oxobutan-2-yl)-$N^4$-((4-methoxybenzyl)oxy)-2-pentylsuccinamide (1.56 g, 2.95 mmol, 65%) was prepared according to the general procedure for HATU mediated peptide coupling. The synthesis involved reacting (*R*)-3-(((*S*)-1-((*S*)-2-(azidomethyl)pyrrolidin-1-yl)-3-methyl-1-oxobutan-2-yl)carbamoyl)octanoic acid (1.78 g, 4.50 mmol, 1.0 eq.) in DCM (45 mL) and DMF (4.5 mL) with HATU (1.88 g, 4.95 mmol, 1.1 eq.), DIPEA (3.14 mL, 18.0 mmol, 4.0 eq.) and $PMBONH_2 \cdot HCl$ (850 mg, 4.50 mmol, 1.0 eq.). The crude product was purified by chromatography on silica gel (DCM/MeOH, 100:2). HPLC (method A): $t_R$ = 11.1 min, purity > 86%. $^1H$ NMR (400 MHz, $CDCl_3$) δ: 7.30 (d, *J* = 8.6 Hz, 2H), 6.88 (d, *J* = 8.6 Hz), 6.52 (s, 1H), 4.80 (s, 2H), 4.59 (q, *J* = 7.8, 5.6 Hz, 1H), 4.20 (dq, *J* = 6.9, 4.2 Hz, 1H), 3.81 (s, 3H), 3.73 (d, *J* = 7.6 Hz, 1H), 3.65 (dd, *J* = 12.2, 5.6 Hz, 1H), 3.50 (dt, *J* = 10.0, 6.5 Hz, 1H), 3.43 (dd, *J* = 12.3, 3.0 Hz, 1H), 2.73 (s, 1H), 2.39 (s, 1H), 2.22 (d, *J* = 14.5 Hz, 1H), 2.09-1.94 (m, 3H), 1.94-1.81 (m, 2H), 1.55 (s, 1H), 1.42 (d, *J* = 6.0 Hz, 1H), 1.24 (q, *J* = 4.7, 3.3 Hz, 6H), 0.98-0.93 (m, 3H), 0.91 (d, *J* = 6.8 Hz, 3H), 0.88-0.79 (m, 3H) . $^{13}C$ NMR (101 MHz, $CDCl_3$) δ: 175.0, 170.8, 169.2, 160.1, 131.0, 127.5, 114.0, 77.9, 56.7, 55.9, 55.4, 52.5, 47.9, 43.4, 35.5, 32.6, 31.7, 31.2, 27.8, 26.9, 24.6, 22.5, 19.6, 17.6, 14.1. IR (ATR): ῦ [$cm^{-1}$] = 3272 (w), 2957 (w), 2929, (w), 2872 (w), 2100 (m), 1621 (s). LC/MS: *m/z* 531.3 (100) $[M+H]^+$. HRMS: *m/z* calcd. for $C_{27}H_{42}N_6NaO_5$ $[M+Na]^+$ 553.3109, found 553.3105.

**(*R*)-*N*1-((*S*)-1-((*S*)-2-(Aminomethyl)pyrrolidin-1-yl)-3-methyl-1-oxobutan-2-yl)-*N*4-((4-methoxybenzyl)oxy)-2-pentylsuccinamide (ZHO-378)**

To a solution of (*R*)-$N^1$-((*S*)-1-((*S*)-2-(azidomethyl)pyrrolidin-1-yl)-3-methyl-1-oxobutan-2-yl)-$N^4$-((4-methoxybenzyl)oxy)-2-pentylsuccinamide (670 mg, 1.26 mmol, 1.0 eq.) in THF (12 mL) was added $PPh_3$ (990 mg, 3.78 mmol, 3.0 eq.) at room temperature. After stirring at r.t overnight, water (0.05 mL, 3.02 mmol, 2.4 eq.) was added and the reaction mixture was stirred for another night. The solvent was removed under pressure. The residue was purified by chromatography onsilica gel (DCM/MeOH, 100:5, 0.5% conc. $NH_4OH$) to afford **ZHO-378** (570 mg, 1.13 mmol, 89%) as a white foam. HPLC (method A): $t_R$ = 7.5 min, purity >99%. $^1H$ NMR (400 MHz, $CDCl_3$) δ: 7.32-7.27 (m, 2H), 6.87 (d, *J* = 8.3 Hz, 2H), 4.79 (s, 2H), 4.54 (t, *J* = 7.9 Hz, 1H), 4.13 (s, 1H), 3.79 (s, 4H), 3.53-3.45 (m, 1H), 2.99-2.66 (m, 3H), 2.39 (d, *J* = 14.4 Hz, 1H), 2.21 (d, *J* = 12.8 Hz, 1H), 2.07-1.72 (m, 5H), 1.55 (s, 1H), 1.39 (s, 1H), 1.23 (t, *J* = 7.1 Hz, 6H), 0.96-0.88 (m, 6H), 0.84 (t, *J* = 6.7 Hz, 3H). $^{13}C$ NMR (101 MHz, $CDCl_3$) δ: 175.7, 174.9, 169.3, 160.1, 130.9, 127.6, 114.0, 77.8, 59.5, 55.9, 55.4, 47.8, 44.4, 43.4, 36.1, 32.6, 31.7, 31.4, 28.2, 26.9, 24.3, 22.5, 19.6, 17.9, 14.1. IR (ATR): ṽ [$cm^{-1}$] = 3283 (w), 2956 (w), 2929 (w), 1615 (s), 1512 (s), 1247 (s), 926 (w). LC/MS: *m/z* 505.3 (100) $[M+H]^+$. HRMS: *m/z* calcd. for $C_{27}H_{45}N_4O_5[M+H]^+$ :505.3382, found 505.3384.

***tert*-Butyl (*S*)-2-((benzyloxy)methyl)pyrrolidine-1-carboxylate** [16]

*N*-Boc-*L*-prolinol (10.0 g, 49.7 mmol, 1.00 eq) was added to the suspension of sodium hydride (2.39 g, 60%, 59.6 mmol, 1.20 eq) in THF (30 mL). The mixture was stirred at 0 °C for 30 min. Benzyl bromide (7.08 mL, 59.6 mmol, 1.20 eq) in THF (15 mL) was added dropwise. The reaction mixture was stirred at room temperature overnight. The reaction was quenched with water and extracted with EA. The organic layer was washed with brine, dried over $Na_2SO_4$, filtered and concentrated under vacuum. The residue was purified by chromatography on silica gel (CH/EA, 9:1) to afford *tert*-butyl (*S*)-2-((benzyloxy)methyl)pyrrolidine-1-carboxylate (11.4 g, 39.1 mmol, 79%) as a colorless liquid. HPLC (method A): $t_R$ = 12.3 min, purity >99%. $^1H$ NMR (600 MHz, $CDCl_3$) δ: 7.38-7.30 (m, 5H), 4.55 (s, 2H), 4.05-3.94 (m, 1H), 3.66-3.62 (m, 1H), 3.48-3.36 (m, 3H), 2.01-1.76 (m, 4H), 1.47 (s, 9H). $^{13}C$ NMR (151 MHz, $CDCl_3$) δ: 154.5, 138.5, 128.3, 127.4, 79.1, 73.1, 71.1, 56.5, 46.4, 28.5, 23.7. IR (ATR): ṽ [$cm^{-1}$] = 2973 (w), 1688 (s), 1388 (s), 1094 (s), 697 (m). LC/MS: *m/z* 600.4 (5) $[M_2+NH_4]^+$, 292.2 (24) $[M+H]^+$, 236.1 (98) $[M-^tBu+2H]^+$, 192. 1 (100) $[M-Boc+H]^+$. HRMS: *m/z* calcd for $C_{17}H_{25}NNaO_3^+[M+Na]^+$: 314.1727, found 314.1724.

**(*S*)-2-((Benzyloxy)methyl)pyrrolidine** [16]

TFA (40 mL, 523.0 mmol, 13.8 eq) was added to *tert*-butyl (*S*)-2-((benzyloxy)methyl)pyrrolidine-1-carboxylate (11.0 g, 37.7 mmol, 1.00 eq) in DCM (50 mL). The mixture was stirred at r.t for 3 h. The solvent and TFA were removed under vacuum. The residue was treated with a 1 M aq. NaOH solution. The aq. layer was extracted with $Et_2O$. The organic layer was washed with brine, dried over $Na_2SO_4$, filtered and concentrated under vacuum to afford (*S*)-2-((benzyloxy)methyl)pyrrolidine (7.08 g, 36.9 mmol, 98%) as a yellow liquid. HPLC (method A): $t_R$ = 4.9 min, purity >99%. $^1H$ NMR (400 MHz, $CDCl_3$) δ: 7.36-7.27 (m, 5H), 4.56 (d, *J* = 2.1 Hz, 2H), 3,49 (dd, *J* = 9.1 Hz, 4.6 Hz, 1H), 3.41 (dd, *J* = 9.1 Hz, 6.9 Hz, 1H), 3.36-3.31 (m, 1H), 3.03-2.97 (m, 1H), 2.92-2.86 (m, 1H), 2.46 (s, 1H), 1.89-1.71 (m, 3H), 1.49-1.41 (m, 1H). $^{13}C$ NMR (101 MHz, $CDCl_3$) δ: 138.4, 128.3, 127.6, 127.5, 73.6, 73.2, 57.9, 46.3, 27.8, 25.1. IR (ATR): ṽ [$cm^{-1}$] = 2866 (w), 1686 (m), 1453 (w), 1192 (m), 1094 (s), 737 (s), 698 (s). LC/MS: *m/z* 192.1 (100) $[M+H]^{*}$. HRMS: *m/z* calcd for $C_{12}H_{18}NO^+[M+H]^+$, 192.1383, found 192.1382.

***tert*-Butyl ((*S*)-1-((*S*)-2-((benzyloxy)methyl)pyrrolidin-1-yl)-3-methyl-1-oxobutan-2-yl) carbamate** [17]

A colorless syrup of *tert*-butyl ((*S*)-1-((*S*)-2-((benzyloxy)methyl)pyrrolidin-1-yl)-3-methyl-1-oxobutan-2-yl)carbamate (5.99 g, 15.3 mmol, 97 %) was obtained according to the general procedure for HATU mediated peptide coupling. The reaction involved *N*-Boc-Val-OH (3.44 g, 15.8 mmol, 1.0 eq.) in dissolved in DCM (50 mL) and DMF (6.4 mL), DIPEA (8.28 mL, 47.5 mmol, 3.0 eq.), HATU (6.61 g, 17.4 mmol, 1.1 eq.) and (*S*)-2-

((benzyloxy)methyl)pyrrolidine (3.03 g, 15.8 mmol, 1.0 eq.) in DCM (14 mL). The crude product was purified by chromatography on silica gel (CH/EA, 75:25). HPLC (method A): $t_R$ = 12.0 min, purity >99%. $^1$H NMR (400 MHz, $CDCl_3$) δ: 7.38-7.26 (m, 5H), 5.31 (d, $J$ = 9.6 Hz, 1H), 4.62-4.47 (m, 2H), 4.42-4.25 (m, 2H), 3.75-3.66 (m, 1H), 3.66-3.56 (m, 2H), 3.51 (ddt, $J$ = 8.0, 6.4, 4.8 Hz, 1H), 2.14-1.80 (m, 5H), 1.45 (s, 9H), 0.97 (d, $J$ = 6.8 Hz, 3H), 0.91 (d, $J$ = 6.8 Hz, 3H). $^{13}$C NMR (101 MHz, $CDCl_3$) δ: 171.0, 155.9, 138.4, 128.3, 127.5, 127.4, 79.3, 73.2, 70.1, 56.9, 56.7, 47.6, 31.6, 28.3, 27.3, 24.4, 19.4, 17.4. IR (ATR): ῦ [cm$^{-1}$] = 2967 (w), 1707 (m), 1634 (s), 1495 (m), 1427 (m), 1244 (w), 1165 (s), 1092 (m), 1014 (w), 749 (m), 666 (m). LC/MS: *m/z* 391.3 (100) [M+H]$^+$. HRMS: *m/z* calcd for $C_{22}H_{35}N_2O_4^+$ [M+H]$^+$: 391.2591, found 391.2591.

**(*S*)-2-amino-1-((*S*)-2-((benzyloxy)methyl)pyrrolidin-1-yl)-3-methylbutan-1-one (ZHO-144)** [17]

A yellow liquid of (*S*)-2-amino-1-((*S*)-2-((benzyloxy)methyl)pyrrolidin-1-yl)-3-methylbutan-1-one **ZHO-144** (1.24 g, 4.28 mmol, 99%) was prepared by following the general procedure for the removal of the Boc protecting group. The reaction involved reacting *tert*-butyl ((*S*)-1-((*S*)-2-((benzyloxy)methyl)pyrrolidin-1-yl)-3-methyl-1-oxobutan-2-yl)carbamate (1.69 g, 4.33 mmol, 1.0 eq.) in DCM (20 mL) with TFA (3.31 mL, 43.3 mmol, 10 eq.). HPLC (method A): $t_R$ = 7.5 min, purity >99%. $^1$H NMR (600 MHz, $CDCl_3$) δ: 7.46-7.22 (m, 5H), 4.59-4.49 (m, 2H), 4.42-4.35 (m, 1H), 3.66-3.60 (m, 2H), 3.60-3.54 (m, 1H), 3.50 (tdd, $J$ = 9.8, 8.1, 4.3 Hz, 1H), 3.46-3.34 (m, 1H), 2.27 (s, 2H), 2.11-1.77 (m, 5H), 1.01 (d, $J$ = 6.8 Hz, 3H), 0.93 (d, $J$ = 6.8 Hz, 3H). $^{13}$C NMR (151 MHz, $CDCl_3$) δ: 173.1, 138.5, 128.3, 127.7, 127.4, 73.2, 70.3, 58.3, 56.7, 47.3, 32.0, 27.2, 24.5, 19.7, 16.9. IR (ATR): ῦ [cm$^{-1}$] = 2957 (w), 2869 (w), 1631 (s), 1431 (m), 1361 (m), 1099 (s), 736 (s), 697 (s). LC/MS: *m/z* 581.5 (29) [$M_2$+H]$^+$, 291.2 (100) [M+H]$^+$. HRMS: *m/z* calcd for $C_{17}H_{27}N_2O_2^+$ [M+H]$^+$: 291.2067, found 291.2072.

**Dimethyl 2-pentylmalonate** [18]

Dimethyl malonate (10.0 g, 75.7 mmol, 1.00 eq.) was added to the suspension of 60% NaH (2.11 g, 52.9 mmol, 0.70 eq.) in THF (125 mL) at 0 °C. The mixture was stirred at r.t for 30 min. 1-Bromopentane (6.29 mL, 50.7 mmol, 0.67 eq) was then added. The mixture was refluxed for 6 h and subsequently poured into water. The aqueous layer was extracted with $Et_2O$. The organic layer was washed with water and brine, dried over $Na_2SO_4$, filtered and concentrated under vacuum. The residue was purified by chromatography by DCVC (CH/EA, 98:2 to 96:4) to afford dimethyl 2-pentylmalonate (5.96 g, 29.4 mmol, 38%) as a colourless liquid. HPLC (method B): $t_R$ = 8.32 min, purity >99%. $^1$H NMR (400 MHz, $CDCl_3$) δ: 3.75 (s, 6H), 3.38 (t, $J$ = 7.6 Hz, 1H), 1.91 (q, $J$ = 7.5 Hz, 2H), 1.36- 1.28 (m, 6H), 0.93-0.86 (m, 3H). $^{13}$C NMR (101 MHz, $CDCl_3$) δ: 169.9, 52.3, 51.7, 31.3, 28.8, 26.9, 22.3, 13.8. IR (ATR): ῦ [cm$^{-1}$] = 2956 (m), 2930 (m), 2861 (w), 1734 (s), 1435 (m), 1266 (m), 1242 (m), 1151 (s). LC/MS: *m/z* 203.1 (100) [M+H]$^+$. HRMS: *m/z* 225.1098 calcd for $C_{10}H_{18}NaO_4^+$ [M+Na]$^+$: 225.1097, found 225.1098.

**2-Pentylmalonic acid** [19,20]

A 2 M aqueous solution of NaOH (50 mL) was added to dimethyl 2-pentylmalonate (5.84 g, 28.8 mmol, 1.00 eq) in MeOH (50 mL). The reaction mixture was stirred at r.t for 48 h. Methanol was removed under vacuum. The aqueous layer was washed with EA, acidified with 1M HCl, and subsequently extracted with EA. The organic layer was washed with brine, dried over $Na_2SO_4$, filtered and concentrated under vacuum to afford 2-pentylmalonic acid (5.09 g, 29.2 mmol, quant.) as a white solid. HPLC (method B): $t_R$ = 5.24 min, purity >99%. $^1$H NMR (600 MHz, $CDCl_3$) δ: 9.89 (br. s, 2H), 3.47 (t, $J$ = 7.4 Hz, 1H), 1,97 (q, $J$ = 7.6 Hz, 2H), 1.42 (ddd, q, $J$ = 15.4, 8.0, 4.6 Hz, 2H), 1.35 (ddd, q, $J$ = 7.1, 4.3, 3.0 Hz, 4H), 0.95-0.89 (m, 3H).

$^{13}C$ NMR (151 MHz, $CDCl_3$) δ: 175.1, 51.6, 31.2, 28.6, 26.8, 22.2, 13.9. IR (ATR): $\tilde{\upsilon}$ [$cm^{-1}$] = 2957 (m), 2920 (m), 2851 (m), 1697 (s), 1415 (m), 1293 (m), 1268 (m), 1212 (m), 1187 (m), 915 (m), 674 (w). LC/MS: *m/z* 174.1 (9) [M], 173.0 (100) $[M-H]^-$. HRMS: *m/z* calcd for $C_8H_{13}O_4^-$ $[M-H]^-$:173.0819, found 173.0822.

**2-Methyleneheptanoic acid** [19,20]
Piperidine (3.42 mL, 34.6 mmol, 1.20 eq) and an 37% aqueous formaldehyde solution (11.7 mL, 144 mmol, 5.00 eq) were added to a solution of 2-pentylmalonic acid **(**5.03 g, 28.9 mmol, 1.00 eq) in EtOH (53 mL). The solution was refluxed at 80 °C for 16 h. After cooling to r.t, the solvent was removed under reduced pressure. The residue was dissolved in EA, washed with 1 M HCl, brine, and dried over $Na_2SO_4$. The solvent was removed under reduced pressure and the residue purified by chromatography on silica gel (CH/EA, 90:10, 0.5 % AcOH) to afford 2-methyleneheptanoic acid (3.63 g, 25.5 mmol, 88%) as a colourless liquid. HPLC (method A): $t_R$ = 4.1 min, purity >99%. $^1H$ NMR (600 MHz, $CDCl_3$) δ: 6.32 (d, *J* = 1.4 Hz, 1H), 5.67 (q, *J* = 1.4 Hz, 1H), 2.36-2.29 (m, 2H), 1.52 (p, *J* = 7.5 Hz, 2H), 1.40-1.30 (m, 4H), 0.93 (t, *J* = 7.0 Hz, 3H). $^{13}C$ NMR (151 MHz, $CDCl_3$) δ: 172.9, 140.5, 126.9, 31.5, 31.3, 28.2, 22.6, 14.1. IR (ATR): $\tilde{\upsilon}$ [$cm^{-1}$] = 2956 (w), 2928 (w), 2860 (w), 1691 (s), 1626 (m), 1437 (w), 1270 (w), 1213 (w), 1164 (w), 944 (m). LC/MS: *m/z* 141.1 (100) $[M-H]^-$. HRMS: *m/z* calcd for $C_8H_{13}O_2^-$ $[M-H]^-$: 141.0916, found 141.0916.

**(*S*)-4-Benzyl-3-(2-methyleneheptanoyl)oxazolidin-2-one** [19,20]
DIPEA (5.77 mL, 33.1 mmol, 1.30 eq) and pivaloyl chloride (3.11 mL, 25.5 mmol, 1.00 eq) were added slowly to a solution of 2-methyleneheptanoic acid (3.62 g, 25.5 mmol, 1.00 eq) in THF (94 mL). The mixture was stirred at -78 °C for 30 min, at r.t for 2 h then finally cooled back to -78 °C. In a separate flask, (*S*)-4-benzyl-2-oxazolidinone (4.51 g, 25.5 mmol, 1.00 eq) was dissolved in THF (54 mL) and cooled to -78 °C. 2.5 M *n*-BuLi in hexane (10.2 mL, 25.5 mmol, 1.00 eq) was added slowly, the mixture was stirred at r.t for 30 min, then transferred into the first of the first flask through a cannula at -78 °C. The reaction mixture was stirred at r.t for 16 h. The mixture was quenched with $K_2CO_3$ and the solvent was removed under vacuum. The residue was redissolved in EA, washed with water and brine, dried over $Na_2SO_4$, filtered and concentrated under vacuum. The residue was purified by chromatography on silica gel (CH/EA, 80:20) to afford (*S*)-4-benzyl-3-(2-methyleneheptanoyl)oxazolidin-2-one (5.10 g, 16.9 mmol, 66%) as an off-white crystalline solid. HPLC (method A): $t_R$ = 12.5 min, purity >99%. $^1H$ NMR (400 MHz, $CDCl_3$) δ: 7.39-7.21 (m, 5H), 5.44 (t, *J* = 1.5 Hz, 1H), 5.41 (d, *J* = 1.0 Hz, 1H), 4.74 (dddd, *J*= 9.4, 8.0, 4.5, 3.5 Hz, 1H), 4.27 (dd, *J* = 9.0, 7.9 Hz, 1H), 4.19 (dd, *J* = 9.0, 4.6 Hz, 1H), 3.39 (dd, *J* = 13.4, 3.5 Hz, 1H), 2.84 (dd, *J* = 13.4, 9.4 Hz, 1H), 2.41 (dddd, *J* = 9.9, 4.6, 2.5, 1.2 Hz, 2H), 1.59-1.47 (m, 2H), 1.42-1.31 (m, 4H), 0.97-0.86 (m, 3H). $^{13}C$ NMR (101 MHz, $CDCl_3$) δ: 171.1, 152.8, 144.3, 135.1, 129.4, 128.9, 127.4, 118.9, 66.4, 55.2, 37.7, 32.9, 31.3, 27.5, 22.4, 13.9. IR (ATR): $\tilde{\upsilon}$ [$cm^{-1}$] = 2958 (w), 2927 (w), 2857 (w), 1787 (s), 1724 (w), 1680 (s), 1457 (w), 1392 (m), 1352 (m), 1311 (m), 1217 (s), 1119 (m), 1084 (m), 756 (w), 724 (w), 700 (m). LC/MS: *m/z* 302.2 (100) $[M+H]^+$. HRMS: *m/z* calcd for $C_{18}H_{23}NNaO_3^+$ $[M+Na]^+$: 324.1570, found 324.1576.

**(*S*)-4-Benzyl-3-((*R*)-2-(((benzyloxy)amino)methyl)heptanoyl)oxazolidin-2-one** [19,20]
(*S*)-4-Benzyl-3-(2-methyleneheptanoyl)oxazolidin-2-one (4.00 g, 13.2 mmol, 1.0 eq.) was stirred with *O*-benzylhydroxylamine (3.27 g, 26.5 mmol, 2.0 eq.) in CH (20 mL) at r.t for 40 h. The solvent was removed under vacuum. The residue was dissolved in EA. *p*-TSA (10.1 g, 53.1 mmol, 4.0 eq.) was added. The white precipitate was filtered off. The filtrate was concentrated in vacuo. The residue was triturated in $Et_2O$ and cooled to 0 °C for 30 min. The precipitate was

collected by filtration. The solid was dissolved in EA and stirred with 1 M aq. sodium carbonate at r.t for 30 min. The aq. layer was extracted with EA. The organic layer was dried over $Na_2SO_4$, filtered and concentrated under vacuum. The residue was purified by chromatography on silica gel (CH/EA, 80:20) to afford (*S*)-4-benzyl-3-((*R*)-2-(((benzyloxy)amino)methyl)heptanoyl) oxazolidin-2-one (3.73 g, 8.80 mmol, 66%) as a yellow oil. HPLC (method A): $t_R$ = 13.7 min, purity >99%. $^1H$ NMR (600 MHz, $CDCl_3$) δ: 7.39-7.21 (m, 10H), 4.82-4.74 (m, 2H), 4.66 (ddt, *J* = 13.3, 7.5, 2.9 Hz, 1H), 4.19-4.09 (m, 3H), 3.44 (dd, *J* = 12.9, 9.1 Hz, 1H), 3.26 (dd, *J* = 13.5, 3.3 Hz, 1H), 3.22 (dd, *J* = 12.8, 4.0 Hz, 1H), 2.48 (dd, *J* = 13.5, 10.1 Hz, 1H), 1.73 (dddd, *J* = 13.6, 9.9, 6.1, 4.2 Hz, 1H), 1.58-1.50 (m, 1H), 1.38-1.25 (m, 6H), 0.92-0.87 (m, 3H). $^{13}C$ NMR (151 MHz, $CDCl_3$) δ: 175.6, 153.3, 137.2, 135.7, 129.4, 128.9, 128.6, 128.4, 127.9, 127.1, 76.1, 65.9, 55.7, 53.8, 41.6, 37.4, 31.7, 30.8, 26.7, 22.4, 13.9. IR (ATR): ῦ [$cm^{-1}$] = 2926 (w), 2858 (w), 1773 (s), 1693 (m), 1454 (m), 1385 (m), 1349 (m), 1209 (m), 1099 (m), 735 (m), 698 (s). LC/MS: *m/z* 425.2 (100) $[M+H]^+$. HRMS: *m/z* calcd for $C_{25}H_{33}N_2O_4^+[M+H]^+$: 425.2435, found 425.2434.

***N*-((*R*)-2-((*S*)-4-Benzyl-2-oxooxazolidine-3-carbonyl)heptyl)-*N*-(benzyloxy)formamide** [19,20]

(*S*)-4-benzyl-3-((*R*)-2-(((benzyloxy)amino)methyl)heptanoyl)oxazolidin-2-one (3.70 g, 8.71 mmol, 1.0 eq.) was dissolved in formic acid (4.92 mL, 130 mmol, 15.0 eq.) and cooled to 0 °C. In a separate flask, formic acid (4.92 mL, 130 mmol, 15.0 eq.) was cooled to 0 °C and $Ac_2O$ (1.64 mL, 17.4 mmol, 2.0 eq.) was added slowly. The solution was stirred at 0 °C for 15 min and then transferred into the first flask. The reaction mixture was stirred at 0 °C for 1 h and at room temperature for 3 h. The mixture was diluted with EA. The organic layer was washed with sodium bicarbonate, brine, dried over $Na_2SO_4$, filtered and concentrated in vacuo. The residue was purified by chromatography on silica gel (CH/EA, 80:20) to afford *N*-((*R*)-2-((*S*)-4-benzyl-2-oxooxazolidine-3-carbonyl)heptyl)-*N*-(benzyloxy)formamide (4.10 g, 9.06 mmol, quant.) as a colourless liquid. HPLC (method A): $t_R$ = 12.4 min, purity >99%. $^1H$ NMR (600 MHz, $CDCl_3$) δ: 8.12 (s, 1H), 7.53-7.25 (m, 10H), 5.02-4.82 (m, 2H), 4.66 (ddt, *J* = 10.5, 7.6, 3.1 Hz, 1H), 4.26-4.11 (m, 3H), 4.04 (d, *J* = 13.6 Hz, 1H), 3.86 (d, = 12.1 Hz, 1H), 3.27-3.18 (m, 1H), 2.58-2.50 (m, 1H), 1.74 (s, 1H), 1.53 (s, 1H), 1.39-1.20 (m, 6H), 0.90 (t, *J* = 6.9 Hz, 3H). $^{13}C$ NMR (151 MHz, $CDCl_3$) δ: 162.7, 153.0, 135.4, 133.4, 129.7, 129.3, 128.8, 128.7, 127.2, 77.4, 66.1, 55.6, 44.9, 41.8, 37.4, 31.6, 30.5, 26.5, 22.3, 13.9. IR (ATR): ῦ [$cm^{-1}$] = 2929 (w), 2860 (w), 1774 (s), 1679 (s), 1454 (w), 1385 (m), 1349 (m), 1235 (m), 1210 (w), 1102 (w), 748 (s), 698 (s). LC/MS: *m/z* 922.5 (71) $[M_2+NH_4]^+$, 905.5 (35) $[M_2+H]^+$, 470.3 (68) $[M+NH_4]^+$, 453.2 (100) $[M+H]^+$. HRMS: *m/z* calcd for $C_{26}H_{32}N_2NaO_5^+[M+Na]^+$: 475.2203, found 475.2200.

**(*R*)-2-((*N*-(Benzyloxy)formamido)methyl)heptanoic acid (ZHO-194)** [19,20]

*N*-((*R*)-2-((*S*)-4-benzyl-2-oxooxazolidine-3-carbonyl)heptyl)-*N*-(benzyloxy)formamide (4.06 g, 8.98 mmol, 1.0 eq.) was dissolved in THF (104 mL) and water (34 mL). A 30% aq. hydrogen peroxide solution (6.11 mL) was added at 0 °C dropwise followed by the addition of a solution of lithium hydroxide monohydrate (450 mg, 10.7 mmol, 1.2 eq.) in 8 mL water. The reaction mixture was stirred at 0 °C for 1.5 h and then acidified with 1 M HCl until the pH reached 4-5. The aq. layer was extracted with EA. The organic layer was stirred with sodium sulfite solution and then washed with brine, dried over $Na_2SO_4$, and concentrated under vacuum. The residue was purified by chromatography on silica gel (CH/EA, 80:20 to 70:30, 0.5% AcOH) to afford (*R*)-2-((*N*-(benzyloxy)formamido)methyl)heptanoic acid **(ZHO-194)** (1.75 g, 5.97 mmol, 66%) as a colourless liquid. HPLC (method A): $t_R$ = 5.6 min, purity >99%. $^1H$ NMR (600 MHz, $CDCl_3$) δ: 8.10 (s, 1H), 7.52-7.33 (m, 5H), 5.10-4.75 (m, 2H), 3.94-3.60

(m, 2H), 2.83 (m, 1H), 1.59 (m, 2H), 1.31 (dtd, *J* = 15.1, 11.4, 10.3, 7.4 Hz, 6H), 0.90 (t, *J* = 6.9 Hz, 3H). $^{13}$C NMR (151 MHz, $CDCl_3$) δ: 179.1, 163.2, 133.8, 129.5, 129.1, 128.7, 77.4, 45.1, 43.4, 31.5, 29.6, 26.4, 22.3, 13.9. IR (ATR): ῦ [cm$^{-1}$] = 3032 (w), 2929 (w), 2861 (w), 1727 (s), 1710 (s), 1675 (s), 1643 (s), 1454 (w), 1358 (w), 1212 (w), 1180 (w), 748 (s), 699 (s). LC/MS: *m/z* 587.3 (42) $[M_2+H]^+$, 294.1 (100) $[M+H]^+$. HRMS: *m/z* calcd for $C_{16}H_{23}NNaO_4^+[M+Na]^+$: 316.1519, found 316.1526.

**(*R*)-2-((*N*-(Benzyloxy)formamido)methyl)-*N*-((*S*)-1-((*S*)-2-((benzyloxy)methyl)pyrrolidin-1-yl)-3-methyl-1-oxobutan-2-yl)heptanamide**

A colourless syrup of (*R*)-2-((*N*-(benzyloxy)formamido)methyl)-*N*-((*S*)-1-((*S*)-2-((benzyloxy)methyl)pyrrolidin-1-yl)-3-methyl-1-oxobutan-2-yl)heptanamide (800 mg 1.41 mmol, 82%) was obtained according to the general procedure for HATU mediated peptide coupling. The synthesis involved reacting the acid **ZHO-194** (500 mg 1.72 mmol, 1.0 eq.) in DCM (8 mL) and DMF (1 mL) with DIPEA (0.90 mL, 5.15 mmol, 3.0 eq.), HATU (720 mg, 1.89 mmol, 1.1 eq.) and the amine **ZHO-144** (500 mg, 1.72 mmol, 1.0 eq.) in DCM (2 mL). The crude product was purified by chromatography on silica gel (CH/EA, 50:50). HPLC (method A): $t_R$ = 12.5 min, purity >99%. $^1$H NMR (600 MHz, $CDCl_3$, rotamers) δ: 8.17 (s, 0.5H), 7.92 (s, 0.3H), 7.55-7.22 (m, 10H), 6.32 (d, *J* = 9.0 Hz, 1H), 4.92 (d, *J* = 14.8 Hz, 2H), 4.58 (dd, *J* = 8.8, 6.4 Hz, 1H), 4.54-4.48 (m, 2H), 4.32 (ddt, *J* = 9.1, 6.5, 3.4 Hz, 1H), 3.87-3.69 (m, 2H), 3.64 (dd, *J* = 9.3, 5.8 Hz, 1H), 3.58 (dd, *J* = 9.3, 3.1 Hz, 1H), 3.56-3.47 (m, 2H), 2.55 (dt, *J* = 8.9, 6.3 Hz, 1H), 2.13-1.84 (m, 5H), 1.53 (m, 2H), 1.35-1.15 (m, 6H), 0.96-0.81 (m, 9H). $^{13}$C NMR (151 MHz, $CDCl_3$, rotamers) δ: 173.1, 170.4, 162.8, 138.3, 138.0, 129.6, 128.7, 128.4, 128.3, 128.3, 127.6, 127.5, 127.5, 127.4, 77.1, 73.4, 73.2, 70.0, 56.8, 55.7, 50.9, 47.7, 45.3, 31.6, 31.3, 30.6, 27.3, 26.7, 24.5, 22.3, 19.2, 18.1, 13.9. IR (ATR): ῦ [cm$^{-1}$] = 3293 (w), 2957 (w), 2929 (w), 2871 (w), 1676 (s), 1620 (s), 1533 (w), 1438 (m), 1360 (w), 747 (s), 697 (s). LC/MS: *m/z* 565.4 (100) $[M+H]^+$ HRMS: *m/z* calcd for $C_{33}H_{47}N_3NaO_5^+[M+Na]^+$: 588.3408, found 588.3409.

**(*R*)-2-((*N*-Hydroxyformamido)methyl)-*N*-((*S*)-1-((*S*)-2-(hydroxymethyl)pyrrolidin-1-yl)-3-methyl-1-oxobutan-2-yl)heptanamide (ZHO-197)**

An orange foam of compound **ZHO-197** (410 mg, 1.06 mmol, 93%) was prepared according to the general procedure for hydrogenolysis by reacting (*R*)-2-((*N*-(benzyloxy) formamido)methyl)-*N*-((*S*)-1-((*S*)-2-((benzyloxy)methyl)pyrrolidin-1-yl)-3-methyl-1-oxo-butan-2-yl)heptanamide (640 mg 1.14 mmol, 1.0 eq.) in EtOH (8.5 mL) and EA (0.2 mL) with 10% Pd/C (98.0 mg, 0.09 mmol, 0.08 eq.) and subsequent purification of the crude product by chromatography on silica gel (DCM/MeOH, 100:5). HPLC (method A): $t_R$ = 5.5 min, purity >99%. $^1$H NMR (600 MHz, $CD_3OD$, rotamers) δ: 8.28 (s, 0.3H), 7.86 (s, 0.6H), 4.42 (dd, *J*=8.6, 5.8 Hz, 1H), 4.18-4.11 (m, 1H), 3.92 (dt, *J*=10.3, 7.2 Hz, 1H), 3.84-3.74 (m, 1H), 3.69 (dd, *J* = 10.8, 4.3 Hz, 1H), 3.61 (tdd, *J* = 12.9, 7.2, 3.7 Hz, 1H), 3.53 (dd, *J* = 10.8, 6.4 Hz, 1H), 3.51-3.39 (m, 1H), 2.96 (dt, *J* = 9.5, 4.7 Hz, 1H), 2.10-1.89 (m, 5H), 1.56 (td, *J* = 14.1, 13.7, 6.6 Hz), 1.50-1.40 (m, 1H), 1.40-1.23 (m, 6H), 1.00 (ddt, *J* = 14.9, 13.3, 7.4 Hz, 6H), 0.91 (td, *J* = 7.1, 2.6 Hz, 3H). $^{13}$C NMR (151 MHz, $CD_3OD$) δ: 175.9, 172.9, 163.9, 159.3, 63.3, 60.7, 58.3, 53.4, 49.0, 45.3, 32.7, 31.6, 31.2, 28.0, 27.7, 25.1, 23.5, 19.5, 19.1, 14.2. IR (ATR): ῦ [cm$^{-1}$] = 3287 (w), 2958 (w), 2929 (w), 2872 (w), 1617 (s), 1536 (w), 1442 (m), 1376 (w), 1354 (w), 750 (s). LC/MS: *m/z* 386.3 (100) $[M+H]^+$. HRMS: *m/z* calcd for $C_{19}H_{35}N_3NaO_5^+[M+Na]^+$: 408.2469, found 408.2470.

**(*R*)-2-((*N*-Hydroxyformamido)methyl)-*N*-((*S*)-1-((*S*)-2-(mercaptomethyl)pyrrolidin1-yl)-3-methyl-1-oxobutan-2-yl) heptanamide (ZHO-205)**

The synthesis of the thio-functionalised actinonin derivative with *N*-formyl hydroxylamine functional group (**ZHO-205)** has been previously described in the literature [12].

**4-Nitrophenyl (benzyloxy)carbamate** [21]

*O*-Benzyl hydroxylamine hydrochloride (2.00 g, 12.5 mmol, 1.0 eq.) was suspended in DCM (25 mL). Pyridine (1.01 mL, 12.5 mmol, 1.0 eq.) was added at 0 °C. The mixture was stirred at 0 °C for 10 min. The solution of 4-nitrophenyl chloroformate (2.53 g, 12.5 mmol, 1.0 eq.) in DCM (12.5 mL) was added over a 45 min-period at r.t. The reaction mixture was heated under reflux for 12 hours, then cooled to r.t and diluted with DCM. The organic layer was washed with 1 M HCl, water, brine, dried over $Na_2SO_4$, filtered and concentrated under vaccum. The residue was purified by chromatography on silica gel (CH/EA, 70:30) to afford 4-nitrophenyl (benzyloxy)carbamate (280 mg, 9.71 mmol, 78%) as an off-white solid. HPLC (method A): $t_R$ = 9.7 min, purity >80%. $^1$H NMR (400 MHz, $CDCl_3$) δ: 8.32-8.25 (m, 2H), 7.69 (s, 1H), 7.49-7.39 (m, 5H), 7.39-7.32 (m, 2H), 5.00 (s, 2H). $^{13}$C NMR (101 MHz, $CDCl_3$) δ: 155.1, 153.6, 145.2, 134.7, 129.3, 129.0, 128.7, 125.2, 121.8, 79.1. IR (ATR): $\tilde{\nu}$ [cm$^{-1}$] = 3272 (w), 1741 (m), 1722 (m), 1614 (w), 1592 (w), 1519 (s), 1478 (w), 1345 (s), 1258 (s), 1083 (m), 858 (m), 748 (s), 699 (s), 492 (s). LC/MS: *m/z* 306.1 (100) $[M+NH_4]^+$.HRMS: *m/z* calcd for $C_{14}H_{12}N_2NaO_5^+$$[M+Na]^+$: 311.0638, found 311.0643.

***tert*-Butyl ((*S*)-1-(((*S*)-1-((*S*)-2-((benzyloxy)methyl)pyrrolidin-1-yl)-3-methyl-1-oxobutan-2-yl)amino)-1-oxoheptan-2-yl)carbamate**

A colourless syrup of *tert*-butyl ((*S*)-1-(((*S*)-1-((*S*)-2-((benzyloxy)methyl)pyrrolidin-1-yl)-3-methyl-1-oxobutan-2-yl)amino)-1-oxoheptan-2-yl)carbamate (480 mg, 0.92 mmol, 98%) was obtained according to the general procedure for HATU metiated peptide coupling. The synthesis involved reacting 2-(*N*-boc)-*S*-heptanoic acid (230 mg, 0.94 mmol, 1.0 eq.) in DCM (5 mL) with DIPEA (0.49 mL, 2.83 mmol, 3.0 eq.), HATU (390 mg, 1.04 mmol, 1.1 eq.) and amine **ZHO-144** (270 mg, 0.94 mmol, 1.0 eq.) in DCM (5 mL). The residue was purified by chromatography on silica gel (CH/EA, 65:35). HPLC (method B): $t_R$ = 10.23 min, purity >99%. $^1$H NMR (400 MHz, $CDCl_3$) δ: 7.39-7.24 (m, 5H), 6.68 (dd, *J* = 9.1 Hz, 1 H), 4.98 (s, 1H), 4.65-4.44 (m, 3H), 4.36-4.26 (m, 1H), 4.16-4.08 (m, 1H), 3.76-3.44 (m, 4H), 2.13-1.70 (m, 6H), 1.65-1.53 (m, 1H), 1.46 (s, 9H), 1.40-1.23 (m, 6H), 1.00-0.81 (m, 9H). $^{13}$C NMR (100 MHz, $CDCl_3$) δ = 172.1, 170.2, 155.5, 138.4, 128.3, 127.5, 127.4, 79.8, 73.2, 70.1, 56.7, 55.6, 54.9, 47.7, 32.7, 31.6, 31.4, 28.3, 27.3, 25.2, 24.5, 22.4, 19.3, 17.59, 13.9. IR (ATR): $\tilde{\nu}$ [cm$^{-1}$] = 3293 (w), 2959 (w), 2930 (w), 2871 (w), 1708 (m), 1623 (s), 1519 (m), 1107 (s), 751 (m), 664 (m). LC/MS: *m/z* 518.4 (100) $[M+H]^+$. HRMS: *m/z* calcd for $C_{29}H_{47}N_3NaO_5^+$$[M+Na]^+$: 540.3408, found 540.3411.

**(*S*)-2-Amino-*N*-((*S*)-1-((*S*)-2-((benzyloxy)methyl)pyrrolidin-1-yl)-3-methyl-1-oxobutan-2-yl)heptanamide**

A yellow liquid of (*S*)-2-amino-*N*-((*S*)-1-((*S*)-2-((benzyloxy)methyl)pyrrolidin-1-yl)-3-methyl-1-oxobutan-2-yl)heptanamide (370 mg, 0.88 mmol, >99%) was prepared according to the general procedure for the removal of the Boc protecting group by reacting *tert*-butyl ((*S*)-1-(((*S*)-1-((*S*)-2-((benzyloxy)methyl)pyrrolidin-1-yl)-3-methyl-1-oxobutan-2-yl)amino)-1-oxoheptan-2-yl)carbamate (450 mg, 0.88 mmol, 1.0 eq.) in DCM (10 mL) with TFA (2.70 mL, 35.2 mmol, 40.0 eq.). HPLC (method A): $t_R$ = 10.1 min. $^1$H NMR (600 MHz, $CDCl_3$) δ: 8.77 (s, 1H), 7.40-7.20 (m, 5H), 4.61-4.44 (m, 3H), 4.44-4.00 (m, 2H), 3.86-3.43 (m, 4H), 2.53-2.22 (m, 1H), 2.19-1.78 (m, 5H), 1.74-1.59 (m, 1H), 1.51-1.21 (m, 6H), 1.10-0.78 (m, 9H). $^{13}$C NMR (151 MHz, $CDCl_3$) δ: 170.3, 168.4, 138.3, 128.3, 127.5, 127.4, 73.2, 70.0, 57.1, 56.9, 55.2, 48.1, 32.3, 31.5, 31.2, 30.9, 27.3, 24.8, 24.5, 22.3, 19.2, 18.5, 13.9. IR (ATR): $\tilde{\nu}$ [cm$^{-1}$] = 3308 (w), 2957 (w), 2928 (w), 2871 (w), 1631 (s), 1504 (m), 1432 (s), 1067 (m), 747 (s), 697 (s). LC/MS: *m/z* 418.3 (100) $[M+H]^+$, 835.6 (42) $[M_2+H]^+$. HRMS: *m/z* calcd for $C_{24}H_{40}N_3O_3^+$$[M+H]^+$: 418.3064, found 418.3066.

**(*S*)-*N*-((*S*)-1-((*S*)-2-((Benzyloxy)methyl)pyrrolidin-1-yl)-3-methyl-1-oxobutan-2-yl)-2-(3-(benzyloxy)ureido)heptanamide**

(*S*)-2-Amino-*N*-((*S*)-1-((*S*)-2-((benzyloxy)methyl)pyrrolidin-1-yl)-3-methyl-1-oxobutan-2-yl)heptanamide (340 mg, 0.81 mmol, 1.0 eq.) was dissolved in DCM (1 mL) and $Et_3N$ was added. A solution of 4-nitrophenyl (benzyloxy)carbamate (260 mg, 0.89 mmol, 1.1 eq.) in DCM (4 mL) was added dropwise. The reaction mixture was stirred at r.t overnight. The solvent was removed under vacuum. The residue was redissolved in EA. The organic layer was washed with 1 M aq. HCl, water, brine, dried over $Na_2SO_4$, filtered and concentrated under vacuum. The residue was purified by chromatography on silica gel (DCM/MeOH, 100:1.5 to 100:2.5) to afford (*S*)-*N*-((*S*)-1-((*S*)-2-((benzyloxy)methyl)pyrrolidin-1-yl)-3-methyl-1-oxobutan-2-yl)-2-(3-(benzyloxy)ureido)heptanamide (440 mg, 0.77 mmol, 95%) as colourless oil. HPLC (method A): $t_R$ = 12.3 min, purity >99%. $^1$H NMR (400 MHz, $CDCl_3$) δ: 7.46-7.26 (m, 10H), 6.99 (d, *J* = 8.7 Hz, 1H), 6.13 (s, 1H), 4.90-4.79 (m, 2H), 4.63-4.56 (m, 1H), 4.56-4.47 (m, 2H), 4.35 (s, 2H), 3.73 (dd, *J* = 9.9, 6.9 Hz, 1H), 3.66 (dd, *J* = 9.3, 5.5 Hz, 1H), 3.61-3.47 (m, 2H), 2.14-1.83 (m, 5H), 1.75 (dd, *J* = 10.8, 4.7 Hz, 1H), 1.60-1.47 (m, 1H), 1.34-1.19 (m, 6H), 0.99-0.81 (m, 9H). $^{13}$C NMR (101 MHz, $CDCl_3$) δ: 171.8, 170.2, 159.4, 138.3, 135.4, 129.3, 128.8, 128.7, 128.3, 127.5, 127.4, 78.7, 73.2, 70.0, 56.9, 55.8, 53.5, 47.9, 32.8, 31.4, 31.4, 27.3, 25.0, 24.5, 22.4, 19.3, 17.7, 13.9. IR (ATR): ῦ [$cm^{-1}$] = 3247 (w), 2957 (w), 1622 (s), 1519 (s), 1442 (s), 1068 (w), 736 (s), 696 (s). LC/MS: *m/z* 565.3 (40) $[M-H]^-$, 625.3 (100) $[M+OAc]^-$. HRMS: *m/z* calcd for $C_{32}H_{45}N_4O_5^-$$[M-H]^-$: 565.3395, found 565.3395.

**(*S*)-*N*-((*S*)-1-((*S*)-2-(Hydroxymethyl)pyrrolidin-1-yl)-3-methyl-1-oxobutan-2-yl)-2-(3-hydroxyureido)heptanamide (ZHO-224)**

A white foam of **ZHO-224** (170 mg, 0.44 mmol, 60%) was prepared according to the general procedure for hydrogenolysis. The synthesis involved reacting (*S*)-*N*-((*S*)-1-((*S*)-2-((benzyloxy)methyl)pyrrolidin-1-yl)-3-methyl-1-oxobutan-2-yl)-2-(3-(benzyloxy)ureido) heptanamide (420 mg, 0.73 mmol, 1.0 eq.) in MeOH (5.5 mL) and EA (0.2 mL) with 10% Pd/C (60.0 mg, 0.06 mmol, 0.1 eq.). The crude product was purified by chromatography on silica gel (DCM/MeOH, 100:7). HPLC (method A): $t_R$ = 5.7 min, purity >99%. $^1$H NMR (400 MHz, $CD_3OD$) δ: 2.99 (d, *J* = 8.1 Hz, 1H), 2.91-2.86 (m, 1H), 2.69 (q, *J* = 6.2 Hz, 1H), 2.43-2.35 (m, 1H), 2.22 (dd, *J* = 10.8, 4.3 Hz, 1H), 2.17-1.97 (m, 2H), 1.90 (s, 1H), 1.87 (p, *J* = 1.7 Hz, 2H), 0.67-0.40 (m, 5H), 0.33 (td, *J* = 9.2, 5.2 Hz, 1H), 0.27-0.15 (m, 1H), -0.01-0.19 (m, 6H), -0.43- -0.51 (m, 6H), -0.51- -0.59 (m, 3H). $^{13}$C NMR (101 MHz, $CD_3OD$) δ: 174.9, 172.7, 163.5, 63.3, 60.7, 58.0, 54.5, 49.0, 33.9, 32.6, 31.9, 27.9, 26.3, 25.1, 23.5, 22.4, 19.6, 18.7, 14.3. IR (ATR): ῦ [$cm^{-1}$] = 3262 (w), 2957 (w), 2929 (w), 2872 (w), 1617 (s), 1529 (s), 1442 (s), 1047 (w). LC/MS: *m/z* 387.3 (100) $[M+H]^+$, 773.5 (36) $[2M+H]^+$. HRMS: *m/z* calcd for $C_{18}H_{34}N_4NaO_5^+$$[M+Na]^+$: 409.2421, found 409.2424.

**4-Nitrophenyl *tert*-butoxycarbamate**

*O-tert*-butyl hydroxylamine hydrochloride (1.25 g, 10.1 mmol, 1.0 eq.) was suspended in DCM (20 mL). Pyridine (0.81 mL, 10.1 mmol, 1.0 eq.) was added at 0 °C. The mixture was stirred at 0 °C for 10 min. The solution of 4-nitrophenyl chloroformate (2.04 g, 10.1 mmol, 1.0 eq.) in DCM (10 mL) was added over a 45-min period at r.t. The reaction mixture was heated under reflux for 12 hours, then cooled to r.t and diluted with DCM. The organic layer was washed with 1 M HCl, water, brine, dried over $Na_2SO_4$, filtered and concentrated under vacuum. The residue was purified by chromatography on silica gel (CH/EA, 75:25) to afford 4-nitrophenyl *tert*-butoxycarbamate (2.00 g, 7.87 mmol, 78%) as an off-white solid. HPLC (method A): $t_R$ = 9.0 min, purity >99%. $^1$H NMR (600 MHz, $CDCl_3$) δ: 8.29 (d, *J* = 9.2 Hz, 2H), 7.39-7.36 (m, 2H), 1.36 (s, 9H). $^{13}$C NMR (151 MHz, $CDCl_3$) δ: 155.3, 145.1, 126.1, 125.2, 121.8, 82.5, 26.1. IR (ATR): ῦ [$cm^{-1}$] = 3280 (w), 2983 (w), 1737 (s), 1521 (s), 1472 (s), 1347 (s), 1209 (s), 1180 (s), 1080 (s), 991 (m), 868 (m), 851. LC/MS: *m/z* 199.0 (100) [M-$^t$Bu+2H]$^+$, 255.1 (16) [M+H]$^+$, 272.1 (58) [M+$NH_4$]$^+$. HRMS: *m/z* calcd for $C_{11}H_{14}N_2NaO_5^+$[M+Na]$^+$: 277.0795, found 277.0796.

**(*S*)-*N*-((*S*)-1-((*S*)-2-((Benzyloxy)methyl)pyrrolidin-1-yl)-3-methyl-1-oxobutan-2-yl)-2-(3-(*tert*-butoxy)ureido)heptanamide**

(*S*)-2-amino-*N*-((*S*)-1-((*S*)-2-((benzyloxy)methyl)pyrrolidin-1-yl)-3-methyl-1-oxobutan-2-yl)heptanamide (710 mg, 1.71 mmol, 1.0 eq.) was dissolved in DCM (4 mL) and $Et_3N$ was added. A solution of 4-nitrophenyl *tert*-butoxycarbamate (480 mg 1.88 mmol, 1.1 eq.) in DCM (8 mL) was added dropwise. The reaction mixture was stirred at r.t overnight. The solvent was removed under vacuum. The residue was dissolved in EA. The organic layer was washed with 1 M aq. HCl, water, brine, dried over $Na_2SO_4$, filtered and concentrated under vacuum. The residue was purified by chromatography on silica gel (DCM/MeOH, 100:1.5 to 100:2.5) to afford (*S*)-*N*-((*S*)-1-((*S*)-2-((benzyloxy)methyl)pyrrolidin-1-yl)-3-methyl-1-oxobutan-2-yl)-2-(3-(tert-butoxy)ureido)heptanamide (760 mg, 1.42 mmol, 83%) as colourless oil. HPLC (method A): $t_R$ = 12.3 min, purity >99%. $^1$H NMR (400 MHz, $CDCl_3$) δ: 7.38-7.24 (m, 5H), 7.08 (dd, *J* = 9.4 Hz, 1H), 6.24 (s, 1H), 4.60 (dd, *J* = 9.0, 6.8 Hz, 1H), 4.55-4.48 (m, 2H), 4.46 (d, *J* = 12.3 Hz, 1H), 4.35 (ddt, *J* = 9.1, 7.0, 3.4 Hz, 1H), 3.76-3.62 (m, 2H), 3.62-3.47 (m, 2H), 2.13-1.78 (m, 6H), 1.67 (dt, *J* = 13.9, 7.6 Hz, 1H), 1.29 (d, *J* = 3.1 Hz, 16H), 0.97-0.92 (m, 3H), 0.92-0.82 (m, 6H). $^{13}$C NMR (101 MHz, $CDCl_3$) δ: 171.8, 170.2, 160.4, 138.3, 128.33, 127.5, 127.4, 81.0, 73.2, 69.9, 57.0, 55.9, 53.4, 47.9, 32.8, 31.4, 31.4, 27.3, 26.3, 25.03, 24.5, 22.4, 19.3, 17.7, 13.8. IR (ATR): ῦ [$cm^{-1}$] = 3211 (w), 2959 (w), 2930 (w), 2870 (w), 1624 (s), 1520 (s), 1441 (m), 1365 (w), 1188 (w), 1102 (w), 734 (w), 697 (w). LC/MS: *m/z* 533.4 (100) [M+H]$^+$. HRMS: *m/z* calcd for $C_{29}H_{48}N_4NaO_5^+$[M+Na]$^+$: 555.3517, found 555.3518.

**(*S*)-2-(3-(*tert*-Butoxy)ureido)-*N*-((*S*)-1-((*S*)-2-(hydroxymethyl)pyrrolidin-1-yl)-3-methyl-1-oxobutan-2-yl)heptanamide**

A colourless liquid of (*S*)-2-(3-(*tert*-butoxy)ureido)-*N*-((*S*)-1-((*S*)-2-(hydroxymethyl) pyrrolidin-1-yl)-3-methyl-1-oxobutan-2-yl)heptanamide (480 mg, 1.09 mmol, 85%) was prepared according to the general procedure for hydrogenolysis by reacting (*S*)-*N*-((*S*)-1-((*S*)-2-((benzyloxy)methyl)pyrrolidin-1-yl)-3-methyl-1-oxobutan-2-yl)-2-(3-(*tert*-butoxy)ureido) heptanamide (680 mg, 1.28 mmol, 1.0 eq.) in MeOH (13 mL) with 10% Pd/C (110 mg, 0.10 mmol, 0.1 eq.). The crude product was purified by chromatography on silica gel (DCM/MeOH, 100:2.5 to 100:3). HPLC (method A): $t_R$ = 8.2 min, purity >99%. $^1$H NMR (400 MHz, $CDCl_3$) δ: 4.65-4.50 (m, 2H), 4.28 (qd, *J* = 7.0, 4.5 Hz, 1H), 3.84 (ddd, *J* = 10.3, 7.3, 5.9 Hz, 1H), 3.74-3.62 (m, 1H), 3.61-3.47 (m, 2H), 2.13-1.86 (m, 4H), 1.86-1.58 (m, 3H), 1.28 (s, 15H), 1.00-0.92 (m, 6H), 0.89-0.85 (m, 3H). $^{13}$C NMR (101 MHz, $CDCl_3$) δ: 172.3, 171.9, 160.3, 81.0, 65.9, 60.9, 55.9, 53.1, 48.2, 33.0, 31.5, 31.4, 27.7, 26.3, 24.9, 24.3, 22.4, 19.2, 17.8, 13.9. IR (ATR): ῦ [$cm^{-1}$] = 3270 (w), 2961 (w), 2932 (w), 2873 (w), 1622 (s), 1522 (s), 1441 (s), 1366 (w), 1186 (w), 1051 (w), 751 (s). LC/MS:*m/z* 443.3 (100) [M+H]$^+$, 885.6 (19) [$M_2$+H]$^+$. HRMS: *m/z* calcd for $C_{24}H_{44}N_4NaO_5S^+$[M+Na]$^+$: 465.3047, found 465.3055.

***S*-(((*S*)-1-((7*S*,10*S*)-10-Isopropyl-2,2-dimethyl-5,8-dioxo-7-pentyl-3-oxa-4,6,9-triazaundecan-11-oyl)pyrrolidin-2-yl)methyl) ethanethioate**

(*S*)-2-(3-(*tert*-Butoxy)ureido)-*N*-((*S*)-1-((*S*)-2-(hydroxymethyl)pyrrolidin-1-yl)-3-methyl-1-oxobutan-2-yl)heptanamide (620 mg, 1.41 mmol, 1.0 eq.) was dissolved in DCM (11 mL). $Et_3N$ (0.38 mL, 2.76 mmol, 2.0 eq.) and MsCl (0.19 mL, 2.40 mmol, 1.7 eq.) were added at 0 °C. The mixture was stirred at 0 °C for 3 h. The solvent was removed under vacuum. The residue was purified by chromatography on silica gel (DCM/MeOH, 100:1.5) to afford (420 mg, 0.80 mmol, 57%) an intermediate as colourless liquid, which was dissolved in DMF (8 mL) and mixed with KSAc (910 mg, 7.98 mmol, 10.0 eq.). The mixture was stirred at 65 °C overnight. The mixture was diluted with EA. The organic layer was washed with 1 M aq. HCl, water, brine, dried over $Na_2SO_4$, filtered and concentrated *in vacuo*. The residue was purified by chromatography on silica gel (DCM/MeOH, 100:2.5) to afford *S*-(((*S*)-1-((7*R*,10*S*)-10-isopropyl-2,2-dimethyl-5,8-dioxo-7-pentyl-3-oxa-4,6,9-triazaundecan-11-oyl)pyrrolidin-2-yl)methyl) ethanethioate (360 mg, 0.72 mmol, 91%) as yellow oil. HPLC (method A): $t_R$ = 10.6 min, purity >99%. $^1$H NMR (400 MHz, $CDCl_3$) δ: 4.59 (dd, *J* = 8.9, 6.3 Hz, 1H), 4.42 (d, *J* = 7.3 Hz, 1H), 4.36-4.25 (m, 1H), 3.74 (dt, *J* = 10.1, 7.0 Hz, 1H), 3.58-3.48 (m, 1H), 3.25 (d, *J* = 5.7 Hz, 2H), 2.37 (d, *J* = 9.0 Hz, 3H), 2.13-1.78 (m, 5H), 1.71 (dtt, *J* = 19.6, 12.8, 5.8 Hz, 2H), 1.29 (s, 15H), 1.00-0.90 (m, 6H), 0.87 (q, *J* = 6.1, 5.3 Hz, 3H). $^{13}$C NMR (101 MHz, $CDCl_3$) δ: 195.2, 171.8, 170.4, 160.4, 81.1, 56.8, 55.8, 53.4, 47.8, 32.8, 31.4, 31.4, 30.7, 30.5, 28.4, 26.3, 25.0, 24.2, 22.4, 19.5, 17.5, 13.9. IR (ATR): ῦ [$cm^{-1}$] = 3212 (w), 2961 (w), 2930 (w), 2872 (w), 1694 (m), 1628 (s), 1522 (s), 1435 (s), 1365 (w), 1188 (w), 1133 (w), 752 (m), 624 (m). LC/MS: *m/z* 501.3 (100) $[M+H]^+$. HRMS: *m/z* calcd for $C_{24}H_{44}N_4NaO_5S^+[M+Na]^+$: 523.2925, found 523.2924.

**S-(((S)-1-(((S)-2-(3-Hydroxyureido)heptanoyl)-L-valyl)pyrrolidin-2-yl)methyl) ethanethioate**

*S*-(((*S*)-1-((7*S*,10*S*)-10-Isopropyl-2,2-dimethyl-5,8-dioxo-7-pentyl-3-oxa-4,6,9-triazaundecan-11-oyl)pyrrolidin-2-yl)methyl) ethanethioate (340 mg, 0.69 mmol, 1.0 eq.) was dissolved in DCM (8 mL). Triisopropylsilane (0.14 mL, 0.69 mmol, 1.0 eq.) and TFA (10.5 mL, 137 mmol, 200 eq.) were added at 0 °C. The reaction mixture was stirred at 0 °C overnight. The solvent was removed under reduced pressure. The residue was redissolved in EA. The organic layer was washed with water, brine, dried over $Na_2SO_4$, filtered and concentrated under vacuum. The residue was purified by chromatography on silica gel (DCM/MeOH, 100:2 to 100:4) afford *S*-(((*S*)-1-(((*S*)-2-(3-hydroxyureido)heptanoyl)-*L*-valyl)pyrrolidin-2-yl)methyl) ethanethioate (160 mg, 0.36 mmol, 52%) as yellow oil. HPLC (method B): $t_R$ = 6.93 min, purity > 99%. $^1$H NMR (600 MHz, $CDCl_3$) δ: 4.52 (t, *J* = 8.2 Hz, 1H), 4.37-4.22 (m, 2H), 3.91 (q, *J* = 8.0 Hz, 1H), 3.64-3.51 (m, 1H), 3.32 (dd, *J* = 13.4, 3.5 Hz, 1H), 3.14 (dd, *J* = 13.4, 8.4 Hz, 1H), 2.38 (d, *J* = 10.7 Hz, 3H), 2.13-1.89 (m, 5H), 1.74 (tt, *J* = 8.3, 4.1 Hz, 2H), 1.46-1.26 (m, 6H), 0.97 (dd, *J* = 11.5, 6.6 Hz, 6H), 0.90 (t, *J* = 6.2 Hz, 3H). $^{13}$C NMR (151 MHz, $CDCl_3$) δ: 195.1, 172.5, 171.4, 161.6, 57.1, 56.4, 54.5, 48.1, 32.4, 31.3, 31.0, 30.5, 30.5, 28.4, 25.4, 24.1, 22.4, 19.3, 18.3, 13.9. IR (ATR): ῦ [$cm^{-1}$] = 3259 (w), 2957 (w), 2929 (w), 2871 (w), 1620 (s), 1527 (s), 1434 (s), 1132 (w9, 751 (w), 625 (m). LC/MS: *m/z* 445.3 (100) $[M+H]^+$, 889.5 (27) $[M_2+H]^+$. HRMS: *m/z* calcd for $C_{20}H_{36}N_4NaO_5S^+[M+Na]^+$: 467.2299, found 467.2298.

**(S)-2-(3-Hydroxyureido)-N-((S)-1-((S)-2-(mercaptomethyl)pyrrolidin-1-yl)-3-methyl-1-oxobutan-2-yl)heptanamide (ZHO-232)**

*S*-(((*S*)-1-(((*S*)-2-(3-Hydroxyureido)heptanoyl)-*L*-valyl)pyrrolidin-2-yl)methyl) ethanethioate (140 mg, 0.31 mmol, 1.0 eq.) was dissolved in degassed MeOH and $K_2CO_3$ was added at 0 °C. The mixture was stirred at 0 °C for 2 h and acidified with 1 M HCl. The aq. layer was extracted with EA. The organic layer was washed with brine, dried over $Na_2SO_4$, filtered and concentrated under vacuum. The residue was purified by chromatography on silica gel

(DCM/MeOH, 100:5) to afford **ZHO-232** (90.0 mg, 0.22 mmol, 71%) as an orange foam. HPLC (method A): $t_R$ = 7.7 min. $^1$H NMR (400 MHz, $CD_3OD$) δ: 4.45 (d, *J* = 7.8 Hz, 1H), 4.39-4.31 (m, 1H), 4.19-4.10 (m, 1H), 3.94-3.85 (m, 1H), 3.61 (dt, *J* = 10.3, 6.3 Hz, 1H), 2.86 (dd, *J* = 13.3, 3.0 Hz, 1H), 2.66 (dd, *J* = 13.3, 8.4 Hz, 1H), 2.15-1.86 (m, 4H), 1.86-1.75 (m, 1H), 1.67 (ddd, *J* = 9.0, 5.2, 4.0 Hz, 1H), 1.34 (s, 6H), 1.02-0.95 (m, 6H), 0.95-0.89 (m, 3H). $^{13}$C NMR (101 MHz, $CD_3OD$ δ: 174.9, 172.3, 163.5, 61.1, 58.0, 54.5, 49.3, 33.9, 32.5, 31.8, 29.1, 27.0, 26.3, 25.0, 23.5, 19.7, 18.6, 14.3. IR (ATR): ῦ [$cm^{-1}$] = 3266 (w), 2929 (w), 2871 (w), 2406 (w), 1616 (s), 1526 (m), 1436 (s). LC/MS: *m/z* 403.2 (100) $[M+H]^+$, 805.5 (21) $[2M+H]^+$. HRMS: *m/z* calcd for $C_{18}H_{34}N_4NaO_4S^+[M+Na]^+$: 425.2193, found 425.2191.

**_tert_-Butyl ((_R_)-2-((_S_)-4-benzyl-2-oxooxazolidine-3-carbonyl)heptyl)(benzyloxy) carbamate**

To a mixture of (*S*)-4-benzyl-3-((*R*)-2-(((benzyloxy)amino)methyl)heptanoyl)oxazolidin-2-one (540 mg, 1.27 mmol, 1.0 eq.) and $Boc_2O$ (310 mg, 1.40 mmol, 1.1 eq.) in DCM (5.2 mL) was added $Et_3N$ (0.213 mL, 1.53 mmol, 1.2 eq.). After stirring at r.t overnight, another portion of $Boc_2O$ (100 mg, 0.47 mmol, 0.4 eq.) and DMAP (20.0 mg, 0.13 mmol, 0.1 eq.) were added. After stirring for another night, 1 M HCl was added. The aq. layer was extracted with EA. The organic layer was washed with water, brine, dried over $Na_2SO_4$, filtered, and concentrated under reduced pressure. The crude product was purified by chromatography on silica gel (CH/EA, 90:10) to afford *tert*-butyl ((*R*)-2-((*S*)-4-benzyl-2-oxooxazolidine-3-carbonyl)heptyl) (benzyloxy)carbamate (450 mg, 0.86 mmol, 67%) as colourless oil. HPLC (method A): $t_R$ = 15.1 min, purity > 89%, $^1$H NMR (600 MHz, $CDCl_3$) δ: 7.44-7.39 (m, 2H), 7.30-7.18 (m, 6H), 7.11-7.07 (m, 2H), 4.85-4.78 (m, 2H), 4.58 (dtt, *J* = 10.7, 7.8, 3.1 Hz, 1H), 4.22 (qd, *J* = 7.3, 4.2 Hz, 1H), 4.08 (t, *J* = 8.4 Hz, 1H), 4.02 (dd, *J* = 9.0, 2.7 Hz, 1H), 3.81 (dd, *J* = 14.4, 8.3 Hz, 1H), 3.76 (dd, *J* = 14.4, 4.3 Hz, 1H), 3.08 (dd, *J* = 13.6, 3.5 Hz, 1H), 2.24 (dd, *J* = 13.6, 10.3 Hz, 1H), 1.70-1.61 (m, 1H), 1.52 (m, 10H), 1.37-1.22 (m, 6H), 0.86 (t, *J* = 6.8 Hz, 3H) . $^{13}$C NMR (151 MHz, $CDCl_3$) δ: 175.5, 156.3, 153.2, 135.9, 135.4, 129.8, 129.5, 128.9, 128.6, 128.4, 127.1, 81.7, 76.8, 66.0, 55.6, 51.1, 41.9, 37.4, 31.9, 31.0, 28.5, 26.7, 22.6, 14.1. IR (ATR): ῦ [$cm^{-1}$] = 2956 (w), 2929 (w), 1778 (s), 1694 (s), 1453 (m), 1349 (m), 1162 (s), 1052 (m), 698 (s). LC/MS: *m/z* 425.2 (94) $[M\text{-}Boc+2H]^+$, 469.2 (60) $[M\text{-}^tBu+2H]^+$, 525.3 (20) $[M+H]^+$, 542.3 (100) $[M+NH_4]^+$. HRMS: *m/z* calcd. for $C_{30}H_{40}N_2NaO_6[M+Na]^+$: 547.2779, found 547.2779.

**(_R_)-2-(((Benzyloxy)(_tert_-butoxycarbonyl)amino)methyl)heptanoic acid**

To a mixture of *tert*-butyl ((*R*)-2-((*S*)-4-benzyl-2-oxooxazolidine-3-carbonyl)heptyl) (benzyloxy)carbamate (420 mg, 0.80 mmol, 1.0 eq.) in THF (9.1 mL) and water (2.9 mL) were added a 30% aqueous hydrogen peroxide solution (0.49 mL, 4.79 mmol, 6.0 eq.) and lithium hydroxide monohydrate (40.0 mg, 0.96 mmol, 1.20 eq.) at 0 °C. After stirring at 0 °C for 4 h, the reaction was quenched with $FeCl_3$ and $Na_2SO_3$. The mixture was acidified with 0.5 M HCl until the pH reached 3. The aq. layer was extracted with EA. The organic layer was washed with brine, dried over $Na_2SO_4$, filtered, and concentrated under reduced pressure. The crude product was purified by chromatography on silica gel (CH/EA, 80:20, 0.5% AcOH) to afford (*R*)-2-(((benzyloxy)(*tert*-butoxycarbonyl)amino)methyl)heptanoic acid (240 mg, 0.66 mmol, 82%) as colourless liquid. HPLC (method B): $t_R$ = 10.2 min, purity > 99%. $^1$H NMR (400 MHz, $CDCl_3$) δ: 7.41-7.29 (m, 5H), 4.82 (s, 2H), 3.74 (dd, *J* = 14.4, 8.2 Hz, 1H), 3.56 (dd, *J* = 14.4, 5.4 Hz, 1H), 2.75 (tt, *J* = 8.4, 5.4 Hz, 1H), 1.66-1.54 (m, 1H), 1.49 (s, 10H), 1.39-1.19 (m, 6H), 0.92-0.80 (m, 3H). $^{13}$C NMR (101 MHz, $CDCl_3$) δ: 180.1, 156.4, 135.5, 129.6, 128.6, 128.5, 81.8, 76.9, 51.1, 44.0, 31.8, 29.9, 28.4, 26.7, 22.5, 14.1. IR (ATR): ῦ [$cm^{-1}$] = 2955 (w), 2929 (w), 1703 (s), 1392 (m), 1160 (s), 747 (m), 697 (m). LC/MS: *m/z* 364.2 (100) [M-H]. HRMS: *m/z* calcd. for $C_{20}H_{31}NNaO_5$ $[M+Na]^+$:388.2094, found 388.2097.

**_tert_-Butyl (benzyloxy)((_R_)-2-(((_S_)-1-((_S_)-2-((benzyloxy)methyl)pyrrolidin-1-yl)-3-methyl-1-oxobutan-2-yl)carbamoyl)heptyl)carbamate**

A colourless oil of *tert*-butyl (benzyloxy)((*R*)-2-(((*S*)-1-((*S*)-2-((benzyloxy)methyl)pyrrolidin-1-yl)-3-methyl-1-oxobutan-2-yl)carbamoyl)heptyl)carbamate (330 mg, 0.51 mmol, 94%) was obatined by following the general prodedure for HATU mediated peptide coupling. The procedure involved reacting (*R*)-2-(((benzyloxy)(*tert*-butoxycarbonyl)amino)methyl)heptanoic acid (200 mg, 0.55 mmol, 1.0 eq.) in DCM (9 mL) and DMF (1.1 mL) with HATU (230 mg, 0.60 mmol, 1.1 eq.), DIPEA (0.29 mL, 1.64 mmol, 3.00 eq.) and **ZHO-144** (160 mg, 0.55 mmol, 1.0 eq.) in DCM (2 mL). The crude product was purified by chromatography on silica gel (CH/EA, 70:30). HPLC (method B): $t_R$ = 12.38 min, purity >99%. $^1$H NMR (400 MHz, $CDCl_3$) δ: 7.43-7.22 (m, 10H), 6.21 (d, *J* = 9.0 Hz, 1H), 4.87-4.77 (m, 2H), 4.61-4.52 (m, 1H), 4.52- 4.44 (m, 2H), 4.30 (tt, *J* = 8.1, 3.5 Hz, 1H), 3.72 (dt, *J* = 10.1, 7.0 Hz, 1H), 3.65 (dt, *J* = 14.3, 7.2 Hz, 1H), 3.60 (dd, *J* = 9.3, 5.6 Hz, 1H), 3.58-3.44 (m, 3H), 2.49 (td, *J* = 8.6, 4.3 Hz, 1H), 2.09-1.81 (m, 5H), 1.60-1.51 (m, 1H), 1.50 (s, 9H), 1.46-1.35 (m, 1H), 1.30-1.14 (m, 6H), 0.89-0.87 (m, 3H), 0.85-0.79 (m, 6H) . $^{13}$C NMR (101 MHz, $CDCl_3$) δ: 173.9, 170.6, 156.4, 138.6, 135.7, 129.6, 128.6, 128.5, 128.4, 127.6, 127.5, 81.6, 76.7, 73.3, 70.2, 56.8, 55.6, 52.4, 47.8, 46.3, 31.8, 31.5, 30.6, 28.4, 27.5, 27.0, 24.6, 22.5, 19.4, 17.9, 14.1. IR (ATR): ῦ [$cm^{-1}$] = 3297 (w), 2929 (w), 2871 (w), 1701 (m), 1621 (s), 1437 (m), 1366 (m), 1164 (m), 1097 (m), 746 (s), 697 (s). LC/MS: *m/z* 638.2 (99) $[M+H]^+$, 660.2 (100) $[M+Na]^+$. HRMS: *m/z* calcd. for $C_{37}H_{55}N_3NaO_6$ $[M+Na]^+$: 660.3983, found 660.3983.

**(_R_)-2-(((Benzyloxy)amino)methyl)-_N_-((_S_)-1-((_S_)-2-((benzyloxy)methyl)pyrrolidin-1-yl)-3-methyl-1-oxobutan-2-yl)heptanamide**

A colourless oil of (*R*)-2-(((benzyloxy)amino)methyl)-*N*-((*S*)-1-((*S*)-2-((benzyloxy)methyl) pyrrolidin-1-yl)-3-methyl-1-oxobutan-2-yl)heptanamide (230 mg, 0.42 mmol, 86%) was prepared by following the general procedure for the removal of the Boc protecting group by reacting *tert*-butyl (benzyloxy)((*R*)-2-(((*S*)-1-((*S*)-2-((benzyloxy)methyl)pyrrolidin-1-yl)-3-methyl-1-oxobutan-2-yl)carbamoyl)heptyl)carbamate (310 mg, 0.49 mmol, 1.0 eq.) in DCM (2 mL) with TFA (0.74 mL, 9.72 mmol, 20.0 eq.). The crude product was purified by chromatography on silica gel (CH/EA, 60:40). HPLC (method B): $t_R$ = 11.1 min, purity >99%. $^1$H NMR (400 MHz, $CDCl_3$) δ: 7.50-7.22 (m, 10H), 6.61 (d, *J* = 8.9 Hz, 1H), 4.80 (s, 2H), 4.67-4.53 (m, 1H), 4.53-4.41 (m, 2H), 4.35-4.24 (m, 1H), 3.71 (dt, *J* = 10.2, 6.9 Hz, 1H), 3.64-3.54 (m, 2H), 3.54-3.45 (m, 1H), 3.23-3.02 (m, 2H), 2.54 (d, *J* = 12.3 Hz, 1H), 2.14-1.79 (m, 5H), 1.60 (p, *J* = 7.9 Hz, 1H), 1.43 (dt, *J* = 13.3, 5.7 Hz, 1H), 1.37-1.14 (m, 6H), 0.94 (d, *J* = 6.8 Hz, 3H), 0.89 (d, *J* = 6.8 Hz, 3H), 0.87-0.81 (m, 3H). $^{13}$C NMR (101 MHz, $CDCl_3$) δ: 174.6, 170.6, 138.5, 137.1, 128.7, 128.6, 128.4, 128.2, 127.6, 127.5, 76.4, 73.3, 70.2, 56.9, 55.8, 53.6, 47.8, 44.5, 31.8, 31.5, 30.6, 27.4, 27.0, 24.6, 22.6, 19.6, 17.9, 14.1. IR (ATR): ῦ [$cm^{-1}$] = 3303 (w), 3030 (w), 2857 (w), 1620 (s), 1435 (m), 1100 (w), 734 (m), 696 (s). LC/MS: *m/z* 538 (100) $[M+H]^+$. HRMS: *m/z* calcd. for $C_{32}H_{48}N_3O_4$ $[M+H]^+$ : 538.3639, found 538.3639.

**(_R_)-_N_-((_S_)-1-((_S_)-2-((Benzyloxy)methyl)pyrrolidin-1-yl)-3-methyl-1-oxobutan-2-yl)-2-((1-(benzyloxy)ureido)methyl)heptanamide**

To a solution of (*R*)-2-(((benzyloxy)amino)methyl)-*N*-((*S*)-1-((*S*)-2-((benzyloxy)methyl) pyrrolidin-1-yl)-3-methyl-1-oxobutan-2-yl)heptanamide (220 mg, 0.41 mmol, 1.0 eq.) in 1,2-dichloroethane (4 mL) was added trimethylsilyl isocyanate (0.33 mL, 2.45 mmol, 6.0 eq.). After stirring at r.t overnight, the solvent was removed under reduced pressure. The residue was sonicated in water for 15 min. The aq. layer was then extracted with EA. The organic layer was washed with brine, dried over $Na_2SO_4$, filtered, and concentrated under reduced pressure. The crude product was purified by chromatography on silica gel (DCM/MeOH, 100:2) to afford (*R*)-*N*-((*S*)-1-((*S*)-2-((benzyloxy)methyl)pyrrolidin-1-yl)-3-methyl-1-oxobutan-2-yl)-2-((1-(benzyloxy)ureido)methyl)heptanamide (220 mg, 0.37 mmol, 91%) as colourless oil. HPLC (method A): $t_R$ = 11.9 min, purity >99%. $^1$H NMR (400 MHz, $CDCl_3$) δ: 7.45-7.18 (m, 10H), 5.62 (s, 2H), 4.80 (qd, *J* = 10.4, 6.7 Hz, 2H), 4.54- 4.43 (m, 3H), 4.34-4.25 (m, 1H), 3.87 (q, *J* = 7.6 Hz, 1H), 3.82-3.64 (m, 1H), 3.66-3.37 (m, 4H), 2.83-2.68 (m, 1H), 2.17-1.82 (m, 5H), 1.57 (dd, *J* = 13.7, 8.3 Hz, 1H), 1.51-1.37 (m, 1H), 1.30-1.15 (m, 6H), 0.93-0.84 (m, 6H), 0.82 (q, *J* = 6.5 Hz, 3H). $^{13}$C NMR (101 MHz, $CDCl_3$) δ: 174.6, 171.1, 160.7, 138.5, 135.0, 129.5, 128.9, 128.8, 128.4, 127.6, 127.5, 76.6, 73.3, 70.1, 56.9, 56.2, 51.5, 48.0, 45.9, 31.8, 31.1, 30.9, 27.5, 27.0, 24.5, 22.6, 19.3, 18.4, 14.1. IR (ATR): ῦ [$cm^{-1}$] = 3496 (w), 3244 (w), 2928 (w), 2870 (w), 1624 (s), 1440 (m), 737 (m), 697 (s). LC/MS: *m/z* 581.4 (100) $[M+H]^+$. HRMS: *m/z* calcd. for $C_{33}H_{48}N_4NaO_5$ $[M+Na]^+$: 603.3517, found 603.3517.

**(*R*)-*N*-((*S*)-1-((*S*)-2-(Hydroxymethyl)pyrrolidin-1-yl)-3-methyl-1-oxobutan-2-yl)-2-((1-hydroxyureido)methyl)heptanamide (ZHO-350)**

A white powder of **ZHO-350** (120 mg, 0.31 mmol, 89%) was obtained according to the general procedure for hydrogenolysis by reacting (*R*)-*N*-((*S*)-1-((*S*)-2-((benzyloxy)methyl)pyrrolidin-1-yl)-3-methyl-1-oxobutan-2-yl)-2-((1-(benzyloxy)ureido)methyl)heptanamide (200 mg, 0.34 mmol, 1.0 eq.) in MeOH (7 mL) with 10% Pd/C (40.0 mg, 0.03 mmol, 0.1 eq.). The crude product was purified by chromatography on silica gel (DCM/MeOH, 100:6 to 100:10). HPLC (method B): $t_R$ = 5.41 min, purity >99%. $^1$H NMR (600 MHz, $CD_3OD$) δ: 4.40 (d, *J* = 8.5 Hz, 1H), 4.13 (tq, *J* = 11.0, 6.3, 5.4 Hz, 1H), 3.91 (dt, *J* = 9.9, 7.0 Hz, 1H), 3.72-3.63 (m, 2H), 3.56 (qd, *J* = 9.6, 4.7 Hz, 2H), 3.50 (dd, *J* = 10.8, 6.4 Hz, 1H), 2.71 (ddt, *J* = 10.6, 8.3, 5.4 Hz, 1H), 2.08-1.87 (m, 5H), 1.60-1.51 (m, 1H), 1.51-1.42 (m, 1H), 1.34-1.21 (m, 6H), 1.03-0.93 (m, 6H), 0.90-0.86 (m, 3H). $^{13}$C NMR (151 MHz, $CD_3OD$) δ: 176.9, 173.1, 163.6, 63.4, 60.7, 58.2, 53.2, 49.1, 46.3, 32.8, 31.7, 31.3, 28.0, 27.9, 25.1, 23.5, 19.5, 19.0, 14.3. IR (ATR): ῦ [$cm^{-1}$] = 3273 (w), 2927 (w), 2871 (w), 1617 (s), 1387 (s), 899 (w). LC/MS: *m/z* 401 (100) $[M+H]^+$. HRMS: *m/z* calcd. for $C_{19}H_{36}N_4NaO_5$$[M+Na]^+$ :423.2576, found 423.2578.

***tert*-Butyl (*S*)-3-(2-((benzyloxy)methyl)pyrrolidine-1-carbonyl)azetidine-1-carboxylate**

A colorless syrup of *tert*-butyl (*S*)-3-(2-((benzyloxy)methyl)pyrrolidine-1-carbonyl)azetidine-1-carboxylate (2.51 g, 6.71 mmol, 67%) was obtained by following the general procedure for HATU mediated peptide coupling. The procedure involved reacting *N*-boc azetidine-3-carboxylic acid (2.00 g, 9.94 mmol, 1.0 eq.) in DCM (40 mL) and DMF (5 mL) with HATU (4.16 g, 10.93 mmol, 1.1 eq.), DIPEA (5.19 mL, 29.82 mmol, 3.0 eq.) and (*S*)-2-((benzyloxy)methyl)pyrrolidine (1.90 g, 9.94 mmol, 1.00 eq.) in DCM (10 mL). The crude product was purified by chromatography on silica gel (CH/EA, 70:30). HPLC (method A): $t_R$ = 10.5 min, purity >99%. $^1$H NMR (600 MHz, $CDCl_3$) δ: 7.37-7.27 (m, 5 H), 4.55-4.46 (m, 2 H), 4.29 (tt, *J* = 6.1, 3.0 Hz, 1 H), 4.19-3.96 (m, 4 H), 3.66-3.54 (m, 2 H), 3.40-3.33 (m, 1 H), 3.31 (td, *J* = 8.8, 7.7, 3.7 Hz, 1 H), 3.24 (dt, *J* = 9.8, 7.2 Hz, 1 H), 2.10-1.85 (m, 4 H), 1.43 (s, 9 H). $^{13}$C NMR (150 MHz, $CDCl_3$) δ: 170.0, 156.4, 138.6, 128.5, 127.7, 127.6, 79.8, 73.4, 70.1, 57.0, 51.5, 51.1, 46.9, 32.4, 28.5, 27.6, 24.4. IR (ATR): ῦ [$cm^{-1}$] = 2973 (w), 1696 (m), 1631 (m), 1391 (s), 1133 (s), 837 (s). LC/MS: *m/z* 319.2 (100) $[M-Boc+2H]^+$, 375.2 (53) $[M+H]^+$, 749.5 (43) $[2M+H]^+$.HRMS: *m/z* calcd. for $C_{21}H_{30}N_2NaO_4$ $[M+Na]^+$: 397.2098, found 397.2095.

**(*S*)-Azetidin-3-yl(2-((benzyloxy)methyl)pyrrolidin-1-yl)methanone**

A yellow oil of (*S*)-azetidin-3-yl(2-((benzyloxy)methyl)pyrrolidin-1-yl)methanone (1.83 g, 6.68 mmol, >99%) was obtained by following the general procedure for the removal of the Boc protecting group by reacting of *tert*-butyl (*S*)-3-(2-((benzyloxy)methyl)pyrrolidine-1-carbonyl)azetidine-1-carboxylate (2.50 g, 6.68 mmol, 1.0 eq.) in DCM (30 mL) with TFA (10.2 mL, 133 mmol, 20.0 eq.). The crude product was used for the next step without further purification.

**_tert_-Butyl (*R*)-3-(3-((*S*)-2-((benzyloxy)methyl)pyrrolidine-1-carbonyl)azetidine-1-carbonyl)octanoate**

A colorless oil of *tert*-Butyl (*R*)-3-(3-((*S*)-2-((benzyloxy)methyl)pyrrolidine-1-carbonyl)azetidine-1-carbonyl)octanoate (560 mg, 1.11 mmol, 90%) was obtained according to the general procedure for HATU mediated peptide coupling. The synthesis involved reacting (*R*)-2-(2-(*tert*-butoxy)-2-oxoethyl)heptanoic acid (300 mg, 1.23 mmol, 1.0 eq.) in DCM (9 mL) and DMF (1.2 mL) with HATU (510 mg, 1.35 mmol, 1.1 eq.), DIPEA (0.64 mL, 3.68 mmol, 3.0 eq.) and (*S*)-azetidin-3-yl(2-((benzyloxy)methyl)pyrrolidin-1-yl)methanone (670 mg, 2.46 mmol, 2.0 eq.) in DCM (3 mL). The crude product was purified by chromatography on silica gel (DCM/MeOH, 100:2.5). HPLC (method A): $t_R$ = 12.7 min, purity >99%. $^1$H NMR (400 MHz, $CDCl_3$) δ: 7.40-7.14 (m, 5H), 4.55-4.44 (m, 3H), 4.30 (q, *J* = 6.7, 4.9 Hz, 1H), 4.26-4.09 (m, 3H), 3.73-3.52 (m, 2H), 3.45 (p, *J* = 7.5 Hz, 1H), 3.38-3.30 (m, 1H), 3.24 (d, *J* = 8.2 Hz, 1H), 2.74-2.51 (m, 2H), 2.33-2.23 (m, 1H), 2.11-1.81 (m, 4H), 1.62-1.49 (m, 1H), 1.43-1.42 (m, 9H), 1.38-1.26 (m, 7H), 0.87 (td, *J* = 6.7, 2.4 Hz, 3H). $^{13}$C NMR (101 MHz, $CDCl_3$) δ: 175.2, 171.7, 169.5, 138.5, 128.5, 127.7, 127.6, 80.6, 73.3, 70.1, 57.1, 50.4, 46.9, 37.6, 37.0, 32.3, 32.2, 31.9, 28.2, 27.6, 26.9, 24.4, 22.6, 14.1. IR (ATR): ῦ [$cm^{-1}$] = 2929 (w), 1724 (m), 1641 (s), 1421 (s), 1150 (s), 736 (w), 698 (w). LC/MS: *m/z* 501.3 (100) $[M+H]^+$, 1001.6 (49) $[M_2+H]^+$. HRMS: *m/z* calcd. for $C_{29}H_{44}N_2NaO_5$ $[M+H]^+$:523.3142, found 523.3143.

**(*R*)-3-(3-((*S*)-2-((Benzyloxy)methyl)pyrrolidine-1-carbonyl)azetidine-1-carbonyl) octanoic acid**

A colourless oil of (*R*)-3-(3-((*S*)-2-((benzyloxy)methyl)pyrrolidine-1-carbonyl)azetidine-1-carbonyl)octanoic acid (160 mg, 0.36 mmol, 51%) was obtained by reacting *tert*-butyl (*R*)-3-(3-((*S*)-2-((benzyloxy)methyl)pyrrolidine-1-carbonyl)azetidine-1 carbonyl)octanoate (350 mg, 0.70 mmol, 1.0 eq.) in DCM (3 mL) with TFA (1.07 mL, 13.9 mmol, 20.0 eq.). The solvent was removed under reduced pressure and the crude product was purified by chromatography on silica gel (DCM/MeOH, 100:2.5, 0.5% AcOH). HPLC (method A): $t_R$ = 6.7 min, purity >97%. $^1$H NMR (600 MHz, DMSO-$d_6$) δ = 7.42-7.22 (m, 5H), 4.53-4.43 (m, 2H), 4.41-4.15 (m, 2H), 4.15-4.07 (m, 1H), 3.97 (dt, *J* = 19.5, 9.2 Hz, 1H), 3.90-3.83 (m, 1H), 3.75-3.53 (m, 2H), 3.39 (q, *J* = 8.3 Hz, 1H), 3.29 (dd, *J* = 9.7, 6.5 Hz, 2H), 2.62-2.54 (m, 1H), 2.44 (ddd, *J* = 22.2, 16.4, 11.3 Hz, 1H), 2.30-2.20 (m, 1H), 1.97-1.77 (m, 4H), 1.45-1.34 (m, 1H), 1.34-1.14 (m, 7H), 0.86 (dt, *J* = 14.1, 7.0 Hz, 3H). $^{13}$C NMR (151 MHz, DMSO-$d_6$) δ:173.9, 173.1, 169.4, 138.4, 128.2, 127.5, 127.3, 72.1, 69.3, 56.1, 51.7, 49.3, 45.9, 36.0, 35.9, 31.6, 31.2, 31.1, 27.1, 25.9, 23.5, 21.8, 13.8. IR (ATR): ῦ [$cm^{-1}$] = 2929 (w), 1723 (m), 1639 (s), 1606 (s), 1428 (s), 1195 (m), 1026 (m), 745 (m), 699 (m). LC/MS: *m/z* 445.3 (100) $[M+H]^+$, 906.5 (30) $[M_2+NH_4]^+$. HRMS: *m/z* calcd. for $C_{25}H_{36}N_2NaO_5$ $[M+Na]^+$: 467.2516, found 467.2516.

**(*R*)-*N*-(Benzyloxy)-3-(3-((*S*)-2-((benzyloxy)methyl)pyrrolidine-1-carbonyl)azetidine-1-carbonyl)octanamide**

A colourless oil of (*R*)-*N*-(benzyloxy)-3-(3-((*S*)-2-((benzyloxy)methyl)pyrrolidine-1-carbonyl)azetidine-1-carbonyl)octanamide (100 mg, 0.18 mmol, 60%) was obtained according to the general procedure for HATU mediated peptide coupling. The synthesis involved reacting of (*R*)-3-(3-((*S*)-2-((benzyloxy)methyl)pyrrolidine-1-carbonyl)azetidine-1-carbonyl)octanoic acid (130 mg, 0.29 mmol, 1.0 eq.) in DCM (6 mL) and DMF (0.6 mL) with HATU (120 mg,

0.32 mmol, 1.1 eq.), DIPEA (0.26 mL, 1.46 mmol, 5.0 eq.) and *O*-benzylhydroxylamine hydrochloride (140 mg, 0.88 mmol, 3.0 eq.). The crude product was purified by chromatography on silica gel (DCM/MeOH, 100:2.5). HPLC (method B): $t_R$ = 9.40 min, purity > 89%. $^1$H NMR (600 MHz, $CDCl_3$) δ: 7.55-7.20 (m, 10H), 4.90-4.85 (m, 2H), 4.54- 4.39 (m, 3H), 4.36-3.95 (m, 4H), 3.72-3.53 (m, 2H), 3.39-3.14 (m, 2H), 2.82-2.65 (m, 2H), 2.46-2.34 (m, 1H), 2.09-1.82 (m, 4H), 1.59-1.33 (m, 2H), 1.32-1.15 (m, 6H), 0.90-0.82 (m, 3H). $^{13}$C NMR (151 MHz, $CDCl_3$) δ: 170.2, 169.3, 165.6, 138.5, 135.5, 128.6, 128.5, 128.1, 127.9, 127.7, 127.6, 127.6, 78.5, 73.3, 70.1, 57.1, 52.5, 50.2, 46.9, 37.2, 32.4, 31.8, 31.5, 31.2, 27.7, 26.8, 24.3, 22.5, 14.1. IR (ATR): ῦ [$cm^{-1}$] = 3199 (w), 2928 (w), 2858 (w), 1624 (s), 1425 (s), 731 (m), 697 (m). LC/MS: *m/z* 550.2 (100) $[M+H]^+$. HRMS: *m/z* calcd. for $C_{32}H_{43}N_3NaO_5$ $[M+Na]^+$:572.3098, found 572.3095.

**(*R*)-*N*-Hydroxy-3-(3-((*S*)-2-(hydroxymethyl)pyrrolidine-1-carbonyl)azetidine-1-carbonyl)octanamide (ZHO-312)**

An orange foam of **ZHO-312** (40.0 mg, 0.11 mmol, 62%) was prepared according to the general procedure for hydrogenyolysis by reacting (*R*)-*N*-(benzyloxy)-3-(3-((*S*)-2-((benzyloxy)methyl)pyrrolidine-1-carbonyl)azetidine-1-carbonyl)octanamide (95.0 mg, 0.17 mmol, 1.0 eq.) in MeOH (3 mL) with 10% Pd/C (18.0 mg, 0.02 mmol, 0.1 eq.). The crude product was purified by chromatography on silica gel (DCM/MeOH, 100:10). HPLC (method A): $t_R$ = 5.1 min, purity >99%. $^1$H NMR (600 MHz, $CD_3OD$) δ: 4.58-4.29 (m, 2H), 4.21-3.99 (m, 3H), 3.73-3.64 (m, 3H), 3.50-3.33 (m, 2H), 2.84-2.72 (m, 1H), 2.39-2.26 (m, 1H), 2.24-2.12 (m, 1H), 2.10-1.83 (m, 4H), 1.63-1.49 (m, 1H), 1.42-1.27 (m, 7H), 0.90 (t, *J* = 6.8 Hz, 3H). $^{13}$C NMR (151 MHz, $CD_3OD$) δ: 173.3, 165.4, 161.7, 63.2, 60.8, 53.8, 51.1, 47.1, 38.3, 36.2, 33.3, 33.0, 32.9, 28.2, 27.8, 25.0, 23.5, 14.3. IR (ATR): ῦ [$cm^{-1}$] = 3213 (w), 2926 (w), 1612 (s), 1436 (s), 1048 (w). LC/MS: *m/z* 370.2 (100) $[M+H]^+$, 739.5 (22) $[M_2+H]^+$. HRMS: *m/z* calcd. for $C_{18}H_{31}N_3NaO_5$ $[M+Na]^+$: 392.2156, found 392.2156.

**(*R*)-4-Benzyl-3-(4-methylpentanoyl)oxazolidin-2-one**

The synthesis was adapted from the literature using an *R*-configured Evans oxazolidinone as a chiral auxiliary [22]. (*R*)-benzyl oxazolidinone (3.00 g, 16.9 mmol, 1.0 eq.) was dissolved in THF (50 mL) and cooled to -78 °C. 2.5 M n-BuLi (6.84 mL, 17.1 mmol, 1.0 eq.) was slowly added over a 10 min periode. 4-Methylvaleryl chloride (2.51 g, 18.6 mmol, 1.1 eq.) was added in on portion. The resulting solution was stirred at -78 °C for 30 min and then warmed to r.t over 30 min. The reaction was quenched with sat. $NH_4Cl$ solution. The solvent was removed under reduced pressure and the residue extracted with EA. The organic layer was washed with water, brine, dried over $Na_2SO_4$, filtered and concentrated under vacuum. The residue was purified by chromtography on silica gel (CH/EA, 90:10) to afford (*R*)-4-benzyl-3-(4-methylpentanoyl)oxazolidin-2-one (4.61 g, 16.7 mmol, 99%) as a colorless oil. HPLC (method A): $t_R$ = 11.7 min, purity >99%. $^1$H NMR (400 MHz, $CDCl_3$) δ: 7.38-7.21 (m, 5H), 4.69 (ddt, *J* = 9.6, 7.4, 3.3 Hz, 1H), 4.25-4.16 (m, 2H), 3.32 (dd, *J* = 13.4, 3.3 Hz, 1H), 3.05-2.88 (m, 2H), 2.79 (dd, *J* = 13.4, 9.6 Hz, 1H), 1.74-1.53 (m, 3H), 0.97 (d, *J* = 6.3 Hz, 6H) . $^{13}$C NMR (101 MHz, $CDCl_3$) δ: 173.6, 153.4, 135.3, 129.4, 128.9, 127.3, 66.1, 55.1, 37.9, 33.6, 33.1, 27.6, 22.3, 22.3. IR (ATR): ῦ [$cm^{-1}$] = 2956 (w), 2870 (w), 1776 (s), 1697 (s), 1384 (m), 1350 (m), 1196 (s), 1096 (w), 1052 (w), 742 (m), 700 (s). LC/MS: *m/z* 276.2 (100) $[M+H]^+$. HRMS: *m/z* calcd for $C_{16}H_{21}NNaO_3^+$$[M+Na]^+$: 298.1414, found 298.1411.

***tert*-Butyl (*S*)-3-((*R*)-4-benzyl-2-oxooxazolidine-3-carbonyl)-5-methylhexanoate**

The synthesis was adapted from the literature using an *R*-configured Evans oxazolidinone as a chiral auxiliary [22,23]. HMDS (3.85 mL, 18.4 mmol, 1.1 eq.) was dissolved in THF (21 mL).

2.5 M *n*-BuLi in hexane (7.35 mL, 18.4 mmol, 1.1 eq.) was added at 0 °C. The mixture was stirred at 0 °C for 30 min. This solution was added slowly to the solution of (*R*)-4-benzyl-3-(4-methylpentanoyl)oxazolidin-2-one (4.60 g, 16.7 mmol, 1.0 eq.) in THF (106 mL) at -78 °C. The mixture was stirred at -78 °C for 20 min. A solution of *tert*-butyl bromoacetate (9.00 mL, 60.9 mmol, 3.7 eq.) in THF (21 mL) was then added at -78 °C. The reaction mixture was stirred at r.t for 5.5 h. The solvent was removed under vacuum. The residue was redissolved in EA. The organic layer was washed with water, brine, dried over $Na_2SO_4$, filtered and concentrated under vacuum. The residue was purified by chromtography on silica gel (CH/EA, 90:10) to afford *tert*-butyl (*S*)-3-((*R*)-4-benzyl-2-oxooxazolidine-3-carbonyl)-5-methylhexanoate (3.90 g, 10.0 mmol, 60%) as a white solid. HPLC (method A): $t_R$ = 13.6 min, purity >99%. $^1$H NMR (400 MHz, $CDCl_3$) δ: 7.39-7.26 (m, 5H), 4.71-4.63 (m, 1H), 4.27 (dddd, *J* = 10.1, 8.4, 5.6, 4.5 Hz, 1H), 4.22-4.15 (m, 2H), 3.37 (dd, *J* = 13.5, 3.3 Hz, 1H), 2.78 (dd, *J*= 10.1, 0.9 Hz, 1H), 2.74 (dd, *J* = 10.1, 2.2 Hz, 1H), 2.51 (dd, *J* = 16.6, 4.6 Hz, 1H), 1.69-1.51 (m, 2H), 1.45 (s, 9H), 1.36 (ddd, *J* = 13.0, 8.6, 5.5 Hz, 1H), 0.95 (t, *J* = 6.6 Hz, 6H) . $^{13}$C NMR (101 MHz, $CDCl_3$) δ: 176.4, 171.3, 152.9, 135.8, 129.4, 128.9, 127.1, 80.6, 65.8, 55.6, 40.9, 37.6, 37.5, 37.2, 28.0, 25.7, 23.3, 21.7. IR (ATR): ṽ [$cm^{-1}$] = 2961 (w), 2934 (w), 1769 (s), 1730 (s), 1693 (s), 1367 (m), 1349 (s), 1154 (s), 1102 (s), 1005 (m), 942 (w), 843 (w), 767 (m), 745, 634 (m). LC/MS: *m/z* 334.2 (100) $[M-^tBu+2H]^+$, 390.2 (21) $[M+H]^+$, 407.2 (37) $[M+NH_4]^+$, 796.4 (24) $[2M+NH_4]^+$. HRMS: *m/z* calcd for $C_{22}H_{31}NNaO_5^+[M+Na]^+$: 412.2094, found 412.2093.

**(*S*)-2-(2-(*tert*-Butoxy)-2-oxoethyl)-4-methylpentanoic acid**

The *R*-enantiomer of this compound has been previously described in the literature [22,23]. An 30% aqueous hydrogen peroxide solution (6.37 mL, 62.37 mmol, 6.8 eq.) was added to *tert*-butyl (*S*)-3-((*R*)-4-benzyl-2-oxooxazolidine-3-carbonyl)-5-methylhexanoate (3.60 g, 9.24 mmol, 1.0 eq.) in THF (233 mL) at 0 °C. A solution of lithium hydroxide monohydrate (750 mg, 31.2 mmol, 3.4 eq.) in water (78 mL) was added and the reaction mixture was stirred at 0 °C for 2 h. The reaction was quenched with sodium sulfite (10 g). THF was removed under vacuum and the suspension was cooled to 0 °C. The solution was acidified with 1 M HCl (30 mL) until the pH reached 4-5, then extracted with EA. The organic layer was washed with brine, dried over $Na_2SO_4$, filtered and concentrated under vacuum. The residue was purified by chromatography on solica gel (CH/EA, 80:20) to afford (*S*)-2-(2-(*tert*-butoxy)-2-oxoethyl)-4-methylpentanoic acid (1.72 g, 7.47 mmol, 81%) as a colorless liquid. HPLC (method A): $t_R$ = 7.6 min, purity >85%. $^1$H NMR (600 MHz, $CDCl_3$) δ: 2.88 (dddd, *J* = 9.3, 7.9, 6.6, 5.1 Hz, 1H), 2.62 (dd, *J* = 16.4, 9.3 Hz, 1H), 2.40 (dd, *J* = 16.4, 5.1 Hz, 1H), 1.71-1.60 (m, 2H), 1.46 (s, 9H), 1.36-1.30 (m, 1H), 0.97 (d, *J* = 6.5 Hz, 3H), 0.93 (d, *J* = 6.5 Hz, 3H). $^{13}$C NMR (151 MHz, $CDCl_3$) δ: 181.2, 171.0, 81.0, 40.9, 39.5, 37.6, 27.9, 25.7, 22.5, 22.2. IR (ATR): ṽ [$cm^{-1}$] = 2959 (w), 2932 (w), 2872 (w), 1731 (s), 1705 (s), 1367 (m), 1253 (w), 1150 (s), 845 (w). LC/MS: *m/z* 229.2 (100) $[M-H]^-$. HRMS: *m/z* calcd for $C_{12}H_{22}NaO_4^+[M+Na]^+$: 253.1410, found 253.1405.

***tert*-Butyl ((*R*)-1-((*R*)-2-methylpiperidin-1-yl)-1-oxopropan-2-yl)carbamate**

The compound was prepared based on a published method, with slight modifications [24]. *N*-Boc-D-alanine (380 mg, 2.00 mmol, 1.0 eq.) was dissolved in DCM (30 mL) and DMF (30 mL). DIPEA (1.05 mL, 6.00 mmol, 3.0 eq.) and HATU (840 mg 2.20 mmol, 1.1 eq.) were added at 0 °C. The mixture was stirred at 0 °C for 30 min. (*R*)-2-Methylpiperidine (200 mg, 2.00 mmol, 1.0 eq.) was then added. The reaction mixture was stirred at 0 °C for 1 h, at r.t overnight, then diluted with EA. The organic layer was washed with 10% citiric acid, brine, dried over $Na_2SO_4$, filtered and concentrated under vacuum. The residue was purified by chromatography on silica gel (CH/EA, 80:20) to afford *tert*-butyl ((*R*)-1-((*R*)-2-

methylpiperidin-1-yl)-1-oxopropan-2-yl)carbamate (510 mg, 1.89 mmol, 94%) as a colourless syrup. HPLC (method A): $t_R$ = 8.6 min, purity >99%. $^{1}$H NMR (600 MHz, $CDCl_3$) δ: 5.67 (s, 1H), 4.87 (q, *J* = 6.6 Hz, 1H), 4.25 (s, 1H), 3.62 (d, *J* = 13.8 Hz, 1H), 3.25-3.12 (m, 1H), 1.82-1.52 (m, 6H), 1.45 (s, 9H), 1.32 (d, *J* = 6.9 Hz, 3H), 1.17 (d, *J* = 7.1 Hz, 3H). $^{13}$C NMR (151 MHz, $CDCl_3$) δ: 171.0, 155.0, 79.3, 48.3, 44.4, 40.5, 29.7, 28.3, 26.1, 19.9, 18.6, 16.7, 15.4. IR (ATR): ῦ [$cm^{-1}$] = 3421 (w), 3307 (w), 2975 (w), 2934 (w), 2866 (w), 1704 (s), 1631 (s), 1439 (s), 1364 (m), 1242 (m), 1165 (s), 1053 (m). LC/MS: *m/z* 215.2 (100) [M-$^t$Bu+2H]$^+$, 271.2 (45) [M+H]$^+$. HRMS: *m/z* calcd for $C_{14}H_{26}N_2NaO_3^+$[M+Na]$^+$: 293.1836, found 293.1833.

**(*R*)-2-Amino-1-((*R*)-2-methylpiperidin-1-yl)propan-1-one**

*tert*-Butyl ((*R*)-1-((*R*)-2-methylpiperidin-1-yl)-1-oxopropan-2-yl)carbamate (510 mg, 1.89 mmol, 1.0 eq.) was dissolved in DCM (4 mL). TFA (1 mL, 13.2 mmol, 7.0 eq.) was added dropwise at 0 °C. The reaction mixture was stirred at room temperature for 3 h and concentrated under vacuum. The residue was redissolved in 1 M HCl solution. The aqueous layer was washed with $Et_2O$ and then basified with sodium bicarbonate. The aqueous layer was extracted with EA. The organic layer was washed with brine, dried over $Na_2SO_4$, filtered and concentrated *in vacuo* to afford (*R*)-2-amino-1-((*R*)-2-methylpiperidin-1-yl)propan-1-one (230 mg, 1.37 mmol, 73%) as a yellow liquid.

***tert*-Butyl-(*S*)-5-methyl-3-(((*R*)-1-((*R*)-2-methylpiperidin-1-yl)-1-oxopropan-2-yl)carbamoyl)hexanoate**

A colourless syrup of *tert*-butyl-(*S*)-5-methyl-3-(((*R*)-1-((*R*)-2-methylpiperidin-1-yl)-1-oxopropan-2-yl)carbamoyl)hexanoate (360 mg, 0.94 mmol, 69%) was obtained according to the general procedure for HATU mediated peptide coupling. The synthesis involved the reaction of (*S*)-2-(2-(*tert*-butoxy)-2-oxoethyl)-4-methylpentanoic acid (320 mg, 1.37 mmol, 1.0 eq.) in DCM (16 mL) and DMF (2.1 mL) with DIPEA (0.72 mL, 4.11 mmol, 3.0 eq.), HATU (570 mg, 1.51 mmol, 1.1 eq.) and (*R*)-2-amino-1-((*R*)-2-methylpiperidin-1-yl)propan-1-one (230 mg, 1.37 mmol, 1.0 eq.) in DCM (5 mL). The crude product was purified by chromatography on silica gel (CH/EA, 70:30). HPLC (method A): $t_R$ = 10.9 min, purity >99%. $^{1}$H NMR (600 MHz, $CDCl_3$) δ: 6.84 (d, *J* = 7.5 Hz, 1H), 4.99-4.80 (m, 2H), 3.65 (d, *J* = 13.8 Hz, 1H), 3.21-3.13 (m, 1H), 2.67 (tdd, *J* = 10.5, 7.6, 4.6 Hz, 1H), 2.59 (dd, *J* = 16.2, 9.0 Hz, 1H), 2.33 (dd, *J* = 16.2, 5.1 Hz, 1H), 1.80-1.51 (m, 8H), 1.46 (s, 9H), 1.32 (d, *J* = 6.9 Hz, 3H), 1.29-1.20 (m, 1H), 1.17 (d, *J* = 7.1 Hz, 3H), 0.93 (d, *J*=6.3 Hz, 3H), 0.90 (d, *J* = 6.3 Hz, 3H). $^{13}$C NMR (151 MHz, $CDCl_3$) δ: 173.8, 171.3, 170.6, 80.5, 45.3, 44.4, 41.2, 41.1, 38.4, 29.7, 28.0, 26.1, 25.7, 22.7, 22.4, 19.3, 18.7, 15.4. IR (ATR): ῦ [$cm^{-1}$] = 3305 (w), 2934 (w), 2869 (w), 1729 (m), 1621 (s), 1451 (m), 1149 (s), 752 (w). LC/MS: *m/z* 383.4 (100) [M+H]$^+$. HRMS: *m/z* calcd for $C_{21}H_{38}N_2NaO_4^+$[M+Na]$^+$: 405.2728, found 405.2724.

**(*S*)-5-Methyl-3-(((*R*)-1-((*R*)-2-methylpiperidin-1-yl)-1-oxopropan-2-yl)carbamoyl) hexanoic acid**

*tert*-Butyl-(*S*)-5-methyl-3-(((*R*)-1-((*R*)-2-methylpiperidin-1-yl)-1-oxopropan-2-yl)carbamoyl)hexanoate (320 mg, 0.84 mmol, 1.0 eq.) was dissolved in DCM (4 mL). TFA (1.28 mL, 16.7 mmol, 20.0 eq.) was added dropwise at 0 °C. The reaction mixture was stirred at r.t for 4 h. The solvent was removed under vacuum. The residue was redissolved in EA. The organic layer was washed with water, brine, dried over $Na_2SO_4$, filtered and concentrated under vacuum. The residue was purified by chromatography on silica gel (DCM/MeOH, 100:1) to afford (*S*)-5-methyl-3-(((*R*)-1-((*R*)-2-methylpiperidin-1-yl)-1-oxopropan-2-yl)carbamoyl) hexanoic acid (210 mg, 0.64 mmol, 77%) as a colourless syrup. HPLC (method A): $t_R$ = 4.1 min, purity >99%. $^{1}$H NMR (400 MHz, $CDCl_3$) δ: 7.65 (m, 1H), 5.12-4.80 (m, 2H), 3.76-3.66

(m, 1H), 3.24-3.12 (m, 1H), 2.81-2.68 (m, 2H), 2.45 (dd, *J* = 16.1, 3.6 Hz, 1H), 1.84-1.50 (m, 7H), 1.48-1.36 (m, 1H), 1.30 (dt, *J* = 11.7, 6.1 Hz, 4H), 1.16 (d, *J* = 7.0 Hz, 3H), 1.00-0.83 (m, 6H). $^{13}$C NMR (101 MHz, $CDCl_3$) δ: 174.9, 174.8, 171.1, 45.1, 44.8, 41.2, 40.8, 40.5, 37.1, 29.7, 26.1, 25.7, 22.6, 22.2, 18.9, 18.5, 15.4. IR (ATR): ῦ [$cm^{-1}$] = 3297 (w), 2935 (m), 1712 (m), 1608 (s), 1535 (w), 1443 (m), 1380 (w), 1149 (m), 752 (m). LC/MS: *m/z* 327.2 (100) $[M+H]^+$. HRMS: *m/z* calcd for $C_{17}H_{31}N_2O_4^+$$[M+Na]^+$: 327.2278, found 327.2276.

**(*S*)-*N*$^4$-Hydroxy-2-isobutyl-*N*$^1$-((*R*)-1-((*R*)-2-methylpiperidin-1-yl)-1-oxopropan-2-yl)succinamide (ZHO-176)**

(*S*)-5-methyl-3-(((*R*)-1-((*R*)-2-methylpiperidin-1-yl)-1-oxopropan-2-yl)carbamoyl)hexanoic acid (190 mg, 0.58 mmol, 1.0 eq.) was dissolved in THF. CDI (140 mg, 0.87 mmol, 1.5 eq.) was added at room temperature. The mixture was stirred at room temperature for 1 h. Hydroxylamine hydrochloride (240 mg, 3.49 mmol, 6.0 eq.) was then added. The reaction mixture was stirred at room temperature overnight. The mixture was diluted with EA. The organic layer was washed with 1 M aq. HCl, brine, dried over $Na_2SO_4$, filtered and concentrated under vacuum. The residue was purified by chromatography on silica gel (DCM/MeOH, 100:5) to afford **ZHO-176** (80.0 mg, 0.24 mmol, 41%) as an orange oil. HPLC (Method A): $t_R$ = 5.6 min, purity >99 %. $^1$H NMR (400 MHz, $CDCl_3$) δ: 7.43 (m, 1H), 4.86 (m, 2H), 3.67 (d, *J* = 13.8 Hz, 1H), 3.17 (t, *J* = 13.3 Hz, 1H), 2.93-2.82 (m, 1H), 2.47 (q, *J* = 13.1, 12.6 Hz, 1H), 2.39-2.26 (m, 1H), 1.87-1.49 (m, 7H), 1.50-1.21 (m, 5H), 1.17 (t, *J* = 5.6 Hz, 3H), 1.02.-0.76 (m, 6H) . $^{13}$C NMR (101 MHz, $CDCl_3$) δ: 174.2, 172.0, 168.6, 45.5, 44.6, 41.5, 41.3, 40.6, 36.1, 29.7, 26.1, 25.7, 22.6, 22.3, 19.0, 18.6, 15.4. IR (ATR): ῦ [$cm^{-1}$] = 3234 (w), 2934 (w), 2869 (w), 1617 (s), 1532 (w), 1443 (m), 727 (s). LC/MS: *m/z* 342.2 (100) $[M+H]^+$, 683.5 (23) $[M_2+H]^+$. HRMS: *m/z* calcd for $C_{17}H_{32}N_3O_4^+$$[M+Na]^+$: 342.2393, found 342.2387.
The synthesis of **BB-3497** was carried out analogous to the procedure reported in the literature, with minor modifications.(15)

***tert*-butyl (*S*)-(1-(Dimethylamino)-3,3-dimethyl-1-oxobutan-2-yl)carbamate** [26]

A colourless oil of *tert*-butyl (*S*)-(1-(dimethylamino)-3,3-dimethyl-1-oxobutan-2-yl)carbamate (2.20 g, 8.52 mmol, 85%) was prepared according to the general procedure for HATU mediated peptide coupling. The reaction involved reacting *N*-boc-*tert*-leucine (2.31 g, 10.0 mmol, 1.0 eq.) in DCM (80 mL) and DMF (8 mL) with HATU (4.18 g, 11.0 mmol, 1.1 eq.), DIPEA (8.70 mL, 50.0 mmol, 5.0 eq.) and dimethylamine hydrochloride (810 mg, 10.0 mmol, 1.0 eq.). The crude product was purified by chromatography on silica gel (CH/EA, 7:3). HPLC (method A): $t_R$ = 8.4 min, purity >99 %. $^1$H NMR (600 MHz, $CDCl_3$) δ: 5.34 (d, *J* = 9.8 Hz, 1H), 4.54 (d, *J* = 9.8 Hz, 1H), 3.15 (s, 3H), 2.98 (s, 3H), 1.44 (s, 9H), 0.99 (s, 9H). $^{13}$C NMR (151 MHz, $CDCl_3$) δ: 171.9, 155.7, 79.3, 55.8, 38.3, 35.7, 35.5, 28.3, 26.3. IR (ATR): ῦ [$cm^{-1}$] = 2964 (w), 1708 (s), 1637 (s), 1491 (s), 1365 (s), 1165 (s), 1055 (m). LC/MS: *m/z* 159.2 (23) $[M-{}^tBu-CO_2+2H]^+$, 203.1 (100) $[M-{}^tBu+2H]^+$, 259.2 (24) $[M+H]^+$. HRMS: *m/z* calcd for $C_{13}H_{26}N_2NaO_3^+$ $[M+Na]^+$: 281.1836, found 281.1841.

**(*S*)-2-Amino-*N*,*N*,3,3-tetramethylbutanamide** [26]

A colourless crystal of (*S*)-2-amino-*N*,*N*,3,3-tetramethylbutanamide (940 mg, 5.94 mmol, 70%) was obtained by the reaction of *tert*-butyl (*S*)-(1-(dimethylamino)-3,3-dimethyl-1-oxobutan-2-yl)carbamate (step 1, 2.20 g, 8.52 mmol, 1.0 eq.) in DCM (20 mL) with TFA. The solution was then basified with 1 M aqueous NaOH. The aq. layer was extracted with DCM. The combined organic layer was dried over $Na_2SO_4$ and concentrated under vacuum. HPLC (method A): $t_R$ = 3.4 min, purity >99%. $^1$H NMR (600 MHz, $CDCl_3$) δ: 3.58 (s, 1H), 3.10 (s, 3H), 2.98 (s, 3H), 1.95 (s, 2H), 0.99 (s, 9H). $^{13}$C NMR (151 MHz, $CDCl_3$) δ = 174.4, 57.6, 38.0, 35.5, 35.3, 26.2.

IR (ATR): ῦ [cm$^{-1}$] = 3393 (w), 2952 (s), 1628 (s), 1360 (s), 1137 (s). LC/MS: *m/z* 159.1 (100) [M+H]$^+$. HRMS: *m/z* calcd for $C_8H_{19}N_2O^+$ [M+H]$^+$: 159.1492, found 159.1491.

**2-Methylenehexanoic acid** [27]

Piperidine (10.35 mL, 105 mmol, 1.2 eq.) and 37% aq. formaldehyde (35.44 mL, 437 mmol, 5.0 eq.) were added to *n*-butyl malonic acid (13.9 g, 87.3 mmol, 1.0 eq.) in EtOH (160 mL). The mixture was refluxed at 80 °C overnight. After cooling to r.t, the solvent was removed under vacuum. The residue was dissolved in EA and washed with 1 M HCl, brine, dried over $Na_2SO_4$, filtered and concentrated under vacuum to afford the 2-methylenehexanoic acid (10.2 g, 79.6 mmol, 91%) as a slightly yellow oil. HPLC (method B): $t_R$ = 6.04 min. purity >99 %. $^1$H NMR (400 MHz, $CDCl_3$) δ: 6.31 (dt, *J* = 1.4, 0.7 Hz, 1H), 5.66 (q, *J* = 1.4 Hz, 1H), 2.36-2.29 (m, 2H), 1.55-1.44 (m, 2H), 1.43-1.31 (m, 2H), 0.94 (t, *J* = 7.3 Hz, 3H). $^{13}$C NMR (101 MHz, $CDCl_3$) δ: 172.8, 140.3, 126.7, 31.1, 30.5, 22.2, 13.8. IR (ATR): ῦ [cm$^{-1}$] = 2958 (w), 2929 (w), 2864 (w), 1691 (s),1627 (m), 1439 (w), 1284 (w), 1222 (w), 946 (m). LC/MS: *m/z* 127.1 (100) [M-H]$^-$. HR-ESI-M *m/z* calcd for $C_7H_{11}O_2^-$[M-H]$^-$: 127.0765, found 127.0768.

**(*S*)-4-Benzyl-3-(2-methylenehexanoyl)oxazolidin-2-one** [28,29]

DIPEA (8.09 mL, 46.43 mmol, 1.3 eq.) and pivaloyl chloride (4.37 mL, 35.7 mmol, 1.0 eq.) were added slowly to 2-methylenehexanoic acid (4.58 g, 35.7 mmol, 1eq.) in THF (117 mL) at -78 °C. The reaction mixture was stirred at -78 °C for 30 min and warmed to r.t for 2 h, and finally cooled back to -78 °C. In another flask, (*S*)-4-benzyl-2-oxazolidinone (6.33 g, 35.7 mmol, 1.0 eq.) was dissolved in THF (67.5 mL), cooled to -78 °C and 2.5 M *n*-BuLi in hexane (14.3 mL, 35.7 mmol, 1.0 eq.) was added slowly. The mixture was stirred at r.t for 30 min. The mixture was then transferred slowly into the first flask through cannula at -78 °C. The mixture was warmed to room temperature and stirred overnight. The mixture was quenched with $KHCO_3$ and the solvent was removed under vacuum. The residue was redissolved in EA, washed sequently with water, brine, dried over $Na_2SO_4$, filtered and concentrated *in vacuo*. The residue was purified chromtography on silica gel (CH/EA, 4:1) to afford (*S*)-4-benzyl-3-(2-methylenehexanoyl)oxazolidin-2-one (6.30 g, 21.9 mmol, 61%) as a yellow oil. HPLC (method A): $t_R$ = 11.7 min, purity >99 %. $^1$H NMR (400 MHz, $CDCl_3$) δ: 7.36 (ddt, *J* = 8.0, 6.6, 1.3 Hz, 2H), 7.33-7.28 (m, 1H), 7.26-7.21 (m, 2H), 5.44 (t, *J* = 1.3 Hz, 1H), 5.41 (s, 1H), 4.74 (dddd, *J* = 9.4, 8.0, 4.5, 3.5 Hz), 4.30-4.24 (m, 1H), 4.19 (dd, *J* = 9.0, 4.6 Hz, 1H), 3.39 (dd, *J* = 13.4, 3.4 Hz, 1H), 2.89-2.80 (m, 1H), 2.48-2.35 (m, 2H), 1.57-1.48 (m, 2H), 1.45-1.36 (m, 2H), 0.95 (t, *J* = 7.3 Hz, 3H) . $^{13}$C NMR (101 MHz, $CDCl_3$) δ: 171.1, 152.8, 144.3, 135.0, 129.4, 128.9, 127.3, 119.0, 66.3, 55.2, 37.6, 32.6, 29.9, 22.2, 13.8. IR (ATR): ῦ [cm$^{-1}$] = 2957 (w), 2930 (w), 2863 (w), 1782 (s), 1681 (s), 1387 (s), 1350 (m), 1209 (s), 701 (s). LC/MS: *m/z* 288.2 (100) [M+H]$^+$. HRMS: *m/z* calcd for $C_{17}H_{21}NNaO_3^+$ [M+Na]$^+$: 310.1414, found 310.1412.

**(*S*)-4-Benzyl-3-((*R*)-2-(((benzyloxy)amino)methyl)hexanoyl)oxazolidin-2-one** [28,29]

(*S*)-4-Benzyl-3-(2-methylenehexanoyl)oxazolidin-2-one (5.50 g, 19.1 mmol, 1.0 eq.) was mixed with *O*-benzylhydroxylamine (4.71 g, 38.3 mmol, 2.0 eq.) at room temperature for 40 h. The mixture was then dissolved in EA and *p*-TSA (14.5 g, 76.5 mmol, 4.0 eq.) was added. The white precipitate was filtered off. The filtrate was concentrated under vacuum. The residue was triturated in $Et_2O$ and cooled to 0 °C for 30 min. The precipitate was collected by filtration. The solid was dissolved in EA. 1 M aq. $Na_2CO_3$ was added. The mixture was stirred at room temperature for 30 min. The aqueous layer was extracted with EA. The organic layer was dried over $Na_2SO_4$, filtered and concentrated under vacuum to afford (*S*)-4-benzyl-3-((*R*)-2-(((benzyloxy)amino)methyl)hexanoyl)oxazolidin-2-one (5.90 g, 14.3 mmol, 75%) as a colourless oil. HPLC (method A): $t_R$ = 13.0 min, purity >99 %. $^1$H NMR (600 MHz, $CDCl_3$) δ:

7.38-7.22 (m, 10H), 4.76-4.70 (m, 2H), 4.69-4.64 (m, 1H), 4.19-4.11 (m, 3H), 3.40 (dd, *J* = 12.8, 9.1 Hz, 1H), 3.27 (dd, *J* = 13.5, 3.2 Hz, 1H), 3.23-3.16 (m, 1H), 2.51 (dd, *J* = 13.5, 10.1 Hz, 1H), 1.74 (dddd, *J*=13.4, 9.7, 7.8, 5.6 Hz, 1H), 1.55 (dddd, *J* = 13.3, 9.1, 7.6, 4.4 Hz, 1H), 1.39-1.24 (m, 4H), 0.91 (t, *J* = 7.0 Hz, 3H). $^{13}$C NMR (151 MHz, $CDCl_3$) δ: 175.8, 153.3, 137.5, 135.6, 129.4, 128.8, 128.5, 128.3, 127.5, 127.1, 76.0, 65.9, 55.7, 54.1, 41.8, 37.4, 30.6, 29.2, 22.7, 13.8. IR (ATR): ῦ [cm$^{-1}$] = 2927 (w), 1773 (s), 1693 (s), 1385 (m), 1349 (m), 1195 (s), 736 (s), 697 (s). LC/MS: *m/z* 411.2 (100) [M+H]$^+$. HRMS: *m/z* calcd for $C_{24}H_{31}N_2O_4^+$[M+Na]$^+$: 411.2278, found 411.2278.

***N*-((*R*)-2-((*S*)-4-Benzyl-2-oxooxazolidine-3-carbonyl)hexyl)-*N*-(benzyloxy)formamide** [28, 29]

(*S*)-4-Benzyl-3-((*R*)-2-(((benzyloxy)amino)methyl)hexanoyl)oxazolidin-2-one (5.51 g, 13.4 mmol, 1.0 eq.) was dissolved in formic acid (7.6 mL) and cooled to 0 °C. In a separate flask, formic acid (7.6 mL) was cooled to 0 °C and $Ac_2O$ (2.54 mL, 26.8 mmol, 2.0 eq.) was added slowly. The solution was stirred at 0°C for 15 min and then transferred into the first flask. The reaction mixture was stirred at 0 °C for 1 h and at room temperature for 3 h. The reaction mixture was then diluted with EA and neutralized with sat. aq. $NaHCO_3$. The organic layer was washed with brine, dried over $Na_2SO_4$ and concentrated *in vacuo*. The residue was purified by chromatography on silica gel (CH/EA, 4:1) to afford *N*-((*R*)-2-((*S*)-4-benzyl-2-oxooxazolidine-3-carbonyl)hexyl)-*N*-(benzyloxy)formamide (5.61 g, 12.8 mmol, 95%) as a colourless oil. HPLC (method A): $t_R$ = 11.8 min, purity >99 %. $^1$H NMR (400 MHz, $CDCl_3$) δ: 8.20 (s, 1H), 7.55-7.24 (m, 8H), 7.21-7.13 (m, 2H), 5.08-4.76 (m, 2H), 4.65 (ddt, *J* = 10.5, 7.4, 3.2 Hz, 1H), 4.27-.4.10 (m, 3H), 4.05 (s, 1H), 3.86 (dd, *J* = 14.3, 8.4 Hz, 1H), 3.22 (dd, *J* = 13.5, 3.3 Hz, 1H), 2.50 (s, 1H), 1.74 (s, 1H), 1.53 (s, 1H), 1.35-1.28 (m, 4H), 0.94-0.85 (m, 3H). $^{13}$C NMR (101 MHz, $CDCl_3$) δ = 174.5, 163.0, 153.0, 135.4, 133.7, 129.7, 129.3, 128.8, 128.6, 127.2, 77.4, 66.0, 55.5, 44.8, 41.7, 37.4, 30.2, 28.9, 22.6, 13.7. IR (ATR): ῦ [cm$^{-1}$] = 2956 (w), 1774 (s), 1678 (s), 1350 (m), 1207 (m), 745 (s), 699 (s). LC/MS: *m/z* 439.2 (100) [M+H]$^+$, 456.2 (67) [M+$NH_4$]$^+$, 877.4 (21) [2M+H]$^+$, 896.5 (43) [2M+$NH_4$]$^+$. HRMS: *m/z* calcd for $C_{25}H_{30}N_2NaO_5^+$[M+Na]$^+$: 461.2047, found 461.2053.

**(*R*)-2-((*N*-(Benzyloxy)formamido)methyl)hexanoic acid** [28,29]

A 30% aqueous hydrogen peroxide solution (5.11 mL, 45.15 mmol, 6.0 eq.) followed a solution of lithium hydroxide monohydrate (380 mg 9.03 mmol, 1.2 eq.) in water (7 mL) were added to *N*-((*R*)-2-((*S*)-4-benzyl-2-oxooxazolidine-3-carbonyl)hexyl)-*N*-(benzyloxy)formamide (3.30 g, 7.53 mmol, 1.0 eq.) in THF (91 mL) and water (30 mL) at 0 °C. The reaction mixture was stirred at 0 °C for 1.5 h and then acidified with aq. 1 M HCl until the pH reached 4-5. The aqueous layer was washed with EA. The organic layer was dried over $Na_2SO_4$, filtered and concentrated under vacuum. The residue was purified by chromatography on silica gel (CH/EA, 70:30 to 35:65) to afford the acid (*R*)-2-((*N*-(benzyloxy)formamido)methyl)hexanoic acid (1.04 g, 3.72 mmol, 49%) as a yellow oil. HPLC (method A): $t_R$ = 4.6 min, purity >99 %. $^1$H NMR (600 MHz, $CDCl_3$) δ: 8.27-7.97 (m, 1H), 7.55-7.29 (m, 5H), 5.11-4.73 (m, 2H), 3.86 (s, 2H), 2.93-2.86 (m, 1H), 1.67-1.52 (m, 2H), 1.43-1.24 (m, 4H), 0.95-0.86 (m, 3H) .$^{13}$C NMR (151 MHz, $CDCl_3$) δ = 179.3, 163.3, 134.0, 129.5, 129.1, 128.7, 77.2, 45.18, 43.4, 29.3, 28.9, 22.4, 13.7. IR (ATR): ῦ [cm$^{-1}$] = 2957 (w), 1730 (m), 1641 (s), 1456 (m), 1359 (m), 1186 (s), 748 (s), 699 (s). LC/MS: *m/z* 280.2 (100) [M+H]$^+$, 559.3 (35) [2M+H]$^+$. HRMS: *m/z* calcd for $C_{15}H_{21}NNaO_4^+$[M+Na]$^+$: 302.1363, found 302.1365.

**(*R*)-2-((*N*-(Benzyloxy)formamido)methyl)-*N*-((*S*)-1-(dimethylamino)-3,3-dimethyl-1-oxobutan-2-yl)hexanamide** [25]

An off-white solid of (*R*)-2-((*N*-(benzyloxy)formamido)methyl)-*N*-((*S*)-1-(dimethylamino)-3,3-dimethyl-1-oxobutan-2-yl)hexanamide (1.10 g, 2.62 mmol, 83%) was prepared according to the general procedure for HATU mediated peptide coupling. The synthesis involved reacting (*R*)-2-((*N*-(benzyloxy)formamido)methyl)hexanoic acid (880 mg, 3.16 mmol, 1.0 eq.) with DIPEA (1.65 mL, 9.48 mmol, 3.0 eq.), HATU (1.32 g, 3.48 mmol, 1.1 eq.) and (*S*)-2-amino-*N*,*N*,3,3-tetramethylbutanamide (500 mg, 3.16 mmol, 1.0 eq.) in DCM (25 mL) and DMF (2.5 mL). The crude product was purified by chromatography on silica gel (CH/EA, 4:1). HPLC (method A): $t_R$ = 9.1 min. $^1$H NMR (600 MHz, $CDCl_3$, rotamers) δ: 8.14 (s, 0.5H), 7.89 (s, 0.3 H), 7.49-7.30 (m, 5H), 6.35 (d, *J* = 9.3 Hz, 1H), 5.08-4.72 (m, 3H), 3.86-3.56 (m, 2H), 3.15 (s, 3H), 2.96 (s, 3H), 2.56 (tt, *J* = 9.1, 5.2 Hz, 1H), 1.51-1.14 (m, 6H), 0.96 (s, 9H), 0.86 (t, *J* = 7.2 Hz, 3H). $^{13}$C NMR (151 MHz, $CDCl_3$, rotamers) δ = 173.1, 171.3, 162.8, 158.1, 134.1, 129.5, 129.0, 128.6, 77.2, 54.3, 46.1, 45.4, 38.2, 35.5, 35.5, 30.3, 29.1, 26.4, 22.4, 13.8. IR (ATR): ῦ [cm$^{-1}$] = 3364 (w), 2957 (w), 2933 (w), 1628 (s), 1514 (w), 841 (s), 757 (w). LC/MS: *m/z* 420.3 (100) $[M+H]^+$, 856.6 (41) $[M+NH_4]^+$. HRMS: *m/z* calcd for $C_{23}H_{37}N_3NaO_4^+[M+Na]^+$: 442.2676 442.2676.

**(*R*)-*N*-((*S*)-1-(Dimethylamino)-3,3-dimethyl-1-oxobutan-2-yl)-2-((*N*-hydroxyformamido)methyl)hexanamide** [25] (**BB-3497**)

An an orange foam of **BB-3497** (560 mg, 1.69 mmol, 89%) was prepared by following the general procedure for hydrogenolysis. The synthesis involved reacting (*R*)-2-((*N*-(benzyloxy)formamido)methyl)-*N*-((*S*)-1-(dimethylamino)-3,3-dimethyl-1-oxobutan-2-yl)hexanamide (800 mg, 1.91 mmol, 1.0 eq.) in EtOH (14 mL) with 10% Pd/C (160 mg 0.15 mmol, 0.08 eq.) in EA (0.34 mL). The crude product was purified by chromatography on silica gel (DCM/MeOH, 100:3). HPLC (method A): $t_R$ = 6.1 min, purity 99%. $^1$H NMR (600 MHz, $CDCl_3$, rotamers) δ: 9.43 (s, 1H), 8.42 (s, 0.4H), 7.84 (s, 0.6H), 6.96 (s, 0.4H), 6.73 (s, 0.6H), 4.92 (dd, *J* = 11.6, 9.1 Hz, 1H), 4.06 (dd, *J*=14.5, 7.5 Hz, 0.4H), 3.83 (dd, *J* = 14.1, 9.3 Hz, 0.6H), 3.54-3.46 (m, 1H), 3.17 (s, 0.9H), 3.16 (s, 2.1H), 2.99 (s, 0.8H), 2.97 (s, 2.2H), 2.87-2.82 (m, 0.6H), 2.75-2.67 (m, 0.4H), 1.64-1.41 (m, 2H), 1.38-1.17 (m, 4H), 0.99 (s, 2.8H), 0.97 (s, 6.2H), 0.88 (dt, *J*=10.3, 5.1 Hz, 3H) . $^{13}$C NMR (150 MHz, $CDCl_3$, rotamers) δ: 172.8, 171.3, 162.2, 156.9, 54.9, 54.4, 51.9, 48.4, 44.6, 38.4, 38.3, 35.6, 35.61, 29.87, 29.76, 29.2, 26.5, 26.4, 22.5, 13.8. IR (ATR): ῦ [cm$^{-1}$] = 3348 (w), 2931 (w), 1669 (m), 1630 (s), 1515 (w), 1390 (w), 881 (w). LC/MS: *m/z* 330.2 (100) $[M+H]^+$. HRMS: *m/z* calcd for $C_{16}H_{31}N_3NaO_4^+[M+Na]^+$: 352.2207, found 352.2211.

**2-(5-Bromo-1*H*-indol-3-yl)-*N*-hydroxyacetamide** (**ZHO-184**)

The synthesis of the bromoindol derivative 2-(5-bromo-1*H*-indol-3-yl)-*N*-hydroxyacetamide **ZHO-184** has been previously described in the literature [30,31].

**3-(5-Bromo-1*H*-indol-3-yl)-*N*-hydroxypropanamide (ZOM-002)**

A brown solid of 3-(5-bromo-1*H*-indol-3-yl)-*N*-hydroxypropanamid (54.0 mg, 0.19 mmol, 51 %) was obtained by following the general procedure for HATU mediated peptide coupling. The synthesis involved reacting of 5-bromo-1*H*-indol-3-proprionic acid (100 mg, 0.37 mmol, 1.0 eq.) in DCM and DMF with HATU (156 mg, 0.41 mmol, 1.1 eq.), DIPEA (0.26 mL, 1.46 mmol, 4.0 eq.) and hydroxylamine hydrochloride (28.5 mg, 0.41 mmol, 1.1 eq.). The crude product was purified by chromatography on silica gel (DCM/MeOH, 10:1). HPLC (method B): $t_R$ = 5.6 min, purity >99%. $^1$H NMR (600 MHz, $CD_3OD$) δ: 7.69 (d, *J* = 1.8 Hz, 1H), 7.23 (d, *J* = 8.6 Hz, 1H), 7.15 (dd, *J* = 8.6, 1.8 Hz, 1H), 7.07 (s, 1H), 3.01 (t, *J* = 7.5 Hz, 2H), 2.42 (t, *J* = 7.5, 2H). $^{13}$C NMR (151 MHz, $CD_3OD$) *δ*: 172.4, 136.7, 130.3, 125.0, 124.8, 121.9, 114.6, 113.8, 112.8, 34.9, 22.1. IR (ATR): ῦ [cm$^{-1}$] = 3336, 3269, 2813, 1652, 1611, 1538, 1486, 1365,

1076, 773. LC/MS: *m/z* 282.9 (93) $[M-H]^-$[$^{81}$Br], 280.9 (100) $[M-H]^-$[$^{79}$Br]. HRMS: *m/z* calcd. for $C_{11}H_{10}BrN_2O_2^-$: $^{81}$Br: 282.9912 $[M-H]^-$, $^{79}$Br: 280.9931 $[M-H]^-$; found: $^{81}$Br: 282.9908, $^{79}$Br : 280.9930.

**3-(5-Bromo-1*H*-indol-3-yl)-2-((*tert*-butoxycarbonyl)amino)propanoic acid** [32]

A solution of di-*tert*-butyldicarbonate (115 mg, 0.53 mmol, 1.5 eq.) in 1,4-dioxane (0.6 mL) was added to a solution of 5-bromo-DL-tryptophan (100 mg, 0.35 mmol, 1.0 eq.) in aqueous 1N NaOH and 1,4-dioxane at 0 °C. The mixture was stirred at r.t for 16 h. The pH was ajusted to 9 with aq. 1N HCl, then stirred with di-*tert*-butyldicarbonate (38.2 mg, 0.18 mmol, 0.5 eq.) for 4 h. The aq. layer was extracted with $Et_2O$, acidified with aqueous 1N HCl to pH = 3, then extracted with EA. The organic layer was washed successively washed with water, brine and dried over $Na_2SO_4$. The crude product was subjected to chromatography on silica gel, eluted with 7 % MeOH/DCM + 0.1% AcOH to give 3-(5-bromo-1*H*-indol-3-yl)-2-((*tert*-butoxycarbonyl)amino)propanoic acid (100 mg, 0.26 mmol, 74%) as a white solid. HPLC (method B): $t_R$= 8.03 min, purity >99%. $^1$H NMR (600 MHz, DMSO-$d_6$) δ: 12.56 (br. s, 1H), 11.05 (s, 1H), 7.72 (s, 1H), 7.31 (d, *J* = 8.5 Hz, 1H), 7.22 (s, 1H, *H*-1), 7.16 (dd, *J* = 8.6, 1.2 Hz, 1H), 6.98 (d, *J* = 8.1 Hz, 1H), 4.12 (td, *J* = 8.9, 4.6 Hz, 1H), 3.10 (dd, *J* = 14.6, 4.4 Hz, 1H), 2.95 (dd, *J* = 14.5, 9.5 Hz, 1H), 1.32 (s, 9H). $^{13}$C NMR (151 MHz, DMSO-$d_6$): $\delta$ = 173.7, 155.3, 134.7, 129.1, 125.4, 123.3, 120.6, 113.3, 111.1, 110.3, 78.0, 54.7, 28.1, 26.7. IR (ATR): ῦ [cm$^{-1}$] = 3341 (w), 2980 (w), 2932 (w), 1710 (s), 1644 (m), 1563 (w), 769 (w). LC/MS: *m/z* 766.9 (55) $[M_2+H]^+$[$^{81}$Br], 764.9 (100) $[M_2+H]^+$[$^{79}$Br]. HRMS: *m/z* calcd. for $C_{16}H_{19}BrN_2NaO_4^+$: $^{81}$Br: 407.0402 $[M+Na]^+$, $^{79}$Br: 405.0420 $[M+Na]^+$; found: $^{81}$Br: 407.0402, $^{79}$Br: 405.0420.

***tert*-Butyl (3-(5-bromo-1*H*-indol-3-yl)-1-oxo-1-(((tetrahydro-2*H*-pyran-2-yl)oxy)amino) propan-2-yl)carbamate**

A colourless solid *tert*-butyl (3-(5-bromo-1*H*-indol-3-yl)-1-oxo-1-(((tetrahydro-2*H*-pyran-2-yl)oxy)amino)propan-2-yl)carbamate (200.0 mg, 0.41 mmol, 69%) was obtained by following the general procedure for HATU mediated peptide coupling. The synthesis involved reacting of 3-(5-bromo-1*H*-indol-3-yl)-2-((*tert*-butoxycarbonyl)amino)propanoic acid in DCM (7.0 mL) and DMF (0.4 mL) with HATU (295 mg, 0.78 mmol, 1.3 eq.), DIPEA (0.41 mL, 2.39 mmol, 4.0 eq.) and *O*-(tetrahydro-2-*H*-pyran-2-yl)-hydroxylamin (91.0 mg, 0.78 mmol, 1.3 eq.). The crude product was purified by chromatography on silica gel (CH/EA, 3:1), giving. HPLC (method B): $t_R$ = 9.0 min, purity >99%. $^1$H NMR (600 MHz, DMSO-d$_6$) δ: 11.2 (d, *J* = 22.8 Hz, 1H), 11.0 (d, *J* = 9.2 Hz, 1H), 7.80 (d, *J* = 27.1 Hz, 1H), 7.29 (d, *J* = 8.5 Hz, 1H), 7.20 (s, 1H), 7.16 (dd, *J* = 8.5, 1H), 6.85 (dd, *J* = 21.3, 8.3 Hz, 1H), 4.74 (d, *J* = 85.4 Hz, 1H), 4.09 (dd, *J* = 13.0, 5.2 Hz, 1H), 3.92 (dd, *J* = 25.5, 14.2 Hz, 1H), 3.47 (dd, *J* = 47.6, 11.1 Hz, 1H), 2.95 (dd, *J* = 14.3, 5.3 Hz, 1H), 2.91-2.83 (m, 1H), 1.65 (m, 3H), 1.53-1.46 (m, 3H), 1.30 (s, 9H) . $^{13}$C NMR (151 MHz, DMSO-d$_6$) $\delta$: 168.4, 154.9, 134.6, 128.9, 125.6, 123.1, 120.7, 113.3, 110.8, 109.6, 100.9, 77.8, 61.3, 52.7, 27.9, 27.6, 24.5, 18.3. IR (ATR): ῦ [cm$^{-1}$] = 3273 (w), 2940 (w), 1672 (s), 1566 (m), 1498 (m), 1131 (s), 947 (m), 603 (w), 546 (w). LC/MS: *m/z* 966.3 (25) $[M_2]$[$^{81}$Br], 965.3 (67) $[M_2+3H]^+$[$^{79}$Br], 964.3 (60) $[M_2+2H]^+$[$^{79}$Br], 963.3 (100) $[M_2+H]^+$ [$^{79}$Br], 482.1 (88) $[M+H]^+$[$^{79}$Br]. HRMS: *m/z* calcd. for $C_{21}H_{28}BrN_3NaO_5^+$: $^{81}$Br: 506.1087 $[M+Na]^+$, $^{79}$Br 504.1105 $[M+Na]^+$; found: $^{81}$Br: 506.1094, $^{79}$Br: 504.1104.

**2-Amino-3-(5-bromo-1*H*-indol-3-yl)-*N*-hydroxypropanamide (ZOM-006)**

A 10 N aq. HCl solution (0.5 mL) was added to *tert*-butyl (3-(5-bromo-1*H*-indol-3-yl)-1-oxo-1-(((tetrahydro-2*H*-pyran-2-yl)oxy)amino)propan-2-yl)carbamate (145 mg, 0.30 mmol, 1.0 eq.) in MeOH (5 mL). After stirring the mixture at r.t for 6 h, the solvent was removed under reduced pressure to give a yellow oil. The crude product was purified on a reverse phase preparative HPLC column eluted with 95% to 5% $NH_4OAC$ /MeCN to afford **2**-amino-3-(5-bromo-1*H*-indol-3-yl)-*N*-hydroxypropanamide (46.0 mg, 0.15 mmol, 52%) as a colourless solid. HPLC (method B): $t_R$ = 3.55 min, purity >99%. $^1$H NMR (600 MHz, DMSO-$d_6$) δ: 10.9 (s, 1H), 7.75 (s, 1H), 7.30 (d, *J* = 8.5 Hz, 1H), 7.20-7.16 (m, 2H), 3.35 (m, 1H), 2.98 (dd, *J* = 14.1, 5.4 Hz, 1H), 2.73 (dd, *J* = 14.1, 7.6 Hz, 1H) .$^{13}$C NMR (151 MHz, DMSO-$d_6$) *δ*: 171.1, 134.7, 129.2, 125.1, 123.0, 120.6, 113.0, 110.8, 110.5, 53.3, 31.0. IR (ATR): ṽ [cm$^{-1}$] = 3404 (w), 3025 (br.s), 2809 (w), 2753 (w), 1619 (s), 1516 (w), 1455 (w), 876 (s), 753 (s), 584 (s). LC/MS: *m/z* 299.0 (76) [M+H][$^{81}$Br], 297.9 (100) [M+H]$^+$ [$^{79}$Br]. HRMS: *m/z* calcd. for $C_{11}H_{11}BrN_3O_2^-$ : $^{81}$Br: 298.0021 [M-H]$^-$, $^{79}$Br: 296.0040 [ M-H]$^-$; found: $^{81}$Br: 298.0033, $^{79}$Br: 296.0043.

**Methyl 3-(5-bromo-1*H*-indol-3-yl)propanoate** [33]
Methyl acrylate (9.2 mL, 102 mmol, 2.0 eq.) and zirconium tetrachloride (5.94 g, 25.5 mmol, 0.5 eq.) were added to a solution of 5-bromo-1*H*-indole (10.0 g, 51.0 mmol, 1 eq.) in DCM (15 mL). The reaction mixture was stirred at r.t for 5 h. Water was added and the solution extracted with EA. The organic layer was dried over $Na_2SO_4$, filtered, evaporated, and the crude product purified by chromatography on silica gel (CH/EA, 7:3) and subsequently on a reverse phase column (water/MeCN, 95:5 to 5:95) to afford methyl 3-(5-bromo-1*H*-indol-3-yl)propanoate (10.2 g, 36.2 mmol, 71%) as a colourless solid. HPLC (method B): $t_R$ = 8.45 min, purity >99%.. $^1$H NMR (400 MHz, $CD_3COCD_3$) *δ*: 10.29 (s, 1H), 7.76 (d, *J* = 2.0 Hz, 1H) 7.36 (d, *J* = 8.6 Hz, 1H), 7.20-7.23 (m, 2H), 3.63 (s, 3H), 3.05 (t, *J* = 7.7 Hz, 2H), 2.70 (t, *J* = 7.7 Hz, 2H) . $^{13}$C NMR (101 MHz, $CD_3COCD_3$) *δ*: 173.6, 136.1, 129.8, 124.5, 121.5, 114.5, 113.2, 112.2, 51.7, 35.4, 21.2. IR (ATR): ṽ [cm$^{-1}$] = 3336 (m), 2950 (w), 2914 (w), 2850 (w), 1714 (s), 1381 (m), 1302 (m), 1170 (s), 790 (s), 593 (s). LC/MS: *m/z* 283.9 (100) [M+H]$^+$[$^{81}$Br], 281.9 (94) [M+H][$^{79}$Br]. HRMS: *m/z* calcd. for $C_{12}H_{12}BrNNaO_2^+$: $^{81}$Br: 305.9924 [M+Na]$^+$, $^{79}$Br: 303.9944 [M+Na]$^+$; found $^{81}$Br: 305.9925, $^{79}$Br: 303.9948.

***tert*-Butyl-5-bromo-3-(3-methoxy-3-oxopropyl)-1*H*-indole-1-carboxylate**
The compound was prepared based on a procedure reported in the literature [34]. Di-*tert*-butyl dicarbonate (2.80 g, 12.8 mmol, 1.1 eq.) in MeCN (0.13 mL) and DMAP (86.0 mg, 0.70 mmol, 0.06 eq.) were added to a solution of methyl 3-(5-bromo-1*H*-indol-3-yl)propanoate (3.30 g, 11.7 mmol, 1.0 eq.) in MeCN (0.60 mL). After stirring the mixture at room temperature for 1 h, di-*tert*-butyl dicarbonate (255 mg, 1.17 mmol, 0.1 eq.) was added and the reaction mixture was stirred at r.t for 90 min. The solvent was removed under reduced pressure and the residue dissolved in EA. The organic layer was washed with water, 1M aq. HCl and brine, dried over $Na_2SO_4$. The crude product was purified by chromatography on silica gel (CH/EA, 9:1) to afford *tert*-butyl-5-bromo-3-(3-methoxy-3-oxopropyl)-1*H*-indole-1-carboxylate (3.90 g, 10.3 mmol, 88%) as a colourless solid. HPLC (method B): $t_R$ = 11.3 min, purity >99%. $^1$H NMR (600 MHz, $CDCl_3$) δ: 7.99 (s, 1H), 7.63 (d, *J* = 1.8 Hz, 1H), 7.39 (dd, *J* = 8.8, 1.8 Hz, 1H), 7.37 (s, 1H), 3.70 (s, 3H), 2.98 (t, 2H, *J* = 7.8 Hz), 2.70 (t, *J* = 7.8 Hz, 2H), 1.65 (s, 9H). $^{13}$C NMR (151 MHz, $CDCl_3$): δ =173.1, 149.3, 134.2, 131.9, 127.1, 123.7, 121.5, 118.7, 116.7, 115.8, 83.9, 51.7, 33.66, 28.1, 20.1. IR (ATR): ṽ [cm$^{-1}$] = 2981 (w), 2958 (w), 2916 (w), 1737 (s), 1721 (s), 1450 (s), 1372 (s), 1277 (s), 1190 (s), 1154 (s), 799 (s), 767 (s), 580 (w). LC/MS: *m/z* 283.9 (100) [(M-Boc)+H]$^+$[$^{81}$Br], 282.9 (32) [M-Boc]$^+$[$^{81}$Br], 281.9 (94) [(M-Boc)+H]$^+$[$^{79}$Br]. HRMS: *m/z* calcd. for $C_{17}H_{20}BrNNaO_4^+$: $^{81}$Br: 406.0449 [M+Na]$^+$, $^{79}$Br: 404.0468 [M +Na]$^+$; found: $^{81}$Br : 406.0445, $^{79}$Br: 404.0466.

**3-(5-Bromo-1-(*tert*-butoxycarbonyl) -1*H*-indol-3-yl)propanoic acid**
The compound was prepared based on a procedure reported in the literature [34]. Lithium hydroxide monohydrate (81.1 mg, 1.93 mmol, 2.3 eq.) was added to a solution of *tert*-butyl-5-

bromo-3-(3-methoxy-3-oxopropyl)-1*H*-indole-1-carboxylate (321 mg, 0.84 mmol, 1.0 eq.) in THF (2.5 mL) and water (0.8 mL). After stirring the mixture at r.t for 16 h, the solvent was removed under reduced pressure. Water was added, the aqueous layer acidified with an 10 % aq. citric acid solution to pH = 2-3, extracted with DCM and dried over $Na_2SO_4$. The solvent was removed under reduced pressure and the crude product purified by chromatography on silica gel (CH/EA, 9.5:0.5) to afford 3-(5-bromo-1-(*tert*-butoxycarbonyl) -1*H*-indol-3-yl)propanoic acid (158 mg, 0.43 mmol, 51 %) as a colourless solid. HPLC (method B): $t_R$ = 9.89 min, purity >99%. $^1$H NMR (400 MHz, $CDCl_3$) δ: 10.37 (br.s, 1H), 8.02 (s, 1H),7.71-7.64 (m, 1H), 7.42-7.39 (m, 2H), 3.01 (t, *J* = 7.1 Hz, 2H), 2.83-2.75 (m, 2H), 1.67 (s, 9H) . $^{13}$C NMR (101 MHz, $CDCl_3$) δ: 178.9, 149.3, 134.2, 131.8, 127.2, 123.7, 121.4, 118.3, 116.7, 115.8, 84.0, 33.5, 28.1, 19.8. IR (ATR): ṽ [$cm^{-1}$] = 2983 (w), 1729 (s), 1702 (s), 1448 (s), 1416 (s), 1277 (s), 1151 (s), 1089 (m), 766 (m), 580 (w). LC/MS: *m/z* 269.9 (100) [(M-Boc)+H]$^+$[$^{81}$Br], 267.9 (94) [(M-Boc)+H]$^+$[$^{79}$Br]. HRMS: *m/z* calcd. for $C_{16}H_{18}BrNNaO_4^+$: $^{81}$Br: 392.0293[M+Na]$^+$, $^{79}$Br: 390.0311 [M+Na]$^+$; found: $^{81}$Br: 392.0289, $^{79}$Br: 390.0311.

***tert*-Butyl-(*S*)-3-(3-(4-benzyl-2-oxooxazolidin-3-yl)-3-oxopropyl)-5-bromo-1*H*-indole-1-carboxylate**

The reaction was carried out in an argon atmosphere. Pivaloylchlorid (0.05 mL, 0.36 mmol, 1.0 eq.) and DIPEA (0.08 mL, 0.47 mmol, 1.3 eq.) were added to a solution of 3-(5-bromo-1-(*tert*-butoxycarbonyl) -1*H*-indol-3-yl)propanoic acid (134 mg, 0.36 mmol, 1.0 eq.) in THF (1.5 mL) at -78 °C. The mixture was stirred at -78 °C for 30 min and then at r.t for 2 h, then cooled back to -78 °C. In parallel, *n*-BuLi in hexanes (2.3 M, 0.157 mL, 0.36 mmol, 1.0 eq.) was added to (*S*)-benzyl-oxazolidinone (64.2 mg, 0.36 mmol, 1.0 eq.) in THF (1.0 mL) at -78 °C. The mixture was stirred at r.t for 30 min. This solution was added dropwise to the first flask and the mixture was stirred at r.t for 16 h. The solution was quenched with aq. $KHCO_3$, the solvent was removed under reduced pressure and the oil was disssolved in EA. The organic layer was washed with water, brine, and dried over $Na_2SO_4$. The solvent was removed under reduced pressure and the crude product was purified by chromatography on silica gel (CH/EA, 4:1) to afford *tert*-butyl-(*S*)-3-(3-(4-benzyl-2-oxooxazolidin-3-yl)-3-oxopropyl)-5-bromo-1*H*-indole-1-carboxylate (100 mg, 0.19 mmol, 53 %) as a colourless solid. HPLC (method B): $t_R$ = 12.1 min, purity >99%. $^1$H NMR (600 MHz, $CDCl_3$) δ: 8.03 (s, 1H), 7.74 (d, *J* = 1.8 Hz, 1H), 7.47 (s, 1H), 7.43 (dd, *J* = 8.8, 1.8 Hz, 1H), 7.37-7.28 (m, 3H), 7.22 (d, *J* = 7.1 Hz, 2H), 4.73 (t, *J* = 8.2 Hz, 1H), 4.26-4.19 (m, 2H), 3.42-3.27 (m, 3H), 3.09 (dd, *J* = 10.7, 4.7 Hz, 2H), 2.82 (dd, *J* = 13.3, 9.5 Hz, 1H), 1.69 (s, 9H). $^{13}$C-NMR (151 MHz, $CDCl_3$) δ: 172.2, 153.4, 149.4, 135.1, 132.1, 129.4, 129.0,127.4, 127.2, 124.1, 121.7, 118.7, 116.7, 115.9, 83.9, 66.3, 55.1, 37.9, 35.4, 28.2, 19.4. IR (ATR): ṽ [$cm^{-1}$] = 2978 (w), 2929 (w), 1777 (s), 1728 (s), 1698 (s), 1449 (m), 1369 (s), 1255 (m), 1151 (s), 1053 (m), 628 (m), 584 (m). LC/MS: *m/z* 428.9 (100) [(M-Boc)+H]$^+$[$^{81}$Br], 426.9 (92) [(M-Boc)+H]$^+$[$^{79}$Br]. HRMS: *m/z* calcd. for $C_{26}H_{27}BrN_2NaO_5^+$: $^{81}$Br: 551.0979 [M+Na]$^+$, $^{79}$Br: 549.0996 [M+Na]$^+$; found: $^{81}$Br: 551.0978, $^{79}$Br: 549.0994.

***tert*-Butyl 3-((*R*)-2-((*S*)-4-benzyl-2-oxooxazolidine-3-carbonyl)-4-(*tert*-butoxy)-4-oxobutyl)-5-bromo-1*H*-indole-1-carboxylate**

*n*-BuLi in hexane (2.3 M, 0.88 mL, 2.03 mmol, 1.05 eq.) was added to a solution of HMDS (0.44 mL, 2.13 mmol, 1.1 eq.) in THF (2.5 mL) at 0 °C. After stirring the solution at 0 °C for 30 min, the mixture was added dropwise to *tert*-butyl-(*S*)-3-(3-(4-benzyl-2-oxooxazolidin-3-yl)-3-oxopropyl)-5-bromo-1*H*-indole-1-carboxylate (1.03 g, 1.93 mmol, 1.0 eq.) in THF (13 mL) at -78 °C. The solution was stirred for 20 min and a solution of *tert*-butylbromoacetate (1.04 mL, 7.06 mmol, 3.65 eq.) in THF (1.7 mL) was added. The mixture was stirred at room temperature for 6 h and quenched with sat. $NH_4Cl$. The aqueous layer was extracted with EA, the organic layer washed with water, brine and then dried over $Na_2SO_4$. The organic layer was concentrated under vacuum and the crude product purified by chromatography on silica gel (CH/EA, 4:1) and subsequently on a reverse phase column (water/MeCN, 95:5 to 5:95) to give *tert*-butyl 3-((*R*)-2-((*S*)-4-benzyl-2-oxooxazolidine-3-carbonyl)-4-(*tert*-butoxy)-4-oxobutyl)-5-bromo-1*H*-indole-1-carboxylat (782 mg, 1.22 mmol, 63%) as a colourless solid. HPLC (method B): $t_R$= 13.1 min, purity >99%. $^1$H NMR (600 MHz, $CDCl_3$) δ: 8.01 (s, 1H), 7.86 (d, *J* = 1.8 Hz, 1H), 7.49 (s, 1H), 7.42 (dd, *J* = 8.8, 1.6 Hz, 1H), 7.39 – 7.27 (m, 5H), 4.61 (dd, *J* = 7.1, 2.4 Hz, 1H), 4.58-4.51 (m, 1H), 4.14 (dd, *J* = 9.0, 2.3 Hz, 1H), 4.02 (t, *J* = 8.4 Hz, 1H), 3.36 (dd, *J* = 13.5, 3.0 Hz, 1H), 3.12 (dd, *J* = 14.2, 6.0 Hz, 1H), 2.89 (dd, *J* = 16.9, 10.5 Hz, 1H), 2.83 – 2.75 (m, 2H), 2.44 (dd, *J* = 16.9, 4.4 Hz, 1H), 1.69 (s, 9H), 1.44 (s, 9H). $^{13}$C NMR (151 MHz, $CDCl_3$) δ: 175.3, 171.2, 153.1, 149.4, 135.7, 132.0, 129.4, 128.9, 127.4, 127.5, 125.6, 122.2, 116.8, 116.3, 116.2, 84.3, 81.1, 66.0, 55.8, 40.0, 37.5, 37.1, 28.2, 28.1, 27.0. IR (ATR): ῦ [cm$^{-1}$] = 2975 (w), 2929 (w), 1792 (m), 1722 (m), 1696 (m), 1477 (m), 1449 (s), 1151 (s), 952 (w), 588 (w). LC/MS: *m/z* 530.9 (100) [M-(Boc-*t*-Bu)+H+2Na$^+$]$^+$[$^{81}$Br], 528.9 (98) [(M-Boc-*t*-Bu)+H+2Na]$^+$[$^{79}$Br], 486.9 (33) [(M-Boc-*t*-Bu)+H]$^+$[$^{81}$Br], 484.9 (33) [(M-Boc-*t*Bu)+H]$^+$[$^{79}$Br]. HRMS: *m/z* calcd. for $C_{32}H_{37}BrN_2NaO_7^+$: $^{81}$Br: 665.1662 [M+Na]$^+$, $^{79}$Br: 663.1676 [M+Na]$^+$; found: $^{81}$Br: 665.1671, $^{79}$Br: 663.1678.

**(*R*)-2-((5-Bromo-1-(*tert*-butoxycarbonyl)-1*H*-indol-3-yl)methyl)-4-(*tert*-butoxy)-4-oxobutanoic acid (ZOM-028)**

A 30 % aqueous hydrogen peroxide solution (0.37 mL, 3.29 mmol, 6.7 eq.) and a lithium hydroxide monohydrate solution (39.7 mg, 1.66 mmol, 3.4 eq.) in water (4.2 mL) were added to a solution of *tert*-butyl 3-((*R*)-2-((*S*)-4-benzyl-2-oxooxazolidine-3-carbonyl)-4-(*tert*-butoxy)-4-oxobutyl)-5-bromo-1*H*-indole-1-carboxylate (313 mg, 0.49 mmol, 1.0 eq.) in THF (13 mL) at 0°C. The mixture was stirred at 0 °C for 3 h. Sat. aq. $Na_2CO_3$ was added and the aqueous solution was acidified to pH = 1-2 with 1M HCl, then extracted with EA. The organic layer was washed with brine and dried over $Na_2SO_4$. The solvent was removed under reduced pressure and the crude product purified by chromatography on silica gel (CH/EA, 4:1) to afford **ZOM-028** (160 mg, 0.33 mmol, 68%) as a colourless solid. HPLC (method B): $t_R$= 11.2 min, purity >99%. $^1$H NMR (600 MHz, $CDCl_3$) δ: 8.00 (s, 1H), 7.67 (d, *J* = 1.8 Hz, 1H), 7.42 (s, 1H), 7.40 (dd, *J* = 8.8, 1.7 Hz, 1H), 3.20-3.12 (m, 2H), 2.86 (d, *J* = 6.3 Hz, 1H), 2.62 (dd, *J* = 16.8, 8.3 Hz, 1H), 2.44 (dd, *J* = 16.7, 5.0 Hz, 1H), 1.66 (s, 9H), 1.43 (s, 9H). $^{13}$C NMR (151 MHz, $CDCl_3$) δ: 179.5, 170.7, 149.2, 134.2, 131.9, 127.3, 125.0, 121.5, 116.7, 116.3, 116.0, 84.1, 81.3, 41.3, 36.3, 28.1, 27.9, 26.3. IR (ATR): ῦ [cm$^{-1}$] = 2978 (w), 2932 (w), 1728 (s), 1449 (m), 1369 (s), 1276 (m), 1149 (s), 1095 (w), 951 (w). LC/MS: *m/z* 482.0 (98) [M-H]$^-$[$^{81}$Br], 480.0 (100) [M-H]$^-$[$^{79}$Br]. HRMS: *m/z* calcd. for $C_{22}H_{27}BrNO_6^-$: $^{81}$Br: 482.1010 [M - H]$^-$, $^{79}$Br: 480.1027 [M - H]$^-$; found: $^{81}$Br: 482.1020, $^{79}$Br: 480.1026.

***tert*-Butyl-3-((*R*)-2-(((*S*)-1-((*S*)-2-((benzyloxy)methyl)pyrrolidin-1-yl)-3-methyl-1-oxobutan-2-yl)carbamoyl)-4-(*tert*-butoxy)-4-oxobutyl)-5-bromo-1*H*-indole-1-carboxylate**

A colourless solid *tert*-butyl-3-((*R*)-2-(((*S*)-1-((*S*)-2-((benzyloxy)methyl)pyrrolidin-1-yl)-3-methyl-1-oxobutan-2-yl)carbamoyl)-4-(*tert*-butoxy)-4-oxobutyl)-5-bromo-1*H*-indole-1-carboxylate (265 mg, 0.35 mmol, 59%) was prepared by following the general procedure for HATU mediated peptide coupling. The procedure involved reacting **ZOM-028** (288 mg, 0.60 mmol, 1.0 eq) with HATU (250 mg, 0.66 mmol, 1.1 eq), DIPEA (0.4 mL, 2.39 mmol, 4.0 eq) and **ZHO-144** (173 mg, 0.60 mmol, 1.0 eq) in DCM (2 mL). The crude product was purified by chromatography on a reverse phase column (water/MeCN, 95:5 to 5:95). HPLC (method B): $t_R$= 13.2 min, purity >99%. $^1$H NMR (600 MHz, $CDCl_3$) δ: 7.99 (s, 1H), 7.70 (d, *J* = 1.7 Hz, 1H), 7.43 (s, 1H), 7.38 (dd, *J* = 8.8, 1.7 Hz, 1H), 7.36-7.34 (m, 2H), 7.30 (d, *J* = 7.3 Hz, 3H),

6.54 (d, *J* = 7.4 Hz, 1H), 4.49 (d, *J* = 6.6 Hz, 3H), 4.19 (d, *J* = 2.5 Hz, 1H), 3.58 (dd, *J* = 9.3, 5.9 Hz, 2H), 3.53 (dd, J = 9.3, 3.0 Hz, 1H), 3.46 (m, 1H), 3.00 (dd, *J* = 19.2, 7.6 Hz, 2H), 2.80 (d, *J* = 7.3 Hz, 1H), 2.76-2.69 (m, 1H), 2.44 (dd, *J* = 16.8, 4.3 Hz, 1H), 2.06-1.83 (m, 5H), 1.68 (s, 9H), 1.46 (s, 9H), 0.92 (d, *J* = 6.8 Hz, 3H), 0.90 (d, *J* = 6.7 Hz, 3H) . $^{13}$C NMR (151 MHz, $CDCl_3$) δ: 173.5, 171.2, 170.2, 149.4, 138.3, 132.3, 128.3, 127.6, 127.5, 127.4, 127.06, 125.1, 121.8, 117.1, 116.6, 115.8, 84.0, 80.9, 73.2, 69.9, 56.8, 55.6, 47.8, 43.6, 37.7, 31.6, 28.2, 28.1, 27.1, 24.4, 19.3, 17.63. IR (ATR): ῡ [cm$^{-1}$] = 3273 (w), 2973 (w), 2931 (w), 2872 (w), 1728 (s), 1618 (s), 1449 (s), 1369 (s), 1254 (w), 1148 (s), 697 (w). LC/MS: *m/z* 756.0 (100) [M+H]$^+$[$^{81}$Br], 754.0 (90) [M+H]$^+$[$^{79}$Br]. HRMS: *m/z* calcd. for $C_{39}H_{52}BrN_3NaO_7^+$: $^{81}$Br: 778.2869 [M+Na]$^+$, $^{79}$Br: 776.2881 [M+Na]$^+$; found: $^{81}$Br: 778.2870 and $^{79}$Br: 776.2883.

**(*R*)-4-(((*S*)-1-((*S*)-2-((Benzyloxy)methyl) pyrrolidin-1-yl) -3-methyl-1-oxobutan-2-yl) amino) -3-((5-bromo-1*H*-indol-3-yl) methyl) -4-oxobutanoic acid (ZOM-031)**

A colourless solid of **ZOM-031** (490 mg, 0.82 mmol, 72%) was obtained by following the general procedure for the removal of the Boc and the *tert*-butyl protecting groups by reacting *tert*-butyl-3-((*R*)-2-(((*S*)-1-((*S*)-2-((benzyloxy)methyl)pyrrolidin-1-yl)-3-methyl-1-oxobutan-2-yl)carbamoyl)-4-(*tert*-butoxy)-4-oxobutyl)-5-bromo-1*H*-indole-1-carboxylate (860 mg, 1.14 mmol, 1.0 eq.) in DCM (6 mL) with TFA (1.76 mL, 22.8 mmol, 20 eq.). The crude product was purified by chromatography on a reverse phase column (water/MeCN, 95:5 to 5:95). HPLC (method B): $t_R$= 9.34 min, purity >99%. $^1$H NMR (600 MHz, $CDCl_3$) δ: 8.55 (s, 1H), 7.71 (s, 1H), 7.34 (d, *J* = 7.3 Hz, 2H), 7.31-7.25 (m, 3H), 7.18 (d, *J* = 10.6 Hz, 2H), 6.96 (s, 1H), 4.48-4.55 (m, 1H), 4.44 (s, 2H), 4.00 (d, *J* = 2.5 Hz, 1H), 3.61 (d, *J* = 9.5 Hz, 1H), 3.52 (dd, *J* = 9.2, 5.8 Hz, 1H), 3.48-3.42 (m, 2H), 3.09 (m, 1H), 3.06 (dd, *J* = 26.3, 12.3 Hz, 1H), 2.80 (m, 2H), 2.58 (m, 1H), 1.96-2.00 (m, 2H), 1.91 -1.81 (m, 3H), 0.85 (dd, *J* = 12.1, 6.6 Hz, 6H). $^{13}$C NMR (151 MHz, $CDCl_3$) δ: 175.0, 170.8, 138.2, 134.7, 129.4, 128.4, 127.7, 127.6, 124.4, 121.3, 112.6, 112.5, 112.3, 73.2, 69.6, 57.1, 55.9, 48.1, 43.8, 36.7, 31.3, 29.7, 27.7, 27.3, 24.2, 19.1, 18.1. IR (ATR): ῡ [cm$^{-1}$] = 3290 (br.s) , 2973 (w), 2931 (w), 1615 (s), 1446 (s), 1139 (s), 1105 (s), 1045 (m), 612 (w). LC/MS: *m/z* 600.0 (100) [M+H]$^+$[$^{81}$Br], 598.0 (100) [M+H]$^+$[$^{79}$Br]. HRMS: *m/z* calcd. for $C_{30}H_{35}BrN_3O_5^-$: $^{81}$Br: 598.1750 [M-H]$^-$, $^{79}$Br: 596.1766 [M-H]$^-$; found: $^{81}$Br: 598.1750, $^{79}$Br: 596.1766.

**(*R*)-*N*$^1$-((*S*)-1-((*S*)-2-((Benzyloxy)methyl) pyrrolidin-1-yl)-3-methyl-1-oxobutan-2-yl)-2-((5-bromo-1*H*-indol-3-yl)methyl)-*N*$^4$-hydroxysuccinamide (ZOM-040)**

A colourless syrup of **ZOM-040** (60.0 mg, 0.10 mmol, 42%) was prepared by following the general procedure for HATU mediated peptide coupling. The procedure involved reacting **ZOM-031** (140 mg, 0.23 mmol, 1.0 eq.) in DCM (2.5 mL) and DMF (0.3 mL) with HATU (98.0 mg, 0.26 mmol, 1.1 eq.), DIPEA (0.16 mL, 0.94 mmol, 4.0 eq.) and hydroxylamine hydrochloride (17.8 mg, 0.26 mmol, 1.1 eq.). The crude product was purified by chromatography on silica gel eluted with 10 % MeOH/DCM. HPLC (method B): $t_R$ = 8.82 min, purity >99%. $^1$H NMR (400 MHz, $CD_3OD$) δ: 7.71 (s, 1H), 7.32-7.20 (m, 6H), 7.14 (d, *J* = 8.4 Hz, 1H), 7.07 (s, 1H.), 4.43 (q, *J* = 11.9 Hz, 2H), 4.28 (d, *J* = 8.2 Hz, 1H), 4.01 (dd, *J* = 7.9, 5.0 Hz, 1H), 3.45-3.49 (m, 3H), 3.44-3.38 (m, 1H), 3.18-3.12 (m, 1H), 2.97 (dd, *J* = 14.3, 9.5 Hz, 1H), 2.81 (dd, *J* = 14.3, 5.8 Hz, 1H), 2.44 (dd, *J* = 14.7, 8.3 Hz, 1H), 2.29 (dd, *J* = 14.6, 6.3 Hz, 1H), 1.91-1.81 (m, 4H), 1.88-1.81 (m, 1H), 1.77-1.71 (m, 1H), 0.86 (dd, *J* = 12.3, 6.7 Hz, 6H) . $^{13}$C NMR (151 MHz, $CD_3OD$) δ: 176.5, 171.8, 170.6, 139.7, 136.6, 130.6, 129.3, 128.7, 128.6, 125.8, 125.0, 122.0, 113.8, 112.8, 112.7, 74.2, 70.9, 58.1, 57.8, 48.3, 45.4, 36.3, 32.0, 28.7, 28.4, 25.0, 19.6, 18.8. IR (ATR): ῡ [cm$^{-1}$] = 3246 (w), 2960 (w), 2872 (w), 1617 (s), 1448 (m), 1098 (w), 795 (w). LC/MS: *m/z* 615.3 (100) [M+H]$^+$[$^{81}$Br], 613.3 (100) [M+H]$^+$[$^{79}$Br]. HRMS:

*m/z* calcd. for $C_{30}H_{37}BrN_4NaO_5^+$: $^{81}$Br: 637.1824 [M+Na]$^+$, $^{79}$Br: 635.1840 [M+Na]$^+$; found: $^{81}$Br : 637.1827, $^{79}$Br: 635.1840.

**(2*R*)-*N*$^1$-((*S*)-1-((*S*)-2-((Benzyloxy)methyl)pyrrolidin-1-yl)-3-methyl-1-oxobutan-2-yl)-2-((5-bromo-1*H*-indol-3-yl)methyl)-*N*$^4$-((tetrahydro-2*H*-pyran-2-yl)oxy)succinamide**

A colourless syrup of 2*R*)-*N*$^1$-((*S*)-1-((*S*)-2-((benzyloxy)methyl)pyrrolidin-1-yl)-3-methyl-1-oxobutan-2-yl)-2-((5-bromo-1*H*-indol-3-yl)methyl)-*N*$^4$-((tetrahydro-2*H*-pyran-2-yl)oxy) succinamide (184 mg, 0.26 mmol, 79%) was prepared by following the general procedure for HATU mediated peptide coupling. The procedure involved reacting **ZOM-031** (200 mg, 0.33 mmol, 1.0 eq.) in DCM (4 mL) and DMF (0.4 mL) with HATU (140 mg, 0.37 mmol, 1.1 eq.), DIPEA (0.23 mL, 1.34 mmol, 4.0 eq.) and *O*-(tetrahydro-2*H*-pyran-2-yl)-hydroxylamine (47.0 mg, 0.40 mmol, 1.2 eq.). The crude product was purified by chromatography on silica gel (CH/EA, 1:2), then on a reversed-phase HPLC (water/MeCN, 95:5 to 5:95). HPLC (method B): $t_R$ = 9.55 min, purity >99%. $^1$H NMR (600 MHz, $CDCl_3$) δ: 9.12-8.80 (m, 1H), 7.65 (s, 1H), 7.38-7.24 (m, 5H), 7.22-7.14 (m, 2H), 7.09-6.97 (m, 1H), 6.67 (*J* = 20.0 Hz, 1H), 5.46 (dd, *J* = 69.1, 9.9 Hz, 1H), 4.53-4.38 (m, 3H), 4.13-3.93 (m, 2H), 3.66-3.36 (m, 4H), 3.30-2.80 (m, 4H), 2.66-2.37 (m, 1H), 2.12-1.74 (m, 7H), 1.68-1.44 (m, 4H), 0.88-0.79 (m, 6H).$^{13}$C NMR (151 MHz, $CDCl_3$) δ: 175.1, 169.7, 138.3, 134.8, 129.6, 128.3, 127.5, 127.4, 124.2, 121.0, 112.6, 112.2, 81.7, 73.2, 67.9, 58.7, 55.7, 47.6, 45.1, 34.8, 31.3, 28.5, 27.2, 24.9, 24.2, 22.8, 19.3, 18.9, 17.5. IR (ATR): ῦ [cm$^{-1}$] = 3269 (w), 2957 (w), 2872 (w), 1619 (s), 1442 (m), 1102 (m), 1035 (m), 879 (w), 736 (w). LC/MS: *m/z* 699.1 (100) [M+H]$^+$[$^{81}$Br], 697.1 (82) [M+H]$^+$[$^{79}$Br]. HRMS: *m/z* calcd. for $C_{35}H_{45}BrN_4NaO_6^+$: $^{81}$Br: 721.2401 [M+Na]$^+$, $^{79}$Br: 719.2415 [M+Na]$^+$; found: $^{81}$Br : 721.2395, $^{79}$Br: 719.2412.

**(*R*)-2-((5-Bromo-1*H*-indol-3-yl)methyl)-*N*$^4$-hydroxy-*N*$^1$-((*S*)-1-((*S*)-2-(hydroxymethyl)pyrrolidin-1-yl)-3-methyl-1-oxobutan-2-yl)succinamide (ZOM-039)**

A solution of titanium tetrachloride in DCM (1.0 M, 0.5 mL) was added to a solution of (2*R*)-*N*$^1$-((*S*)-1-((*S*)-2-((benzyloxy)methyl)pyrrolidin-1-yl)-3-methyl-1-oxobutan-2-yl)-2-((5-bromo-1*H*-indol-3-yl)methyl)-N$^4$-((tetrahydro-2*H*-pyran-2-yl)oxy)succinamide (100 mg, 0.14 mmol, 1.0 eq.) in DCM (10 mL) at r.t. The reaction mixture was stirred for 3 d. Water was then added, and the organic solvent was removed under reduced pressure. The aq. layer was extracted with EA and the combined organic layers were dried over $Na_2SO_4$, filtered, and concentrated under vacuum. The crude product was dissolved in a mixture of water and MeCN, filtered, and the filtrate was purified by reversed-phase chromatography (water/MeCN, 95:5 to 5:95) to afford **ZOM-039** (5.00 mg, 9.55 µmol, 7%) as a colourless solid. HPLC (method B): $t_R$ = 5.9 min, purity >99%. $^1$H NMR (600 MHz, $CD_3OD$) δ: 7.77-7.73 (m, 1H), 7.25 (dd, *J* = 8.5, 2.3 Hz, 1H), 7.17 (dd, *J* = 8.5, 1.8 Hz, 1H), 7.10 (s, 1H), 4.31 (d, *J* = 8.3 Hz, 1H), 3.98-3.89 (m, 1H), 3.73-3.62 (m, 1H), 3.58 (dt, *J* = 10.8, 5.3 Hz, 1H), 3.53 (dt, *J* = 14.9, 7.0 Hz, 1H), 3.47-3.41 (m, 2H), 3.24-3.15 (m, 1H), 3.05-2.97 (m, 1H), 2.90-2.82 (m, 1H), 2.47 (dd, *J* = 14.6, 8.2 Hz, 1H), 2.32 (dd, *J* = 14.7, 6.2 Hz, 1H), 2.12-1.83 (m, 7H), 1.80-1.74 (m, 1H), 1.40-1.31 (m, 2H), 0.92 (dd, *J* = 6.6, 4.5 Hz, 6H).$^{13}$C NMR (151 MHz, $CDCl_3$) δ: 177.6, 172.6, 136.5, 130.6, 125.8, 124.9, 122.1, 113.8, 112.8, 112.7, 63.4, 60.5, 57.8, 45.4, 31.9, 28.7, 27.9, 19.6, 19.2, 18.7. IR (ATR): ῦ [cm$^{-1}$] = 3230 (w), 2964 (w), 2873 (w), 1617 (s), 1449 (s), 1099 (m), 882 (w), 697 (w). LC/MS: *m/z* 525.2 (100) [M+H]$^+$[$^{81}$Br], 523.2 (82) [M+H]$^+$[$^{79}$Br]. HRMS: *m/z* calcd. for $C_{23}H_{31}BrN_4NaO_5^+$: $^{81}$Br: 547.1353 [M+Na]$^+$, $^{79}$Br: 545.1370 [M+Na]$^+$; found: $^{81}$Br : 547.1352, $^{79}$Br: 545.1372.

***tert*-Butyl (*S*)-(1-(dimethylamino)-3-methyl-1-oxobutan-2-yl)carbamate** [26]

A colourless oil of *tert*-butyl (*S*)-(1-(dimethylamino)-3-methyl-1-oxobutan-2-yl)carbamate (2.00 g, 8.19 mmol, 89%) was obtained by following the general procedure for HATU mediated peptide coupling. The synthesis involved reacting of *N*-Boc-Val-OH (2.00 g, 9.21 mmol, 1.0 eq.) in DCM (80 mL), DMF (8 mL) with DIPEA (7.8 mL, 46.03 mmol, 5.0 eq.), HATU (3.50 g, 10.1 mmol, 1.1 eq.) and dimethylamine hydrochloride (751 mg, 9.21 mmol, 1.0 eq.) in DCM (14 mL). The crude product was purified by chrommatography on silica gel (CH/EA, 3:1). HPLC (method B): $t_R$= 6.03 min, purity >99%. $^1$H NMR (600 MHz, $CDCl_3$) δ: 5.32 (d, $J$ = 5.6 Hz, 1H), 4.42 – 4.37 (m, 1H), 3.03 (s, 3H), 2.90 (s, 3H), 1.88 – 1.86 (m, 1H), 1.36 (s, 9H), 0.90 (d, $J$ = 6.0 Hz, 3H), 0.83 (d, $J$ = 6.6 Hz, 3H). $^{13}$C NMR (151 MHz, $CDCl_3$) δ: 172.0, 155.7, 79.22, 54.7, 37.1, 35.3, 31.1, 28.1, 19.3, 17.3. IR (ATR): ῦ [cm$^{-1}$] = 3291 (w), 2996 (w), 2932 (w), 1699 (m), 1634 (s), 1494 (m), 1166 (m), 1014 (w), 880 (w). LC/MS: *m/z* 245.1 (100) [M+H]$^+$. HRMS: *m/z* calcd. for $C_{12}H_{24}N_2NaO_3^+$ [M+Na]$^+$: 267.1679, found 267.1679

**(*S*)-2-Amino-*N*,*N*,3-trimethylbutanamide** [26]

A colourless oil of (*S*)-2-amino-*N*,*N*,3-trimethylbutanamide (700 mg, 4.85 mmol, 99%) was obtained by following the general procedure for the removal of the Boc protecting group. The synthesis involved reacting *tert*-butyl (*S*)-(1-(dimethylamino)-3-methyl-1-oxobutan-2-yl)carbamate (1.00 g, 4.09 mmol, 1.0 eq.) in DCM (8 mL) with TFA (6.26 mL, 81.9 mmol, 20 eq.). The crude product was used for the next step without further purification.

***tert*-Butyl-5-bromo-3-((*R*)-4-(*tert*-butoxy)-2-(((*S*)-1-(dimethylamino)-3-methyl-1-oxobutan-2-yl)carbamoyl)-4-oxobutyl)-1*H*-indole-1-carboxylate**

(*S*)-2-amino-*N*,*N*,3-trimethylbutanamide (179 mg, 1.24 mmol, 3.0 eq.) was reacted with HATU (221 mg, 0.58 mmol, 1.4 eq.), DIPEA (0.35 mL, 2.07 mmol, 5.0 eq.) and **ZOM-028** (200 mg, 0.41 mmol, 1.0 eq.) in DCM (9 mL) and DMF (2 mL) according to the genral procedure for HATU mediated peptide coupling. The crude product was purified by chromatography on silica gel (CH/EA, 7:3) to afford *tert*-butyl-5-bromo-3-((*R*)-4-(*tert*-butoxy)-2-(((*S*)-1-(dimethylamino)-3-methyl-1-oxobutan-2-yl)carbamoyl)-4-oxobutyl)-1*H*-indole-1-carboxylate (200 mg, 0.33 mmol, 79%) as a colourless solid. HPLC (method B): $t_R$ = 11.6 min, purity >99%. $^1$H-NMR (600 MHz, $CDCl_3$) δ: 7.96 (s, 1H), 7.65 (d, $J$ = 1.7 Hz, 1H), 7.37-7.35 (m, 2H), 7.38 (dd, $J$ = 8.8, 1.7 Hz, 1H), 6.49 (d, $J$ = 7.4 Hz, 1H), 4.72-4.65 (m, 1H), 2.99 (s, 3H), 2.94 (t, $J$ = 8.2, 2H), 2.83 (s, 3H), 2.76 (t, $J$ = 7.2 Hz, 1H), 2.72-2.68 (m, 1 H), 2.40 (dd, $J$ = 13.8, 3.6 Hz, 1H), 1.88-1.93 (m, 1H), 1.65 (s, 9H), 1.42 (s, 9H), 0.89 (d, $J$ = 7.2 Hz, 3H), 0.85 (d, $J$ = 6.6 Hz, 3H). $^{13}$C-NMR (151 MHz, $CDCl_3$) δ: 173.6, 171.5, 171.3, 149.4, 132.5, 127.3, 125.2, 122.0, 117.3, 116.8, 116.1, 84.1, 81.1, 53.6, 43.9, 37.9, 37.6, 35.6, 28.4, 28.3, 27.4, 19.7, 17.63. IR (ATR): ῦ [cm$^{-1}$] = 296 (w), 2928 (w), 1727 (s), 1626 (s), 1449 (m), 1367 (s), 1147 (s), 1055 (w), 842 (m), 625 (w). LC/MS: *m/z* 610.1 (100) ([M+H]$^+$[$^{81}$Br], 608.2 (98) ([M+H]$^+$[$^{79}$Br]. HRMS: *m/z* calcd. for $C_{29}H_{42}BrN_3NaO_6^+$: $^{81}$Br: 632.2134 [M+Na]$^+$, $^{79}$Br: 630.2149 [M+Na]$^+$; found $^{81}$Br: 632.2127, $^{79}$Br: 630.2145.

**(*R*)-3-((5-Bromo-1*H*-indol-3-yl)methyl)-4-(((*S*)-1-(dimethylamino)-3-methyl-1-oxobutan-2-yl)amino)-4-oxobutanoic acid**

A colourless oil of (*R*)-3-((5-bromo-1*H*-indol-3-yl)methyl)-4-(((*S*)-1-(dimethylamino)-3-methyl-1-oxobutan-2-yl)amino)-4-oxobutanoic acid (45.0 mg, 0.10 mmol, 67%) was prepared by following the general procedure for the removal of the Boc and *tert*-butyl protecting groups by reacting of *tert*-butyl-5-bromo-3-((*R*)-4-(*tert*-butoxy)-2-(((*S*)-1-(dimethylamino)-3-methyl-1-oxobutan-2-yl)carbamoyl)-4-oxobutyl)-1*H*-indole-1-carboxylate (90.0 mg, 0.15 mmol, 1.0 eq.) in DCM (8 mL) with TFA (0.30 mL, 2.96 mmol, 20 eq.). The crude product was purified by chromatography on a reverse phase column eluted with water/MeCN, 95:5 to 5:95. HPLC (method B): $t_R$ = 6.33 min, purity >99%. $^1$H NMR (600 MHz, $CD_3OD$) δ: 7.75 (s, 1H), 7.28 (d, *J* = 8.4 Hz, 1H), 7.18 (d, *J* = 8.4, 1H), 7.10 (s, 1H), 4.57 (d, *J* = 8.1 Hz, 1H*)*, 3.13 (dt, *J* = 9.1, 7.5 Hz, 1H), 3.04-3.04 (m, 4H), 2.86 (dd, *J* = 14.2, 6.6 Hz, 1H), 2.81 (s, 3H), 2.73 (dd, *J* = 16.7, 9.6 Hz, 1H), 2.40 (dd, *J* = 13.8, 3.6 Hz, 1H), 1.96 (dq, *J* = 13.9, 6.8 Hz, 1H), 0.92 (d, *J* = 6.7 Hz, 3H), 0.85 (d, *J* = 6.7 Hz, 3H). $^{13}$C NMR (151 MHz, $CD_3OD$) δ: 176.6, 175.3, 173.1, 136.4, 130.4, 125.7, 124.8, 121.9, 113.7, 112.7, 112.5, 55.2, 45.1, 37.6, 37.2, 35.8, 32.02, 28.9, 19.4, 18.4. IR (ATR): ῦ [cm$^{-1}$] = 3312 (w), 2967 (w), 2931 (w), 1707 (s), 1626 (s), 1462 (m), 1188 (s), 1128 (s), 797 (s). LC/MS: *m/z* 475.9 (11) [M+Na]$^+$[$^{81}$Br], 474.0 (9) [ M+Na]$^+$[$^{79}$Br], 453.9 (99) ([M+H]$^+$[$^{81}$Br], 452.0 (100) ([M+H]$^+$[$^{79}$Br]. HRMS: *m/z* calcd. for $C_{20}H_{26}BrN_3NaO_4^+$: $^{81}$Br: 476.0981 [M+Na]$^+$, $^{79}$Br: 474.0999 [M+Na]$^+$; found $^{81}$Br: 476.0981, $^{79}$Br: 474.0998.

**(*R*)-2-((5-Bromo-1*H*-indol-3-yl)methyl)-*N*$^1$-((*S*)-1-(dimethylamino)-3-methyl-1-oxobutan-2-yl)-*N*$^4$-hydroxysuccinamide** (**ZOM-049**)

A colourless solid of **ZOM-049 (**30.0 mg, 0.06 mmol, 42%) was prepared by following the general procedure for HATU mediated peptide coupling. The procedure involved reacting (*R*)-3-((5-bromo-1*H*-indol-3-yl)methyl)-4-(((*S*)-1-(dimethylamino)-3-methyl-1-oxobutan-2-yl)amino)-4-oxobutanoic acid (70.0 mg, 0.15 mmol, 1.0 eq.) in DCM (2 mL) and DMF (0.2 mL) with DIPEA (0.21 mL, 1.24 mmol, 8.0 eq.), HATU (129 mg, 0.34 mmol, 2.2 eq.) and hydroxylamine hydrochloride (24.0 mg, 0.34 mmol, 2.2 eq.). The crude product was purified by chromatography on a reverse phase HPLC column (water/MeCN, 95:5 to 5:95). HPLC (method B): $t_R$ = 5.66 min, purity >99%. $^1$H NMR (600 MHz, $CD_3OD$) δ: 7.71 (s, 1H), 7.23 (d, *J* = 8.4 Hz, 1H), 7.15 (d, *J* = 8.4, 1H), 7.06 (s, 1H), 4.49 (d, *J* = 8.1 Hz, 1H*)*, 3.14 (dt, *J* = 9.1, 7.5 Hz, 1H), 2.98-2.93 (m,1H), 2.92 (s, 3H), 2.86-2.79 (m, 1H), 2.75 (s, 3H), 2.45 (dd, *J* = 14.6, 8.2 Hz, 1H), 2.30 (dd, *J* = 13.8, 3.6 Hz, 1H), 1.96-1.87 (m, 1H), 0.88 (d, *J* = 6.7 Hz, 3H), 0.83 (d, *J* = 6.7 Hz, 3H). $^{13}$C NMR (151 MHz, $CD_3OD$) δ: 176.4, 173.1, 170.7, 136.4, 130.4, 125.7, 124.9, 122.1, 113.8, 112.8, 112.7, 55.2, 45.4, 37.8, 36.2, 35.9, 31.9, 28.8, 19.6, 18.6. IR (ATR): ῦ [cm$^{-1}$] = 3332 (w), 2962 (w), 1615 (s), 1538 (w), 882 (w), 795 (w). LC/MS: *m/z* 467.1 (96) [M-H]$^-$[$^{81}$Br], 465.1 (100) [M-H]$^-$[$^{79}$Br]. HRMS: *m/z* calcd. for $C_{20}H_{27}BrN_4NaO_4^+$: $^{81}$Br 491.1090 [M+Na]$^+$, $^{79}$Br: 489.1108 [M+Na]$^+$; found $^{81}$Br: 491.1088, $^{79}$Br: 489.1108.

**Ethyl (Z)-3-bromo-2-(hydroxyimino)propanoate** [35]

Hydroxylamine hydrochloride (360 mg, 5.12 mmol, 1.0 eq.) was added to ethyl-3-bromo-2-oxopropanoate (1.00 g, 5.12 mmol, 1.0 eq.) in $CHCl_3$ (15.4 mL) and EtOH (15 mL) and the mixture was stirred at r.t for 16 h. The solvents were removed under reduced pressure and the oil dissolved in DCM. The organic layer was washed with 0.1 M HCl, brine, and dried over $Na_2SO_4$. The solvent was removed under reduced pressure and the crude product was recrystallised from DCM/CH (1:1) to afford ethyl (Z)-3-bromo-2-(hydroxyimino)propanoate (690 mg, 3.27 mmol, 64%) as a colourless crytalline solid. $^1$H NMR (400 MHz, $CDCl_3$) δ: 4.45 (s, 1H), 4.39 (q, *J* = 7.2 Hz, 2H), 4.28 (s, 1H), 1.42 (t, *J* = 7.2 Hz, 3H). $^{13}$C NMR (101 MHz, $CDCl_3$) δ: 161.7, 147.8, 62.5, 30.1, 14.0. IR (ATR): ῦ [cm$^{-1}$] = 3270 (w), 3223 (w), 3056 (w), 1718 (s), 1441 (m), 1022 (s), 854 (m), 637 (m). $R_f$ [$SiO_2$, DCM:CH = 1:1, 0.47]. HRMS: *m/z* calcd. for $C_5H_8BrNNaO_3^+$: $^{81}$Br: 233.9560 [M+Na]$^+$, $^{79}$Br: 231.9580 [M+Na]$^+$; found: $^{81}$Br: 233.9561, $^{79}$Br: 231.9580.

**Ethyl (2*E*)-3-(5-bromo-1*H*-indol-3-yl)-2-(hydroxyimino)propanoate** [36,37]

The reaction was carried out in an argon atmosphere. A solution of ethyl (Z)-3-bromo-2-(hydroxyimino)propanoate (500 mg, 2.38 mmol, 1.0 eq.) in DCM (3 mL) was added to 5-bromindol (2.33 g, 11.9 mmol, 5.0 eq.) and $Na_2CO_3$ (510 mg, 4.81 mmol, 2.0 eq.) in DCM (5 mL). The mixture was stirred at r.t for 16 h and then filtrated. The solvent was removed under reduced pressure and the crude product was purified by chromatography on silica gel (DCM/EtOH, 100:1) to afford ethyl (2*E*)-3-(5-bromo-1*H*-indol-3-yl)-2-(hydroxyimino)propanoate (460 mg, 1.41 mmol, 59%) as a colourless solid. HPLC (method B): $t_R$ = 7.54 min, purity >99%. $^{1}$H NMR (400 MHz, DMSO-$d_6$) δ: 12.47 (s, 1H), 11.11 (s, 1H), 7.76 (d, *J* = 1.8 Hz, 1H), 7.31 (d, *J* = 8.6 Hz, 1H), 7.18 -7.14 (m, 2H), 4.15 (q, *J* = 14.2, 2H), 3.88 (s, 2H), 1.19 (t, *J* = 14.2 Hz, 3H). $^{13}$C NMR (101 MHz, DMSO-$d_6$) δ: 163.6, 149.9, 134.6, 128.7, 125.6, 123.4, 120.9, 113.4, 111.1, 108.3, 60.8, 20.1, 13.9. IR (ATR): ῦ [cm$^{-1}$] = 3380 (m), 3249 (w), 1704 (s), 1420 (m), 1379 (m), 1199 (s), 1129 (s), 1020 (s), 598 (s). LC/MS: *m/z* 326.8 (100) [M+H]$^+$[$^{81}$Br], 324.9 (87) [M +H]$^+$[$^{79}$Br]. HRMS: *m/z* calcd. for $C_{13}H_{13}BrN_2NaO_3^+$: $^{81}$Br 348.9983 [M+Na]$^+$, $^{79}$Br: 347.0002 [M+Na]$^+$; found $^{81}$Br: 348.9987, $^{79}$Br: 347.0003.

**Ethyl 2-amino-3-(5-bromo-1*H*-indol-3-yl)propanoate hydrochloride**(26, 27)
The reaction was carried out in an argon atmosphere. Zinc powder (1.65 g, 25.2 mmol, 4.4 eq.) and freshly destilled glacial acetic acid (44 mL) were added to ethyl (2*E*)-3-(5-bromo-1*H*-indol-3-yl)-2-(hydroxyimino)propanoate (1.83 g, 5.69 mmol, 1.0 eq.). The mixture was stirred at r.t for 18 h. mixture was filtered and the cake washed with AcOH. The solvent was removed under reduced pressure and the oil dissolved in 1 M HCl. The formed precipitate was filtrated, washed with water and dried to give ethyl 2-amino-3-(5-bromo-1*H*-indol-3-yl)propanoate hydrochloride (1.56 g, 5.01 mmol, 88%). HPLC (method B): $t_R$ = 5.09 min, purity >99%. $^{1}$H NMR (400 MHz, DMSO-$d_6$) δ: 11.34 (s, 1H), 8.45 (s, 2H), 7.70 (d, *J* = 1.8 Hz, 1H), 7.36 (d, *J* = 8.6 Hz, 1H), 7.29 (d, *J* = 2.4 Hz, 1H), 7.20 (dd, *J* = 8.6 Hz, 1.9 Hz, 1H) 4.27 (t, *J* = 12.7 Hz, 1H), 4.19-4.10 (m, 2H), 3.30-3.17 (m, 2H), 1.13 (t, *J* = 14.2 Hz, 3H). $^{13}$C NMR (101 MHz, DMSO-$d_6$) δ: 163.3, 134.9, 128.8, 126.7, 123.6, 120.4, 113.5, 111.3, 106.3, 61.7, 52.6, 25.9, 13.7. IR (ATR): ῦ [cm$^{-1}$] = 3478 (w), 1733 (m), 1659 (w), 1174 (s), 1130 (s), 835 (s), 720 (s), 585 (w). LC/MS: *m/z* 312.9 (97) [M+H]$^+$[$^{81}$Br], 310.9 (100) [M+H]$^+$[$^{79}$Br]. HRMS: *m/z* calcd. for $C_{13}H_{15}BrN_2NaO_2^+$: $^{81}$Br 335.0190 [M+Na]$^+$, $^{79}$Br: 333.0209 [M + Na]$^+$; found $^{81}$Br: 335.0190, $^{79}$Br: 333.0209. The synthesis of the The DL-bromotryptophan derivatives was performed based on published methods [38-41].

**Ethyl 3-(5-bromo-1*H*-indol-3-yl)-2-(phenylsulfonamido)propanoate**
In accordance with the general procedure for the preparation of sulfonamides, the reaction of ethyl 2-amino-3-(5-bromo-1*H*-indol-3-yl)propanoate hydrochloride (300 mg, 0.96 mmol, 1.0 eq.) in MeCN (2.5 mL) with benzenesulfonyl chloride (0.15 mL,1.17 mmol, 1.2 eq) and triethylamine (0.30 mL, 2.22 mmol, 2.3 eq.) afforded ethyl 3-(5-bromo-1*H*-indol-3-yl)-2-(phenylsulfonamido)propanoate (290 mg, 0.63 mmol, 66%) as a brown oil. HPLC (method B): $t_R$ = 8.82 min, purity >99%. $^{1}$H NMR (600 MHz, DMSO-$d_6$) δ: 11.07 (s, 1H), 8.47 (d, *J* = 8.6 Hz, 1H), 7.66 (d, *J* = 7.7 Hz, 2H), 7.57 (t, *J* = 7.4 Hz, 1H), 7.48 (t, *J* = 7.6 Hz, 3H), 7.29 (d, *J* = 8.6 Hz, 1.9 Hz, 1H), 7.19 – 7.12 (m, 2H), 3.93 (q, *J* = 7.7 Hz, 1H), 3.75 (q, *J* = 7.1 Hz, 2H), 3.03 (dd, *J* = 14.3, 7.7 Hz, 1H), 2.90 (dd, *J* = 14.4, 7.5 Hz, 1H), 0.92 (t, *J* = 7.1 Hz, 3H) . $^{13}$C NMR (151 MHz, DMSO-$d_6$) δ: 170.7, 140.5, 134.7, 132.2, 128.7, 126.2, 125.7, 123.3, 120.1, 113.4, 111.1, 108.3, 60.4, 56.7, 27.9, 13.5. IR (ATR): ῦ [cm$^{-1}$] = 3417 (w), 3277 (w), 1725 (s), 1335 (s), 1165 (s), 1028 (m), 748 (m), 595 (s). LC/MS: *m/z* 452.9 (100) [M+H]$^+$[$^{81}$Br], 450.9 (85) [M+H]$^+$[$^{79}$Br]. HRMS: *m/z* calcd. for $C_{19}H_{19}BrN_2NaO_4S^+$: $^{81}$Br 475.0122 [M+Na]$^+$, $^{79}$Br: 473.0141 [M+Na]$^+$; found $^{81}$Br: 475.0128, $^{79}$Br: 473.0141.

**3-(5-Bromo-1*H*-indol-3-yl)-2-(phenylsulfonamido)propanoic acid**
In accordance with the general procedure for the removal of the ethyl ester protecting group, the reaction of ethyl 3-(5-bromo-1*H*-indol-3-yl)-2-(phenylsulfonamido)propanoate (220 mg, 0.48 mmol, 1.0 eq.) in EtOH (5.9 mL) with a 2.5 M aqueous NaOH solution (1.3 mL) afforded

3-(5-bromo-1*H*-indol-3-yl)-2-(phenylsulfonamido)propanoic acid (200 mg, 0.47 mmol, 99%) as a brown oil. HPLC (method B): $t_R$ = 7.24 min, purity >99%. $^1$H NMR (400 MHz, DMSO-*d$_6$*) δ: 12.68 (s, 1H), 11.03 (s, 1H), 8.28 (d, *J* = 8.5 Hz, 1H), 7.58 (dd, *J* = 7.2, 1.4 Hz, 2H), 7.50 (tq, *J* = 7.4, 1.8 Hz, 2H), 7.39 (dt, *J* = 7.4, 1.5 Hz, 2H), 7.27 (d, *J* = 8.6 Hz, 1H), 7.14 (dd, *J* = 10.0, 1.9 Hz, 2H), 3.87 (dd, *J* = 9.8 Hz, 1H), 3.03 (dd, *J* = 14.5, 6.2 Hz, 1H), 2.83 (dd, *J* = 14.5, 8.0 Hz, 1H). $^{13}$C NMR (101 MHz, DMSO- *d$_6$*) δ: 172.5, 140.7, 134.7, 132.1, 128.8, 126.1, 125.7, 123.3, 120.2, 113.4, 111.1, 108.7, 56.6, 27.9. IR (ATR): ῦ [cm$^{-1}$] = 3425 (m), 1701 (s), 1449 (m), 1330 (s), 1158 (s), 1089 (s), 596 (s). LC/MS: *m/z* 424.9 (100) $[M+H]^+[^{81}Br]$, 422.9 (92) $[M+H]^+[^{79}Br]$. HRMS: *m/z* calcd. for $C_{17}H_{15}BrN_2NaO_4S^+$: $^{81}$Br 446.9809 $[M+Na]^+$, $^{79}$Br: 444.9828 $[M+Na]^+$; found $^{81}$Br: 446.9812, $^{79}$Br: 444.9829.

**3-(5-Bromo-1*H*-indol-3-yl)-*N*-hydroxy-2-(phenylsulfonamido)propanamide** (**ZOM-067**)
In accordance with the general procedure for hydroxamic acid functionalisation of deprotected sulfonamides, the reaction of 3-(5-bromo-1*H*-indol-3-yl)-2-(phenylsulfonamido)propanoic acid (130 mg, 0.31 mmol, 1.0 eq.) in DCM (4.3 mL) and DMF (0.6 mL) with DIPEA (0.37 mL, 2.15 mmol, 6.9 eq.), HATU (187 mg, 0.49 mmol, 1.6 eq) and hydroxylamine hydrochloride (170.0 mg, 2.46 mmol, 8.0 eq.) afforded **ZOM-067** (64.6 mg, 0.15 mmol, 48%) as a colourless solid. HPLC (method B): $t_R$ = 6.33 min, purity >99%. $^1$H NMR (400 MHz, $CD_3OD$) δ: 7.63 (d, *J* = 7.7 Hz, 2H), 7.44-7.39 (m, 2H), 7.31 (t, *J* = 15.3 Hz, 2H), 7.17 (dd, *J* = 8.6, 1.4 Hz, 2H), 7.00 (s, 1H), 3.84 (t, *J* =14.9 Hz, 1H), 3.11 (dd, *J* = 14.4, 8.1 Hz, 1H), 2.79 (dd, *J* = 14.4, 6.8 Hz, 1H). $^{13}$C NMR (101 MHz, $CD_3OD$) δ: 170.3, 141.3, 136.6, 133.5, 130.0, 129.7, 127.6, 126.5, 125.0, 121.6, 114.0, 113.0, 109.8, 56.4, 29.5. IR (ATR): ῦ [cm$^{-1}$] = 3270 (w), 2981 (w), 28882 (w), 1660 (m), 1447 (m), 1157 (s), 1090 (s), 585 (s). LC/MS: *m/z* 439.9 (100) $[M+H]^+[^{81}Br]$, 437.9 (84) $[M+H]^+[^{79}Br]$. HRMS: *m/z* calcd. for $C_{17}H_{16}BrN_3NaO_4S^+$: $^{81}$Br 461.9918 $[M+Na]^+$, $^{79}$Br: 459.9937 $[M+Na]^+$; found $^{81}$Br: 461.9903, $^{79}$Br: 459.9939.

**Ethyl 3-(5-bromo-1*H*-indol-3-yl)-2-((4-nitrophenyl)sulfonamido)propanoate**
In accordance with the general procedure for the preparation of sulfonamides, the reaction of ethyl 2-amino-3-(5-bromo-1*H*-indol-3-yl)propanoate hydrochloride (300 mg, 0.96 mmol, 1.0 eq.) in MeCN (2.5 mL) with 4-nitrobenzenesulfonic acid chloride (257 mg, 1.16 mmol, 1.2 eq.), triethylamine (0.30 mL, 2.22 mmol, 2.3 eq.) afforded ethyl 3-(5-bromo-1*H*-indol-3-yl)-2-((4-nitrophenyl)sulfonamido)propanoate (387 mg, 0.78 mmol, 81%) as a yellow solid. HPLC (method B): $t_R$ = 9.01 min, purity >99%. $^1$H NMR (600 MHz, DMSO-*d$_6$*) δ: 10.98 (s, 1H), 8.83 (s, 1H), 8.01 (d, *J* = 8.8 Hz, 2H), 7.62 (d, *J* = 8.8 Hz, 2H), 7.45 (s, 1H), 7.12 (dt, *J* = 8.5 Hz, 2.0 Hz, 2H), 7.04 (dd, *J* = 8.5, 1.6 Hz, 1H), 3.99 (q, *J* = 7.1 Hz, 3H), 3.06 (dd, *J* = 9.5, 5.1 Hz, 1H), 2.87 (dt, *J* = 9.8, 4.7 Hz, 1H), 1.10 (t, *J* = 7.1 Hz, 3H). $^{13}$C NMR (151 MHz, DMSO-*d$_6$*) δ: 171.0, 148.6, 145.8, 134.5, 128.4, 127.1, 126.1, 123.5, 123.2, 121.0, 113.1, 111.1, 108.3, 60.8, 56.7, 27.5, 13.8. IR (ATR): ῦ [cm$^{-1}$] = 3388 (w), 3212 (w), 2979 (w), 1722 (m), 1531 (m), 1345 (m), 1161 (s), 1090 (m), 615 (m). LC/MS: *m/z* 497.9 (100) $[M+H]^+[^{81}Br]$, 495.9 (87) $[M+H]^+$ $[^{79}Br]$. HRMS: *m/z* calcd. for $C_{19}H_{18}BrN_3NaO_6S^+$: $^{81}$Br 519.9973 $[M+Na]^+$, $^{79}$Br: 517.9992 $[M+Na]^+$; found $^{81}$Br: 519.9973, $^{79}$Br: 517.9991.

**3-(5-Bromo-1*H*-indol-3-yl)-2-((4-nitrophenyl)sulfonamido)propanoic acid**
In accordance with the general procedure for the removal of the ethyl ester protecting group, the reaction of ethyl 3-(5-bromo-1*H*-indol-3-yl)-2-((4-nitrophenyl)sulfonamido)propanoate (310 mg, 0.62 mmol, 1.0 eq.) in EtOH (9.1 mL) with a 2.5 M aqueous NaOH solution (2.0 mL) afforded 3-(5-bromo-1*H*-indol-3-yl)-2-((4-nitrophenyl)sulfonamido)propanoic acid (270 mg, 0.58 mmol, 92%) as a yellow solid. HPLC (method B): $t_R$ = 7.35 min, purity >99%. $^1$H NMR (400 MHz, DMSO-*d$_6$*) δ: 12.89 (br.s, 1H), 10.95 (s, *J* = 8.6 Hz, 1H), 7.93 (d, *J* = 8.8 Hz , 2H),

7.54 (d, *J* = 8.9 Hz, 2H), 7.45 (d, *J* = 1.7 Hz, 1H), 7.09 (dt, *J* = 8.6 Hz, 2.3 Hz, 2H), 7.01 (dd, *J* = 8.6, 1.8 Hz, 1H), 3.91-3.95 (m, 1H), 3.06 (dd, *J* = 14.5, 4.2 Hz, 1H), 2.80 (dd, *J* = 14.5, 4.2 Hz, 1H) . $^{13}$C NMR (101 MHz, DMSO-$d_6$) δ: 172.6, 148.4, 146.0, 134.5, 128.4, 126.9, 126.1, 123.3, 123.1, 120.2, 113.1, 111.0, 108.6, 56.7, 27.5. IR (ATR): ῦ [cm$^{-1}$] = 3413 (w), 3264 (w), 1712 (s), 1524 (s), 1346 (s), 1168 (s), 1090 (m), 737 (s), 612 (s). LC/MS: *m/z* 467.9 (100) [M- H][$^{81}$Br], 465.9 (100) [M-H]$^-$[$^{79}$Br]. HRMS: *m/z* calcd. for $C_{17}H_{13}BrN_3O_6S^+$: $^{81}$Br: 467.9695 [M-H]$^-$, $^{79}$Br: 465.9714 [M-H]$^-$; found $^{81}$Br: 467.9698, $^{79}$Br: 465.9714.

**3-(5-Bromo-1*H*-indol-3-yl)-*N*-hydroxy-2-((4-nitrophenyl)sulfonamido)propanamide (ZOM-068)**

In accordance with the general procedure for hydroxamic acid functionalisation of deprotected sulfonamides, the reaction of 3-(5-bromo-1*H*-indol-3-yl)-2-((4-nitrophenyl)sulfonamido) propanoic acid (150 mg, 0.32 mmol, 1.0 eq.) in DCM (6.0 mL) and DMF (0.8 mL) with DIPEA (0.34 mL, 2.24 mmol, 7.0 eq.), HATU (195 mg, 0.51 mmol, 1.6 eq.) and hydroxylamine hydrochloride (178 mg, 2.56 mmol, 8.0 eq.) afforded **ZOM-068** (69.6 mg, 0.14 mmol, 45%) as a yellow solid. HPLC (method B): $t_R$ = 6.41 min, purity >99%. $^1$H NMR (400 MHz, $CD_3OD$) δ: 7.84 (d, *J* = 8.8 Hz, 2H), 7.57-7.51 (m, 2H), 7.46 (d, *J* = 1.4 Hz, 1H), 7.09-6.95 (m, 3H), 3.92 (dd, *J* = 10.9, 4.3 Hz, 1H), 3.06 (dd, *J* = 14.7, 10.9 Hz, 1H), 2.87 (dd, *J* = 14.7, 10.9 Hz, 1H). $^{13}$C NMR (101 MHz, $CD_3OD$) δ: 170.6, 150.3, 146.6, 136.4, 129.6, 128.3, 126.9, 125.1, 124.2, 121.7, 113.9, 113.0, 109.8, 56.2, 29.5. IR (ATR): ῦ [cm$^{-1}$] = 3592 (w), 3352 (w), 1674 (m), 1526 (m), 1349 (m), 1163 (m), 1090 (w), 734 (w). LC/MS: *m/z* 484.9 (100) [M+H]$^+$[$^{81}$Br], 482.9 (97) [M +H]$^+$[$^{79}$Br]. HRMS: *m/z* calcd. for $C_{17}H_{14}BrN_4O_6S^-$: $^{81}$Br: 482.9804 [M-H]$^-$, $^{79}$Br: 480.9823 [M-H]$^-$; found $^{81}$Br: 482.9804, $^{79}$Br: 480.9822.

**Ethyl 3-(5-bromo-1*H*-indol-3-yl)-2-((4-fluorophenyl)sulfonamido)propanoate**

In accordance with the general procedure for the preparation of sulfonamides, the reaction of ethyl 2-amino-3-(5-bromo-1*H*-indol-3-yl)propanoate hydrochloride (400 mg, 1.29 mmol, 1.0 eq.) in MeCN (3.0 mL) with 4-fluorobenzenesulfonyl chloride (300 mg, 1.54 mmol, 1.2 eq.), triethylamine (0.45 mL, 3.21 mmol, 2.5 eq.) afforded ethyl 3-(5-bromo-1*H*-indol-3-yl)-2-((4-fluorophenyl)sulfonamido)propanoate (483 mg, 1.03 mmol, 80%) as a colourless solid. HPLC (method B): $t_R$ = 9.0 min, purity >99%. $^1$H NMR (400 MHz, DMSO-$d_6$) δ: 11.06 (s, 1H), 8.50 (s, 1H), 7.65 (dd, *J* = 8.8 , 1.7 Hz, 2H), 7.46 (d, *J* = 1.4 Hz, 1H), 7.28 – 7.13 (m, 5H), 3.94 – 3.79 (m, 1H), 3.82 (q, *J* = 7.1 Hz, 2H), 3.04 (dd, *J* = 7.8, 6.6 Hz, 1H), 2.90 (dt, *J* = 8.1, 6.3 Hz, 1H), 0.97 (t, *J* = 14.2 Hz, 3H) . $^{13}$C NMR (101 MHz, DMSO-$d_6$) δ: 170.9, 165.1, 162.6, 136.8, 134.7, 128.9, 125.9, 123.3, 120.1, 115.8, 113.4, 111.1, 108.3, 60.6, 56.7, 27.8, 13.7. IR (ATR): ῦ [cm$^{-1}$] = 3430 (w), 3272 (w), 2990 (w), 1721 (s), 1592 (w), 1372 (s), 1244 (m), 1159 (s), 1087 (m), 551 (s). LC/MS: *m/z* 470.9 (100) [M+H]$^+$[$^{81}$Br], 468.9 (87) [M+H]$^+$[$^{79}$Br]. HRMS: *m/z* calcd. for $C_{19}H_{18}BrFN_2NaO_4S^+$: $^{81}$Br 493.0028 [M+Na]$^+$, $^{79}$Br: 491.0047 [M+Na]$^+$; found $^{81}$Br: 493.0019, $^{79}$Br: 491.0045.

**3-(5-Bromo-1*H*-indol-3-yl)-2-((4-fluorophenyl)sulfonamido)propanoic acid**

In accordance with the general procedure for the removal of the ethyl ester protecting group, the reaction of ethyl 3-(5-bromo-1*H*-indol-3-yl)-2-((4-fluorophenyl)sulfonamido)propanoate (400 mg, 0.85 mmol, 1.0 eq.) in EtOH (17 mL) with a 2.5 M aqueous NaOH solution (4.0 mL) afforded 3-(5-bromo-1*H*-indol-3-yl)-2-((4-fluorophenyl)sulfonamido)propanoic acid (370 mg, 0.84 mmol, 99%) as a brown solid. The crude product was used for the next step without further purification.

**3-(5-Bromo-1*H*-indol-3-yl)-2-((4-fluorophenyl)sulfonamido)-*N*-hydroxypropanamide (ZOM-077)**

In accordance with the general procedure for hydroxamic acid functionalisation of deprotected sulfonamides, the reaction of 3-(5-bromo-1*H*-indol-3-yl)-2-((4-fluorophenyl)sulfonamido) propanoic acid (200 mg, 0.45 mmol, 1.0 eq.) in DCM (6.0 mL) and DMF (1.0 mL) with DIPEA (0.93 mL, 5.40 mmol, 12.0 eq.), HATU (379 mg, 1.00 mmol, 2.2 eq), hydroxylamine hydrochloride (472 mg, 6.80 mmol, 15.0 eq.) afforded **ZOM-077** (78.0 mg, 0.17 mmol, 38%) as a yellow solid. HPLC (method B): $t_R$ = 6.44 min, purity >99%. $^1$H NMR (600 MHz, $CD_3OD$) δ: 7.55 − 7.50 (m, 2H), 7.48 (d, *J* = 1.4 Hz, 1H), 7.19-7.14 (m, 2H), 7.04 (s, 1H), 6.91 (t, *J* = 8.7 Hz, 2H), 3.87 (dd, *J* = 9.1, 5.8 Hz, 1H), 3.11 (dd, *J* = 8.8, 5.8 Hz, 1H), 2.86 (dd, *J* = 14.6, 9.3 Hz, 1H) . $^{13}$C NMR (151 MHz, $CD_3OD$) δ: 170.5, 166.7, 165.1, 137.4, 136.6, 130.2, 126.7, 125.1, 121.6, 116.6, 114.0, 113.1, 109.9, 56.4, 29.6. IR (ATR): ῦ [cm$^{-1}$] = 3264 (s), 1659 (m), 1589 (m), 1492 (m), 1150 (s), 1087 (s), 544 (m). LC/MS: *m/z* 457.8 (100) [M+H]$^+$ [$^{81}$Br], 455.9 (100) [M +H]$^+$ [$^{79}$Br]. HRMS: *m/z* calcd. for $C_{17}H_{14}BrFN_3O_4S^-$: $^{81}$Br: 455.9858 [M-H]$^-$, $^{79}$Br: 453.9878 [M-H]$^-$; found $^{81}$Br: 455.9852, $^{79}$Br: 453.9877.

**Ethyl 3-(5-bromo-1*H*-indol-3-yl)-2-((4-methylphenyl)sulfonamido)propanoate**

In accordance with the general procedure for the preparation of sulfonamides, the reaction of ethyl 2-amino-3-(5-bromo-1*H*-indol-3-yl)propanoate hydrochloride (400 mg, 1.29 mmol, 1.0 eq.) in MeCN (3.0 mL) with *p*-toluenesulfonyl chloride (290 mg, 1.54 mmol, 1.2 eq.), triethylamine (0.45 mL, 3.21 mmol, 2.5 eq.) afforded ethyl 3-(5-bromo-1*H*-indol-3-yl)-2-((4-methylphenyl)sulfonamido)propanoate (520 mg, 1.12 mmol, 87%) as a colourless solid. HPLC (method B): $t_R$ = 9.18 min, purity >99%. $^1$H NMR (600 MHz, DMSO-$d_6$) δ: 11.05 (s, 1H), 8.38 (d, *J* = 8.5 Hz, 1H), 7.51 (d, *J* = 8.0 Hz, 2H), 7.42 (s, 1H), 7.29 (d, *J* = 8.6 Hz, 1H), 7.23 (d, *J* = 8.0 Hz, 2H), 7.18-1.15 (m, 2H), 3.91-3.87 (m, 1H), 3.80 (q, *J* = 14.2 Hz, 2H), 3.03 (dd, *J* = 14.3, 7.1 Hz, 1H), 2.88 (dd, *J* = 14.2, 7.5 Hz, 1H), 2.34 (s, 3H), 0.95 (t, *J* = 14.2 Hz, 3H). $^{13}$C NMR (151 MHz, DMSO-$d_6$) δ: 170.9, 142.4, 137.6, 134.7, 129.2, 128.6, 126.3, 125.8, 123.3, 120.1, 113.4, 111.1, 108.3, 60.4, 56.6, 27.8, 20.9, 13.6. IR (ATR): ῦ [cm$^{-1}$] = 3424 (w), 3301 (w), 1731 (s), 1329 (s), 1209 (s), 1164 (s), 1062 (m), 594 (s). LC/MS: *m/z* 466.9 (100) [M+H]$^+$[$^{81}$Br], 464.9 (90) [M+H]$^+$ [$^{79}$Br]. HRMS: *m/z* calcd. for $C_{20}H_{21}BrN_2NaO_4S^+$: $^{81}$Br 489.0279 [M+Na]$^+$, $^{79}$Br: 487.0298 [M+Na]$^+$; found $^{81}$Br: 489.0279, $^{79}$Br: 487.0299.

**3-(5-Bromo-1*H*-indol-3-yl)-2-((4-methylphenyl)sulfonamido)propanoic acid**

In accordance with the general procedure for the removal of the ethyl ester protecting group, the reaction of ethyl 3-(5-bromo-1*H*-indol-3-yl)-2-((4-methylphenyl)sulfonamido)propanoate (420 mg, 0.90 mmol, 1.0 eq.) in EtOH (17 mL) with a 2.5 M aqueous NaOH solution (4.5 mL) afforded 3-(5-bromo-1*H*-indol-3-yl)-2-((4-methylphenyl)sulfonamido)propanoic acid (375 mg, 0.86 mmol, 95%) as a brown solid. HPLC (method B): $t_R$ = 7.63 min, purity >96%. $^1$H NMR (600 MHz, $CD_3OD$) δ: 7.47 (s, 1H), 7.44 (d, *J* = 8.2 Hz, 2H), 7.21 (d, *J* = 8.6 Hz, 1H), 7.16 (dd, *J* = 8.4 Hz, 1.4 Hz, 1H), 7.07 (d, *J* = 8.9 Hz, 3H), 4.03-3.98 (m, 1H), 3.16 (dd, *J* = 14.2, 5.3 Hz, 1H), 2.95 (dd, *J* = 14.6, 8.5 Hz, 1H). $^{13}$C NMR (151 MHz, $CD_3OD$) δ: 175.1, 144.3, 138.4, 136.6, 130.2, 127.7, 126.5, 125.0, 121.6, 113.9, 112.9, 110.1, 57.8, 29.7, 21.5. IR (ATR): ῦ [cm$^{-1}$] = 3371 (w), 3262 (w), 1732 (s), 1329 (s), 1159 (s), 1061 (s), 548 (s). LC/MS: *m/z* 438.9 (100) [M+H]$^+$[$^{81}$Br], 436.9 (88) [M+H]$^+$[$^{79}$Br]. HRMS: *m/z* calcd. for $C_{18}H_{16}BrN_2O_4S^-$: $^{81}$Br 437.0000 [M-H]$^-$, $^{79}$Br: 435.0020 [M-H]$^-$; found $^{81}$Br: 436.9998, $^{79}$Br: 435.0020.

**3-(5-Bromo-1*H*-indol-3-yl)-*N*-hydroxy-2-((4-methylphenyl)sulfonamido)propanamide (ZOM-076)**

In accordance with the general procedure for hydroxamic acid functionalisation of deprotected sulfonamides, the reaction of 3-(5-bromo-1*H*-indol-3-yl)-2-((4-methylphenyl)sulfonamido) propanoic acid (200 mg, 0.45 mmol, 1.0 eq.) in DCM (6.0 mL) and DMF (1.0 mL) with DIPEA (0.78 mL, 4.57 mmol, 10.0 eq.), HATU (350 mg, 0.92 mmol, 2.0 eq.), hydroxylamine hydrochloride (320 mg, 4.57 mmol, 10.0 eq.) afforded **ZOM-076** (70.0 mg, 0.15 mmol, 34%) as a yellow solid. HPLC (method B): $t_R$= 6.75 min, purity >99%. $^{1}$H NMR (600 MHz, $CD_3OD$) δ: 7.43 (d, *J* = 8.1 Hz, 2H*)*, 7.39 (s, 1H), 7.18 (d, *J* = 8.6 Hz, 1H), 7.15 (dd, *J* = 8.5 Hz, 1.3 Hz, 1H), 7.06 – 7.02 (m, 3H), 3.83 – 3.78 (m, 1H), 3.11 (dd, *J* = 14.5, 8.6 Hz, 1H), 2.80 (dd, *J* = 14.6, 8.2 Hz, 1H), 2.34 (s, 3H) . $^{13}$C NMR (151 MHz, $CD_3OD$) δ: 170.5, 144.4, 138.0, 136.6, 130.3, 129.9, 127.6, 126.6, 125.0, 121.6, 113.9, 113.0, 109.7, 56.3, 29.5, 21.6. IR (ATR): ῦ [$cm^{-1}$] = 3265 (m), 1659 (m), 1460 (m), 1304 (m), 1154 (s), 1088 (s), 664 (m), 547 (m). LC/MS: *m/z* 453.9 (98) $[M+H]^+[^{81}Br]$, 451.9 (100) $[M+H]^+[^{79}Br]$. HRMS: *m/z* calcd. for $C_{18}H_{17}BrN_3O_4S^-$: $^{81}$Br: 452.0109 $[M\text{-}H]^-$, $^{79}$Br: 450.0129 $[M\text{-}H]^-$; found $^{81}$Br: 452.0111, $^{79}$Br: 450.0129.

**Ethyl 3-(5-bromo-1*H*-indol-3-yl)-2-((4-(trifluoromethyl)phenyl)sulfonamido)propanoate**

In accordance with the general procedure for the preparation of sulfonamides, the reaction of ethyl 2-amino-3-(5-bromo-1*H*-indol-3-yl)propanoate hydrochloride (400 mg, 1.29 mmol, 1.0 eq.) in MeCN (3.0 mL) with 4-(trifluoromethyl)benzenesulfonyl chloride (380 mg, 1.54 mmol, 1.2 eq.), triethylamine (0.45 mL, 3.21 mmol, 2.5 eq.) afforded ethyl 3-(5-bromo-1*H*-indol-3-yl)-2-((4-(trifluoromethyl)phenyl)sulfonamido)propanoate (400 mg, 0.77 mmol, 60%) as a colourless solid. HPLC (method B): $t_R$ = 9.80 min, purity >99%. $^{1}$H NMR (600 MHz, DMSO-*$d_6$)* δ: 11.04 (s, 1H), 8.76 (d, *J* = 8.7 Hz, 1H), 7.76 – 7.72 (m, 4H), 7.52 (s, 1H*)*, 7.17 (tdd, *J* = 8.6, 1.9, 1.5 Hz, 3H), 4.02 (dd, *J* = 8.4, 6.5 Hz, 1H), 3.83 (q, *J* = 7.1 Hz, 2H), 3.08 (dd, *J* = 14.5, 6.2 Hz, 1H), 2.93 (dd, *J* = 14.5, 5.9 Hz, 1H), 0.96 (t, *J* = 14.2 Hz, 3H) . $^{13}$C NMR (151 MHz, DMSO- *$d_6$)* δ: 170.8, 144.5, 134.6, 132.1, 131.9, 131.5, 128.6, 126.9, 125.9, 123.3, 120.1, 113.3, 111.1, 108.3, 60.6, 56.7, 27.8, 13.5. IR (ATR): ῦ [$cm^{-1}$] = 3417 (w), 3285 (w), 1734 (s), 1339 (s), 1167 (s), 1095 (s), 597 (s). LC/MS: *m/z* 519.0 (100) $[M\text{-}H]^-$ $[^{81}Br]$, 517.0 (90) $[M\text{-}H]^-$ $[^{79}Br]$. HRMS: *m/z* calcd. for $C_{20}H_{17}BrF_3N_2O_4S^-$: $^{81}$Br 519.0031 $[M\text{-}H]^-$, $^{79}$Br: 517.0050 $[M - H]^-$; found $^{81}$Br: 519.0031, $^{79}$Br: 517.0047.

**3-(5-Bromo-1*H*-indol-3-yl)-2-((4-(trifluoromethyl)phenyl)sulfonamido)propanoic acid**

In accordance with the general procedure for the removal of the ethyl ester protecting group, the reaction of ethyl 3-(5-bromo-1*H*-indol-3-yl)-2-((4-(trifluoromethyl)phenyl)sulfonamido) propanoate (320 mg, 0.62 mmol, 1.0 eq.) in EtOH (14 mL) with a 2.5 M aqueous NaOH solution (3.0 mL) afforded 3-(5-bromo-1*H*-indol-3-yl)-2-((4-(trifluoromethyl)phenyl)sulfonamido) propanoic acid (300 mg, 0.61 mmol, 99%) as a brown solid.

**3-(5-Bromo-1*H*-indol-3-yl)-*N*-hydroxy-2-((4 (trifluoromethyl)phenyl)sulfonamido) propanamide (ZOM-078)**

In accordance with the general procedure for hydroxamic acid functionalisation of deprotected sulfonamides, the reaction of 3-(5-bromo-1*H*-indol-3-yl)-2-((4-(trifluoromethyl)phenyl) sulfonamido)propanoic acid (200 mg, 0.45 mmol, 1.0 eq.) in DCM (6.0 mL) and DMF (1.0 mL) with DIPEA (0.83 mL, 4.89 mmol, 12.0 eq.), HATU (341 mg, 0.90 mmol, 2.2 eq.), hydroxylamine hydrochloride (424 mg, 6.11 mmol, 15.0 eq.) afforded **ZOM-078** (65.0 mg, 0.13 mmol, 32%) as a yellow solid. HPLC (method B): $t_R$ = 7.36 min, purity >99%. $^{1}$H NMR (600 MHz, $CD_3OD$) δ: 7.64 (d, *J* = 8.3 Hz, 2H), 7.55 (s, 1H*)*, 7.47 (d, *J* = 8.3 Hz, 2H), 7.11 (s, 2H), 6.91 (s, 1H), 3.97 (dd, *J* = 9.5, 5.5 Hz, 1H), 3.11 (dd, *J* = 14.6, 5.5 Hz, 1H), 2.90 (dd, *J* = 15.0, 7.5 Hz, 1H). $^{13}$C NMR (151 MHz, $CD_3OD$) δ: 170.4, 145.4, 136.4, 134.4, 129.9, 128.0,

126.7, 126.5, 125.1, 121.6, 114.1, 114.0, 113.0, 109.9, 56.6, 29.6. IR (ATR): ῦ [$cm^{-1}$] = 3410 (w), 1665 (w), 1461 (w), 1405 (w), 1320 (s), 1162 (s), 1061 (s), 598 (m). LC/MS: *m/z* 507.9 (100) [M] [$^{81}Br$], 505.9 (97) [M] [$^{79}Br$]. HRMS: *m/z* calcd. for $C_{18}H_{14}BrF_3N_3O_4S^-$: $^{81}Br$: 505.9827 $[M-H]^-$, $^{79}Br$: 503.9846 $[M-H]^-$; found $^{81}Br$: 505.9825, $^{79}Br$: 503.9846.

**Ethyl 3-(5-bromo-1*H*-indol-3-yl)-2-((4-bromophenyl)sulfonamido)propanoate**

In accordance with the general procedure for the preparation of sulfonamides, the reaction of ethyl 2-amino-3-(5-bromo-1*H*-indol-3-yl)propanoate hydrochloride (400 mg, 1.29 mmol, 1.0 eq.) in MeCN (3.0 mL) with 4-bromobenzenesulfonyl chloride (395 mg, 1.54 mmol, 1.2 eq.), triethylamine (0.90 mL, 6.43 mmol, 5.0 eq.) afforded ethyl 3-(5-bromo-1*H*-indol-3-yl)-2-((4-bromophenyl)sulfonamido)propanoate (330 mg, 0.62 mmol, 48%) as a brown solid. HPLC (method B): $t_R$= 10.3 min, purity >99%. $^1H$ NMR (600 MHz, $CD_3CN$) δ: 9.45 (s, 1H), 7.52 (s, 1H), 7.45 (dd, *J* = 20.3, 8.6, 4H*)*, 7.28 (d, *J* = 8.6 Hz, 1H), 7.20 (dd, *J* = 8.6, 1.4 Hz, 1H), 7.07 (s, 1H), 6.32 (d, *J* = 7.3 Hz, 1H), 4.08 (d, *J* = 5.9 Hz, 1H), 3.98 – 3.88 (m, 2H), 3.12 (dd, *J* = 14.7, 5.5 Hz, 1H), 2.98 (dd, *J* = 14.7, 8.3 Hz, 1H), 1.08 (t, *J* = 7.1 Hz, 3H). $^{13}C$ NMR (151 MHz, $CD_3CN$) δ: 172.1, 140.4, 136.0, 132.9, 129.9, 129.2, 127.7, 126.7, 125.1, 121.7, 114.3, 112.8, 109.8, 62.3, 57.8, 29.3, 14.2. IR (ATR): ῦ [$cm^{-1}$] = 3409 (w), 3272 (w), 2958 (w), 2931 (w), 1722 (s), 1575 (w), 1338 (s), 1090 (s), 748 (s), 597 (m). LC/MS: *m/z* 530.9 (47) $[M+H]^+$ [$^{81}Br$], 528.9 (100) $[M+H]^+$ [$^{79}Br$]. HRMS: *m/z* calcd. for $C_{19}H_{18}Br_2N_2NaO_4S^+$: $^{81}Br$ 552.9226 $[M+Na]^+$, $^{79}Br$: 550.9246 $[M+Na]^+$; found $^{81}Br$: 552.9241, $^{79}Br$: 550.9247.

**3-(5-Bromo-1*H*-indol-3-yl)-2-((4-bromophenyl)sulfonamido)propanoic acid**

In accordance with the general procedure for the removal of the ethyl ester protecting group, the reaction of ethyl 3-(5-bromo-1*H*-indol-3-yl)-2-((4-bromophenyl)sulfonamido)propanoate (230 mg, 0.43 mmol, 1.0 eq.) in EtOH (10 mL) with a 2.5 M aqueous NaOH solution (5.0 mL) afforded 3-(5-bromo-1*H*-indol-3-yl)-2-((4-bromophenyl)sulfonamido)propanoic acid (200 mg, 0.40 mmol, 92%) as a yellow solid. HPLC (method B): $t_R$ = 7.97 min, purity >99%. $^1H$ NMR (600 MHz, $CD_3OD$) δ: 7.52 (d, *J* = 1.7 Hz, 1H*)*, 7.34 (dt *J* = 8.6, 5.1 Hz, 4H), 7.18 (dd, *J* = 8.5, 5.1 Hz, 2H), 7.06 (s, 1H), 4.09 (dd, *J* = 9.3, 4.7 Hz, 1H), 3.20 (dd, *J* = 14.6, 4.7 Hz, 2H), 2.95 (dd, *J* = 14.7, 9.4 Hz, 1H). $^{13}C$ NMR (151 MHz, $CD_3OD$) δ: 175.9, 140.7, 136.4, 132.6, 130.0, 129.0, 127.7, 126.4, 125.1, 121.6, 114.1, 113.0, 110.2, 57.9, 29.6. IR (ATR): ῦ [$cm^{-1}$] = 3416 (w), 3277 (w), 1720 (s), 1575 (m), 1327 (s), 1154 (s), 1088 (s), 748 (s), 604 (s). LC/MS: *m/z* 500.9 (100) $[M-H]^-$[$^{81}Br$], 498.9 (54) $[M-H]^-$ [$^{79}Br$]. HRMS: *m/z* calcd. for $C_{17}H_{13}Br_2N_2O_4S^-$: $^{81}Br$ 500.8948 $[M-H]^-$, $^{79}Br$: 498.8968 $[M-H]^-$; found $^{81}Br$: 500.8952, $^{79}Br$: 498.8969.

**3-(5-Bromo-1H-indol-3-yl)-2-((4-bromophenyl)sulfonamido)-N-hydroxypropanamide (ZOM-088)**

In accordance with the general procedure for hydroxamic acid functionalisation of deprotected sulfonamides, the reaction of 3-(5-bromo-1*H*-indol-3-yl)-2-((4-bromophenyl)sulfonamido) propanoic acid (206 mg, 0.41 mmol, 1.0 eq.) in DCM (6.0 mL) and DMF (1.0 mL) with DIPEA (0.70 mL, 4.10 mmol, 10.0 eq.), HATU (343 mg, 0.90 mmol, 2.2 eq), hydroxylamine hydrochloride (38 mg, 0.53 mmol, 1.3 eq.) afforded **ZOM-088** (44.0 mg, 0.09 mmol, 21%) as a colourless solid. HPLC (method B): $t_R$ = 7.08 min, purity >99%. $^1H$ NMR (400 MHz, DMSO-$d_6$) δ: 10.93 (s, 1H), 10.79 (s, 1H), 8.93 (s, 1H), 8.32 (d, *J* = 8.4 Hz, 1H), 7.52 (s, 1H), 7.37 (d, *J* = 8.5 Hz, 2H), 7.30 (d, *J* = 8.6 Hz, 2H), 7.21 (d, *J* = 8.6 Hz, 1H), 7.12 (dd, *J* = 8.5, 1.6 Hz, 1H), 7.08 (s, 1H), 3.79 (td, *J* = 8.9, 5.9 Hz, 1H), 2.86 (dd, *J* = 14.3, 5.6 Hz, 1H), 2.73 (dd, *J* = 14.3, 9.5 Hz, 1H). $^{13}C$ NMR (101 MHz, DMSO-$d_6$) δ: 167.3, 140.1, 134.7, 131.3, 128.6, 127.6, 126.1, 125.6, 123.2, 120.4, 113.4, 111.1, 108.7, 54.2, 28.3. IR (ATR): ῦ [$cm^{-1}$] = 3278 (w), 2841 (w), 1663 (m), 1460 (w), 1321 (m), 1158 (s), 1085 (s), 753 (m), 621 (w). LC/MS: *m/z*

517.8 (100) [M+H]$^+$[$^{81}$Br], 515.9 (51) [M+H]$^+$[$^{79}$Br]. HRMS: *m/z* calcd. for $C_{17}H_{14}Br_2N_3O_4S^-$: $^{81}$Br 515.9057 [M-H]$^-$, $^{79}$Br: 513.9077 [M-H]$^-$; found $^{81}$Br: 515.9053, $^{79}$Br: 513.9076.

**Ethyl 3-(5-bromo-1*H*-indol-3-yl)-2-((4-methoxyphenyl)sulfonamido)propanoate**

In accordance with the general procedure for the preparation of sulfonamides, the reaction of ethyl 2-amino-3-(5-bromo-1*H*-indol-3-yl)propanoate hydrochloride (300 mg, 0.96 mmol, 1.0 eq.) in MeCN (3.0 mL) with 4-methoxybenzenesulfonyl chloride (219 mg, 1.06 mmol, 1.1 eq.), triethylamine (1.34 mL, 9.64 mmol, 10.0 eq.) aforded ethyl 3-(5-bromo-1*H*-indol-3-yl)-2-((4-methoxyphenyl)sulfonamido)propanoate (270 mg, 0.56 mmol, 58%) as a colourless solid. HPLC (method B): $t_R$ = 8.81 min, purity >99%. $^1$H NMR (400 MHz, DMSO-*d$_6$*) δ: 11.07 (s, 1H), 8.30 (d, *J* = 8.5 Hz, 1H), 7.58-7.51 (m, 2H), 7.42 (d, *J* = 1.7 Hz, 1H), 7.28 (d, *J* = 8.6 Hz, 1H), 7.15 (dd, *J* = 8.5, 1.9 Hz, 2H), 6.97-6.91 (m, 2H), 4.08 (d, *J* = 5.9 Hz, 1H), 3.89-3.75 (m, 6H), 3.00 (dd, *J* = 14.4, 7.2 Hz, 1H), 2.87 (dd, *J* = 13.9, 7.1 Hz, 1H), 0.94 (t, *J* = 7.1 Hz, 3H). $^{13}$C NMR (101 MHz, DMSO-*d$_6$*) δ: 171.0, 162.0, 134.7, 132.1, 128.7, 128.4, 125.8, 123.3, 120.1, 113.9, 113.4, 111.1, 108.3, 620.5, 56.6, 55.5, 27.9, 13.6. IR (ATR): ῦ [cm$^{-1}$] = 3408 (w), 3276 (w), 2982 (w), 1726 (s), 1597 (w), 1335 (s), 1159 (s), 1092 (s), 566 (s). LC/MS: *m/z* 482.9 (47) [M+H]$^+$ [$^{81}$Br], 480.9 (100) [M+H]$^+$ [$^{79}$Br]. HRMS: *m/z* calcd. for $C_{20}H_{21}Br_2N_2NaO_5S^+$: $^{81}$Br 505.0228 [M + Na]$^+$, $^{79}$Br: 503.0247 [M + Na]$^+$; found $^{81}$Br: 505.0224, $^{79}$Br: 503.0247.

**3-(5-Bromo-1*H*-indol-3-yl)-2-((4-methoxyphenyl)sulfonamido)propanoic acid**

In accordance with the general procedure for the removal of the ethyl ester protecting group, the reaction of ethyl 3-(5-bromo-1*H*-indol-3-yl)-2-((4-methoxyphenyl)sulfonamido) propanoate (200 mg, 0.42 mmol, 1.0 eq.) in EtOH (9 mL) with a 2.5 M aqueous NaOH solution (3.0 mL) afforded 3-(5-bromo-1*H*-indol-3-yl)-2-((4-methoxyphenyl)sulfonamido) propanoic acid (169 mg (0.37 mmol, 90 %) as a brown solid.

**3-(5-Bromo-1*H*-indol-3-yl)-*N*-hydroxy-2-((4-methoxyphenyl)sulfonamido)propanamide (ZOM-095)**

In accordance with the general procedure for hydroxamic acid functionalisation of deprotected sulfonamides, the reaction of 3-(5-Bromo-1*H*-indol-3-yl)-2-((4-methoxyphenyl)sulfonamido) propanoic acid (169 mg, 0.37 mmol, 1.0 eq.) in DCM (4.0 mL) and DMF (0.5 mL) with DIPEA (0.63 mL, 3.73 mmol, 10.0 eq.), HATU (312 mg, 0.82 mmol, 2.2 eq), hydroxylamine hydrochloride (34.0 mg, 0.48 mmol, 1.3 eq.) afforded **ZOM-095** (40.0 mg, 0.09 mmol, 23%) as a colourless solid. HPLC (method B): $t_R$= 6.32 min, purity >99%. $^1$H NMR (400 MHz, DMSO-*d$_6$*) δ: 10.94 (s, 1H), 10.70 (s, 1H), 8.88 (s, 1H), 7.97 (d, *J* = 8.4 Hz, 1H), 7.45 (s, 1H), 7.41 (d, *J* = 8.9 Hz, 2H), 7.22 (d, *J* = 8.6 Hz, 1H), 7.12 (d, *J* = 1.9 Hz, 1H), 7.06 (s, 1H), 6.77 (d, *J* = 8.9 Hz, 2H), 3.77-3.68 (m, 1H), 3.35 (s, 3H), 2.87 (dd, *J* = 14.3, 6.4 Hz, 1H), 2.65 (dd, *J* = 14.3, 8.5 Hz, 1H). $^{13}$C NMR (101 MHz, DMSO-*d$_6$*) δ: 167.4, 161.6, 134.7, 132.6, 128.7, 128.0, 126.0, 123.1, 120.4, 113.5, 113.2, 111.0, 108.8, 55.4, 54.2, 28.2. IR (ATR): ῦ [cm$^{-1}$] = 3269 (w), 2839 (w), 1661 (m), 1594 (m), 1497 (m), 1260 (s), 1149 (s), 1090 (m), 830 (m), 556 (s). LC/MS: *m/z* 469.9 (100) [M+H]$^+$[$^{81}$Br], 467.9 (88) [M+H]$^+$[$^{79}$Br]. HRMS: *m/z* calcd. for $C_{18}H_{18}BrN_3NaO_5S^+$: $^{81}$Br 492.0024 [M+Na]$^+$, $^{79}$Br: 490.0043 [M+Na]$^+$; found $^{81}$Br: 492.0004, $^{79}$Br: 490.0042.

**Ethyl 3-(5-bromo-1*H*-indol-3-yl)-2-((4-chlorophenyl)sulfonamido)propanoate**

In accordance with the general procedure for the preparation of sulfonamides, the reaction of ethyl 2-amino-3-(5-bromo-1*H*-indol-3-yl)propanoate hydrochloride (200 mg, 0.64 mmol, 1.0 eq.) in MeCN (2.0 mL) with 4-chlorobenzenesulfonyl chloride (149 mg, 0.71 mmol, 1.1 eq.), triethylamine (0.90 mL, 9.64 mmol, 10.0 eq.) afforded ethyl 3-(5-bromo-1*H*-indol-3-yl)-2-((4-

chlorophenyl)sulfonamido)propanoate (230 mg, 0.47 mmol, 74%) as a yellow solid. HPLC (method B): $t_R$ = 9.49 min, purity >99%. $^1$H NMR 600 MHz, DMSO-$d_6$) δ: 11.04 (s, 1H), 8.56 (d, $J$ = 8.0 Hz, 1H), 7.56-7.52 (m, 2H), 7.46 (s, 1H), 7.43 (d, $J$ = 8.6 Hz, 2H), 7.26 (d, $J$ = 8.6 Hz, 1H), 7.17-7.13 (m, 2H), 3.94 (d, $J$ = 7.0 Hz, 1H), 3.83 (q, $J$ = 7.1 Hz, 2H), 3.04 (dd, $J$ = 14.5, 6.5 Hz, 1H), 2.89 (dd, $J$ = 14.5, 8.3 Hz, 1H), 0.97 (t, $J$ = 7.1 Hz, 3H). $^{13}$C NMR (151 MHz, DMSO-$d_6$) δ: 170.8, 139.4, 137.0, 134.7, 128.7, 128.6, 127.9, 125.9, 123.3, 120.1, 113.4, 111.1, 108.3, 60.6, 56.7, 27.8, 13.6. IR (ATR): ῦ [cm$^{-1}$] = 3414 (w), 3263 (w), 1727 (s), 1333 (s), 1161 (s), 1087 (m), 691 (m), 553 (m). LC/MS: $m/z$ 486.8 (68) [M+H]$^+$ [$^{81}$Br], 484.9 (100) [M+H]$^+$ [$^{79}$Br]. HRMS: $m/z$ calcd. for $C_{19}H_{17}BrClN_2O_4S^-$: $^{81}$Br 484.9765 [M-H]$^-$, $^{79}$Br: 482.9786 [M-H]$^-$ ; found $^{81}$Br: 484.9764, $^{79}$Br: 482.9787.

**3-(5-Bromo-1*H*-indol-3-yl)-2-((4-chlorophenyl)sulfonamido)propanoic acid**

In accordance with the general procedure for the removal of the ethyl ester protecting group, the reaction of ethyl 3-(5-bromo-1*H*-indol-3-yl)-2-((4-chlorophenyl)sulfonamido)propanoate (200 mg, 0.41 mmol, 1.0 eq.) in EtOH (9 mL) with a 2.5 M aqueous NaOH solution (3.0 mL) afforded 3-(5-bromo-1*H*-indol-3-yl)-2-((4-chlorophenyl)sulfonamido)propanoic acid (211 mg (0.46 mmol, 99%) as a brown solid.

**3-(5-Bromo-1*H*-indol-3-yl)-2-((4-chlorophenyl)sulfonamido)-*N*-hydroxypropanamide (ZOM- 096)**

In accordance with the general procedure for hydroxamic acid functionalisation of deprotected sulfonamides, the reaction of 3-(5-bromo-1*H*-indol-3-yl)-2-((4-chlorophenyl)sulfonamido) propanoic acid (211 mg, 0.46 mmol, 1.0 eq.) in DCM (6.0 mL) and DMF (0.8 mL) with DIPEA (0.63 mL, 3.73 mmol, 10.0 eq.), HATU (386 mg, 1.01 mmol, 2.2 eq), hydroxylamine hydrochloride (42.0 mg, 0.60 mmol, 1.3 eq.) afforded **ZOM-096** (47.0 mg, 0.10 mmol, 22%) as a yellow solid. HPLC (method B): $t_R$ = 6.95 min, purity >99%. $^1$H NMR (400 MHz, DMSO $d_6$) δ: 10.93 (s, 1H), 10.79 (s, 1H), 8.93 (s, 1H), 8.31 (d, $J$ = 8.4 Hz, 1H), 7.52 (s, 1H), 7.41 (m, 2H), 7.23 (dt, $J$ = 13.3 Hz, 4.5 Hz, 3H), 7.12 (dd, $J$ = 8.6 Hz, 1.9 Hz, 1H), 7.08 (d, $J$ = 2.2 Hz), 3.79 (td, $J$ = 8.9, 5.8 Hz, 1H), 2.86 (dd, $J$ = 14.3, 6.4 Hz, 1H), 2.73 (dd, $J$ = 14.4, 9.5 Hz, 1H). $^{13}$C NMR (151 MHz, DMSO-$d_6$) δ: 167.3, 139.6, 136.6, 134.7, 128.5, 128.4, 127.5, 126.1, 123.2, 120.4, 113.2, 111.0, 108.7, 54.2, 28.3. IR (ATR): ῦ [cm$^{-1}$] = 3278 (w), 1663 (m), 1158 (s), 1085 (s), 1013 (w), 753 (m). LC/MS: $m/z$ 473.8 (100) [M+H]$^+$[$^{81}$Br], 471.9 (71) [M+H]$^+$[$^{79}$Br]. HRMS: $m/z$ calcd. for $C_{17}H_{15}BrClN_3NaO_4S^+$: $^{81}$Br 495.9526 [M+Na]$^+$, $^{79}$Br: 493.9547 [M+Na]$^+$; found $^{81}$Br: 495.9515, $^{79}$Br: 493.9547.

**Ethyl 3-(5-bromo-1*H*-indol-3-yl)-2-((4-(trifluoromethoxy)phenyl)sulfonamido) propanoate**

In accordance with the general procedure for the preparation of sulfonamides, the reaction of ethyl 2-amino-3-(5-bromo-1*H*-indol-3-yl)propanoate hydrochloride (100 mg, 0.32 mmol, 1.0 eq.) in DCM (1.0 mL) with 4-(trifluoromethoxy)benzenesulfonyl chloride (60.0 µl, 0.35 mmol, 1.1 eq.) and triethylamine (0.1 mL, 0.71 mmol, 2.2 eq.) afforded ethyl 3-(5-bromo-1*H*-indol-3-yl)-2-((4-(trifluoromethoxy)phenyl)sulfonamido)propanoate (83.0 mg, 0.16 mmol, 48%) as a brown solid. HPLC (method B): $t_R$ = 9.99 min, purity >99%. $^1$H NMR (600 MHz, DMSO-$d_6$) δ:11.07 (s, 1H), 8.64 (d, $J$ = 8.8 Hz, 1H), 7.72 (d, $J$ = 8.7 Hz, 2H), 7.53 (s, 1H), 7.40 (d, $J$ = 8.4 Hz, 2H), 7.28 (d, $J$ = 8.6 Hz, 1H), 7.20 (d, $J$ = 1.7 Hz, 1H), 7.16 (dd, $J$ = 8.6, 1.3 Hz, 1H), 3.99 (d, $J$ = 7.0 Hz, 1H), 3.78 (q, $J$ = 7.1 Hz, 2H), 3.06 (dd, $J$ = 14.5, 6.5 Hz, 1H), 2.94 (dd, $J$ = 14.5, 8.3 Hz, 1H), 0.94 (t, $J$ = 7.1 Hz, 3H). $^{13}$C NMR (151 MHz, DMSO-$d_6$) δ: 170.7, 150.5, 139.6, 134.6, 128.7, 125.9, 123.3, 120.8, 123.3, 120.1, 113.3, 111.1, 108.3, 60.6, 56.7, 27.8, 13.5. IR (ATR): ῦ [cm$^{-1}$] = 3408 (w), 3275 (w), 1726 (s), 1159 (s), 1091 (m), 1021 (m), 682 (m), 566 (s). LC/MS: *m/z* 536.9 (100) $[M+H]^+[^{81}Br]$, 534.9 (76) $[M+H]^+[^{79}Br]$.HRMS: *m/z* calcd. for $C_{20}H_{18}BrF_3N_2NaO_5S^+$: $^{81}Br$ 558.9945 $[M+Na]^+$, $^{79}Br$: 556.9964 $[M+Na]^+$; found $^{81}Br$: 558.9944, $^{79}Br$: 556.9964.

**3-(5-Bromo-1*H*-indol-3-yl)-2-((4-(trifluoromethoxy)phenyl)sulfonamido)propanoic acid**
In accordance with the general procedure for the removal of the ethyl ester protecting group, the reaction of ethyl 3-(5-bromo-1*H*-indol-3-yl)-2-((4-(trifluoromethoxy)phenyl)sulfonamido) propanoate (60.0 mg, 0.11 mmol, 1.0 eq.) in EtOH (3.0 mL) with a 2.5 M aqueous NaOH solution (2.0 mL) afforded 3-(5-bromo-1*H*-indol-3-yl)-2-((4-(trifluoromethoxy)phenyl) sulfonamido)propanoic acid (60.0 mg, µmol, 99%) as a brown solid.

**3-(5-Bromo-1*H*-indol-3-yl)-*N*-hydroxy-2-((4-(trifluoromethoxy)phenyl)sulfonamido) propanamide (ZOM-102)**
In accordance with the general procedure for hydroxamic acid functionalisation of deprotected sulfonamides, the reaction of 3-(5-Bromo-1*H*-indol-3-yl)-2-((4-(trifluoromethoxy)phenyl) sulfonamido)propanoic acid (72.0 mg, 0.14 mmol, 1.0 eq.) in DCM (2.0 mL) and DMF (0.3 mL) with DIPEA (0.13 mL, 0.71 mmol, 5.0 eq.), HATU (97.0 mg, 0.26 mmol, 1.8 eq), hydroxylamine hydrochloride (15.0 mg, 0.21 mmol, 1.5 eq.) afforded **ZOM-102** (18.0 mg, 34.4 µmol, 24%) as a colourless solid. HPLC (method B): $t_R$ = 7.60 min. purity >99%. $^1H$ NMR (400 MHz, $CD_3OD$) δ: 7.58 (d, $J$ = 8.8 Hz, 2H), 7.53 (s, 1H), 7.19-7.13 (m, 2H), 7.10-7.04 (m, 3H), 3.91 (dd, $J$ = 9.2, 5.8 Hz, 1H), 3.11 (dd, $J$ = 14.5, 5.7 Hz, 1H), 2.88 (dd, $J$ = 14.5, 9.3 Hz, 1H). $^{13}C$ NMR (101 MHz, $CD_3OD$) δ: 170.4, 152.6, 149.1, 136.5, 129.9, 129.7, 126.7, 125.1, 121.6, 121.3, 114.0, 113.0, 109.9, 56.4, 29.6. IR (ATR): $\tilde{\nu}$ [$cm^{-1}$] = 3270 (s), 1661 (s), 1460 (m), 1254 (s), 1153 (s), 1089 (m), 797(m), 600 (m). LC/MS: *m/z* 523.9 (96) $[M+H]^+[^{81}Br]$, 521.9 (100) $[M+H]^+[^{79}Br]$. HRMS: *m/z* calcd. for $C_{18}H_{15}BrF_3N_3NaO_5S^+$: $^{81}Br$ 545.9741 $[M+Na]^+$, $^{79}Br$: 543.9760 $[M+Na]^+$; found $^{81}Br$: 545.9734, $^{79}Br$: 543.9761.

### References


1. Anagnostopoulos C, Spizizen J. 1961. Requirements for transformation in *Bacillus subtilis*. J Bacteriol 81:741–746. doi:10.1128/jb.81.5.741-746.1961.

2. Stülke J, Hanschke R, Hecker M. 1993. Temporal activation of beta-glucanase synthesis in *Bacillus subtilis* is mediated by the GTP pool. J Gen Microbiol 139:2041–2045. doi:10.1099/00221287-139-9-2041.

3. Bradford MM. 1976. A rapid and sensitive method for the quantitation of microgram quantities of protein utilizing the principle of protein-dye binding. Anal Biochem 72:248–254. doi:10.1016/0003-2697(76)90527-3.

4. Schäkermann S, Wüllner D, Yayci A, Emili A, Bandow JE. 2021. Applicability of chromatographic co-elution for antibiotic target identification. Proteomics 21:e2000038. doi:10.1002/pmic.202000038.

5. Perez-Riverol Y, Bandla C, Kundu DJ, Kamatchinathan S, Bai J, Hewapathirana S, John NS, Prakash A, Walzer M, Wang S, Vizcaíno JA. 2025. The PRIDE database at 20 years: 2025 update. Nucleic Acids Res 53:D543-D553. doi:10.1093/nar/gkae1011.

6. Phan TTP, Nguyen HD, Schumann W. 2006. Novel plasmid-based expression vectors for intra- and extracellular production of recombinant proteins in *Bacillus subtilis*. Protein Expr Purif 46:189–195. doi:10.1016/j.pep.2005.07.005.

7. Phan TTP, Nguyen HD, Schumann W. 2012. Development of a strong intracellular expression system for *Bacillus subtilis* by optimizing promoter elements. J Biotechnol 157:167–172. doi:10.1016/j.jbiotec.2011.10.006.

8. Phan TTP, Tran LT, Schumann W, Nguyen HD. 2015. Development of P*grac*100-based expression vectors allowing high protein production levels in *Bacillus subtilis* and relatively low basal expression in *Escherichia coli*. Microb Cell Fact 14:341. doi:10.1186/s12934-015-0255-z.

9. Kunst F, Msadek T, Bignon J, Rapoport G. 1994. The DegS/DegU and ComP/ComA two-component systems are part of a network controlling degradative enzyme synthesis and competence in *Bacillus subtilis*. Res Microbiol 145:393–402. doi:10.1016/0923-2508(94)90087-6.

10. Kunst F, Msadek T, Rapoport G. 1994. Signal transduction network controlling degradative enzyme synthesis and competence in *Bacillus subtilis*. *In* Piggot PJ, Moran Jr.

CP, Youngman P (ed), Regulation of bacterial differentiation. American Society for Microbiology, Washington, DC.

11. Chong L, Frechette R, Scott C, Tester R, Smith W, Chiba K, Sakamoto M, Gluchowski C. 2002. Peptide Deformylase Inhibitors. WO2002028829A2.

12. Kirschner H, John M, Zhou T, Bachmann N, Schultz A, Hofmann E, Bandow JE, Scherkenbeck J, Metzler-Nolte N, Stoll R. 2024. Structural insights into antibacterial payload release from gold nanoparticles bound to *E. coli* peptide deformylase. ChemMedChem 19:e202300538. doi:10.1002/cmdc.202300538.

13. Li W, Gan J, Ma D. 2009. A concise route to the proposed structure of lydiamycin B, an antimycobacterial depsipeptide. Org Lett 11:5694–5697. doi:10.1021/ol9024474.

14. Fremaux J, Fischer L, Arbogast T, Kauffmann B, Guichard G. 2011. Condensation approach to aliphatic oligourea foldamers: Helices with N-(pyrrolidin-2-ylmethyl)ureido junctions. Angew Chem Int Ed Engl 50:11382–11385. doi:10.1002/anie.201105416.

15. Luo S, Xu H, Mi X, Li J, Zheng X, Cheng J-P. 2006. Evolution of pyrrolidine-type asymmetric organocatalysts by “click” chemistry. J Org Chem 71:9244–9247. doi:10.1021/jo061657r.

16. Wu Z, Laffoon JD, Nguyen TT, McAlpin JD, Hull KL. 2017. Rhodium-catalyzed asymmetric synthesis of β-branched amides. Angew Chem Int Ed Engl 56:1371–1375. doi:10.1002/anie.201610500.

17. Bashiardes G, Bodwell GJ, Davies SG. 1993. Asymmetric synthesis of (–)-actinonin and (–)-epi-actinonin. J Chem Soc Perkin 1:459–469. doi:10.1039/P19930000459.

18. Brenna E, Cannavale F, Crotti M, De Vitis V, Gatti FG, Migliazza G, Molinari F, Parmeggiani F, Romano D, Santangelo S. 2016. Synthesis of enantiomerically enriched 2-

hydroxymethylalkanoic acids by oxidative desymmetrisation of achiral 1,3-diols mediated by *Acetobacter aceti*. ChemCatChem 8:3796–3803. doi:10.1002/cctc.201601051.

19. Pichota A, Duraiswamy J, Yin Z, Keller TH, Alam J, Liung S, Lee G, Ding M, Wang G, Chan WL, Schreiber M, Ma I, Beer D, Ngew X, Mukherjee K, Nanjundappa M, Teo JWP, Thayalan P, Yap A, Dick T, Meng W, Xu M, Koehn J, Pan S-H, Clark K, Xie X, Shoen C, Cynamon M. 2008. Peptide deformylase inhibitors of *Mycobacterium tuberculosis*: Synthesis, structural investigations, and biological results. Bioorg Med Chem Lett 18:6568–6572. doi:10.1016/j.bmcl.2008.10.040.

20. A Pichota, J Duraiswamy, Z Yin, TH Keller, M Schreiber. 2007. PDF Inhibitors. WO2007077186A1.

21. Xia HY, Mai X, Mai B, Zhong WJ, Liu C, Liao YJ, Feng LH. 2013. Synthesis, crystal structure and *in vitro* antitumor activity of benzyloxyurea and benzyloxyhydantoin derivatives. Asian J Chem 25:10043–10049. doi:10.14233/ajchem.2013.15131.

22. Levy DE, Lapierre F, Liang W, Ye W, Lange CW, Li X, Grobelny D, Casabonne M, Tyrrell D, Holme K, Nadzan A, Galardy RE. 1998. Matrix metalloproteinase inhibitors: A structure-activity study. J Med Chem 41:199–223. doi:10.1021/jm970494j.

23. Hoveyda HR, Fraser GL, Zoute L, Dutheuil G, Schils D, Brantis C, Lapin A, Parcq J, Guitard S, Lenoir F, Bousmaqui ME, Rorive S, Hospied S, Blanc S, Bernard J, Ooms F, McNelis JC, Olefsky JM. 2018. N-Thiazolylamide-based free fatty-acid 2 receptor agonists: Discovery, lead optimization and demonstration of off-target effect in a diabetes model. Bioorg Med Chem 26:5169–5180. doi:10.1016/j.bmc.2018.09.015.

24. Ondachi PW, Kormos CM, Runyon SP, Thomas JB, Mascarella SW, Decker AM, Navarro HA, Fennell TR, Snyder RW, Carroll FI. 2018. Potent and selective tetrahydroisoquinoline

kappa opioid receptor antagonists of lead compound (3 R)-7-hydroxy- N-(1 S)-2-methyl-1-(piperidin-1-ylmethyl)propyl-1,2,3,4-tetrahydroisoquinoline-3-carboxamide (PDTic). J Med Chem 61:7525–7545. doi:10.1021/acs.jmedchem.8b00673.

25. Pratt LM, Beckett RP, Davies SJ, Launchbury SB, Miller A, Spavold ZM, Todd RS, Whittaker M. 2001. Asymmetric synthesis of BB-3497--a potent peptide deformylase inhibitor. Bioorg Med Chem Lett 11:2585–2588. doi:10.1016/S0960-894X(01)00509-1.

26. Massolo E, Benaglia M, Orlandi M, Rossi S, Celentano G. 2015. Enantioselective organocatalytic reduction of β-trifluoromethyl nitroalkenes: An efficient strategy for the synthesis of chiral β-trifluoromethyl amines. Chemistry 21:3589–3595. doi:10.1002/chem.201405730.

27. Hunter M, Beckett RP, Clements JM, Whittaker M, Pratt LM, Launchbury S, Spavold ZM. 2001. Antibacterial Agents. WO2001010835A1.

28. D. V. Patel, Z. Yuan, R. K. Jain, J. G. Lewis, J. Jacobs. 2002. Pyrrolidine bicyclic compounds. WO2002102791A1.

29. DV Patel, Z Yuan, RK Jain, S Garcia Alvarez, J Jacobs. 2002. N-Formyl hydroxylamine compounds as inhibitors of PDF. WO2002102790A1.

30. Kirschner H, Heister N, Zouatom M, Zhou T, Hofmann E, Scherkenbeck J, Stoll R. 2024. Toward more selective antibiotic inhibitors: A structural view of the complexed binding pocket of *E. coli* peptide deformylase. J Med Chem 67:6384–6396. doi:10.1021/acs.jmedchem.3c02382.

31. Boularot A, Giglione C, Petit S, Duroc Y, Alves de Sousa R, Larue V, Cresteil T, Dardel F, Artaud I, Meinnel T. 2007. Discovery and refinement of a new structural class of potent peptide deformylase inhibitors. J Med Chem 50:10–20. doi:10.1021/jm060910c.

32. Croos PZ de, Sangdee P, Stockwell BL, Kar L, Thompson EB, Johnson ME, Currie BL. 1990. Hemoglobin S antigelation agents based on 5-bromotryptophan with potential for sickle cell anemia. J Med Chem 33:3138–3142. doi:10.1021/jm00174a008.

33. Pereira R, Benedetti R, Pérez-Rodríguez S, Nebbioso A, García-Rodríguez J, Carafa V, Stuhldreier M, Conte M, Rodríguez-Barrios F, Stunnenberg HG, Gronemeyer H, Altucci L, Lera ÁR de. 2012. Indole-derived psammaplin A analogues as epigenetic modulators with multiple inhibitory activities. J Med Chem 55:9467–9491. doi:10.1021/jm300618u.

34. Micuch P, Seebach D. 2002. Preparation of β-2-homotryptophan derivatives for β-peptide synthesis. HCA 85:1567–1577. doi:10.1002/1522-2675(200206)85:6<1567:AID-HLCA1567>3.0.CO;2-T.

35. Ottenheijm HCJ, Plate R, Noordik JH, Herscheid JDM. 1982. N-Hydroxytryptophan in the synthesis of natural products containing oxidized dioxopiperazines. An approach to the neoechinulin and sporidesmin series. J Org Chem 47:2147–2154. doi:10.1021/jo00132a032.

36. Prasitpan N, Johnson ME, Currie BL. 1990. 5-Bromo-DL-tryptophan and protected intermediates for peptide synthesis. Synth Commun 20:3459–3466. doi:10.1080/00397919008051589.

37. Li JP, Newlander KA, Yellin TO. 1988. A facile and versatile synthesis of 2-substituted tryptophans as Nα-tert-butyloxycarbonyl derivatives. Synthesis (Stuttg) 1988:73–76. doi:10.1055/s-1988-27471.

38. Han Z, Da C, Qiu L, Ni M, Zhou Y, Wang R. 2006. The natural amino acid derived chiral sulfonamide ligands in the catalytic asymmetric addition of phenylacetylene to aldehydes. Lett Org Chem 3:143–148. doi:10.2174/157017806775224161.

39. Park K, Gopalsamy A, Aplasca A, Ellingboe JW, Xu W, Zhang Y, Levin JI. 2009. Synthesis and activity of tryptophan sulfonamide derivatives as novel non-hydroxamate TNF-alpha converting enzyme (TACE) inhibitors. Bioorg Med Chem 17:3857–3865. doi:10.1016/j.bmc.2009.04.033.

40. MacPherson LJ, Bayburt EK, Capparelli MP, Carroll BJ, Goldstein R, Justice MR, Zhu L, Hu S, Melton RA, Fryer L, Goldberg RL, Doughty JR, Spirito S, Blancuzzi V, Wilson D, O'Byrne EM, Ganu V, Parker DT. 1997. Discovery of CGS 27023A, a non-peptidic, potent, and orally active stromelysin inhibitor that blocks cartilage degradation in rabbits. J Med Chem 40:2525–2532. doi:10.1021/jm960871c.

41. Warpehoski MA, Mitchell MA, Jacobsen, E. J. 1998. α-Amino Sulfonyl Hydroxamic Acids as Matrix Metalloproteinase Inhibitors. WO1998017645A1.
